# Incomplete remyelination via endogenous or therapeutically enhanced oligodendrogenesis is sufficient to recover visual cortical function

**DOI:** 10.1101/2024.02.21.581491

**Authors:** Gustavo Della-Flora Nunes, Lindsay A Osso, Johana A Haynes, Amanda Morris, Lauren Conant, Michael E Stockton, Michael A Thornton, Jeffrey A Vivian, Rohan Gandhi, Daniel J Denman, Ethan G Hughes

**Author notes:** These authors contributed equally (Gustavo Della-Flora Nunes, Lindsay A Osso). These authors contributed equally (Daniel J Denman, Ethan G Hughes).

## Abstract

Myelin loss induces deficits in action potential propagation that result in neural dysfunction and contribute to the pathophysiology of neurodegenerative diseases, injury conditions, and aging. Because remyelination is often incomplete, better understanding endogenous remyelination and developing remyelination therapies that seek to restore neural function are clinical imperatives. Here, we used *in vivo* two-photon microscopy and electrophysiology to study the dynamics of endogenous and therapeutic-induced cortical remyelination and functional recovery after cuprizone-mediated demyelination in mice. We focused on the visual pathway, which is uniquely positioned to provide insights into structure-function relationships during de/remyelination. We show that endogenous remyelination is driven by recent oligodendrocyte loss and is highly efficacious following mild demyelination, but fails to restore the oligodendrocyte population when high rates of oligodendrocyte loss occur too quickly. Testing a novel thyromimetic compared to clemastine fumarate, we find it better enhances oligodendrocyte gain during remyelination and hastens recovery of neuronal function. Surprisingly, its therapeutic benefit was temporally restricted, and it acted exclusively following moderate to severe demyelination to eliminate endogenous remyelination deficits. However, complete remyelination is unnecessary as partial oligodendrocyte restoration was sufficient to recover visual neuronal function. These findings advance our understanding of remyelination and its impact on functional recovery to inform future therapeutic strategies.

---

Myelin, made by oligodendrocytes enwrapping axons with lipid-rich membranes, is essential for proper central nervous system (CNS) function. Loss of oligodendrocytes and myelin – known as demyelination – induces severe delay and failure of action potential propagation^1,2^, leaves neurons and their axons vulnerable to degeneration^3–5^, and causes motor, sensory, and cognitive impairment^6,7^. Demyelination occurs in white and gray matter in several pathologies, including inflammatory demyelinating diseases like multiple sclerosis^8^, traumatic CNS injury^9,10^, stroke^11^, Alzheimer’s disease^12^, and aging^13,14^. In particular, neocortical gray matter demyelination is highly correlated with physical and cognitive disability in multiple sclerosis^15^, emphasizing the clinical importance of understanding the role of neocortical myelin in neuronal function^16^.

Demyelination is typically followed by a period of heightened new myelin formation known as remyelination, which can restore action potential propagation and prevent neurodegeneration^2,4,5^. Remyelination is carried out by newly formed oligodendrocytes differentiating from oligodendrocyte precursor cells (OPCs)^17,18^ as well as – in some instances – by oligodendrocytes that survive the demyelinating injury^19–22^. However, the endogenous remyelination response is often insufficient, resulting in chronic demyelination^23,24^ and limited functional recovery^25,26^. Thus, understanding the drivers and limitations of endogenous remyelination and developing methods to enhance it are clinical imperatives for many demyelinating conditions. Despite substantial progress in identifying compounds that improve remyelination in recent years, there is still no FDA-approved remyelination therapy. Furthermore, independent of specific therapeutic strategies, we require a deeper understanding of fundamental aspects of therapeutic-induced remyelination, such as the dynamics and constraints of therapeutic action and the magnitude of remyelination required to recover neuronal function.

The afferent visual pathway is well-suited to investigate the relationship between neocortical myelin and neuronal function throughout de/remyelination^27^. The circuits of primary visual cortex (V1) are sensitive to input spike precision^28^ and contain precise and reliable sensory-evoked activity^29–31^, important for action potential transmission and visual coding^32,33^. Moreover, perturbations in the timing of sensory-evoked activity in V1 have previously been observed in patients and animal models during de/remyelination^25,34–36^. Here, we used longitudinal in vivo two-photon imaging of oligodendrocytes and high-density electrical recordings with single neuron resolution in V1 to study the dynamics of endogenous and therapeutic-induced neocortical remyelination and the relationship between remyelination and functional recovery.

Demyelination was induced with cuprizone, and mice were treated with two remyelination drugs: a new thyroid hormone mimetic, LL-341070, and a clinically validated therapeutic, clemastine fumarate.

Treatment with cuprizone induced oligodendrocyte loss and a concomitant increase in visual response latency. This was followed by a rapid and robust endogenous remyelination response that was driven by recent oligodendrocyte loss. Endogenous remyelination was highly efficacious at mild demyelination levels, but when moderate or severe demyelination occurred too quickly, endogenous remyelination failed to restore the oligodendrocyte population. LL-341070 treatment substantially increased oligodendrocyte gain during remyelination, acting more quickly and robustly than clemastine fumarate, and hastened neuron functional recovery. Surprisingly, its therapeutic benefit was temporally restricted as well as loss-dependent, exclusively impacting remyelination after moderate or severe demyelination to eliminate the endogenous remyelination deficit. However, complete remyelination was unnecessary as partial restoration of the oligodendrocyte population was sufficient to recover neuronal function.

## Results

### Severity of demyelination determines the extent of oligodendrocyte replacement

To induce demyelination, 9-10-week-old mice were fed chow containing 0.2% cuprizone. Cuprizone is a copper chelator widely used to cause specific oligodendrocyte cell death and demyelination, although its mechanism of action is incompletely understood^37,38^. After a subacute exposure (3.5 weeks versus commonly used exposure of 6 weeks^38^), mice were returned to their normal diet for seven additional weeks to allow for remyelination (**Fig. 1A**). Using histology, we did not detect demyelination in the optic nerve, indicating preservation of myelin in the anterior visual pathway with this cuprizone exposure protocol (**Fig. 1B, C**). To examine cortical demyelination, we used longitudinal *in vivo* two-photon imaging of V1 in *Mobp-EGFP* mice (**Fig. 1D**). These mice express *EGFP* under the promoter/enhancer for myelin-associated oligodendrocyte basic protein (*Mobp*), specifically labeling all oligodendrocytes and their associated myelin sheaths with EGFP (**Fig. 1E**)^39^. *In vivo* imaging throughout this period enabled the longitudinal tracking of oligodendrocytes in individual mice over time (**Fig. 1A, D-F**).

**Fig. 1.**
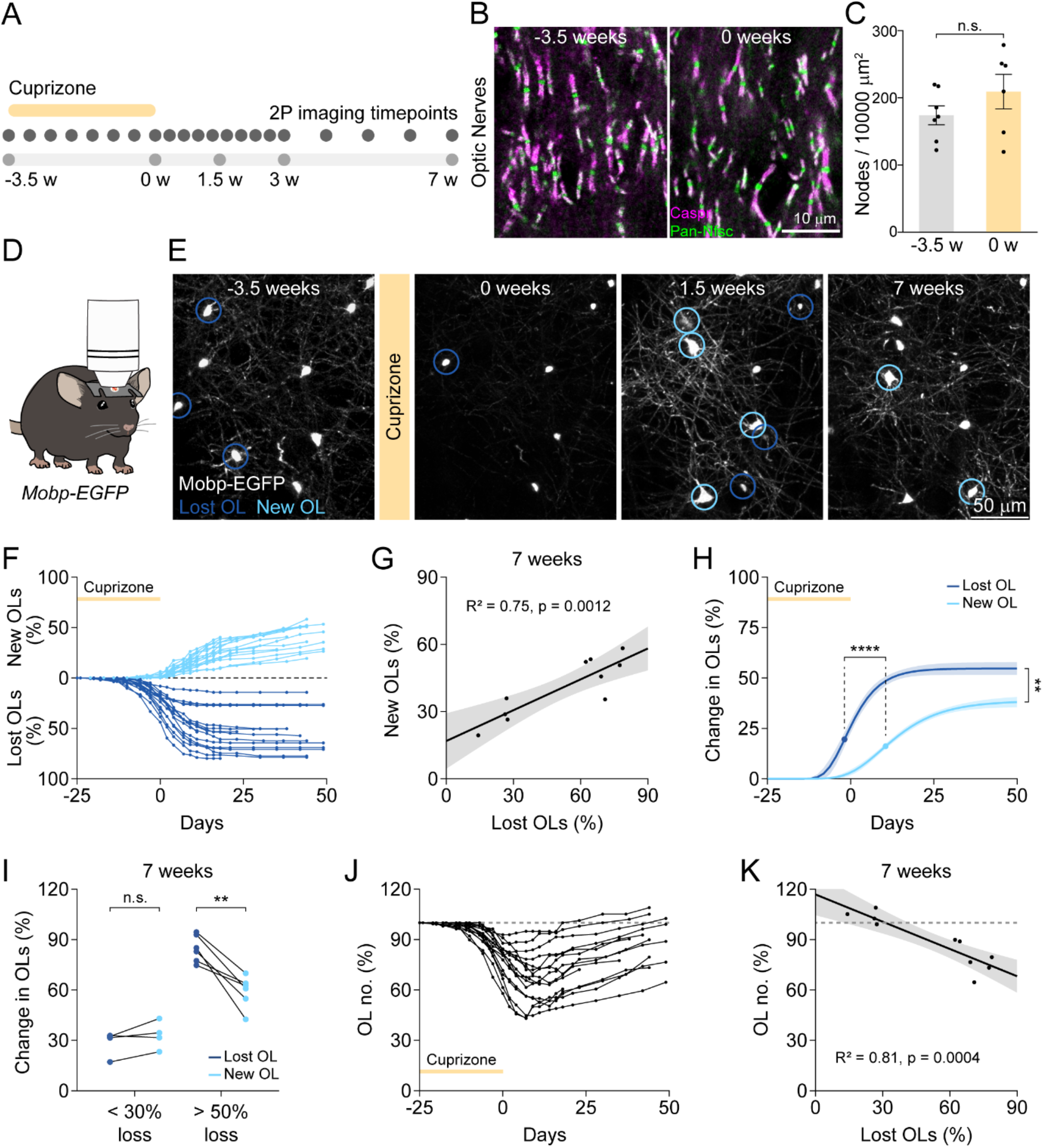
Endogenous remyelination fails to restore the oligodendrocyte population after moderate or severe demyelination. **(A)** Experimental timeline of cuprizone-mediated demyelination and longitudinal *in vivo* two-photon imaging. **(B)** Representative figures illustrating nodes of Ranvier and paranodes stained with Contactin-associated protein (Caspr, in magenta, localizing to paranodes) and an antibody labeling all neurofascin isoforms (Pan-Nfsc, in green, localizing to nodes of Ranvier and paranodes). **(C)** Cuprizone feeding does not alter the density of nodes of Ranvier in the optic nerve (bilaterally expressing Caspr and Pan-Nfsc). **(D)** *Mobp-EGFP* mice underwent craniotomy above primary visual cortex (V1) to allow for longitudinal *in vivo* two-photon imaging of oligodendrocytes (OLs). **(E)** Representative images of V1 OLs in one mouse at baseline (−3.5 weeks), at the end of cuprizone (0 weeks), and at two points during remyelination (1.5 and 7 weeks). Lost OLs (dark blue) and new OLs (light blue) are encircled. **(F)** Cumulative OL loss and gain as a percentage of baseline OLs in individual mice over time (n = 16). **(G)** Magnitude of OL gain by 7 weeks is tightly correlated with magnitude of OL loss by 7 weeks (n = 10, linear regression with 95% CI). **(H)** Three parameter Gompertz growth curve (95% CI) fit to group cumulative OL loss and gain shown for visualization. Statistical comparisons were performed on parameters derived from three parameter growth curves fit to individual mice (Supplementary Fig. 1, A to B). Only mice with imaging data out to at least 3 weeks post-cuprizone were used in growth curve modeling. Time of maximum growth rate occurred earlier for loss than gain. Gain asymptote is lower than loss asymptote. **(I)** At < 30% OL loss, mice make as many new OLs as they lose by 7 weeks. At > 50% OL loss, mice make fewer new OLs than they lose. **(J)** OL number as a percentage of baseline OLs in individual mice over time (n = 16). Dashed line at 100%. **(K)** OL number at 7 weeks is tightly correlated with magnitude of OL loss by 7 weeks (n = 10, linear regression with 95% CI). Dashed line at 100%. In **C**, Mean (−3.5 weeks: 173.4±14.08, n = 7; 0 weeks: 208.6±25.66, n = 6; p = 0.23, two-tailed unpaired t-test). In **H**, Max. growth rate (lost OL: -1.86±0.60, new OL: 10.52± 0.91, n = 15, t(14) = 10.12, **** p < 0.0001, paired t test). Asymptote (loss: 55.04±5.62, gain: 40.69±2.69, n = 15, t(14) = 3.64, ** p = 0.0027, paired t test). In **I**, Change in OLs at < 30% loss (lost OL: 23.71±3.15, new OL: 27.63±3.41, n = 4, t(3) = 1.874, p = 0.16, paired t test) and at > 50% loss (lost OL: 70.5±2.72, new OL: 49.27±3.23, n = 6, t(5) = 5.4, ** p = 0.0029, paired t test). n.s. not significant, ** p < 0.01, **** p < 0.0001, mean±SEM.

Cuprizone treatment resulted in overlapping periods of oligodendrocyte loss and gain (**Fig. 1E, F**) with sigmoidal dynamics that were well modeled by three parameter Gompertz growth curves (**Fig. 1F and Supplementary Fig. 1A, B**), allowing us to derive loss and gain asymptotes and timing of maximum rates (**Fig. 1H**).

Oligodendrocyte loss began during the cuprizone period and plateaued approximately three weeks following cuprizone removal (**Fig. 1F, H, and Supplementary Fig. 1A, D**), with a maximum loss rate 2±2 (SD) days prior to cuprizone removal (**Fig. 1H**). Oligodendrocyte gain commenced at the end of cuprizone (**Fig. 1F, H, and Supplementary Fig. 1B, E**) and its rate peaked 11±4 (SD) days after cuprizone removal, significantly later than loss (**Fig. 1H**). Between individual mice, we observed a large range in the extent of oligodendrocyte loss in the visual cortex (14% to 80% at 3 weeks) (**Fig. 1F and Supplementary 1A**), which was tightly positively correlated with the magnitude and rate of oligodendrocyte gain throughout remyelination (**Fig. 1G and Supplementary Fig. 2A, E**). However, the relationship between the magnitude of oligodendrocyte loss and gain was not one-to-one (**Fig. 1G**). Oligodendrocyte gain was insufficient to replace all lost oligodendrocytes by seven weeks of remyelination (**Fig. 1H**), consistent with findings in other cortical regions following cuprizone^19,40^. Interestingly, this deficit in endogenous cortical remyelination depended on the magnitude of oligodendrocyte loss incurred. Mice that lost fewer than 30% of oligodendrocytes generated as many new oligodendrocytes by seven weeks as they had lost, while mice that lost greater than 50% of oligodendrocytes failed to gain sufficient numbers of new oligodendrocytes to replace them (**Fig. 1I**). Thus, by seven weeks, there was a strong negative correlation between oligodendrocyte loss magnitude and oligodendrocyte number, with only mice with low levels of oligodendrocyte loss regenerating their original oligodendrocyte population while mice with high levels of oligodendrocyte loss exhibited a regeneration deficit (**Fig. 1J, K**). It is likely that this regeneration gap between mice incurring low and high levels of oligodendrocyte loss will persist since oligodendrocyte gain rate was no longer correlated with oligodendrocyte loss magnitude by seven weeks of remyelination (**Supplementary Fig. 2F**).

### Oligodendrocyte gain during remyelination is driven by recent oligodendrocyte loss

Elevated oligodendrocyte gain rates in response to oligodendrocyte loss is widely observed in animal models of demyelination (**Supplementary Fig. 1E**)^13,19,22,40–42^. Yet, we have a limited understanding of the factors initiating this endogenous response. Elevated gain rates correlate with loss levels (**Supplementary Fig. 2, C-E**) and subside shortly following the end of loss (**Supplementary Fig. 1E**), indicating both a scale and temporal dependence of oligodendrocyte gain on loss. However, it is unknown whether oligodendrocyte gain is induced in response to (a) a reduction in oligodendrocyte number initiating a homeostatic-like drive to restore the population, or (b) acute signaling around the loss of oligodendrocytes. Exploiting the lengthy, overlapping periods of oligodendrocyte loss and gain with varying levels between mice (**Fig. 1F**), we tested which parameters best predicted oligodendrocyte gain rate during remyelination. We modeled our longitudinal data to assess parameters continuously (**Supplementary Fig. 1A-E**).

If restoration of the oligodendrocyte population drives oligodendrocyte gain rate, we would expect gain rate to be higher the further oligodendrocyte numbers are from pre-cuprizone levels. To test this, we assessed the correlation between oligodendrocyte number and gain rate (**Fig. 2A, B**). We found that oligodendrocyte gain rate at 14 days post-cuprizone was weakly predicted by oligodendrocyte number at 7 days post-cuprizone (**Fig. 2A**). In assessing oligodendrocyte gain rate at 7, 14, 21, and 28 days post-cuprizone, we found that there was no period at which oligodendrocyte number predicted oligodendrocyte gain rate (**Fig. 2B**), indicating that a drive to reestablish oligodendrocyte numbers does not underlie oligodendrocyte gain during remyelination.

**Fig. 2.**
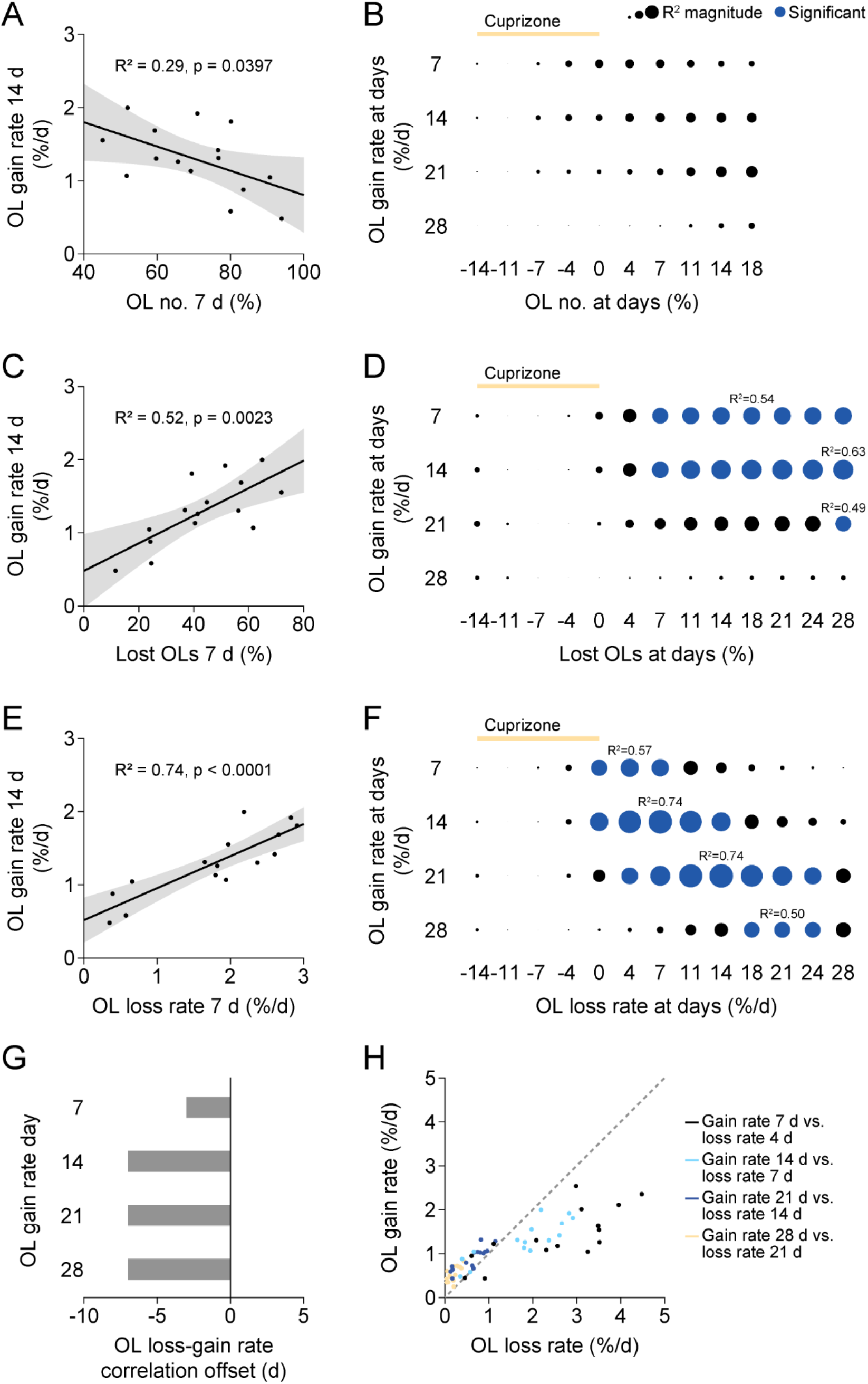
Endogenous remyelination is driven by recent oligodendrocyte loss. **(A** and **C** and **E)** OL gain rate (as a percentage of baseline OLs per day) at 14 days is weakly correlated with OL number (as a percentage of baseline OLs) at 7 days (A), correlated with cumulative OL loss (as a percentage of baseline OLs) at 7 days (C), and strongly correlated with OL loss rate (as a percentage of baseline OLs per day) at 7 days (E) (n = 15, linear regression with 95% CI). **(B** and **D** and **F)** OL gain rate at 7, 14, 21, or 28 days does not correlate well with OL number at -14, -11, -7, 0, 4, 7, 11, 14, or 18 days (B), correlates well with cumulative OL loss at several days during remyelination (D), and correlates best with OL loss rate approximately 7 days prior (F) (n = 15, linear regression with 95% CI). Dot size represents R^2^ magnitude. R^2^ value is indicated for the strongest significant correlation. Dark blue dots signify significant correlations given Bonferroni correction for multiple comparisons, p < 0.05/10 = 0.005 (B) or p < 0.05/13 = 0.0038 (D and F). **(G)** OL gain rate at 7, 14, 21, and 28 days is best predicted by OL loss rate 4, 7, 7, and 7 days prior, respectively. **(H)** Plot of strongest correlations from (F): OL gain rate at 7, 14, 21, and 28 days versus OL loss rate at 4, 7, 14, and 21 days, respectively (black, light blue, dark blue, yellow, respectively). Dashed line of equality.

By contrast, cumulative oligodendrocyte loss was more strongly predictive of oligodendrocyte gain rate (**Fig. 2C, D**). Oligodendrocyte loss at 7 days post-cuprizone predicted oligodendrocyte gain rate at 14 days post-cuprizone (**Fig. 2C**). More broadly, cumulative loss throughout the remyelination period was predictive of oligodendrocyte gain rate at several time points (**Fig. 2D**). Thus, oligodendrocyte gain rate is more responsive to the loss of oligodendrocytes than to oligodendrocyte number.

However, we observed that cumulative loss of oligodendrocytes up until the end of cuprizone treatment (45±3 % of total loss) did not predict oligodendrocyte gain rate at any point during remyelination (**Fig. 2D**), suggesting that the timing of oligodendrocyte loss was an important factor in the ability to induce future oligodendrocyte gain. To specifically test this, we assessed the correlation between oligodendrocyte loss rate and oligodendrocyte gain rate (**Fig. 2E, F**). We found that the best predictor of oligodendrocyte gain rate at any time point was the oligodendrocyte loss rate approximately seven days prior (**Fig. 2E-G**). Oligodendrocyte gain rates at 7, 14, 21, and 28 days were best predicted by the oligodendrocyte loss rates at 4, 7, 14, and 21 days, respectively, and were poorly predicted by loss rates at other time points (**Fig. 2E, F**). For example, while the oligodendrocyte loss rate at 7 days strongly predicted the oligodendrocyte gain rate at 14 days, it did not predict the oligodendrocyte gain rate at 28 days (**Fig. 2E, F**). Similarly, the oligodendrocyte loss rate at 21 days strongly predicted the gain rate at 28 days but did not predict the gain rate at 7 days (**Fig. 2F**). These data support the existence of acute signaling that occurs around the death of oligodendrocytes that induces new oligodendrocyte formation approximately one week later (**Fig. 2G**). Thus, recent oligodendrocyte loss – not oligodendrocyte population restoration – drives a temporally limited endogenous remyelination response.

Similar to cumulative oligodendrocyte loss and gain (**Fig. 1G**), we noticed that the relationship between oligodendrocyte loss rate at 7 days and gain rate at 14 days was not one-to-one (**Fig. 2E**). Thus, we next sought to determine how the scale of this relationship varied across the magnitude of oligodendrocyte loss or the phase of remyelination. To do this, we plotted oligodendrocyte gain rate at 7, 14, 21, and 28 days versus oligodendrocyte loss rate at 4, 7, 14, and 21 days, respectively (**Fig. 2H**). At low rates of oligodendrocyte loss (below approximately 1.5% per day), oligodendrocyte gain rate equaled or exceeded the rate of loss (**Fig. 2H**), regardless of the phase of remyelination (**Fig. 2H**). By contrast, at high rates of oligodendrocyte loss (greater than 1.5% per day), which subsided approximately one and a half weeks post-cuprizone (**Supplementary Fig. 1D**), oligodendrocyte gain rate could not keep up, never exceeding approximately 2.5% per day despite substantially higher loss rates (**Fig. 2H and Supplementary Fig. 1E**). Thus, while oligodendrocyte gain during remyelination is induced by recent oligodendrocyte loss, the number of new oligodendrocytes formed per lost oligodendrocyte depends on the rate at which the oligodendrocytes were lost.

### Thyromimetic treatment enhances oligodendrocyte gain during remyelination

Given the limited magnitude (**Fig. 1H-K**) and period (**Fig. 2E,F**) of endogenous remyelination and the overwhelming presence of chronically demyelinated lesions in multiple sclerosis patients^24,43,44^, identifying exogenous methods to enhance remyelination is a clinical imperative. Thus, we sought to evaluate the ability of a novel thyroid hormone receptor beta agonist (the thyromimetic LL-341070), to improve oligodendrocyte gain during remyelination. Thyroid hormone and thyroid hormone receptor beta agonists are potent inducers of OPC differentiation^45–47^ and enhancers of remyelination^48–52^, but their use as remyelination therapeutics has been precluded by peripheral side effects. However, the development of fatty acid amide hydrolase (FAAH)-targeted prodrugs has enabled preferential delivery of systemic doses to CNS tissues^53,54^. LL-341070 enters the central nervous system as an inactive prodrug, where it is hydrolyzed into its active form by FAAH and is thus able to act locally on thyroid hormone beta receptors (**Fig. 3A**). Following 3.5 weeks of 0.2% cuprizone administration, *Mobp-EGFP* mice were treated daily for the first three weeks of remyelination with 0.3 mg/kg or 0.1 mg/kg LL-three weeks of remyelination with 0.3 mg/kg or 0.1 mg/kg LL-341070 or vehicle (**Fig. 3B, C**). Mice underwent longitudinal *in vivo* two-photon imaging of visual cortex to allow for tracking of oligodendrocyte loss and gain throughout the demyelination and remyelination periods (**Fig. 3B, D**).

**Fig. 3.**
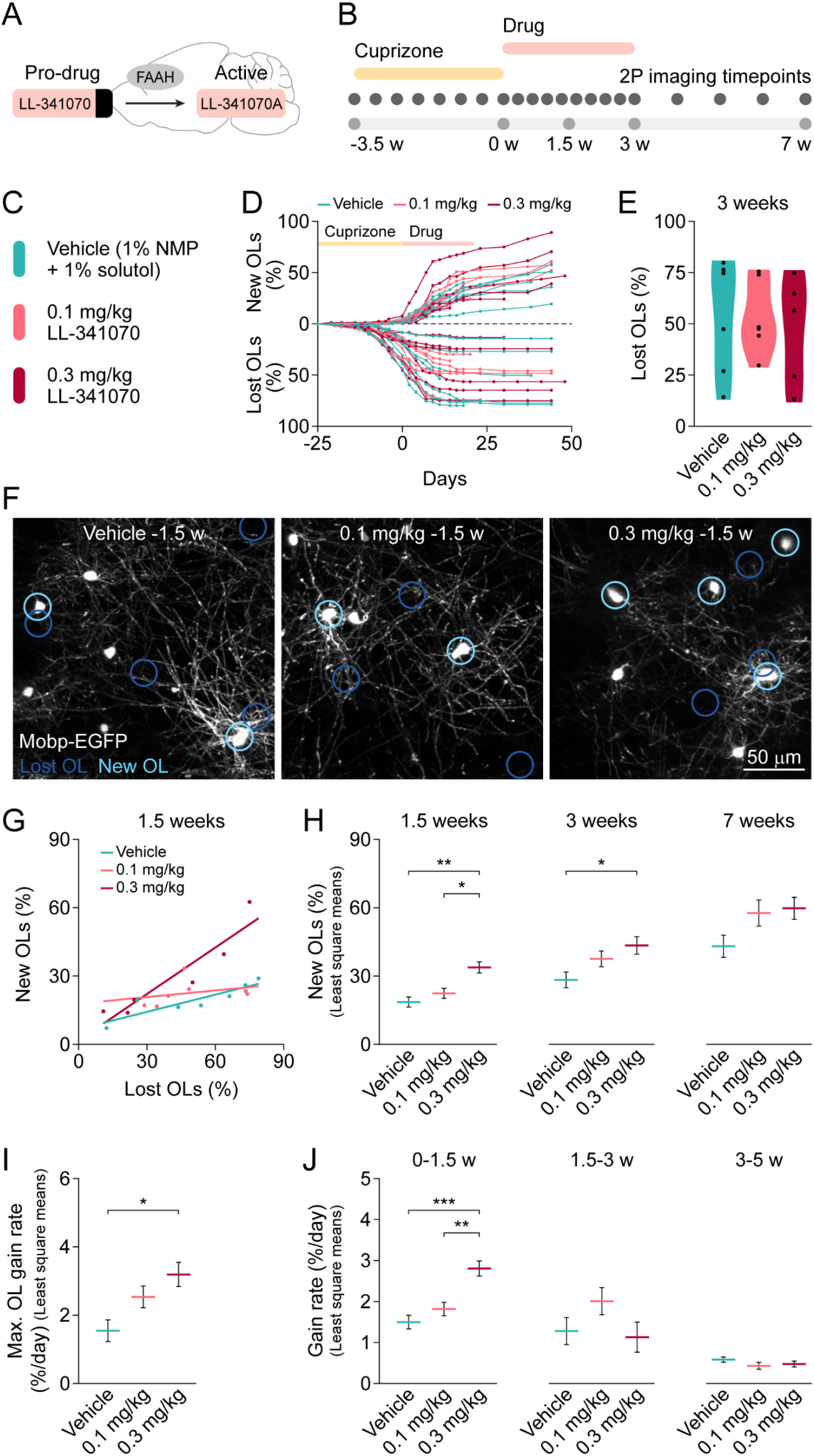
Thyromimetic treatment enhances oligodendrocyte gain during remyelination. **(A)** LL-341070 is a novel thyromimetic that enters the CNS as an inactive prodrug, where it is hydrolyzed by fatty acid amide hydrolase (FAAH) into its active form and targets thyroid receptors. **(B)** Experimental timeline of cuprizone-mediated demyelination, drug dosing, and longitudinal *in vivo* two-photon imaging. **(C)** Mice were dosed with either vehicle, 0.1 mg/kg LL-341070, or 0.3 mg/kg LL-341070. **(D)** Cumulative OL loss and gain as a percentage of baseline OLs in individual mice over time (vehicle: n = 7, 0.1 mg/kg: n = 7, 0.3 mg/kg: n = 6). **(E)** OL loss by 3 weeks did not differ between groups in mean or variance. **(F)** Representative images of V1 OLs at 1.5 weeks post-cuprizone. Locations of lost OLs (dark blue) and new OLs (light blue) are encircled. Images of different timepoints are shown in Supplementary Fig. 3. **(G)** Analysis of covariance with unequal slopes shows effect of treatment, OL loss, and interaction between OL loss and treatment on OL gain at 1.5 weeks. **(H)** High dose of LL-341070 increases OL gain at 1.5 weeks and 3 weeks. No effect detected on OL gain at 7 weeks. **(I)** High dose of LL-341070 increases the maximum OL gain rate during remyelination. **(J)** High dose of LL-341070 increases OL gain rate from 0-1.5 weeks. No effect detected on OL gain rate from 1.5-3 weeks or from 3-5 weeks. In **E**, Mean (vehicle: 53.24±10, n = 6; 0.1 mg/kg: 53.27±10, n = 6; 0.3 mg/kg: 46.8±10.96, n = 5; F(2, 14) = 0.12, p = 0.89, ANOVA). Variance (F(2, 14) = 0.89, p = 0.43, Brown-Forsythe). In **G**, Model (F(5, 14) = 12.66, **** p < 0.0001, ANOVA). Effects: treatment ((F(2) = 11.22, ** p = 0.0012), OL loss (F(1) = 28.98, **** p < 0.0001), interaction (F(2) = 7.65, ** p = 0.0057). Vehicle: n = 7, 0.1 mg/kg: n = 7, 0.3 mg/kg: n = 6. In **H**, OL gain at 1.5 weeks (vehicle: 18.56±2.21, n = 7; 0.1 mg/kg: 22.39±2.21, n = 7; 0.3 mg/kg: 33.77±2.45, n = 6; 0.3 mg/kg versus vehicle ** p = 0.0011, 0.3 mg/kg versus 0.1 mg/kg * p = 0.01, 0.1 mg/kg versus vehicle p = 0.46, Tukey HSD), 3 weeks (vehicle: 28.28±3.44, n = 6; 0.1 mg/kg: 37.53±3.45, n = 6; 0.3 mg/kg: 43.46±3.83, n = 5; 0.3 mg/kg versus vehicle * p = 0.033, 0.3 mg/kg versus 0.1 mg/kg p = 0.51, 0.1 mg/kg versus vehicle p = 0.19, Tukey HSD), and 7 weeks (treatment: (F(2) = 3.3713, p = 0.12; vehicle: n = 4, 0.1 mg/kg: n = 3, 0.3 mg/kg: n = 4). In **I**, Max. OL gain rate (vehicle: 1.54±0.32, n = 6; 0.1 mg/kg: 2.54±0.32, n = 6; 0.3 mg/kg: 3.19±0.35, n = 5; 0.3 mg/kg versus vehicle * p = 0.013, 0.3 mg/kg versus 0.1 mg/kg p = 0.38, 0.1 mg/kg versus vehicle p = 0.11, Tukey HSD). In **J**, OL gain rate 0-1.5 weeks (vehicle: 1.50±0.17, n = 7; 0.1 mg/kg: 1.82±0.17, n = 7; 0.3 mg/kg: 2.80±0.18, n = 6; 0.3 mg/kg versus vehicle *** p = 0.0003, 0.3 mg/kg versus 0.1 mg/kg ** p = 0.0036, 0.1 mg/kg versus vehicle p = 0.38, Tukey HSD), 1.5-3 weeks (vehicle: n = 6, 0.1 mg/kg: n = 6, 0.3 mg/kg: n = 5, F(5, 11) = 1, p = 0.46, ANOVA), and 3-5 weeks (vehicle: n = 5, 0.1 mg/kg: n = 3, 0.3 mg/kg: n = 4, F(5, 6) = 3.82, p = 0.067, ANOVA). * p < 0.05, ** p < 0.01, *** p < 0.001, least square mean ± SEM.

As observed in control mice (**Fig. 1D**), all groups experienced a range in the level of oligodendrocyte loss (**Fig. 3D, E**), with no difference in the mean or variance between groups (**Fig. 3E, F and Supplementary Fig. 3A**). Since oligodendrocyte gain depends on loss (**Fig. 1E and Supplementary Fig. 2A, B**), it was important to account for the level of loss incurred by each mouse when testing the role of thyromimetic treatment on oligodendrocyte gain. To do this, we used an analysis of covariance with unequal slopes to test the effect of treatment on oligodendrocyte gain while considering the effect of oligodendrocyte loss and the interaction between oligodendrocyte loss and treatment (**Fig. 3G and Supplementary Fig. 4A, B**).

At 1.5 weeks, we found a significant effect of treatment, oligodendrocyte loss, and the interaction of oligodendrocyte loss and treatment on oligodendrocyte gain (**Fig. 3G**). To compare the treatment groups, we used the modeled means of oligodendrocyte gain for each treatment group (least square means) at the average level of oligodendrocyte loss (**Fig. 3H**). We found that mice treated with the high dose of LL-341070 gained substantially more new oligodendrocytes by 1.5 weeks as compared to mice treated with the low dose or vehicle (**Fig. 3F, H, and Supplementary Fig. 3A**). Similarly, at the end of drug treatment (3 weeks of remyelination), mice treated with the high dose of LL-341070 had gained more new oligodendrocytes than did mice treated with vehicle (**Fig. 3H and Supplementary Fig. 4A**). By 7 weeks, there were no detectable differences between groups (**Fig. 3H and Supplementary Fig. 4B**). Thus, thyromimetic treatment enhances oligodendrocyte gain during remyelination.

Given the failure of endogenous remyelination to surpass an oligodendrocyte gain rate of approximately 2.5% per day (**Fig. 2H and Supplementary Fig. 1E**), we next sought to understand how thyromimetic treatment impacted oligodendrocyte gain rate. Like oligodendrocyte gain, oligodendrocyte gain rate depends on oligodendrocyte loss (**Fig. 2C, D and Supplementary Fig. 2C-E**); thus, an analysis of covariance with unequal slopes was used in the same manner to isolate the effects of treatment on oligodendrocyte gain rate. We found that the high dose of LL-341070 increased the maximum oligodendrocyte gain rate during remyelination by two-fold (**Fig. 3I and Supplementary Fig. 4C-E**). On average, mice treated with the high dose of LL-341070 had a maximum gain rate of 3.2±0.35 (SEM) % per day, with multiple mice exceeding the observed endogenous limit of 2.5% per day (**Fig. 3I, Supplementary Fig. 4D, E**).

Next, to understand the dynamics with which thyromimetic treatment impacted oligodendrocyte gain, we analyzed oligodendrocyte gain rate across different phases of remyelination. We found that the high dose of LL-341070 increased oligodendrocyte gain rate specifically during the first 1.5 weeks of its administration (**Fig. 3H and Supplementary Fig. 4F**). Unexpectedly, during the second half of treatment (1.5-3 weeks) or following treatment (3-5 weeks), there was no impact of LL-341070 on oligodendrocyte gain rate (**Fig. 3H and Supplementary Fig. 4G, H**). Thus, thyromimetic treatment accelerates oligodendrocyte gain during remyelination by transiently increasing oligodendrocyte gain rate during the first half of treatment.

### Severity of demyelination determines efficacy of remyelination therapy

In our analysis of covariance with unequal slopes, we found that the effect of treatment on oligodendrocyte gain was significantly modulated by the level of oligodendrocyte loss (**Fig. 3G**). To further dissect this finding, we assessed the ability of thyromimetic treatment to augment oligodendrocyte gain at various levels of oligodendrocyte loss. We found that at 25% oligodendrocyte loss, LL-341070 treatment had no significant impact on oligodendrocyte gain by 1.5 weeks (**Fig. 4A**), while at 75% oligodendrocyte loss, the high dose of LL-341070 more than doubled the amount of oligodendrocyte gain (**Fig. 4A**).

**Fig. 4.**
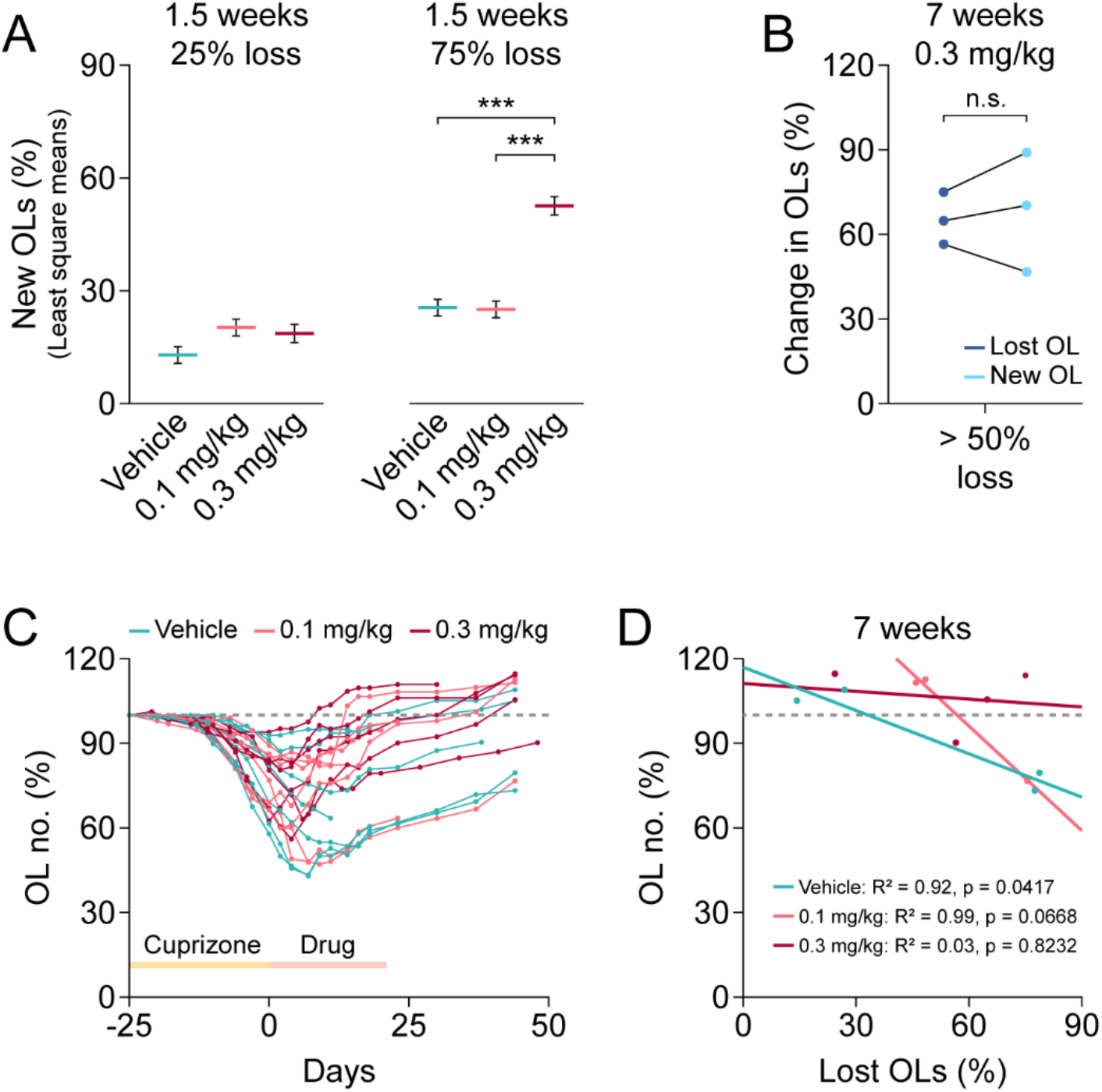
Thyromimetic LL-341070 enhances oligodendrocyte gain to a greater extent after moderate or severe demyelination. **(A)** At 1.5 weeks, high dose of LL-341070 does not increase OL gain at 25% OL loss but dramatically improves OL gain at 75% OL loss. **(B)** At > 50% OL loss, mice treated with the high dose of LL-340170 make as many new OLs as they lose by 7 weeks. **(C)** OL number as a percentage of baseline OLs in individual mice over time (vehicle: n = 7, 0.1 mg/kg: n = 7, 0.3 mg/kg: n = 6). Dashed line at 100%. **(D)** OL number at 7 weeks is not correlated with magnitude of OL loss by 7 weeks in mice treated with high dose of LL-341070 (n = 4, linear regression) or low dose of LL-341070 (n = 3, linear regression) but is correlated in mice treated with vehicle (n = 4, linear regression). Dashed line at 100%. * p < 0.05. In **A**, OL gain at 25% loss (vehicle: 12.97 ± 2.21, n = 7; 0.1 mg/kg: 20.24 ± 2.21, n = 7; 0.3 mg/kg: 18.69 ± 2.45, n = 6; 0.3 mg/kg versus vehicle F(1, 14) = 1.7405, p = 0.21; 0.3 mg/kg versus 0.1 mg/kg F(1, 14) = 0.1, p = 0.75; 0.1 mg/kg versus vehicle F(1, 14) = 2.06, p = 0.17; post-hoc contrast at 25% loss with Bonferroni correction) and 75% loss (vehicle: 25.57 ± 2.21, n = 7; 0.1 mg/kg: 25.07 ± 2.21, n = 7; 0.3 mg/kg: 52.66 ± 2.45, n = 6; 0.3 mg/kg versus vehicle F(1, 14) = 26.62, *** p = 0.0001; 0.3 mg/kg versus 0.1 mg/kg F(1, 14) = 22.65, *** p = 0.0003; 0.1 mg/kg versus vehicle F(1, 14) = 0.0095, p = 0.92; post-hoc contrast at 75% loss with Bonferroni correction). *** p < 0.001/3 = 0.00033, least square mean ± SEM. In **B**, Change in OLs at > 50% loss (lost OL: 65.45 ± 5.34, new OL: 68.71 ± 12.24, n = 3, t(2) = 0.47, p = 0.69, paired t test). n.s. not significant, * p < 0.05, mean ± SEM.

Given this preferential improvement of remyelination at high loss levels, we sought to determine whether thyromimetic treatment would ameliorate the regeneration deficit observed at high loss levels in control mice (**Fig. 1G-I**). Unlike control mice, mice treated with the high dose of LL-341070 successfully gained as many new cells as they lost even at high levels of loss (**Fig. 4B**), restoring oligodendrocyte numbers across all loss levels (**Fig. 4C, D**) and eliminating the relationship between oligodendrocyte loss and the number of oligodendrocytes at seven weeks (**Fig. 4D**). Thus, endogenous remyelination fails to restore oligodendrocyte numbers in mice with high levels of oligodendrocyte loss, while thyromimetic treatment specifically ameliorates this deficit.

### Regenerative oligodendrogenesis is enhanced more quickly and robustly by LL-341070 compared to clemastine fumarate

Considering these promising results, we sought to compare the efficacy of LL-341070 in promoting oligodendrocyte gain during remyelination to the pioneering remyelination therapeutic clemastine fumarate^4,35^. In parallel cohorts, mice were treated with clemastine fumarate (10 mg/kg) or vehicle (10% DMSO) for the first three weeks of remyelination and underwent longitudinal *in vivo* two-photon imaging of V1 to track oligodendrocyte loss and gain over time (**Fig. 5A, B**). Mice treated with vehicles for clemastine and LL-341070 were indistinguishable (**Supplementary Fig. 5A-I**) and thus were combined for statistical comparisons (**Fig. 5A**). Comparisons were made to mice treated with the high dose of LL-341070 (**Fig. 3, 4**).

**Fig. 5.**
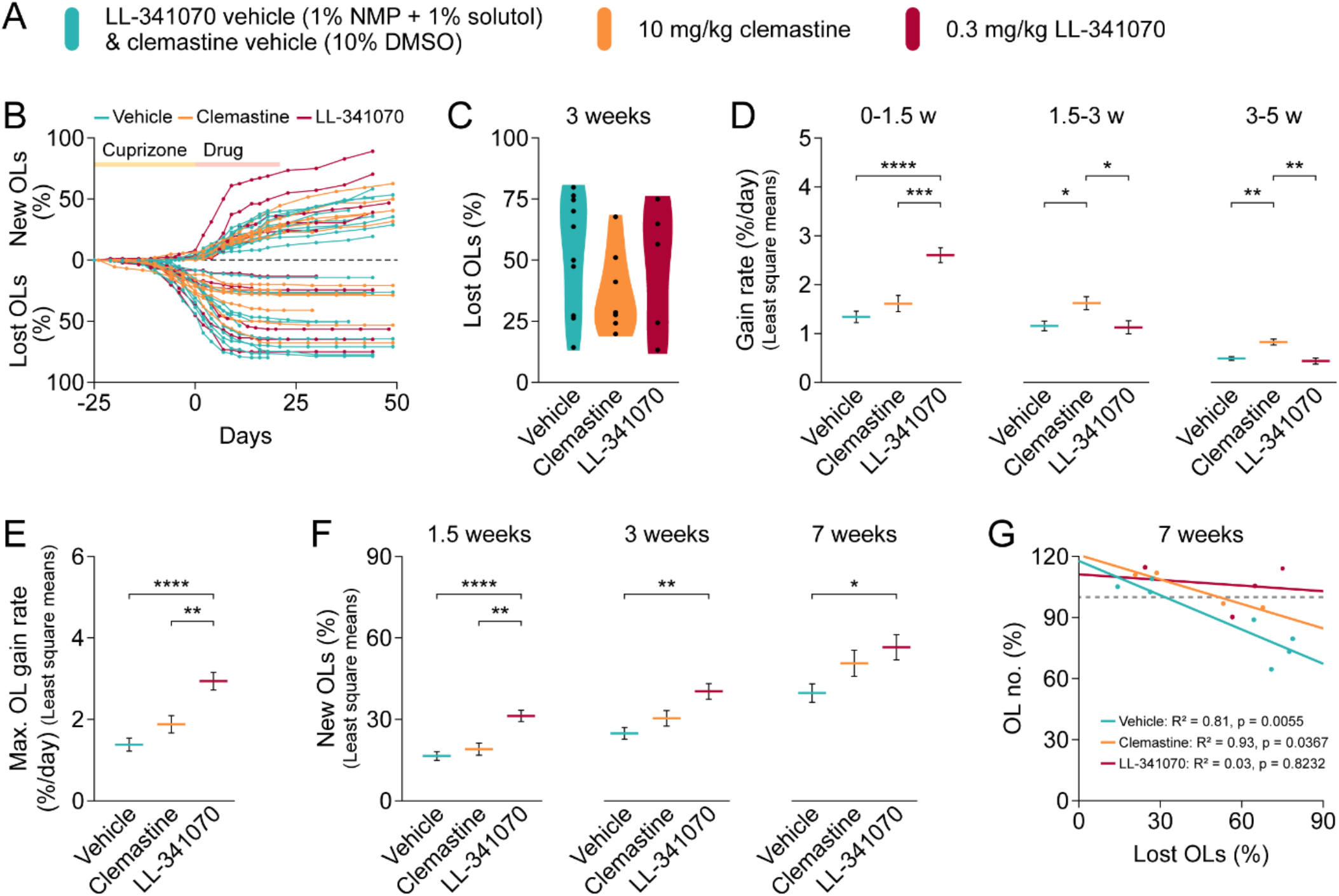
Thyromimetic enhances oligodendrocyte gain during remyelination more quickly and robustly than clemastine fumarate. **(A)** Mice were dosed with either vehicle, 10 mg/kg clemastine fumarate, or 0.3 mg/kg LL-341070. Mice treated with vehicle for LL-341070 and clemastine fumarate were combined for statistical comparisons. **(B)** Cumulative OL loss and gain as a percentage of baseline OLs in individual mice over time (vehicle: n = 11, clemastine: n = 7, LL-341070: n = 6). **(C)** OL loss by 3 weeks did not differ between groups in mean or variance. **(D)** LL-341070 increases OL gain rate from 0-1.5 weeks. Clemastine increases OL gain rate from 1.5-3 weeks and from 3-5 weeks. **(E)** LL-341070 increases maximum OL gain rate. **(F)** LL-341070 increases OL gain at 1.5 weeks, 3 weeks, and 7 weeks. **(G)** OL number at 7 weeks is not correlated with magnitude of OL loss by 7 weeks in mice treated with LL-341070 (n = 4, linear regression) but is correlated in mice treated with clemastine (n = 4, linear regression) and vehicle (n = 7, linear regression). Dashed line at 100%. In **C**, Mean (vehicle: 52.95 ± 7.15, n = 10; clemastine: 37.07 ± 19.19, n = 7; LL-341070: 46.80 ± 10.11, n = 5; F(2, 19) = 1.02, p = 0.3806, ANOVA). Variance (F(2, 19) = 0.6904, p = 0.5135, Brown-Forsythe). In **D**, OL gain rate 0-1.5 weeks (vehicle: 1.34 ± 0.12, n = 11; clemastine: 1.61 ± 0.17, n = 7; LL-341070: 2.60 ± 0.15, n = 6; LL-341070 versus vehicle **** p < 0.0001, LL-341070 versus clemastine *** p = 0.0009, clemastine versus vehicle p = 0.39, Tukey HSD), 1.5-3 weeks (vehicle: 1.16 ± 0.10, n = 10; clemastine: 1.62 ± 0.13, n = 7; LL-341070: 1.12 ± 0.13, n = 5; clemastine versus LL-341070 * p = 0.045, clemastine versus vehicle * p = 0.031, LL-341070 versus vehicle p = 0.98, Tukey HSD), and 3-5 weeks (vehicle: 0.49 ± 0.04, n = 9; clemastine: 1.62 ± 0.13, n = 5; LL-341070: 1.12 ± 0.13, n = 4; clemastine versus LL-341070 ** p = 0.002, clemastine versus vehicle ** p = 0.0014, LL-341070 versus vehicle p = 0.76, Tukey HSD). In **E**, Max. OL gain rate (vehicle: 1.38 ± 0.16, n = 10; clemastine: 1.88 ± 0.21, n = 7; LL-341070: 2.94 ± 0.22, n = 5; LL-341070 versus vehicle **** p < 0.0001, LL-341070 versus clemastine ** p = 0.0080, clemastine versus vehicle p = 0.18, Tukey HSD). In **F**, OL gain 1.5 weeks (vehicle: 16.50 ± 1.61, n = 11; clemastine: 19.03 ± 2.26, n = 7; LL-341070: 31.25 ± 2.08, n = 6; LL-341070 versus vehicle **** p < 0.0001, LL-341070 versus clemastine ** p = 0.0024, clemastine versus vehicle p = 0.64, Tukey HSD), 3 weeks (vehicle: 24.84 ± 2.12, n = 10; clemastine: 30.39 ± 2.82, n = 7; LL-341070: 40.32 ± 2.88, n = 5; LL-341070 versus vehicle ** p = 0.0014, LL-341070 versus clemastine p = 0.063, clemastine versus vehicle p = 0.29, Tukey HSD), and 7 weeks (vehicle: 39.70 ± 3.40, n = 7; clemastine: 50.65 ± 4.83, n = 4; LL-341070: 56.59 ± 4.66, n = 4; LL-341070 versus vehicle *p = 0.04, LL-341070 versus clemastine p = 0.66, clemastine versus vehicle p = 0.21, Tukey HSD). *p < 0.05, ** p < 0.01, *** p < 0.001, **** p < 0.0001, least square mean ± SEM.

First, we assessed oligodendrocyte loss between groups. Mice treated with vehicle, clemastine fumarate, or LL-341070 exhibited a wide range in loss, with no differences in mean or variance between groups (**Fig. 5C**). Next, we compared oligodendrocyte gain during remyelination between groups and found differences in the timing and magnitude. While LL-341070 treatment robustly increased oligodendrocyte gain rate during the first half of treatment, clemastine fumarate more subtly increased oligodendrocyte gain rate over a longer period, beginning during drug administration and continuing after treatment ended (**Fig. 5D and Supplementary Fig. 6A-C**). Nevertheless, the maximum oligodendrocyte gain rate induced by clemastine fumarate did not reach that induced by LL-341070 (**Fig. 5E and Supplementary Fig. 6D-F**).

These dynamics resulted in more new oligodendrocytes in mice treated with LL-341070 than in mice treated with clemastine fumarate or vehicle at 1.5 weeks (**Fig. 5F and Supplementary Fig. 6G**). By three and seven weeks, despite the prolonged impact of clemastine fumarate on oligodendrocyte gain rate, cumulative oligodendrocyte gain was not detectably higher in mice treated with clemastine fumarate than in mice treated with vehicle; meanwhile, thyromimetic-treated mice maintained higher levels (**Fig. 5F and Supplementary Fig. 6H, I**).

Finally, we assessed whether clemastine fumarate – like LL-341070 – was able to ameliorate the regeneration deficit at high oligodendrocyte loss levels. We found that this was not the case; rather, a strong correlation persisted between oligodendrocyte loss and oligodendrocyte number at seven weeks in mice treated with clemastine fumarate (**Fig. 5G and Supplementary Fig. 6J**), indicating that clemastine fumarate is unable to fully eliminate the regeneration deficit experienced by mice with high levels of oligodendrocyte loss. Overall, while clemastine fumarate enhances oligodendrocyte gain rate, the tested dose of LL-341070 is more effective at augmenting oligodendrocyte gain rate, cumulative oligodendrocyte gain, and restoration of oligodendrocyte numbers.

### The recovery of single-neuron latency is accelerated by LL-341070

Given the robust effects of LL-341070 in promoting remyelination, we sought to test whether this treatment could also enhance the restoration of neuronal function following demyelination. Demyelination delays visual-induced population responses in the visual cortex known as visual evoked potentials (VEPs), while remyelination rescues VEP latency deficits^25,34,55^. Thus, VEP latency measurements are often used as supportive evidence to help confirm a clinical diagnosis of multiple sclerosis as well as a biomarker for functional improvement in clinical trials^35,56^. However, how demyelination and remyelination affect the visual responses of individual neurons *in vivo* has remained unexplored. To this end, we used Neuropixels, high-density extracellular recording probes that allow for the simultaneous evaluation of hundreds of neurons, implanted into V1 (**Fig. 6A**). To capture different aspects of visual processing, animals were presented with brief (50 ms), full visual-field luminance changes towards both dark and bright (**Supplementary Fig. 7A, B**) during the electrophysiological recording. We probed the timing of visually-evoked neural activity in healthy mice, demyelinated mice (end of cuprizone), and mice treated with thyromimetic or vehicle at the end of treatment (three weeks post-cuprizone) or four weeks after the end of the treatment (seven weeks post-cuprizone) (**Fig. 6B**).

**Fig. 6.**
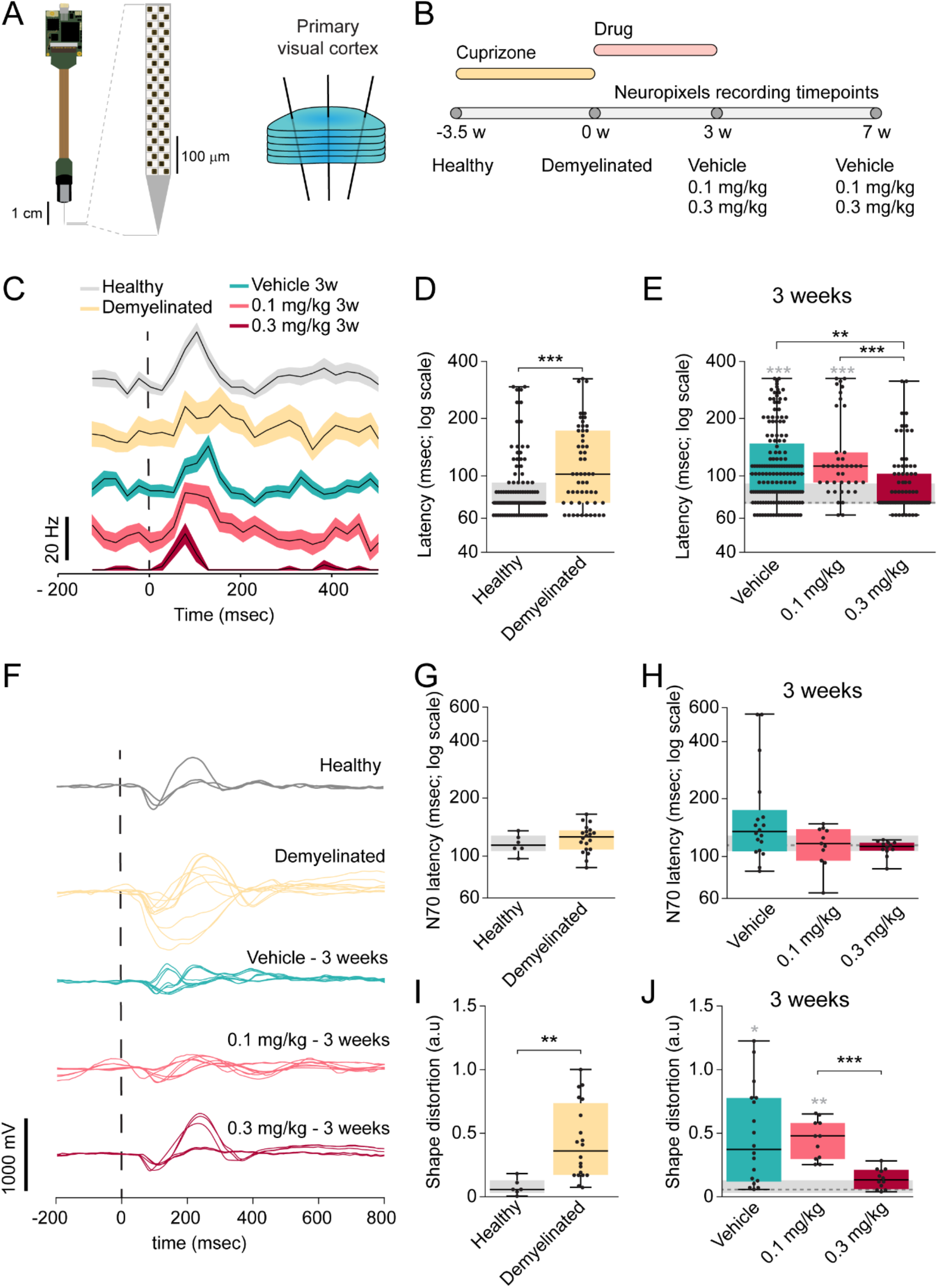
LL-341070 treatment accelerates recovery of visual function after demyelination. **(A)** Neuronal activity was recorded during visual stimuli presentation using three *Neuropixels* probes inserted in the primary visual cortex. **(B)** Recordings were terminal procedures and were performed in different animals to capture the effects of demyelination and endogenous or therapeutically-induced remyelination. **(C)** Representative traces (black lines) and SEM (shaded regions) of single-neuron responses to dark flash. Dashed line represents stimulus onset. **(D)** Single-neuron responses to dark or bright flash are delayed after cuprizone-induced demyelination. **(E)** Treatment with high dose LL-341070 for three weeks can reestablish the typical neuronal latencies. **(F)** Representative visual evoked potentials (VEPs) in response to a dark flash. Dashed line represents stimulus onset. **(G)** VEP N70 latency to dark or bright flash is not significantly altered by cuprizone. **(H)** All treatment groups show similar VEP N70 latency after three weeks of remyelination: **(I)** VEP shape in response to both dark and bright flash is distorted by cuprizone treatment. **(J)** VEP shape distortion is rescued by three-week treatment with high dose of LL-341070. In **D**, Median (IQR) [healthy: 72.25 (72.25-92.33), n = 104 responses from 6 probes from 3 mice; demyelinated: 102.37 (72.25-172.65), n = 55 responses from 7 probes from 5 mice; Z = 4.37, p < 0.0001 Wilcoxon]. In **E**, Median (IQR) [vehicle: 102.37 (72.25-147.55), n = 157 responses from 8 probes from 3 mice; 0.1 mg/kg: 112.41 (92.33-132.49), n = 40 responses from 6 probes from 4 mice; 0.3 mg/kg: 72.25 (72.25-102.37), n = 76 responses from 4 probes from 2 mice; F(3)= 43.17; p < 0.0001, Kruskal-Wallis; vehicle vs healthy *** p < 0.0001, 0.1 mg/kg vs healthy *** p < 0.0001, 0.3 mg/kg vs 0.1 mg/kg *** p = 0.0002, 0.3 mg/kg vs vehicle *** p = 0.0022, Steel-Dwass]. In **G**, Median (IQR) [healthy: 112.925(105.42-126.73), n = 6 VEPs from 3 probes from 2 mice; demyelinated: 124.73 (107.42-134.93), n = 20 VEPs from 10 probes from 4 mice; Z = -0.79, p = 0.4287, Wilcoxon]. In **H**, Median (IQR) [vehicle: 133.53 (105.12-173.05), n = 18 VEPs from 9 probes from 3 mice; 0.1 mg/kg: 115.73 (94.41-137.34), n = 11 VEPs from 8 probes from 3 mice; 0.3 mg/kg: 11.73 (106.22-116.83), n = 12 VEPs from 6 probes from 2 mice; F(3)= 6.1077; p = 0.1065, Kruskal-Wallis]. In **I**, Median (IQR) [healthy: 0.0567 (0.03-0.13), n = 6 measurements from 3 probes from 2 mice; demyelinated: 0.36 (0.17-0.74), n = 20 measurements from 10 probes from 4 mice; Z = -3.1342, p = 0.0017, Wilcoxon]. In **J**, Median (IQR) [vehicle: 0.37 (0.12-0.78), n = 18 measurements from 9 probes from 3 mice; 0.1 mg/kg: 0.48 (0.3-0.58), n = 11 measurements from 8 probes from 3 mice; 0.3 mg/kg: 0.13 (0.059-0.21), n = 12 measurements from 6 probes from 2 mice; F(3)= 20.38; p = 0.0001, Kruskal-Wallis; vehicle vs healthy # p = 0.024, 0.1 mg/kg vs healthy ## p = 0.006, 0.3 mg/kg vs 0.1 mg/kg *** p = 0.0005, Steel-Dwass].

Taking advantage of the multiple recording channels in the *Neuropixels* probes, we isolated single neurons and analyzed how their response latencies were altered by demyelination and remyelination therapy (**Fig. 6C**). Single-neuron responses were delayed after demyelination (**Fig. 6D**). Three-week treatment with high dose LL-341070 restored the latency to the flashed stimulus, while vehicle-treated mice maintained delayed responses (**Fig. 6E**). When assessing luminance changes to dark or bright individually, we found that responses to both stimuli were delayed by cuprizone (**Supplementary Fig. 8A, D**) and rescued by high dose LL-341070 for 3 weeks (**Supplementary Fig. 8B, E**). Meanwhile, at this timepoint, responses to dark remained delayed in vehicle-treated mice, while responses to bright were still slowed in mice treated with low dose LL-341070 (**Supplementary Fig. 8B, E**). Overall, these data support an effect of thyromimetic treatment in accelerating the recovery of single-neuron response latencies after demyelination.

### Restoration of visual evoked potentials is accelerated by LL-3411070

VEPs capture the summed input and neuronal network response to a visual stimulus in cortex, which may reflect the underlying deficits in neuronal activity and connectivity in V1. Therefore, we generated the VEP data from the local field potentials (LFPs) recorded by the Neuropixels probes (**Supplementary Fig. 7C**). We first assessed the timing and amplitude of the VEP (**Fig. 6F**). We found no significant differences between groups in latency of the first negative peak (N70), which is thought to correlate to the arrival of the thalamic input in V1 (**Fig. 6G, H and Supplementary Fig. 9A, B, D and E**). Similarly, we only detected small changes in amplitude of the first positive peak (P100) in relation to N70 (**Supplementary Fig. 10, A, B, D, E, G and H**). However, we observed that VEP structure appeared less organized in demyelinated mice compared to healthy controls (**Fig. 6F**). To quantify these differences, we developed a shape distortion metric, which quantifies the spatial distance correlation between a given trial VEP and the average VEP of healthy mice (see **Methods**). Using this metric, we found that demyelinated mice exhibited altered VEP shape as compared to healthy controls (**Fig. 6I and Supplementary Fig. 9G, J**). Importantly, the demyelination-triggered distortion in VEP shape was rescued by three weeks post-cuprizone in mice treated with the high dose of LL-341070, while vehicle-treated mice maintained an altered VEP shape (**Fig. 6J**). Therefore, in concert with accelerating oligodendrocyte regeneration, thyromimetic treatment enhances the recovery of the V1 neuronal population response to visual stimuli.

### Partial restoration of oligodendrocyte population is sufficient to recover delayed visual responses

Since visual responses were still impaired in vehicle- and low dose-treated mice after three weeks of remyelination, when remyelination was still ongoing, we assessed visual responses four weeks later, after seven weeks of remyelination. Between three and seven weeks post-cuprizone, single neuron responses improved in mice treated with vehicle or low dose LL-341070, while neuronal responses of mice treated with high dose LL-341070 did not change (**Fig. 7A**). Thus, mice treated with high dose LL-341070 reached a plateau of recovery by three weeks while the other groups continued to improve. By seven weeks of remyelination, all groups reached this recovery plateau. At this time, we found no differences between the treatment groups in any of the evaluated parameters and responses were similar in range to those of healthy mice (**Fig. 7B-D, Supplementary Fig. 8C, F, Supplementary Fig. 9C, F, I and L, and Supplementary Fig. 10C, F and I**). However, this recovery plateau may not represent complete restoration as we observed some small differences from healthy mice for single neuron latency of the vehicle group and for the shape distortion of the VEP responses (**Fig. 7B, D**).

**Fig. 7.**
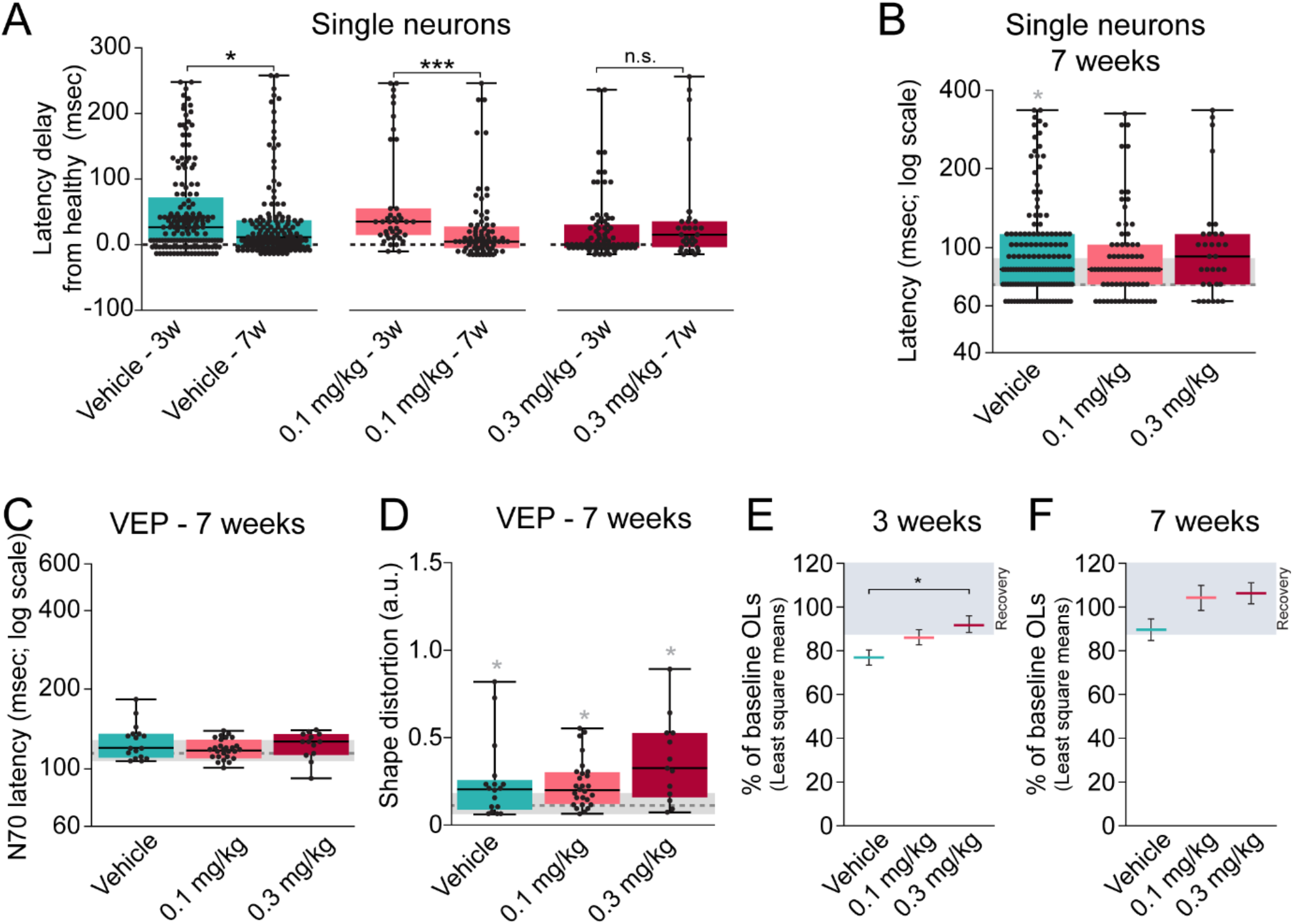
Functional rescue of visual neuronal function is attained with partial restoration of the oligodendrocyte population. **(A)** Single-neuron latency improves between 3 weeks and 7 weeks in animals treated with vehicle or low-dose LL-341070, while it is already recovered at 3 weeks by treatment with high-dose LL-341070. **(B)** Single-neuron responses to dark or bright flash are equivalent in all groups after seven weeks of remyelination. **(C)** Seven weeks post-demyelination, there are no differences between groups in VEP N70 latency. **(D)** Seven weeks post-demyelination, there are no differences between groups in the VEP shape. **(E)** Oligodendrocyte population has been restored to at least ∼87% (% of baseline) after three weeks of high dose LL-341070, but remains below this threshold in mice that received vehicle or low dose LL-341070 (left). **(F)** After seven weeks of remyelination, oligodendrocyte population surpassed ∼87% (% of baseline) in all groups (right). Shaded area signifies recovery plateau. In **A**, Median (IQR) vehicle [3-weeks: 25.1 (5.02-70.28), n = 157 responses from 8 probes from 3 mice; 7-weeks: 10.04 (−5.02-35.14), n = 144 responses from 11 probes from 4 mice; Z = -2.3, p = 0.022, Wilcoxon]; Median (IQR) 0.1 mg/kg [3-weeks: 35.14 (15.06-55.22), n = 40 responses from 6 probes from 4 mice; 7-weeks: 5.02 (−5.02-27.61), n = 77 responses from 8 probes from 3 mice; Z = 4.05, p < 0.0001, Wilcoxon]; and Median (IQR) 0.3 mg/kg [3-weeks: 0 (−5.02-30.12), n = 76 responses from 4 probes from 2 mice; 7-weeks: 15.06 (−3.76-35.14), n = 32 responses from 3 probes from 1 mouse; Z = 1.18, p = 0.238, Wilcoxon]. In **B**, Median (IQR) [vehicle: 82.29 (72.25-112.41), n = 144 responses from 11 probes from 4 mice; 0.1 mg/kg: 82.29 (72.25-102.37), n = 77 responses from 8 probes from 3 mice; 0.3 mg/kg: 92.33 (72.25-112.41), n = 32 responses from 3 probes from 1 mouse; F(3)= 10.8514; p = 0.0126, Kruskal-Wallis; vehicle vs healthy * p = 0.0133, Steel-Dwass]. In **C**, Median (IQR) [vehicle: 118.53 (108.92-134.33), n = 8 probes from 3 mice; 0.1 mg/kg: 116.13 (108.52-127.73), n = 14 probes from 5 mice; 0.3 mg/kg: 125.33 (111.72-134.13), n = 7 probes from 3 mice; F(3)= 3.322; p = 0.3446, Kruskal-Wallis]. In **D**, Median (IQR) [vehicle: 0.21 (0.09-0.26), n = 17 measurements from 9 probes from 3 mice; 0.1 mg/kg: 0.2 (0.12-0.3), n = 27 measurements from 14 probes from 5 mice; 0.3 mg/kg: 0.32 (0.16-0.53), n = 13 measurements from 9 probes from 3 mice; F(3)= 13.22; p = 0.0042, Kruskal-Wallis; vehicle vs healthy * p = 0.043, 0.1 mg/kg vs healthy * p = 0.012, 0.3 mg/kg vs healthy * p = 0.023, Steel-Dwass]. In **E**, OL number at 3 weeks as a percentage of baseline OLs (Least square mean ± SEM): (vehicle: 77.19 ± 3.48, n = 6; 0.1 mg/kg: 86.52 ± 3.49, n = 6; 0.3 mg/kg: 92.5 ± 3.87, n = 5; 0.3 mg/kg versus vehicle * p = 0.033, Tukey HSD). In **F**, OL number at 7 weeks as a percentage of baseline OLs (Least square mean ± SEM): (vehicle: 89.77 ± 4.91, n = 4; 0.1 mg/kg: 104.44 ± 5.77, n = 3; 0.3 mg/kg: 106.51 ± 4.88, n = 4).

Given that thyromimetic treatment accelerated the restoration of both oligodendrocyte numbers and neuronal function, we sought to investigate the level of remyelination necessary to recover neuronal function. Three weeks post-cuprizone, mice treated with vehicle, low, and high dose of LL-341070 had 77±3, 86±3, and 92±4 (least square means±SEM) % of their baseline oligodendrocyte populations, respectively (**Fig. 7E**). After seven weeks of remyelination, when recovery of single-neuron function was equivalent across groups, mice treated with vehicle, low, and high doses of LL-341070 had 90±5, 104±6 and 106±5 (least square means±SEM) % of their baseline oligodendrocyte populations, respectively (**Fig. 7F**). Thus, neuronal responses recovered when the oligodendrocyte number was restored to approximately 87% of the baseline population and did not improve above this threshold. This indicates that partial restoration of oligodendrocytes is sufficient to achieve recovery of neuronal responses, while further generation of oligodendrocytes past this threshold does not provide additional benefit.

## Discussion

Myelin loss, including in visual gray matter, is a common feature of several neurodegenerative diseases and injury conditions, and is present in normal aging^8–14^. In addition, myelin is malformed or present in insufficient levels in several neurodevelopmental and neuropsychiatric disorders^57–62^. By promoting formation of new oligodendrocytes and myelin, remyelination therapies may be clinically important for numerous neurological conditions. However, progress may be hampered by our incomplete understanding of the remyelination process and lack of clinically available remyelination therapies.

In this study, we used *in vivo* microscopy and neurophysiology approaches to unravel important aspects of remyelination in the visual system. We found that oligodendrocyte gain during remyelination is driven by recent oligodendrocyte loss and demonstrated that oligodendrocyte regeneration fails when high rates of demyelination occur too quickly. Advancing our understanding of therapeutic-induced remyelination, we found that distinct drugs stimulate oligodendrocyte gain with different temporal dynamics and that remyelination therapy is most effective when demyelination is moderate or severe. Importantly, we demonstrated that the novel thyromimetic LL-341070 is a highly potent remyelination therapeutic, which accelerates oligodendrocyte regeneration and recovery of neuronal function. Moreover, our data showed that even partial remyelination can restore visual neuronal function. Several of these findings may have important implications for future therapeutic strategies.

While remyelination is widely observed following a demyelinating injury^19,22,41,42^, the cellular and molecular triggers of this response are unknown. We found that oligodendrocyte gain rate during remyelination is driven by recent oligodendrocyte loss rather than a drive to reestablish oligodendrocyte numbers (**Fig. 2E-G**), indicating that acute signaling around the loss of myelinating oligodendrocytes induces new oligodendrocyte formation. However, the exact source of the signal is unknown. Previous work found that new oligodendrocyte generation temporally correlates better with myelin loss and microglial activation than with the laser ablation of oligodendrocyte cell bodies^41^. Together with these findings, our work suggests that the pro-remyelinating signal(s) may derive from microglial activation or phagocytosis, oligodendrocyte/myelin degradation or debris, or acute signaling from recently demyelinated axons. Identification of the molecular triggers of remyelination could have important clinical implications, as it may further illuminate causes of remyelination failure and provide additional targets to enhance remyelination.

Despite the occurrence of endogenous remyelination, it is often insufficient, leading to chronic demyelination^24^. We have described two factors that may contribute to this phenomenon. First, we found evidence of a critical time window for endogenous remyelination shortly following the loss of oligodendrocytes (**Fig. 2E-G**). For endogenous remyelination to be maximally efficacious, it is essential that new oligodendrocytes can form and survive in this period. However, prolonged inhibition extending beyond the end of oligodendrocyte loss may occur during demyelinating disease, precluding any endogenous remyelination response. Second, we observed deficits in endogenous remyelination in response to high rates of oligodendrocyte loss (**Fig. 2H**). Mice with moderate or severe demyelination failed to restore their initial oligodendrocyte populations (**Fig. 1K**) due to extended periods where oligodendrocyte loss exceeded 1.5% per day (**Supplementary Fig. 2D**). At these loss rates, endogenous rates could not keep up and never exceeded 2.5% per day, despite substantially higher loss rates (**Fig. 2H**). It is unclear what limits the endogenous remyelination response specifically at high loss rates in this non-inflammatory demyelination model, though several non-inflammatory remyelination inhibitors have been identified including accumulation of extracellular matrix molecules, myelin debris, and Nogo receptor 1 signaling^26,63,64^. Regardless, enhancing OPC differentiation using LL-341070 induced oligodendrocyte gain rates greatly exceeding 2.5% per day (**Fig. 3I, J, and Supplementary Fig. 4E, F**) and restored oligodendrocyte numbers (**Fig. 4C, D**), supporting the longstanding idea that there is untapped differentiation potential of OPCs in endogenous remyelination^65^, which we find limits the ability of the endogenous system to sufficiently respond to high rates of oligodendrocyte loss.

Interestingly, while remyelination therapy was incredibly effective in augmenting oligodendrocyte gain in mice with moderate or severe demyelination, thyromimetic treatment did not substantially impact oligodendrocyte gain in mice with mild demyelination (**Fig. 4A, D, and Fig. 5G**). Why is remyelination therapy less effective at low oligodendrocyte loss levels? Previous studies have indicated that the survival of differentiating OPCs is approximately 20% in healthy adult mouse cortex^39^. One possibility is that during mild demyelination, a similarly low rate of survival could substantially dampen the effects of therapeutic-induced OPC differentiation on oligodendrocyte gain, acting as a safeguard against excessive remyelination. Future investigation into OPC differentiation and survival rates will be critical to understand this finding and other intricacies of endogenous and therapeutic-induced remyelination more fully.

In comparing two remyelinating compounds, we observed differences in the timing of their effects on oligodendrocyte gain (**Fig. 5D-F**). LL-341070 induced a profound but transient increase in oligodendrocyte gain rate during the first half of its administration, while clemastine fumarate had a subtler impact over a longer period that continued post-administration (**Fig. 5D, E**). The enduring effect of clemastine fumarate on oligodendrocyte gain rate beyond the treatment period (**Fig. 5D**) is unlikely to be entirely explained by residual drug since clemastine fumarate is cleared fairly rapidly^66,67^. More likely, clemastine fumarate induces OPC differentiation during the treatment period, but as differentiating OPCs take several days to mature^68^, we first observe *Mobp*-EGFP expression in some cells during the post-treatment period. The temporally limited impact of LL-341070 treatment could have several explanations. First, thyroid hormone receptor beta could be inactivated with prolonged agonism. However, this is unlikely as there is sustained gene expression changes after three weeks of LL-341070 administration in rats exposed to cuprizone^54^. Second LL-341070 could restore sufficient numbers of oligodendrocytes within the first 1.5 weeks to preclude the need for more. However, we found that a drive to restore oligodendrocyte number does not control oligodendrocyte gain rate (**Fig. 2A, B**), and so enhanced restoration of the oligodendrocyte population (87±2 (SEM) % with 0.3 mg/kg LL-341070 at 1.5 weeks) should not inhibit future gain. Third, the high magnitude of differentiation induced by LL-341070 could cause a depletion in OPCs available for differentiation. Cortical OPCs maintain their density by elevating proliferation in response to differentiation events^68^ but it is currently unknown how OPC numbers and dynamics are modulated by prolonged, high magnitude differentiation rates or the duration over which such rates are sustainable. Future investigation into such questions will be important to optimize therapeutic dosing strategies for remyelination.

Visual impairments are a common manifestation of demyelinating diseases such as multiple sclerosis^69^. Here, we utilized a model with preferential cortical demyelination (**Fig. 1B, C**) and high-density extracellular *in vivo* recordings to investigate the impact of demyelination on visual neuronal function. We found that demyelination delayed the responses of individual V1 neurons to visual stimuli (**Fig. 6D**) but did not detect statistically significant changes in the latency of aggregated V1 activity (VEP N70; **Fig. 6G**). Previous studies that have found demyelination-mediated changes in VEP N70 latency have induced more severe demyelination, particularly in the anterior visual pathway; in contrast, we did not observe demyelination the optic nerve (**Fig. 1B, C**). These studies have also monitored VEP non-invasively in the same animals before and after demyelination^25,34,55^, which may be important to reduce variability given that VEPs have large inter-individual differences^70^. Though VEP N70 latency was unaltered, we did find that demyelination altered VEP shape (**Fig. 6I**), perhaps due to the added dependence of VEP shape on cortical neuron responses in addition to input from the anterior visual pathway. Intriguingly, this finding suggests that cortical demyelination may contribute to the well-described changes in VEP shape in multiple sclerosis patients^71^.

After seven weeks of endogenous or therapeutically enhanced remyelination, all groups reached a plateau of functional recovery similar to healthy levels (**Fig. 7A-D**). Furthermore, remyelination therapy accelerated this restoration of V1 neuronal function (**Fig. 6E, J**). These findings are in line with previous reports indicating that remyelination can restore demyelination-mediated deficits in stimulus-evoked neuronal latencies leading to behavioral improvements^25,34^, highlighting the imperative to bring remyelinating therapeutics to patients. Remyelination restores conduction properties^2^, synaptic transmission^72^, and neuron excitability^19,72^ disrupted by demyelination. Interestingly, we found that partial restoration of oligodendrocyte numbers was sufficient to recover neuronal function impaired by demyelination, observing no improvements in neuronal function beyond restoration of ∼87% of the oligodendrocyte population (**Fig. 7A, E and F**). New oligodendrocytes formed during remyelination myelinate both demyelinated axon segments and previously unmyelinated regions, leaving some regions demyelinated and resulting in rearrangement of the myelin landscape^19,40^. Interestingly, it is the more heavily myelinated axons that are prioritized for remyelination^13,41,73^, and these axons become remyelinated more quickly than other axons^13^. Together, these results suggest the possibility that there may be a subset of axons – those of which are heavily myelinated – whose remyelination restores neuronal function and is prioritized, such that partial restoration of oligodendrocytes is sufficient for their requisite remyelination. Future interrogation of the network implications of axon-specific demyelination and remyelination will shed light on this intriguing possibility.

## Supporting information

Source Data and Statistics

## Acknowledgements

We thank A. Chavez for assistance with genotyping and maintenance of the mouse colony, Laboratory Animal Resources at CU Anschutz for assistance with mouse drug treatments, and PW Hosokawa for statistical consulting. Michael Liebling developed the PoorMan3D registration plugin for Image Fiji.

## Author contributions

Conceptualization: GDFN, LAO, EGH, DJD, JAV, RG Investigation: GDFN (surgeries, drug treatment, in vivo imaging, immunohistochemistry, image analysis, statistical analysis), LAO (surgeries, drug treatment, in vivo imaging, image analysis, statistical analysis), DJD (electrophysiology, electrophysiology analysis), JAH (pilot experiments), AM (immunohistochemistry, image analysis), LC (in vivo imaging, image analysis), MES (surgeries, image analysis), MAT (surgeries).

Visualization: GDFN, LAO, EGH, DJD

Funding acquisition: EGH, DJD

Project administration: GDFN, LAO, EGH, DJD

Supervision: EGH, DJD

Writing – original draft: GDFN, LAO

Writing – review & editing: GDFN, LAO, JAV, DJD, EGH

## Data and material availability

All data that support the findings, tools and reagents will be shared on an unrestricted basis; requests should be directed to the corresponding authors.

## List of Supplementary Materials

Supplementary Figs. 1 to 10 Source Data

## Methods

### Animals

All animal experiments were approved by the Institutional Animal Care and Use Committee (IACUC) of the University of Colorado Anschutz Medical Campus. Animals were housed in sex-segregated individually ventilated cages, in groups up to five. Food and water were provided ad libitum. The room was kept at 72 °F ± 2 °F temperature with 50±20% humidity and was on a 14h light 10h dark cycle. The Tg(Mobp-EGFP)IN1Gsat/Mmucd mouse line (referred throughout the text as *Mobp*-EGFP) was described previously^39,74^. Mobp-EGFP mice were from a C57BL/6 genetic background. Both male and female Mobp-EGFP hemizygous mice were used for experiments, with experimental groups assigned randomly.

### Genotyping

Genotyping for Mobp-EGFP was performed from genomic DNA, using genotyping primers F: 5′ GGTTCCTCCCTCACATGCTGTTT 3′ and R: 5′ TAGCGGCTGAAGCACTGCA 3′ and PCR conditions of 94 °C for 4 min, [94 °C for 30 s, 73-68 °C (0.5 °C decrease per cycle) for 30 s and 72 °C for 30 sec] for 10 cycles, (94 °C for 30 s, 68 °C for 30 s and 72 °C for 30 sec) for 28 cycles, and 72 °C for 7 min, yielding a ∼300 bp band.

### Cranial window implantation surgery

6-8-week-old mice were anesthetized with isoflurane inhalation (induction, 5%; maintenance, 1.5–2.0%, mixed with 0.5 L/ min O_2_) and kept at 37 °C body temperature with a thermostat-controlled heating plate. After removal of the skin over the right cerebral hemisphere, the skull was cleaned and a 2 × 2 mm region of skull centered over the primary visual cortex (1.5 mm anterior to lambda– 0.5 mm posterior to lambda and 1.5–3.5 mm lateral to bregma) was removed using a high-speed dental drill. A piece of cover glass (VWR, No. 1) was then placed in the craniotomy and sealed with Vetbond (3M) and then dental cement (C&B Metabond). A 5 mg/kg dose of carprofen was subcutaneously administered before awakening and for three additional days for analgesia. For head stabilization, a custom metal plate with a central hole was attached to the skull.

### Remyelination therapies

LL-341070 was provided by Autobahn Therapeutics as a solution in 1% 1-methyl-2-pyrrolidone (Sigma 270458) and 1% Kolliphor HS 15 (Sigma 42966) to a final concentration of 20 □g/ml or 60 □g/mL. Clemastine fumarate (Tocris 1453) was solubilized in a 10% DMSO solution to a final concentration of 1 mg/mL. Remyelination therapies or respective vehicles were administered once daily for three weeks by gavage at 10 mL/kg for clemastine and 5 mL/kg for LL-341070, resulting in doses of 0.1 mg/kg or 0.3 mg/kg for LL-341070 and 10 mg/kg for clemastine. During drug administration, experimenters were blinded to the experimental groups. Mice treated with LL-341070 and clemastine or their respective vehicles were run in simultaneous cohorts by the same experimenters and in some instances were run in the same cohort. Since we did not observe differences between untreated and vehicle-treated mice (**Supplementary Fig. 5**), these mice were combined to study endogenous remyelination (**Fig. 1, 2**).

### Cuprizone-mediated demyelination

Demyelination was induced in 9-10-week-old mice using 0.2% Bis(cyclohexanone)oxaldihydrazone (cuprizone; Sigma C9012 or Sigma 14690), stored in a glass desiccator at 4 °C. Cuprizone was added to powdered chow (Envigo T.2918M.15), mixed for ∼10 minutes to ensure homogeneity, and provided to mice in custom feeders (designed to minimize exposure to moisture) for 24-25 days on an ad libitum basis. Feeders were refilled every 2–3 days, and fresh cuprizone chow was prepared weekly. Healthy mice received normal powdered chow in identical feeders. Regular chow was removed and powdered chow was introduced 2-3 days before cuprizone to ensure acclimation to the powdered food. Cages were changed weekly to avoid build-up of cuprizone chow in bedding, and to minimize reuptake of cuprizone chow following cessation of diet via coprophagia.

### Immunofluorescence

Mice were anesthetized with an intraperitoneal injection of sodium pentobarbital (100 mg per kg body weight) and transcardially perfused with 4% paraformaldehyde in 0.1 M phosphate buffer (pH 7.0–7.4). Optic nerves were postfixed in 4% paraformaldehyde overnight at 4 °C, transferred to 30% sucrose solution in PBS (pH 7.4) and stored at 4 °C until processing. Tissue was positioned in cryomolds with Tissue-Tek OCT, frozen, and 10-μm thick sections were obtained using a cryostat. Immunostaining was performed with sections on a slide. Sections were first permeabilized with acetone for 5 min, preincubated in blocking solution (5% normal donkey serum, 0.3% Triton X-100 in PBS, pH 7.4) for 1h at room temperature, then incubated overnight at 4 °C in primary antibody. Primary antibodies were Rabbit anti-Caspr (Abcam ab34151) and Chicken anti-pan-Nfsc (R&D Biosystems AF3235), both diluted 1:500 in blocking buffer. Secondary antibody incubation was performed at room temperature for 1 h. Secondary antibodies were Alexa Fluor® 488 Donkey anti-chicken IgY (Jackson Immunoresearch Laboratories 703-545-155) and Alexa Fluor™ 568 Donkey anti-rabbit IgG (Thermo Fischer A10042). Sections were mounted on slides with Vectashield antifade reagent (Vector Laboratories). Images were acquired using a laser-scanning confocal microscope (Leica SP8) at a magnification of 63X.

### Two-photon imaging

In vivo imaging sessions began three weeks post-surgery and took place 1-3 times per week (Fig. 3B). During imaging sessions, mice were anesthetized with isoflurane and immobilized by attaching the head plate to a custom stage. Images were collected using a Zeiss LSM 7MP microscope equipped with a BiG GaAsP detector using a mode-locked Ti:sapphire laser (Coherent Ultra) tuned to 920 nm. The average power at the sample during imaging was 5-30 mW. Vascular and cellular landmarks were used to identify the same cortical region over longitudinal imaging sessions. MOBP–EGFP image stacks were acquired using a Zeiss W plan-apochromat ×20/1.0 NA water immersion objective giving a volume of 425 μm × 425 μm × 336 μm (1,024 × 1,024 pixels; corresponding to layers I-III, 0–336 μm from the meninges) from the cortical surface.

### Image processing and analysis

Image stacks and time series were analyzed using ImageJ Fiji ^75^. For quantification of nodes of Ranvier, three fields of view of 10000 7m were analyzed per animal. Longitudinal image stacks were registered using the Fiji plugins Correct 3D drift ^76^ and PoorMan3DReg. Oligodendrocyte cell tracking was performed manually. A custom Fiji script ^39^ enabled recording of oligodendrocyte state (new, lost, or stable EGFP+ soma) across timepoints. Oligodendrocyte gain and loss are reported relative to the baseline cell population to account for variations in baseline cell number. The rate of oligodendrocyte gain was quantified as the percentage change in gain over the amount of time elapsed. During data analysis, experimenters were blinded to the experimental groups. For presentation in figures, image brightness and contrast levels were adjusted for clarity.

### Visual stimulation

Stimuli were displayed on the inner surface of a custom-designed spherical dome enclosure (**Supplementary Fig. 7A**). The enclosure was a clear 24” diameter acrylic sphere (California Quality Plastics, Ontario, CA) coated with a custom UV-reflective paint. The paint had 50% absorbance to reduce interference caused by internal reflections and reflection was spectrally flat from 350 – 600 nm (Twilight Labs Inc., Grand Forks, ND). The animal was placed on a horizontal disc suspended near the center of the sphere, head-fixed such that it could run freely, and positioned with the right eye at the center of the sphere. This created a spherical coordinate system with the viewing eye at the origin; based on previous measurements the optical axis projects to 22º in elevation and 60º in azimuth in the dome coordinate system. A custom digital light processing (DLP) projector (DLi Innovations, Austin, TX) with mouse photoreceptor-optimized LED sources placed outside of the enclosure projected, at a refresh rate of 60 Hz, through a small opening in the dome onto a silver-coated aluminum mirror (Wagner Collaborative Metal Works, Milwaukee, WI) suspended beneath the mouse platform.

Visual stimuli covered 200º in azimuth and 100º in elevation. Visual stimuli were generated on a PC using custom written software based on the PsychoPy (http://www.psychopy.org) platform. Flashed stimuli consisted of 50 msec temporary increases (bright flash) or decreases (dark flash) in full-field luminance, ∼1.8 cd/m^2^ from a mean luminance of 3.6 cd/m^2^.

### *In vivo* electrophysiology

To record from neuronal populations across primary visual cortex (V1) we used three Neuropixels^77^ probes simultaneously. These are high-density electrophysiology probes with 384 contiguous channels on a single shank covering 3.84 mm. Immediately before the recording, the animals were anesthetized, the cranial window was removed, and a partial durectomy was performed to allow for probe insertion. After head-fixation inside a visual stimulus environment, probes were inserted through the opening in the skull. Recordings were grounded to the animal. Following insertion, the exposed skull was covered with 1-2% agar to improve recording stability and maintain brain health. For all recordings, several hundred channels were left outside of the brain but within the covering agar to facilitate data-driven probe localization. The probe was allowed to settle for at least 15, but typically 30 minutes, before responses were recorded.

Each probe averaged 196 +/- 165 (std. dev) visual cortical neurons per recording. A post-hoc assignment of both unit depth and cortical layer was made by taking the distance between the center of the waveform amplitude distribution along the shank and the channel estimated at just beyond the pial surface. Neurons were recorded from all cortical layers, though distributions were biased towards deeper regions.

### *In vivo* electrophysiology: spike sorting and pre-processing

Electrophysiological data from each channel was recorded in two bands: a “spike” band sampled at 30kHz and hardware filtered at 300 – 10000Hz, and a “field potential” band sampled at 2.5kHz and hardware filtered at 0.1 – 500 Hz (**Supplementary Fig. 7C**).

To recover the spike times from individual neurons recorded in the spike band, we used a workflow that included algorithmic and manual refinement steps. We began with spike time extraction and putative single unit isolation using Kilosort, and manually refined the results using phy (github.com/cortexlab/phy). During manual refinement, most decisions were merging of clusters that had been algorithmically split due to drift in amplitude on the timescale of the entire recording; in addition, the first spikes in a burst also needed to be manually merged. Each unit was tagged as multi-unit (MUA) or single-unit based on the waveform and shapes of neighboring waveforms and the middle few milliseconds of the spike time auto-correlograms. For each recording, the quality of isolation of each unit was quantified using several metrics: maximum signal-to-noise, purity of the auto-correlogram, and distance of the mean waveform from other units in both feature and waveform space. These metrics were combined into a single quality metric using a Fisher’s linear discriminant projection based on the manual labels. Finally, the depth of each unit from the pial surface was calculated based on features of the neural data. We calculated the power in a range of frequency bands for each channel for a short epoch. The distribution of several bands revealed features of the underlying structure; from the alpha-band (8-14 Hz), we were able to identify the first channel above the pial surface. This channel served as the reference channel for measuring the depth of each unit. The center of each unit was calculated based on the mean waveform, weighted by the amplitude of the waveform on each channel. Using the spike trains from these large populations of single neurons, we compare latencies to visual responses between conditions.

To analyze visually evoked potentials, we computed average potentials on each of the hundreds of electrodes in the visual cortex. Unfiltered local field potentials were trial-averaged to generate evoked potentials; a linear sub-array of channels corresponding to those in visual cortex was selected from the recorded data. Since visual evoked potentials (VEPs) are usually recorded non-invasively from scalp electrodes placed over the occipital region, electrical activity from superficial cortex is overrepresented in these recordings. To match this effect for comparison to clinical VEPs, we applied a log-weight according to depth of the electrode and summed all the evoked potentials. Because nominal amplitude values can be affected by the degree of tissue damage caused by probe placement, as well as the geometry of the electrode relative to large potential sources such as the apical dendrites of large neurons, VEP amplitude data was normalized by the standard deviation of VEP signals preceding the stimuli for each Neuropixels probe.

To calculate the shape distortion metric, we first averaged the computed VEP of healthy animals to create a VEP shape template. We then calculated a point-by-point spatial distance correlation between the conditional VEPs and the VEP shape template (scipy.spatial.distance.correlation). Since this is a comparative measure that is bounded between 0 and 2, VEP responses to bright and dark flash were considered to be independent and were analyzed together.

The VEP signal was manually inspected by an investigator blinded to the experimental conditions, and signals deemed to have low quality were excluded from further analyses.

### Statistics and modeling

Summarized statistical information is present in the legend of each figure. Detailed statistical information is in Table S1. Sample sizes were not predetermined using statistical methods, but were comparable to relevant publications ^19,39^. Experimental groups were replicated in multiple cohorts with multiple experimental groups per cohort. Statistical analyses were conducted using JMP Pro 16 (SAS). Normality was assessed in all datasets using the Shapiro–Wilk test. Normality was satisfied in all longitudinal imaging experiments, so parametric statistics were used. To compare group means, paired two-tailed Student’s *t* test or ANOVA with Tukey’s post hoc were used. To compare variance between different groups, the Brown– Forsythe test was used. To investigate the relationship between two variables, simple linear regression was used; the coefficient of determination (r^2^) and the p-value of an ANOVA testing the null-hypothesis that there is no linear relationship between the tested variables were reported. Given that oligodendrocyte gain depends on the level of oligodendrocyte loss, we used an analysis of covariance with unequal slopes to take this relationship into account when testing for drug effects. We used a standard least square personality of the model, with oligodendrocyte gain (or gain rate) as the independent variable, and treatment, oligodendrocyte loss (or loss rate), and loss by treatment interaction as model effects. We reported the least square means adjusted for the average oligodendrocyte loss (or loss rate) unless otherwise specified. An ANOVA tested for the effects of oligodendrocyte loss (or loss rate), treatment, and their interaction. Treatment effects were further investigated using Tukey’s HSD post-hoc tests when significance was found with ANOVA.

For temporal analysis of oligodendrocyte gain rates (in Figs. 3J and 5D), oligodendrocyte gain rates were determined during specific time windows by dividing the number of new oligodendrocytes by the number of days. To model the continuous dynamics of oligodendrocyte gain or loss, we used Gompertz three-parameter modeling:

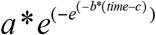

where:

a = asymptote

b = curve’s growth rate

c = inflection point

For Fig. 2, we used a quintic polynomial curve to model the variation in oligodendrocyte population (% of baseline) throughout demyelination and remyelination.

Oligodendrocyte gain and loss rates in Fig. 2 were obtained from the first derivative of the modeled dynamics of oligodendrocyte gain or loss. Maximum oligodendrocyte gain rate was the oligodendrocyte gain rate at the inflection point of the modeled oligodendrocyte gain curve.

For the electrophysiology results, normality was violated; therefore, we used nonparametric tests: Wilcoxon signed-rank test for comparisons between two groups and Kruskal-Wallis test followed by Steel-Dwass post-hoc tests to compare more than two groups.

For data visualization, all error bars represent the standard error of the mean, all bar graphs denote means and all box plots illustrate medians and IQRs unless otherwise specified. Significance was determined by P < 0.05, except when a Bonferroni correction was used for multiple comparisons (indicated in figure legends).

