## Supplemental Figures for "Incomplete remyelination via endogenous or therapeutically enhanced oligodendrogenesis is sufficient to recover visual cortical function"

**­­**

**Supplementary Figures**

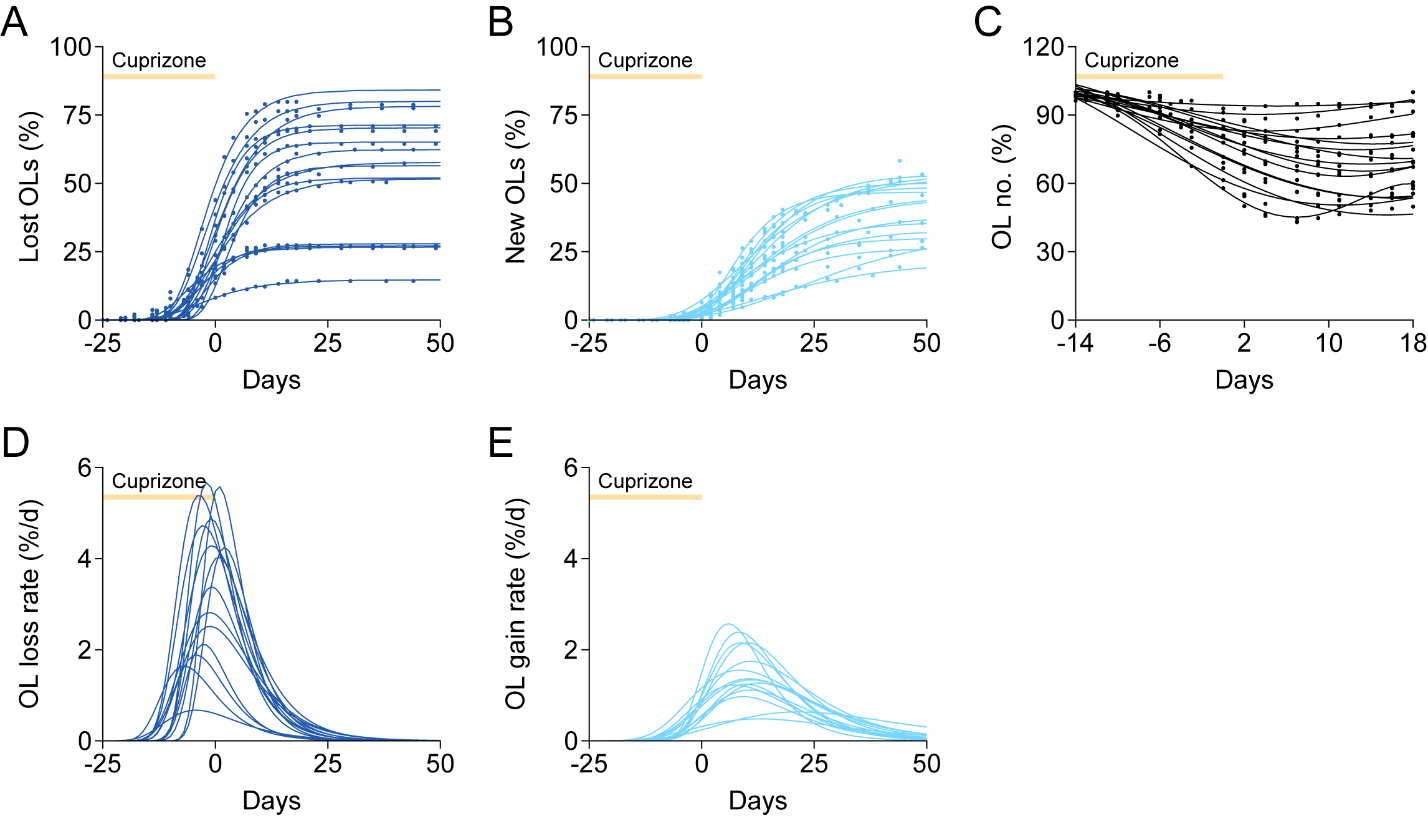

**Supplementary Fig.1. Modeling oligodendrocyte dynamics over time.**

**(A** to **B)** Three parameter Gompertz growth curves fit to cumulative OL loss (A) and gain (B) over time for individual mice. R^2^ ≥ 0.98 for all mice. Only mice with imaging data out to at least 3 weeks post-cuprizone were used in growth curve modeling (n = 15).

**(C)** Quintic curves fit to OL number over time for individual mice. R^2^ ≥ 0.86 for all mice. Only mice with imaging data out to at least 3 weeks post-cuprizone were used in quintic curve modeling (n = 15). Modeling tracked the data most accurately between -14 days and 18 days so modeled values were used from only this period.

**(D** to **E)** First derivatives of three parameter Gompertz growth curves for cumulative OL loss (D) and gain (E) for individual mice (n = 15) represent OL loss (D) and gain (E) rates over time.

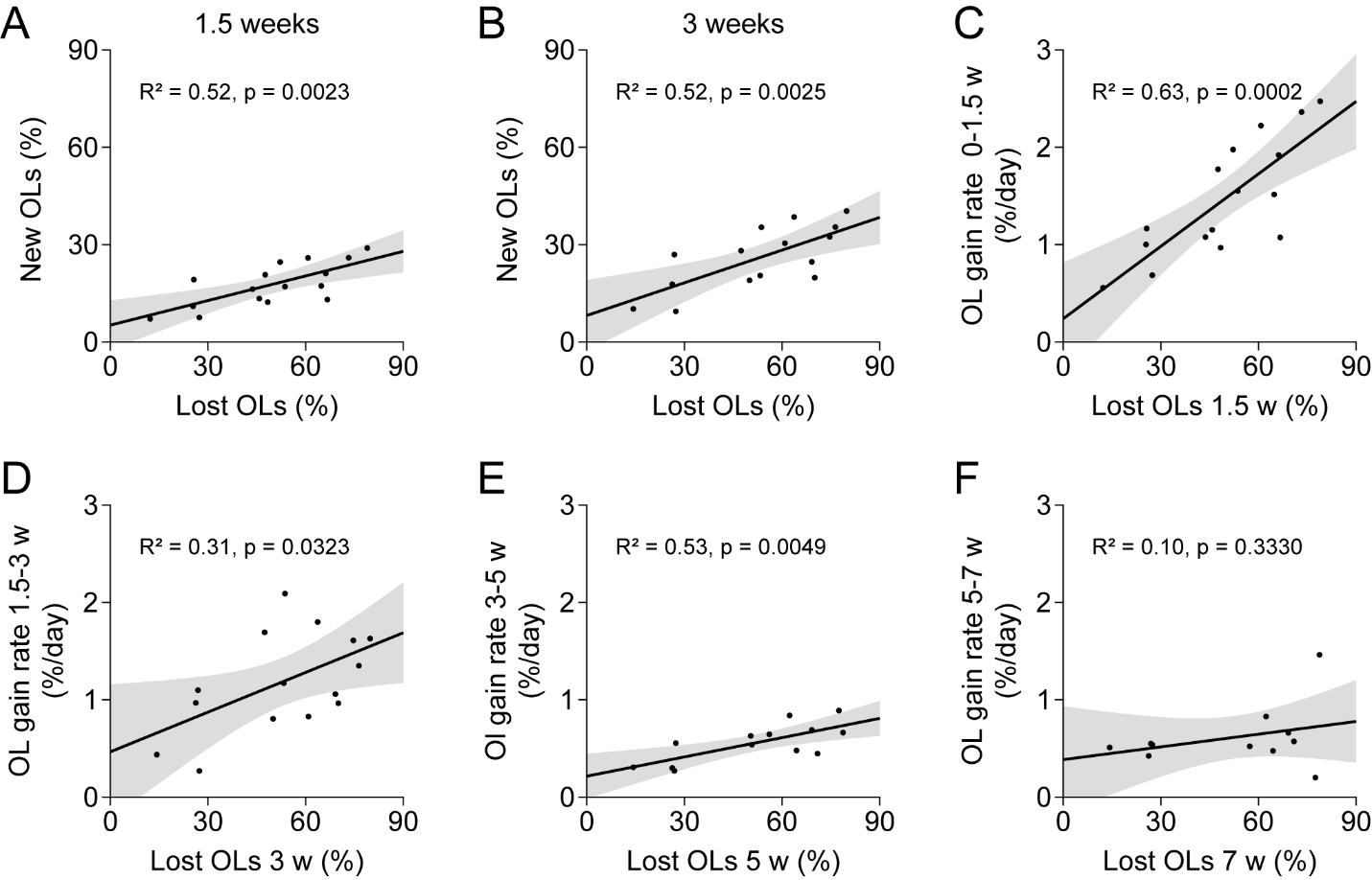

**Supplementary Fig.2. Oligodendrocyte gain magnitude and rate correlate with oligodendrocyte loss magnitude.**

**(A)** Magnitude of OL gain by 1.5 weeks is correlated with magnitude of OL loss by 1.5 weeks (n = 16).

**(B)** Magnitude of OL gain by 3 weeks is correlated with magnitude of OL loss by 3 weeks (n = 15).

**(C)** OL gain rate from 0 to 1.5 weeks is correlated with magnitude of OL loss by 1.5 weeks (n = 16).

**(D)** OL gain rate from 1.5 to 3 weeks is correlated with magnitude of OL loss by 3 weeks (n = 15).

**(E)** OL gain rate from 3 to 5 weeks is correlated with magnitude of OL loss by 5 weeks (n = 13).

**(F)** OL gain rate from 5 to 7 weeks does not correlate with magnitude of OL loss by 7 weeks (n = 11).

Linear regression with 95% CI.

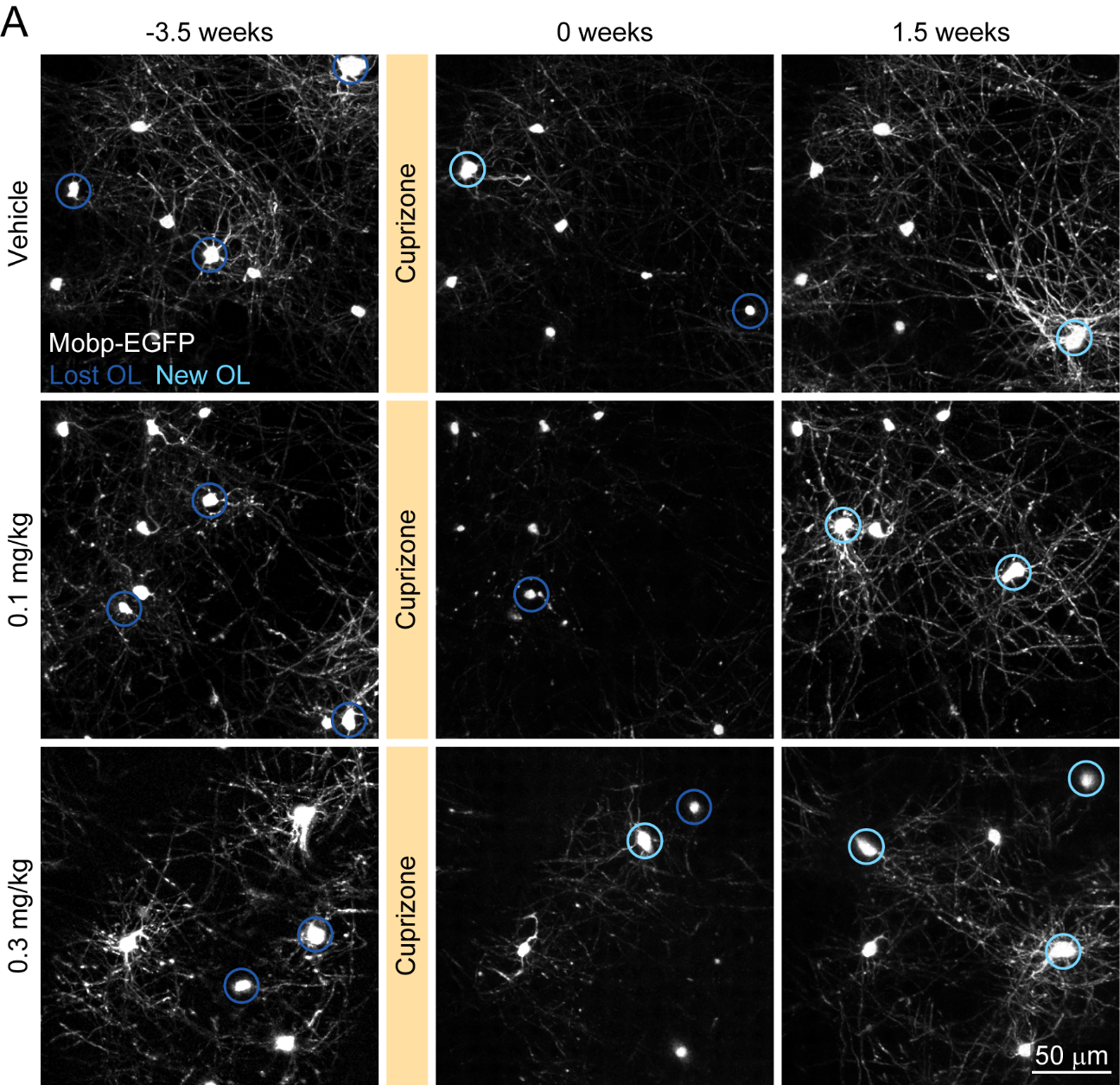

**Supplementary Fig.3. Representative images of V1 oligodendrocytes in thyromimetic- and vehicle-treated mice.**

**(A)** Representative images of V1 OLs in mice treated with vehicle, 0.1 mg/kg LL-341070, or 0.3 mg/kg LL-341070 at baseline (-3.5 weeks), at the end of cuprizone (0 weeks), and at 1.5 weeks post-cuprizone. Lost OLs (dark blue) and new OLs (light blue) are encircled.

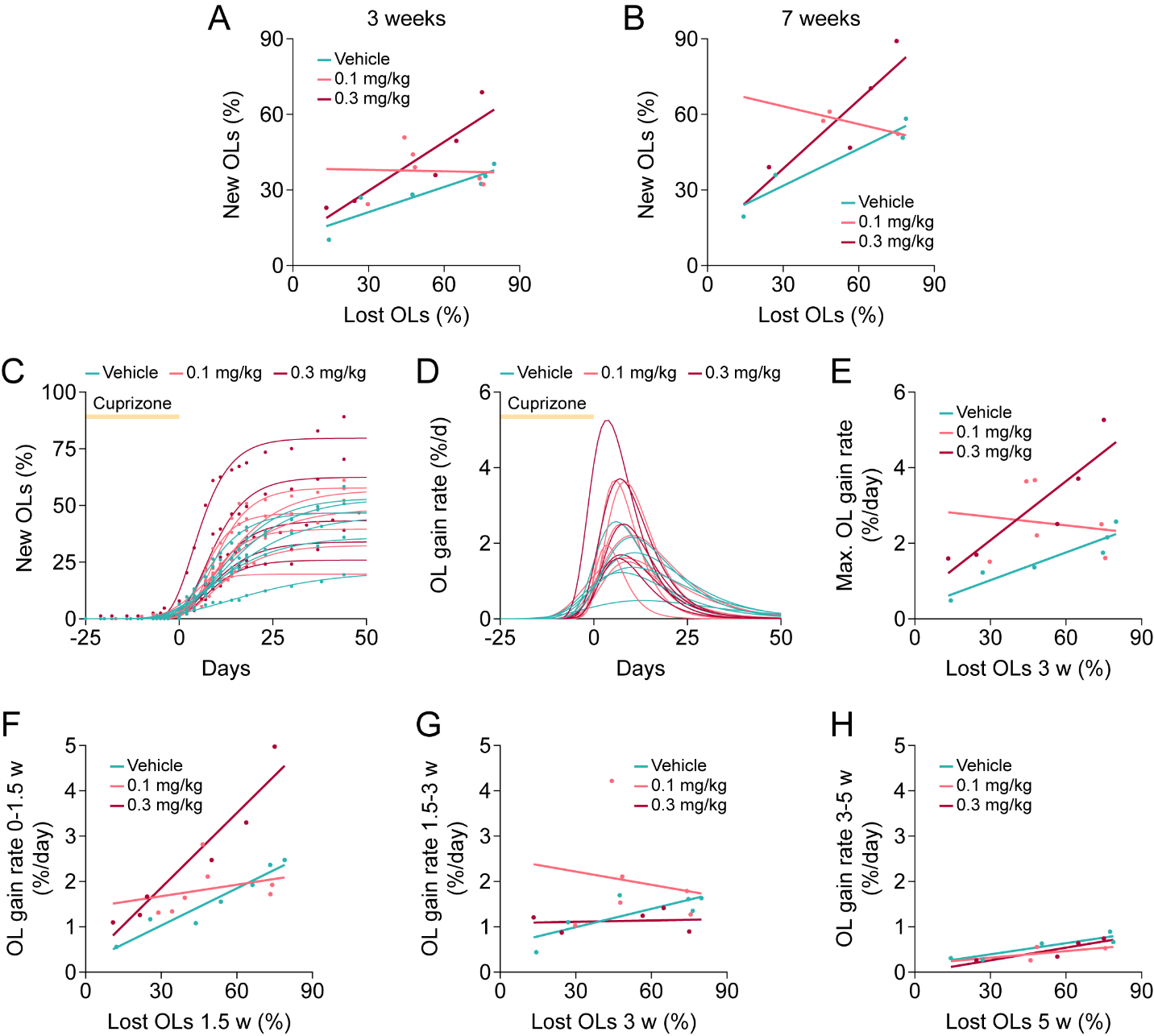

**Supplementary Fig.4. Supporting evidence that thyromimetic treatment enhances oligodendrocyte gain during remyelination.**

**(A)** Multiple linear regression model shows effect of treatment and OL loss but not the interaction between OL loss and treatment on OL gain at 3 weeks.

**(B)** Multiple linear regression model shows no effect of treatment, OL loss, or interaction between OL loss and treatment on OL gain at 7 weeks.

**(C)** Three parameter Gompertz growth curves fit to cumulative OL gain over time for individual mice. R^2^ > 0.97 for all mice. Only mice with imaging data out to at least 3 weeks post-cuprizone were used in growth curve modeling (vehicle: n = 6, 0.1 mg/kg: n = 6, 0.3 mg/kg: n = 5).

**(D)** First derivatives of three parameter Gompertz growth curves for cumulative OL gain for individual mice represent OL gain rates over time (vehicle: n = 6, 0.1 mg/kg: n = 6, 0.3 mg/kg: n = 5) and were used to derive maximum OL gain rate.

**(E)** Multiple linear regression model shows effect of treatment and OL loss but not the interaction between OL loss and treatment on maximum OL gain rate.

**(F)** Multiple linear regression model shows effect of treatment , OL loss, and interaction between OL loss and treatment on OL gain rate between 0 and 1.5 weeks

**(G)** No effect detected in multiple linear regression model for effect of treatment, OL loss, and interaction between treatment and OL loss on OL gain rate between 1.5 and 3 weeks.

**(H)** No effect detected in multiple linear regression model for effect of treatment, OL loss, and interaction between treatment and OL loss on OL gain rate between 3 and 5 weeks.

In **A**, Model (F(5, 11) = 5.77, ** p = 0.0075, ANOVA). Effects: treatment (F(2) = 4.5174, * p = 0.037), OL loss (F(1) = 10.68, ** p = 0.0075), interaction (F(2) = 3.31, p = 0.075). Vehicle: n = 6, 0.1 mg/kg: n = 6, 0.3 mg/kg: n = 5.

In **B**, Model (F(5, 5) = 6.27, * p = 0.033, ANOVA). Effects: treatment (F(2) = 3.37, p = 0.12), OL loss (F(1) = 5.09, p = 0.074), interaction (F(2) = 2.84, p = 0.15). Vehicle: n = 4, 0.1 mg/kg: n = 3, 0.3 mg/kg: n = 4.

In **E**, Model (F(5, 11) = 5.22, * p = 0.011, ANOVA). Effects: treatment (F(2) = 6.26, * p = 0.015), OL loss (F(1) = 6.54, * p = 0.027), interaction (F(2) = 3.12, p = 0.085). Vehicle: n = 6, 0.1 mg/kg: n = 6, 0.3 mg/kg: n = 5.

In **F**, Model (F(5, 14) = 16.42, **** p < 0.0001, ANOVA). Effects: treatment (F(2) = 14.82, *** p = 0.0003), OL loss (F(1) = 40.1741, **** p < 0.0001), interaction (F(2) = 7.64, ** p = 0.0057). Vehicle: n = 7, 0.1 mg/kg: n = 7, 0.3 mg/kg: n = 6.

In **G**, Model (F(5, 11) = 1, p = 0.46, ANOVA). Vehicle: n = 6, 0.1 mg/kg: n = 6, 0.3 mg/kg: n = 5.

In **H**, Model (F(5, 6) = 3.82, p = 0.067, ANOVA). Vehicle: n = 5, 0.1 mg/kg: n = 3, 0.3 mg/kg: n = 4.

* p < 0.05, ** p < 0.01, *** p < 0.001, **** p < 0.0001, least square mean ± SEM.

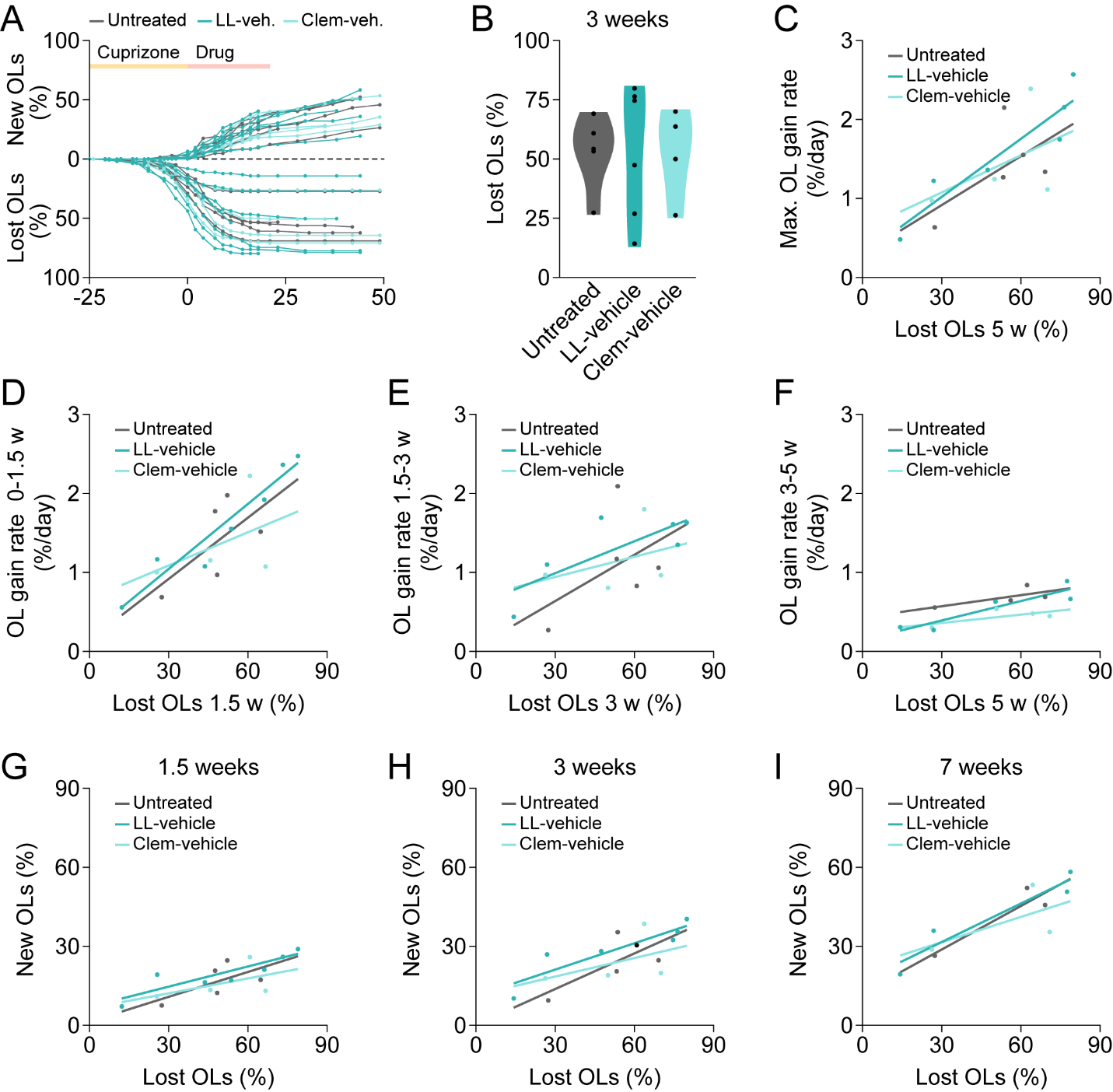

**Supplementary Fig.5. Magnitude and dynamics of oligodendrocyte loss and gain are comparable between untreated and vehicle-treated mice.**

**(A)** Cumulative OL loss and gain as a percentage of baseline OLs in individual mice over time (untreated: n = 5, LL-341070 vehicle: n = 7, clemastine vehicle: n = 4).

**(B)** OL loss by 3 weeks did not differ between groups in mean or variance.

**(C)** No effect detected in multiple linear regression model for effect of treatment, OL loss, and interaction between treatment and OL loss on maximum OL gain rate.

**(D)** Multiple linear regression model shows effect of OL loss but not treatment or the interaction between OL loss and treatment on OL gain rate from 0-1.5 weeks

**(E)** No effect detected in multiple linear regression model for effect of treatment, OL loss, and interaction between treatment and OL loss on OL gain rate between 1.5 and 3 weeks **(F)** Multiple linear regression model shows effect of shows effect of treatment and OL loss but no effect of interaction between OL loss and treatment on OL gain rate from 3-5 weeks. Difference between untreated and clemastine vehicle.

**(G)** No effect detected in multiple linear regression model for effect of treatment, OL loss, and interaction between treatment and OL loss on OL gain at 1.5 weeks.

**(H)** No effect detected in multiple linear regression model for effect of treatment, OL loss, and interaction between treatment and OL loss on OL gain at 3 weeks

**(I)** No effect detected in multiple linear regression model for effect of treatment, OL loss, and interaction between treatment and OL loss on OL gain at 7 weeks

In **B**, Mean (untreated: 52.86 ± 10.04, n = 5; LL-341070 vehicle: 53.24 ± 9.168, n = 6; clemastine vehicle: 52.20 ± 11.23, n = 4; F(2, 12) = 0.0013, p = 1, ANOVA). Variance (F(2, 12) = 1.82, p = 0.2, Brown-Forsythe).

In **C**, Model (F(5, 9) = 2.72, p = 0.092, ANOVA). Untreated: n = 5, LL-341070 vehicle: n = 6, clemastine vehicle: n = 4.

In **D**, Model (F(5, 10) = 4.21, * p = 0.026, ANOVA). Effects: treatment (F(2) = 0.33, p = 0.72), OL loss (F(1) = 9.93, * p = 0.01), interaction (F(2) = 0.41, p = 0.67). Vehicle: Untreated: n = 5, LL-341070 vehicle: n = 7, clemastine vehicle: n = 4.

In **E**, Model (F(5, 9) = 1.017, p = 0.4612, ANOVA). Untreated: n = 5, LL-341070 vehicle: n = 6, clemastine vehicle: n = 4.

In **F**, Model (F(5, 7) = 6.57, * p = 0.0141, ANOVA). Effects: treatment (F(2) = 4.83, * p = 0.048), OL loss (F(1) = 10.83, * p = 0.013), interaction (F(2) = 1.016, p = 0.41). Post hoc (untreated: 0.67 ± 0.05, n = 4; LL-341070 vehicle: 0.57 ± 0.05, n = 5; clemastine vehicle: 0.44 ± 0.05, n = 4; untreated versus LL-341070 vehicle p = 0.38, untreated versus clemastine vehicle * p = 0.041, LL-341070 vehicle versus clemastine vehicle p = 0.2223, Tukey HSD).

In **G**, Model (F(5, 10) = 2.75, p = 0.082, ANOVA). Untreated: n = 5, LL-341070 vehicle: n = 7, clemastine vehicle: n = 4.

In **H**, Model (F(5, 9) = 2.69, p = 0.0933, ANOVA). Untreated: n = 5, LL-341070 vehicle: n = 6, clemastine vehicle: n = 4.

In **I**, Model (F(5, 4) = 2.96, p = 0.1580, ANOVA). Untreated: n = 3, LL-341070 vehicle: n = 4, clemastine vehicle: n = 3.

* p < 0.05, least square mean ± SEM.

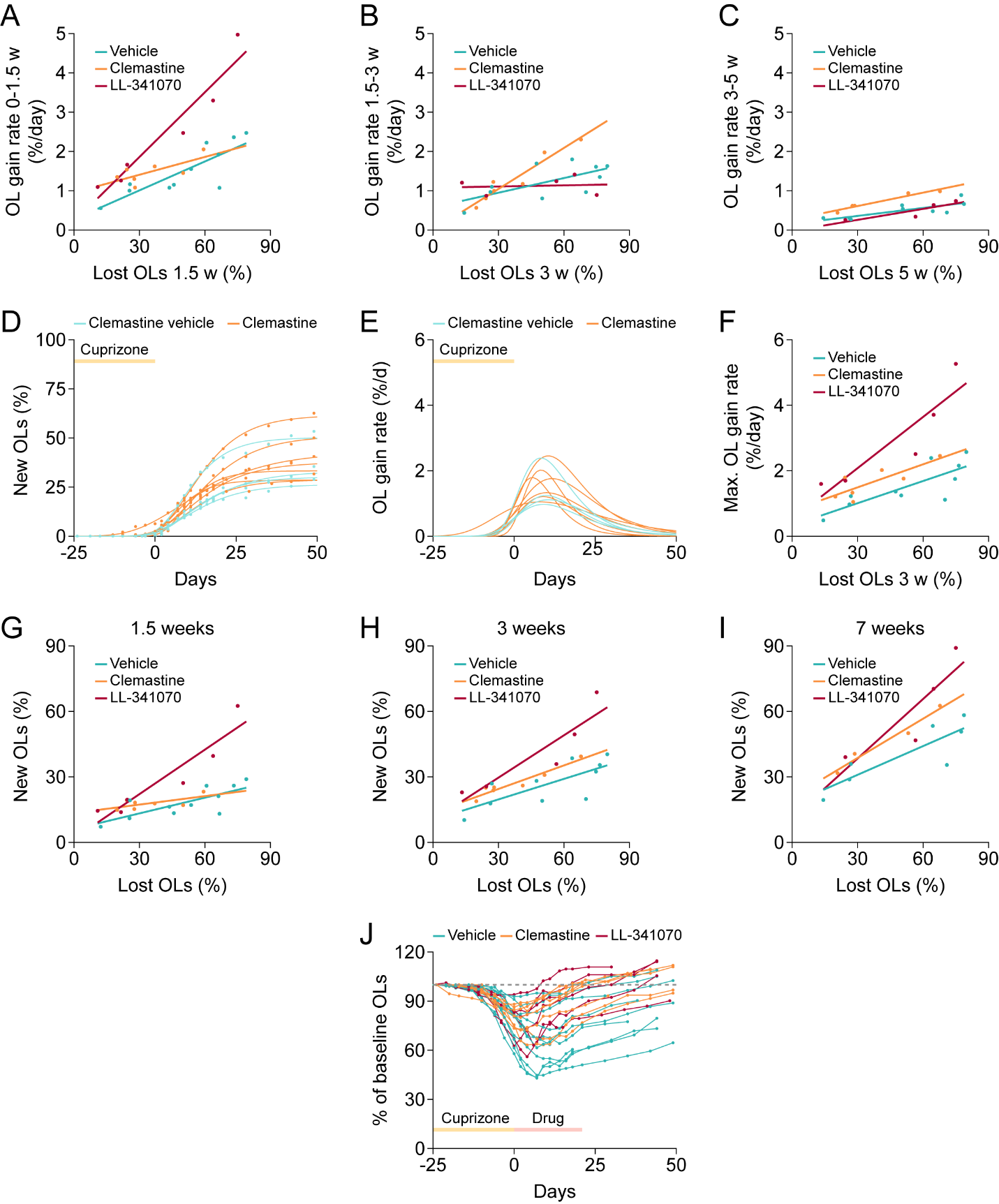

**Supplementary Fig.6. Supporting evidence that LL-341070 enhances oligodendrocyte gain during remyelination more quickly and robustly than clemastine fumarate**

**(A)** Multiple linear regression model shows effect of treatment, OL loss, and interaction between OL loss and treatment on OL gain rate from 0-1.5 weeks.

**(B)** Multiple linear regression model shows effect of treatment, OL loss, and interaction between OL loss and treatment on OL gain rate from 1.5-3 weeks.

**(C)** Multiple linear regression model shows effect of treatment and OL loss but no effect of interaction between OL loss and treatment on OL gain rate from 3-5 weeks.

**(D)** Three parameter Gompertz growth curves fit to cumulative OL gain over time for individual mice. R^2^ > 0.98 for all mice. Only mice with imaging data out to at least 3 weeks post-cuprizone were used in growth curve modeling (clemastine vehicle: n = 4, clemastine: n = 7).

**(E)** First derivatives of three parameter Gompertz growth curves for cumulative OL gain for individual mice represent OL gain rates over time (clemastine vehicle: n = 4, clemastine: n = 7).

**(F)** Multiple linear regression model shows effect of treatment, OL loss, and interaction between OL loss and treatment on maximum OL gain rate.

**(G)** Multiple linear regression model shows effect of treatment, OL loss, and interaction between OL loss and treatment on OL gain rate at 1.5 weeks.

**(H)** Multiple linear regression model shows effect of treatment, OL loss, but not the interaction between OL loss and treatment on OL gain at 3 weeks

**(I)** Multiple linear regression model shows effect of treatment and OL loss but no effect of interaction between OL loss and treatment on OL gain at 7 weeks.

**(J)** OL number as a percentage of baseline OLs in individual mice over time (vehicle: n = 7, clemastine: n = 4, LL-341070: n = 4). Dashed line at 100%.

In **A**, Model (F(5, 18) = 25.14, **** p < 0.0001, ANOVA). Effects: treatment (F(2) = 21.97, **** p < 0.0001), OL loss (F(1) = 50.036, **** p < 0.0001), interaction (F(2) = 8.46, ** p = 0.0026). Vehicle: n = 11, clemastine: n = 7, LL-341070: n = 6.

In **B**, Model (F(5, 16) = 6.7, ** p = 0.0015, ANOVA). Effects: treatment (F(2) = 4.78, * p = 0.024), OL loss (F(1) = 23.15, *** p = 0.0002), interaction (F(2) = 6.96, ** p = 0.0067). Vehicle: n = 10, clemastine: n = 7, LL-341070: n = 5.

In **C**, Model (F(5, 12) = 10.071, *** p = 0.0006, ANOVA). Effects: treatment (F(2) = 13.4, *** p = 0.0009), OL loss (F(1) = 34.35, *** p < 0.0001), interaction (F(2) = 0.96, p = 0.41). Vehicle: n = 9, clemastine: n = 5, LL-341070: n = 4.

In **F**, Model (F(5, 21) = 15.93, **** p < 0.0001, ANOVA). Effects: treatment (F(2) = 16.82, *** p = 0.0001), OL loss (F(1) = 37.12, *** p < 0.0001), interaction (F(2) =3.69, ** p = 0.048). Vehicle: n = 10, clemastine: n = 7, LL-341070: n = 5.

In **G**, Model (F(5, 18) = 18.77, **** p < 0.0001, ANOVA). Effects: treatment (F(2) = 16.38, **** p < 0.0001), OL loss (F(1) = 32.91, **** p < 0.0001), interaction (F(2) = 9.13, ** p = 0.0018). Vehicle: n = 11, clemastine: n = 7, LL-341070: n = 6.

In **H**, Model (F(5, 16) = 12.71, **** p < 0.0001, ANOVA). Effects: treatment (F(2) = 9.39, ** p = 0.002), OL loss (F(1) = 37.93, **** p < 0.0001), interaction (F(2) = 2.54, p = 0.1103). Vehicle: n = 10, clemastine: n = 7, LL-341070: n = 5.

In **I**, Model (F(5, 9) = 9.19, ** p = 0.0024, ANOVA). Effects: treatment (F(2) = 4.71, * p = 0.04), OL loss (F(1) = 28.94, *** p = 0.0004), interaction (F(2) = 1.51, p = 0.27). Vehicle: n = 7, clemastine: n = 4, LL-341070: n = 4.

* p < 0.05, ** p < 0.01, *** p < 0.001, **** p < 0.0001, least square mean ± SEM.

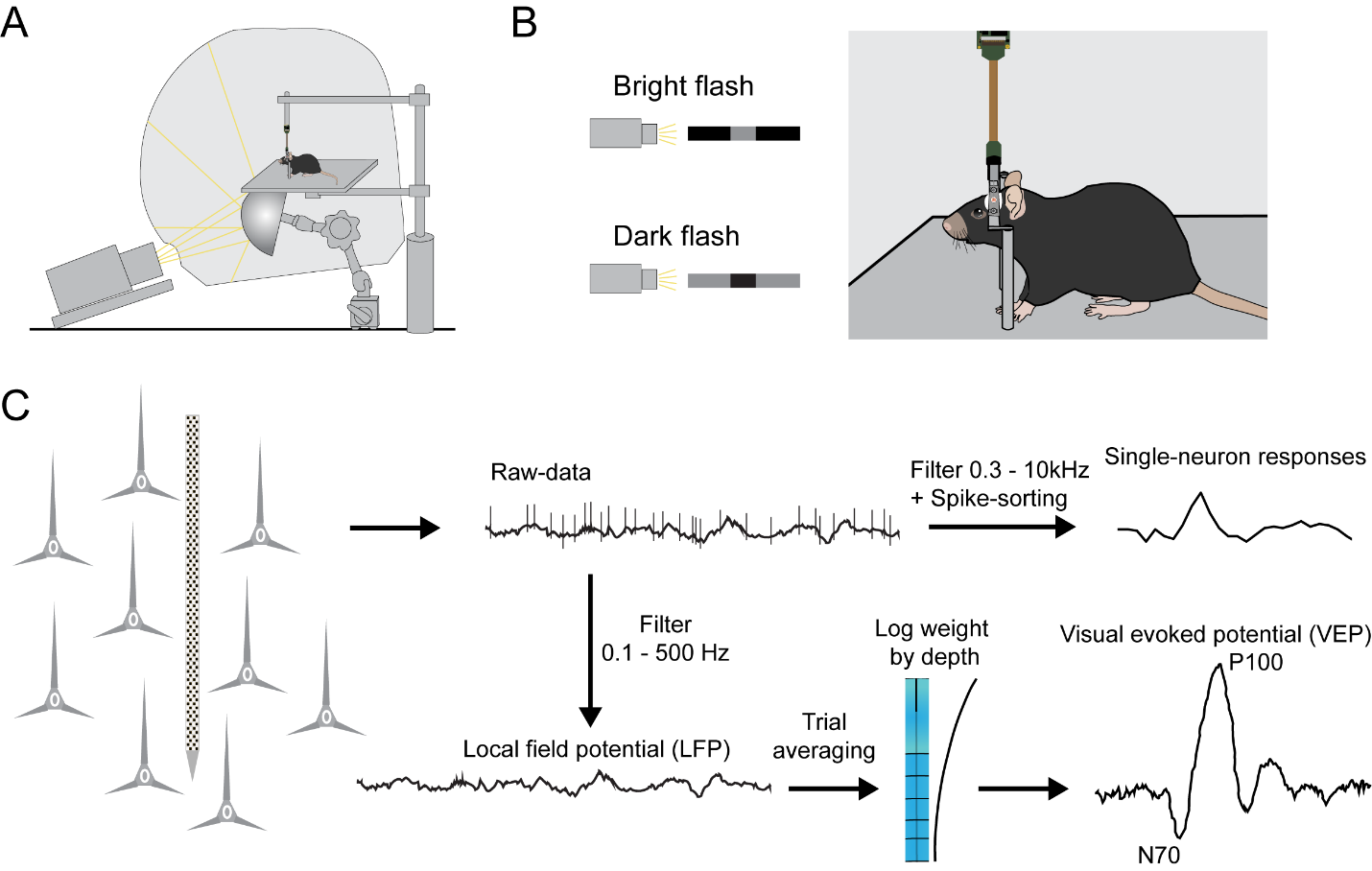

**Supplementary Fig.7. Neuronal recordings using *Neuropixel* probes**

**(A)** Schematic representation of the immersive visual stimulation dome.

**(B)** During the recording, mice are presented with a bright flash stimulus (50 ms increase in luminance) and a dark flash stimulus (50 ms decrement in luminance).

**(C)** Two signals can be recovered from raw *Neuropixels* recordings: single-neuron responses are identified through spike sorting (top), while visual evoked responses (VEPs) are obtained from a sum of trial-averaged local field potentials weighted by depth.

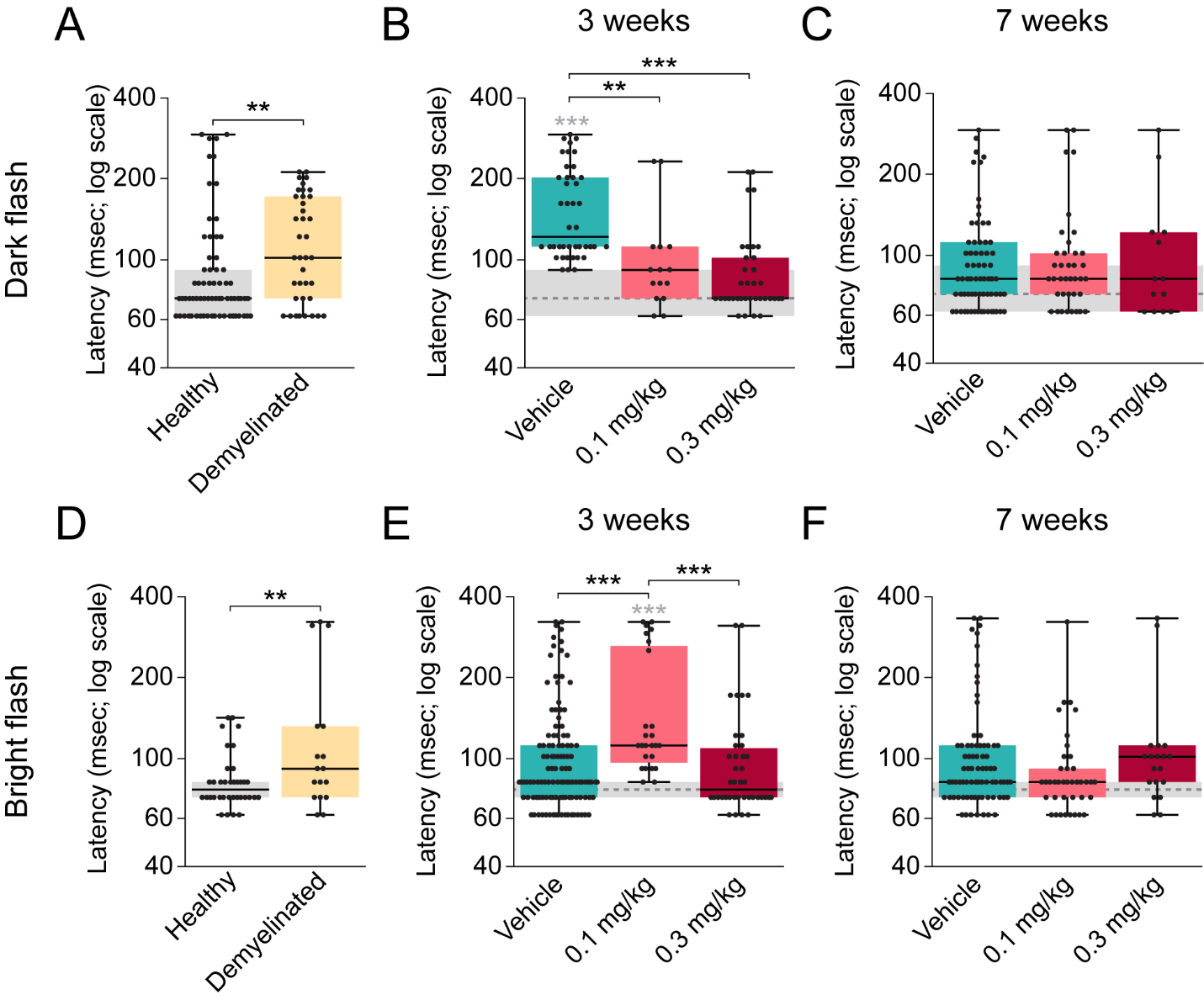

**Supplementary Fig.8. Single neuron responses to bright and dark flash**

**(A)** Single-neuron responses to dark flash are delayed after cuprizone-induced demyelination.

**(B)** Treatment with either low or high dose of LL-341070 for three weeks can reestablish the typical neuronal latencies to dark flash.

**(C)** Single-neuron responses to dark flash are normalized in all groups after seven weeks of remyelination.

**(D)** Single-neuron responses to bright flash are delayed after cuprizone-induced demyelination.

**(E)** Typical neuronal latencies to bright flash were restored after three weeks of treatment with vehicle or high dose LL-341070, while latency delays remained in mice treated with low dose LL-341070.

**(F)** Latency delays were recovered in all groups after seven weeks of remyelination.

In **A**, Median (IQR) [healthy: 72.25 (62.21-92.33), n = 68 neurons from 5 probes from 3 mice; demyelinated: 102.37 (72.25-172.65), n = 38 neurons from 7 probes from 5 mice; Z = 3.391, p = 0.0007, Wilcoxon].

In **B**, Median (IQR) [vehicle: 122.45 (112.41-202.77), n = 42 neurons from 5 probes from 2 mice; 0.1 mg/kg: 92.33 (72.25-112.41), n = 15 neurons from 4 probes from 4 mice; 0.3 mg/kg: 72.25 (72.25-102.37), n = 36 neurons from 4 probes from 2 mice; F(3)= 48.3; p < 0.0001, Kruskal-Wallis; vehicle vs healthy *** p < 0.0001, 0.1 mg/kg vs vehicle ** p = 0.002, 0.3 mg/kg vs vehicle *** p < 0.0001, Steel-Dwass].

In **C**, Median (IQR) [vehicle: 82.29 (72.25-112.41), n = 68 neurons from 10 probes from 4 mice; 0.1 mg/kg: 82.29 (72.25-102.37), n = 38 neurons from 8 probes from 3 mice; 0.3 mg/kg: 82.29 (72.25-122.45), n = 13 neurons from 1 probe from 1 mouse; F(3)= 3.1; p = 0.3763, Kruskal-Wallis].

In **D**, Median (IQR) [healthy: 77.27 (72.25-82.29), n = 36 neurons from 6 probes from 3 mice; demyelinated: 92.33 (72.25-132.49), n = 17 neurons from 4 probes from 4 mice; Z = 2.5782, p = 0.0099, Wilcoxon].

In **E**, Median (IQR) [vehicle: 82.33 (72.25-112.41), n = 115 neurons from 8 probes from 3 mice; 0.1 mg/kg: 112.41 (97.35-263.01), n = 25 neurons from 6 probes from 4 mice; 0.3 mg/kg: 77.27 (72.25-109.9), n = 40 neurons from 4 probes from 2 mice; F(3)= 25.84; p < 0.0001, Kruskal-Wallis; 0.1 mg/kg vs healthy *** p < 0.0001, vehicle vs 0.1 mg/kg *** p = 0.0005, 0.3 mg/kg vs 0.1 mg/kg *** p = 0.0004, Steel-Dwass].

In **F**, Median (IQR) [vehicle: 82.29 (72.25-112.41), n = 76 neurons from 11 probes from 4 mice; 0.1 mg/kg: 82.29 (72.25-92.33), n = 39 neurons from 6 probes from 3 mice; 0.3 mg/kg: 102.37 (82.29-112.41), n = 19 neurons from 3 probes from 1 mouse; F(3)= 9.6; p = 0.022, Kruskal-Wallis].

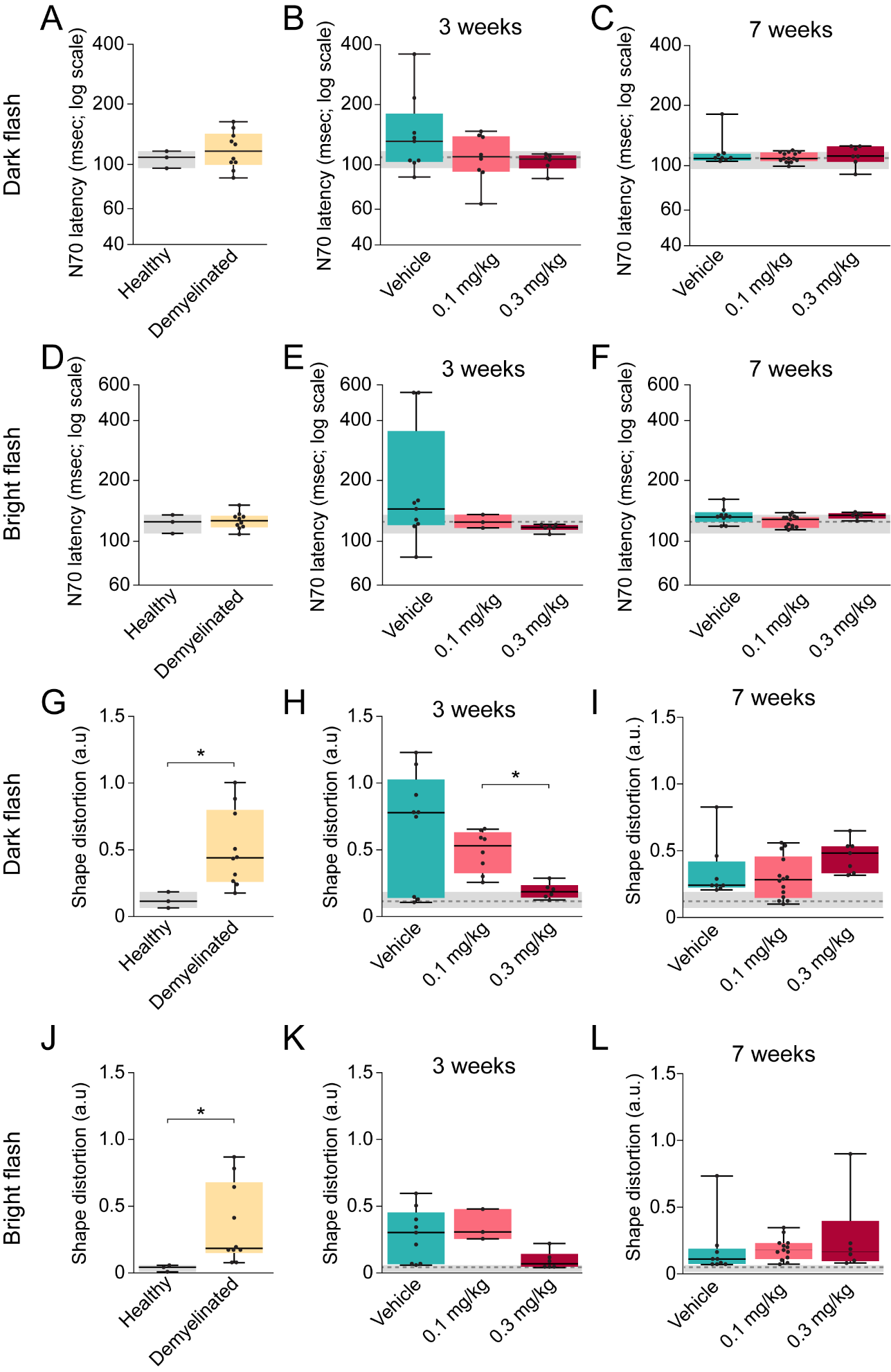

**Supplementary Fig.9. VEP responses to a dark or bright flash**

**(A)** VEP N70 latency to dark flash is not significantly altered by demyelination.

**(B)** All treatment groups show similar VEP N70 latencies to dark flash after 3 weeks of remyelination.

**(C)** There are no differences between groups in VEP N70 latencies to dark flash after 7 weeks of remyelination.

**(D)** Demyelination does not significantly delay VEP N70 latency to bright flash.

**(E)** VEP N70 latencies to bright flash are similar across groups after 3 weeks of remyelination.

**(F)** There are no differences between groups in VEP N70 latencies to bright flash after 7 weeks of remyelination.

**(G)** Demyelination alters the VEP response to dark flash

**(H)** After three weeks of remyelination, high-dose treatment with LL-341070 provides a larger benefit to altered VEP shape than low-dose treatment.

**(I)** VEP shape in response to dark flash is equivalent across groups seven weeks post-cuprizone

**(J)** Cuprizone alters the shape of VEP responses to bright flash

**(K)** Three weeks post cuprizone, there is no difference between groups in the shape of the VEP elicited by bright.

**(L)** The shape of VEPs in response to dark flash is equivalent across groups after seven weeks of remyelination

In **A**, Median (IQR) [healthy: 108.92 (96.12-116.93), n = 3 probes from 2 mice; demyelinated: 116.93 (99.92-143.04), n = 10 probes from 4 mice; Z = -0.42, p = 0.67, Wilcoxon].

In **B**, Median (IQR) [vehicle: 130.93 (103.32-179.75), n = 9 probes from 3 mice; 0.1 mg/kg: 109.52 (92.32-138.54), n = 8 probes from 3 mice; 0.3 mg/kg: 106.72 (95.52-111.12), n = 6 probes from 2 mice; F(3)= 1.66; p = 0.64, Kruskal-Wallis].

In **C**, Median (IQR) [vehicle: 108.92 (106.62-115.23), n = 8 probes from 3 mice; 0.1 mg/kg: 108.92 (106.02-117.03), n = 14 probes from 5 mice; 0.3 mg/kg: 112.12 (104.92-125.33), n = 7 probes from 3 mice; F(3)= 0.863; p = 0.83, Kruskal-Wallis].

In **D**, Median (IQR) [healthy: 124.13 (108.52-134.53), n = 3 probes from 2 mice; demyelinated: 0.13 (0.12-0.13), n = 10 probes from 4 mice; Z = -0.084, p = 0.93, Wilcoxon].

In **E**, Median (IQR) [vehicle: 144.14 (119.73-353.02), n = 9 probes from 3 mice; 0.1 mg/kg: 123.73 (115.73-134.93), n = 3 probes from 3 mice; 0.3 mg/kg: 116.73 (113.12-119.43), n = 6 probes from 2 mice; F(3)= 6.026; p = 0.11, Kruskal-Wallis].

In **F**, Median (IQR) [vehicle: 131.33 (124.13-138.74), n = 9 probes from 3 mice; 0.1 mg/kg: 127.73 (115.93-131.33), n = 13 probes from 5 mice; 0.3 mg/kg: 134.13 (129.23-137.34), n = 6 probes from 3 mice; F(3)= 6.28; p = 0.099, Kruskal-Wallis].

In **G**, Median (IQR) [healthy: 0.1104 (0.06-0.18), n = 3 probes from 2 mice; demyelinated: 0.43 (0.26-0.8), n = 10 probes from 4 mice; Z = -2.28, p = 0.022, Wilcoxon].

In **H**, Median (IQR) [vehicle: 0.77 (0.14-1.02), n = 9 probes from 3 mice; 0.1 mg/kg: 0.53 (0.32-0.63), n = 8 probes from 3 mice; 0.3 mg/kg: 0.18 (0.14-0.23), n = 6 probes from 2 mice; F(3)= 9.68; p = 0.022, Kruskal-Wallis; 0.3 mg/kg vs 0.1 mg/kg * p = 0.019, Steel-Dwass].

In **I**, Median (IQR) [vehicle: 0.23 (0.22-0.41), n = 8 probes from 3 mice; 0.1 mg/kg: 0.27 (0.14-0.45), n = 14 probes from 5 mice; 0.3 mg/kg: 0.48 (0.32-0.53), n = 7 probes from 3 mice; F(3)= 11.46; p = 0.0095, Kruskal-Wallis].

In **J**, Median (IQR) [healthy: 0.038 (0.0041-0.053), n = 3 probes from 2 mice; demyelinated: 0.18 (0.14-0.67), n = 10 probes from 4 mice; Z = -2.45, p = 0.014, Wilcoxon].

In **K**, Median (IQR) [vehicle: 0.3 (0.066-0.453), n = 9 probes from 3 mice; 0.1 mg/kg: 0.31 (0.26-0.48), n = 3 probes from 3 mice; 0.3 mg/kg: 0.069 (0.047-0.14), n = 6 probes from 2 mice; F(3)= 11.2; p = 0.0107, Kruskal-Wallis].

In **L**, Median (IQR) [vehicle: 0.10 (0.067-0.18), n = 9 probes from 3 mice; 0.1 mg/kg: 0.17 (0.1-0.22), n = 13 probes from 5 mice; 0.3 mg/kg: 0.159 (0.088-0.39), n = 6 probes from 3 mice; F(3)= 9.85; p = 0.02, Kruskal-Wallis].

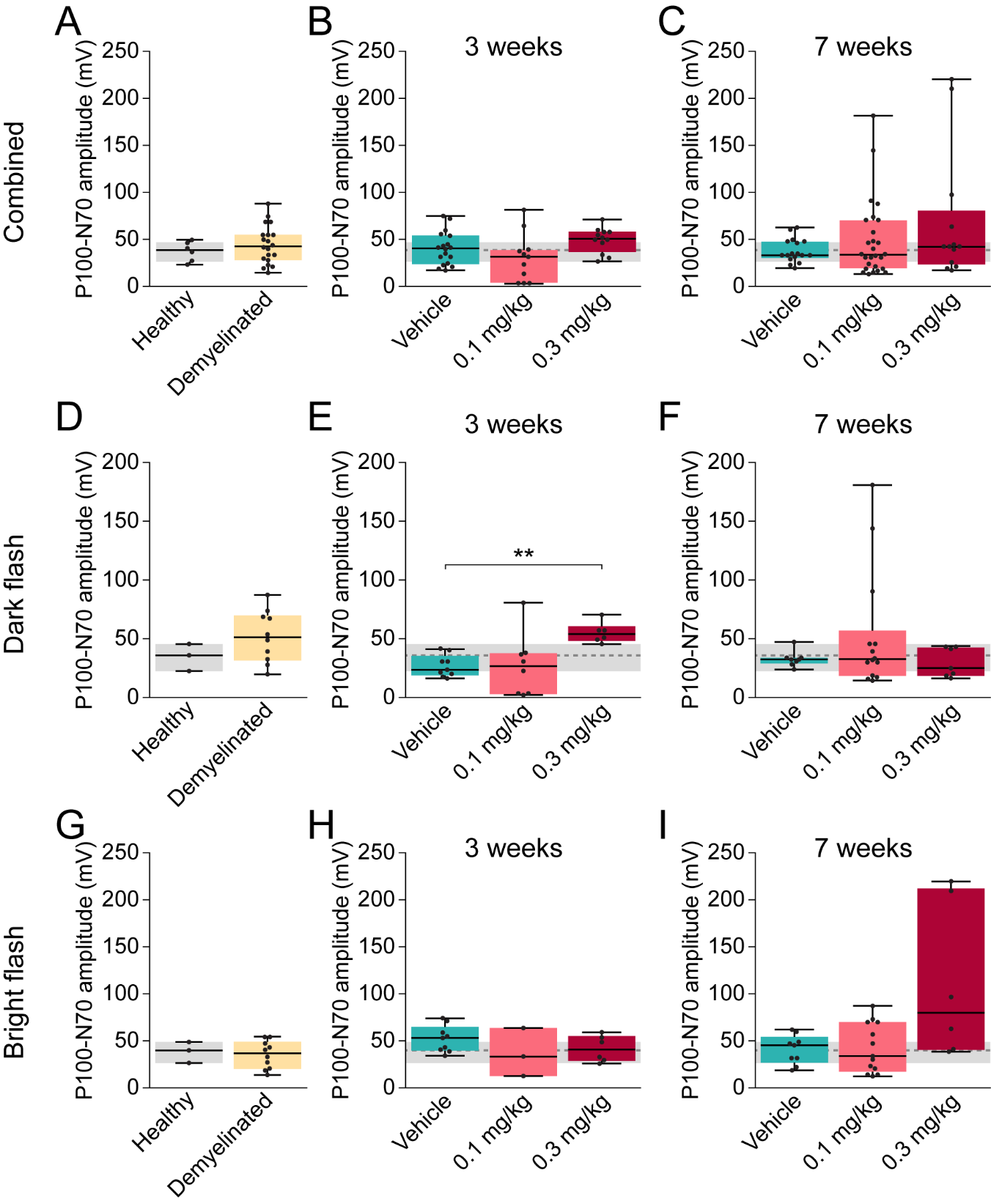

**Supplementary Fig.10. The amplitude of VEP responses is only minimally affected by demyelination and remyelination**

**(A)** The difference in amplitude of P100 and N70 VEP peaks in flash response is not significantly altered by demyelination.

**(B)** There is no difference between groups in P100-N70 VEP after three weeks of remyelination.

**(C)** There is no difference between groups in P100-N70 amplitude seven weeks after cuprizone.

**(D)** Demyelination does not affect the P100-N70 VEP amplitude in response to dark flash.

**(E)** Treatment with high dose LL-341070 for the first three weeks of remyelination increased the P100-N70 VEP amplitude in response to dark flash in comparison to the amplitude of vehicle-treated mice.

**(F)** There are no differences between groups in P100-N70 amplitude to dark flash after seven weeks of remyelination.

**(G)** The P100-N70 VEP amplitude in response to bright flash is not affected by cuprizone.

**(H)** There are no differences between groups in P100-N70 amplitude to bright flash after three weeks of remyelination.

**(I)** After seven weeks of remyelination, P100-N70 VEP amplitude in response to bright flash is similar across groups.

In **A**, Median (IQR) [healthy: 37.9 (25.51-46.35), n = 6 VEPs from 3 probes from 2 mice; demyelinated: 41.77 (27.31-54.31), n = 20 VEPs from 10 probes from 4 mice; Z = -0.7, p = 0.48, Wilcoxon].

In **B**, Median (IQR) [vehicle: 39.52 (22.88-53.39), n = 18 VEPs from 9 probes from 3 mice; 0.1 mg/kg: 30.65 (3.09-37.87), n = 11 VEPs from 3 probes from 3 mice; 0.3 mg/kg: 50.06 (36.01-57.27), n = 12 VEPs from 6 probes from 2 mice; F(3)= 6.22; p = 0.10, Kruskal-Wallis].

In **C**, Median (IQR) [vehicle: 32.49 (29.43-46.83), n = 17 VEPs from 9 probes from 3 mice; 0.1 mg/kg: 33.07 (18.67-69.5), n = 27 VEPs from 13 probes from 5 mice; 0.3 mg/kg: 41.41 (22.63-79.75), n = 13 VEPs from 6 probes from 3 mice; F(3)= 0.78; p = 0.85, Kruskal-Wallis].

In **D**, Median (IQR) [healthy: 35.9338 (22.5313-45.5413), n = 3 probes from 2 mice; demyelinated: 51.25 (31.37-69.87), n = 10 probes from 4 mice; Z = -1.1, p = 0.27, Wilcoxon].

In **E**, Median (IQR) [vehicle: 23.61 (18.72-35.46), n = 9 probes from 3 mice; 0.1 mg/kg: 26.65 (2.73-37.57), n = 8 probes from 3 mice; 0.3 mg/kg: 53.87 (48.26-60.67), n = 6 probes from 2 mice; F(3)= 11.03; p = 0.012, Kruskal-Wallis; 0.3 mg/kg vs vehicle ** p = 0.0097, Steel-Dwass].

In **F**, Median (IQR) [vehicle: 32.38 (28.82-33.74), n = 8 probes from 3 mice; 0.1 mg/kg: 32.49 (18.32-56.87), n = 14 probes from 5 mice; 0.3 mg/kg: 24.93 (18.47-42.54), n = 7 probes from 3 mice; F(3)= 0.79; p = 0.85, Kruskal-Wallis].

In **G**, Median (IQR) [healthy: 39.86 (26.5-48.78), n = 3 probes from 2 mice; demyelinated: 36.69 (20.07-48.95), n = 10 probes from 4 mice; Z = 0.084, p = 0.93, Wilcoxon].

In **H**, Median (IQR) [vehicle: 53.096 (39.73-64.99), n = 9 probes from 3 mice; 0.1 mg/kg: 33.41 (12.6-63.67), n = 3 probes from 3 mice; 0.3 mg/kg: 40.75 (28.58-55.08), n = 6 probes from 2 mice; F(3)= 3.22; p = 0.36, Kruskal-Wallis].

In **I**, Median (IQR) [vehicle: 45.38 (26.79-54.25), n = 9 probes from 3 mice; 0.1 mg/kg: 33.75 (17.24-69.9), n = 13 probes from 5 mice; 0.3 mg/kg: 79.75 (40.51-212.03), n = 6 probes from 3 mice; F(3)= 5.3; p = 0.15, Kruskal-Wallis].
