## Supplementary material for "Incomplete remyelination via endogenous or therapeutically enhanced oligodendrogenesis is sufficient to recover visual cortical function": Source Data and Statistics

**Figure 1**

**1C**

##### Oneway Analysis of Full nodes By Group

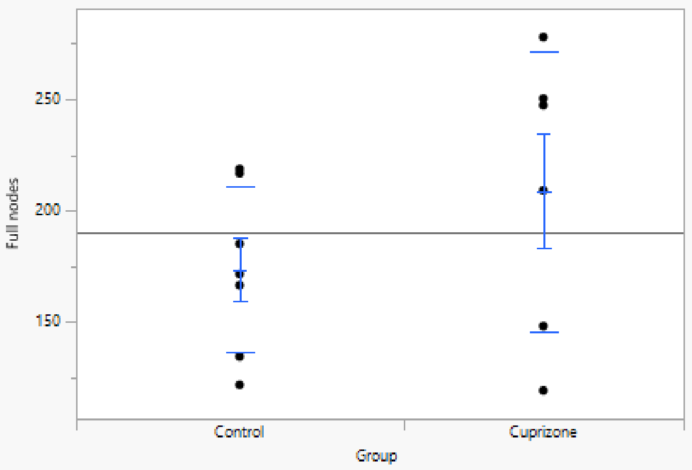

##### Means and Std Deviations

| Level | Number | Mean | Std Dev | Std Err Mean | Lower 95% | Upper 95% | Std Dev Lower 95% | Std Dev Upper 95% |
|---|---|---|---|---|---|---|---|---|
| Control | 7 | 173.42758 | 37.239684 | 14.075278 | 138.98662 | 207.86855 | 23.997007 | 82.004246 |
| Cuprizone | 6 | 208.61111 | 62.863314 | 25.663841 | 142.64011 | 274.58211 | 39.239784 | 154.17944 |

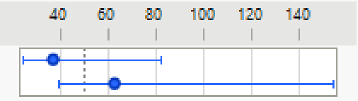

##### t Test

Cuprizone-Control

Assuming unequal variances

| | | | |
|---|---|---|---|
| Difference | 35.18 | t Ratio | 1.202024 |
| Std Err Dif | 29.27 | DF | 7.867169 |
| Upper CL Dif | 102.88 | Prob > \|t\| | 0.2643 |
| Lower CL Dif | -32.51 | Prob > t | 0.1321 |
| Confidence | 0.95 | Prob < t | 0.8679 |

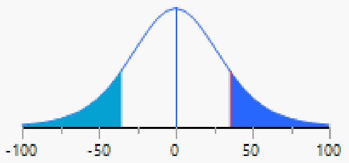

**1H**

| Lost asymptote | Gain asymptote | | | |
|---|---|---|---|---|
| 62.350722 | 52.264042 | | | |
| 57.692211 | 37.884252 | | | |
| 51.639086 | 45.444816 | | | |
| 14.699219 | 21.079175 | | | |
| 27.106146 | 35.978007 | | | |
| 78.19109 | 53.107853 | | | |
| 84.158801 | 46.805868 | | | |
| 80.031454 | 52.400282 | | | |
| 56.468931 | 48.592752 | | | |
| 51.998985 | 30.059736 | | | |
| 27.871313 | 32.4931 | | | |
| 70.32215 | 44.830646 | | | |
| 65.089514 | 50.158102 | | | |
| 71.296404 | 32.894871 | | | |
| 26.648747 | 26.322967 | | paired t test | 0.002676025 |
| | | | unpaired t test | 0.028972426 |

| Lost inflection point | Gain inflection point | | | |
|---|---|---|---|---|
| 0.708785 | 8.3455686 | | | |
| -1.326652 | 12.649627 | | | |
| -1.179215 | 10.44124 | | | |
| -4.333861 | 12.90742 | | | |
| -2.600023 | 7.3100664 | | | |
| -0.817762 | 10.865324 | | | |
| -3.613168 | 5.9314901 | | | |
| -2.726373 | 10.425748 | | | |
| 2.033625 | 8.569284 | | | |
| -0.813147 | 9.8532138 | | | |
| -4.222406 | 21.352316 | | | |
| -0.95256 | 11.046446 | | | |
| 0.802375 | 8.2522214 | | | |
| -1.911749 | 10.678233 | | | |
| -6.963465 | 9.1416281 | | paired t te | 1.5323E-06 |
| | | | unpaired t | 3.53904E-10 |
| -1.861039733 | 10.51798843 | | | |

**1I**

| | Lost | Gain | | | |
|---|---|---|---|---|---|
| < 30% loss | 27.35849 | 26.41509 | | | |
| | 26.92308 | 35.89744 | | | |
| | 26.27119 | 28.81356 | | < 30% loss | |
| | 14.28571 | 19.38776 | | paired t test | 0.15768 |
| | | | | unpaired t test | 0.430575 |
| 60-90% loss | 62.31884 | 52.17391 | | | |
| | 64.44444 | 53.33333 | | | |
| | 69.1358 | 45.67901 | | | |
| | 70.88608 | 35.44304 | | | |
| | 77.46479 | 50.70423 | | 60-90% loss | |
| | 78.74016 | 58.26772 | | paired t test | 0.002937 |
| | | | | unpaired t test | 0.000513 |
| | | | | overall | |
| | | | | paired t test | 0.043311 |
| | | | | unpaired t test | 0.226736 |

**Figure 3**

**3E**

##### Oneway Analysis of Running lost (% of baseline) By Manipulation

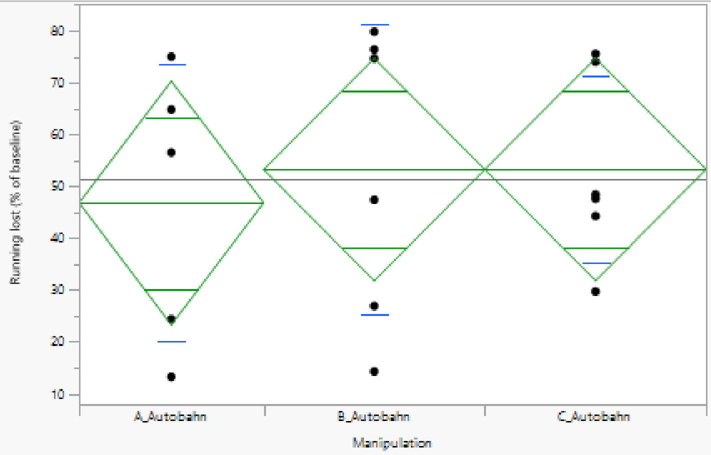

##### Oneway Anova

###### Summary of Fit

| | |
|---|---|
| Rsquare | 0.017209 |
| Adj Rsquare | -0.12319 |
| Root Mean Square Error | 24.50329 |
| Mean of Response | 51.35841 |
| Observations (or Sum Wgts) | 17 |

###### Analysis of Variance

| Source | DF | Sum of Squares | Mean Square | F Ratio | Prob > F |
|---|---|---|---|---|---|
| Manipulation | 2 | 147.1851 | 73.593 | 0.1226 | 0.8856 |
| Error | 14 | 8405.7578 | 600.411 | | |
| C. Total | 16 | 8552.9429 | | | |

###### Means for Oneway Anova

| Level | Number | Mean | Std Error | Lower 95% | Upper 95% |
|---|---|---|---|---|---|
| A_Autobahn | 5 | 46.8000 | 10.958 | 23.297 | 70.303 |
| B_Autobahn | 6 | 53.2444 | 10.003 | 31.789 | 74.700 |
| C_Autobahn | 6 | 53.2710 | 10.003 | 31.816 | 74.726 |

Std Error uses a pooled estimate of error variance

##### Tests that the Variances are Equal

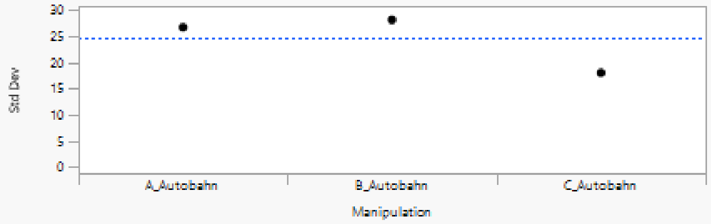

| Level | Count | Std Dev | MeanAbsDif to Mean | MeanAbsDif to Median |
|---|---|---|---|---|
| A_Autobahn | 5 | 26.65810 | 22.38272 | 20.43838 |
| B_Autobahn | 6 | 28.08714 | 23.70570 | 23.70570 |
| C_Autobahn | 6 | 17.99280 | 14.35066 | 12.73400 |

| Test | F Ratio | DFNum | DFDen | Prob > F |
|---|---|---|---|---|
| O'Brien[.5] | 1.2479 | 2 | 14 | 0.3172 |
| Brown-Forsythe | 0.8909 | 2 | 14 | 0.4323 |
| Levene | 1.6359 | 2 | 14 | 0.2299 |
| Bartlett | 0.4784 | 2 | . | 0.6198 |

Warning: Small sample sizes. Use Caution.

###### Welch's Test

Welch Anova testing Means Equal, allowing Std Devs Not Equal

| F Ratio | DFNum | DFDen | Prob > F |
|---|---|---|---|
| 0.1074 | 2 | 8.5206 | 0.8994 |

Excluded Rows 675

**3G**

#### Response Running gain (% of baseline)

##### Regression Plot

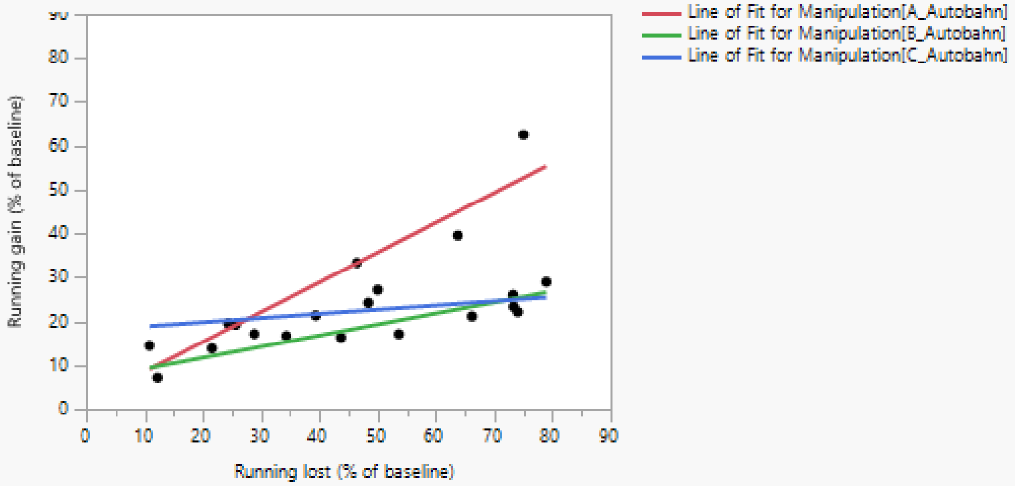

##### Effect Summary

| Source | Logworth | | PValue |
|---|---|---|---|
| Running lost (% of baseline) | 4.016 | | 0.00010 |
| Manipulation | 2.908 | | 0.00123 |
| Manipulation*Running lost (% of baseline) | 2.246 | | 0.00568 |

Remove Add Edit FDR

##### Summary of Fit

| | |
|---|---|
| RSquare | 0.818901 |
| RSquare Adj | 0.754223 |
| Root Mean Square Error | 5.793877 |
| Mean of Response | 23.54699 |
| Observations (or Sum Wgts) | 20 |

##### Analysis of Variance

| Source | DF | Sum of Squares | Mean Square | F Ratio |
|---|---|---|---|---|
| Model | 5 | 2125.1123 | 425.022 | 12.6612 |
| Error | 14 | 469.9661 | 33.569 | Prob > F |
| C. Total | 19 | 2595.0784 | | <.0001* |

##### Parameter Estimates

| Term | Estimate | Std Error | t Ratio | Prob>\|t\| |
|---|---|---|---|---|
| Intercept | 8.7360191 | 3.285473 | 2.66 | 0.0187* |
| Manipulation[A_Autobahn] | 8.8643211 | 1.936098 | 4.58 | 0.0004* |
| Manipulation[B_Autobahn] | -6.344155 | 1.839203 | -3.45 | 0.0039* |
| Running lost (% of baseline) | 0.3426447 | 0.063646 | 5.38 | <.0001* |
| Manipulation[A_Autobahn]*(Running lost (% of baseline)-47.1919) | 0.3366804 | 0.086159 | 3.91 | 0.0016* |
| Manipulation[B_Autobahn]*(Running lost (% of baseline)-47.1919) | -0.090556 | 0.084066 | -1.08 | 0.2996 |

##### Effect Tests

| Source | Nparm | DF | Sum of Squares | F Ratio | Prob > F |
|---|---|---|---|---|---|
| Manipulation | 2 | 2 | 753.39017 | 11.2215 | 0.0012* |
| Running lost (% of baseline) | 1 | 1 | 972.93016 | 28.9830 | <.0001* |
| Manipulation*Running lost (% of baseline) | 2 | 2 | 513.86842 | 7.6539 | 0.0057* |

#### Effect Details

##### Manipulation

###### Least Squares Means Table

| Level | Least Sq Mean | Std Error | Mean |
|---|---|---|---|
| A_Autobahn | 33.770389 | 2.4480733 | 29.5084 |
| B_Autobahn | 18.561913 | 2.2126084 | 19.4002 |
| C_Autobahn | 22.385901 | 2.2064433 | 22.5840 |

###### LSMeans Differences Tukey HSD

α= 0.050 Q= 2.61728

LSMean[j]

| LSMean[i] | Mean[i]-Mean[j] / Std Err Dif / Lower CL Dif / Upper CL Dif | A_Autobahn | B_Autobahn | C_Autobahn |
|---|---|---|---|---|
| A_Autobahn | Mean[i]-Mean[j] | 0 | 15.20848 | 11.38449 |
| | Std Err Dif | 0 | 3.299803 | 3.295672 |
| | Lower CL Dif | 0 | 6.571966 | 2.758789 |
| | Upper CL Dif | 0 | 23.84499 | 20.01019 |
| B_Autobahn | Mean[i]-Mean[j] | -15.2085 | 0 | -3.82399 |
| | Std Err Dif | 3.299803 | 0 | 3.124744 |
| | Lower CL Dif | -23.845 | 0 | -12.0023 |
| | Upper CL Dif | -6.57197 | 0 | 4.354345 |
| C_Autobahn | Mean[i]-Mean[j] | -11.3845 | 3.823988 | 0 |
| | Std Err Dif | 3.295672 | 3.124744 | 0 |
| | Lower CL Dif | -20.0102 | -4.35434 | 0 |
| | Upper CL Dif | -2.75879 | 12.00232 | 0 |

| Level | | | Least Sq Mean |
|---|---|---|---|
| A_Autobahn | A | | 33.770389 |
| C_Autobahn | | B | 22.385901 |
| B_Autobahn | | B | 18.561913 |

Levels not connected by same letter are significantly different.

| Level | - Level | Difference | Std Err Dif | Lower CL | Upper CL | p-Value |
|---|---|---|---|---|---|---|
| A_Autobahn | B_Autobahn | 15.20848 | 3.299803 | 6.57197 | 23.84499 | 0.0011* |
| A_Autobahn | C_Autobahn | 11.38449 | 3.295672 | 2.75879 | 20.01019 | 0.0101* |
| C_Autobahn | B_Autobahn | 3.82399 | 3.124744 | -4.35434 | 12.00232 | 0.4592 |

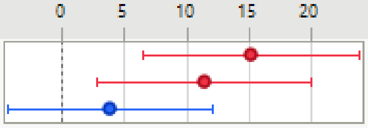

###### Least Squares Means Plot

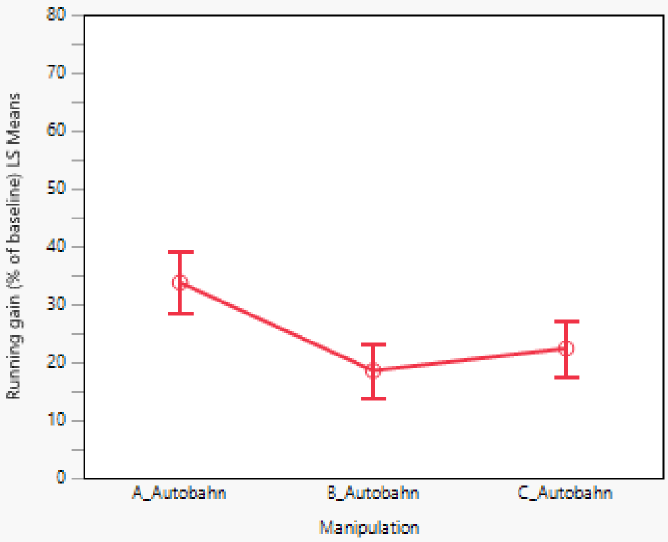

##### Running lost (% of baseline)

##### Manipulation*Running lost (% of baseline)

**3H**

#### Response Running gain (% of baseline)

##### Regression Plot

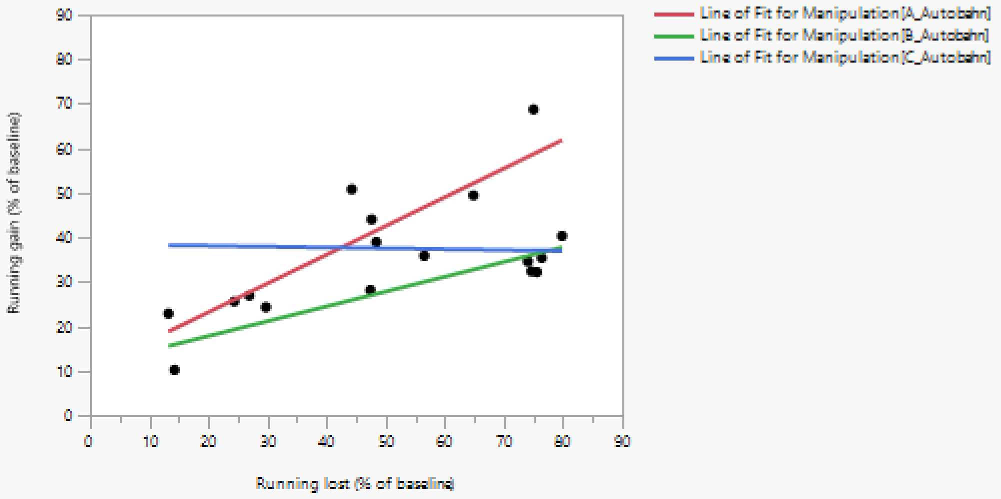

##### Effect Summary

| Source | Logworth | PValue |
|---|---|---|
| Running lost (% of baseline) | 2.125 | 0.00749 |
| Manipulation | 1.432 | 0.03697 |
| Manipulation*Running lost (% of baseline) | 1.126 | 0.07480 |

Remove Add Edit ☐ FDR

##### Summary of Fit

| | |
|---|---|
| RSquare | 0.723905 |
| RSquare Adj | 0.598407 |
| Root Mean Square Error | 8.403394 |
| Mean of Response | 35.35304 |
| Observations (or Sum Wgts) | 17 |

##### Analysis of Variance

| Source | DF | Sum of Squares | Mean Square | F Ratio |
|---|---|---|---|---|
| Model | 5 | 2036.6897 | 407.338 | 5.7683 |
| Error | 11 | 776.7874 | 70.617 | Prob > F |
| C. Total | 16 | 2813.4771 | | 0.0075* |

##### Parameter Estimates

| Term | Estimate | Std Error | t Ratio | Prob>\|t\| |
|---|---|---|---|---|
| Intercept | 19.982814 | 5.442919 | 3.67 | 0.0037* |
| Manipulation[A_Autobahn] | 7.0361033 | 3.024358 | 2.33 | 0.0401* |
| Manipulation[B_Autobahn] | -8.144566 | 2.865532 | -2.84 | 0.0160* |
| Running lost (% of baseline) | 0.3201401 | 0.097963 | 3.27 | 0.0075* |
| Manipulation[A_Autobahn]*(Running lost (% of baseline)-51.3584) | 0.3262556 | 0.133707 | 2.44 | 0.0328* |
| Manipulation[B_Autobahn]*(Running lost (% of baseline)-51.3584) | 0.0132619 | 0.124757 | 0.11 | 0.9173 |

##### Effect Tests

| Source | Nparm | DF | Sum of Squares | F Ratio | Prob > F |
|---|---|---|---|---|---|
| Manipulation | 2 | 2 | 638.01242 | 4.5174 | 0.0370* |
| Running lost (% of baseline) | 1 | 1 | 754.16209 | 10.6796 | 0.0075* |
| Manipulation*Running lost (% of baseline) | 2 | 2 | 467.84876 | 3.3126 | 0.0748 |

#### Effect Details

##### Manipulation

###### Least Squares Means Table

| Level | Least Sq Mean | Std Error | Mean |
|---|---|---|---|
| A_Autobahn | 43.460801 | 3.8261729 | 40.5143 |
| B_Autobahn | 28.280132 | 3.4399402 | 28.9089 |
| C_Autobahn | 37.533160 | 3.4538519 | 37.4961 |

###### LSMeans Differences Tukey HSD

α= 0.050 Q= 2.70081

| LSMean[i] | Mean[i]-Mean[j]<br>Std Err Dif<br>Lower CL Dif<br>Upper CL Dif | A_Autobahn | B_Autobahn | C_Autobahn |
|---|---|---|---|---|
| A_Autobahn | | 0<br>0<br>0<br>0 | 15.18067<br>5.145171<br>1.284559<br>29.07678 | 5.927641<br>5.154483<br>-7.99362<br>19.8489 |
| B_Autobahn | | -15.1807<br>5.145171<br>-29.0768<br>-1.28456 | 0<br>0<br>0<br>0 | -9.25303<br>4.874657<br>-22.4185<br>3.912475 |
| C_Autobahn | | -5.92764<br>5.154483<br>-19.8489<br>7.993618 | 9.253029<br>4.874657<br>-3.91248<br>22.41853 | 0<br>0<br>0<br>0 |

| Level | | | Least Sq Mean |
|---|---|---|---|
| A_Autobahn | A | | 43.460801 |
| C_Autobahn | A | B | 37.533160 |
| B_Autobahn | | B | 28.280132 |

Levels not connected by same letter are significantly different.

| Level | - Level | Difference | Std Err Dif | Lower CL | Upper CL | p-Value |
|---|---|---|---|---|---|---|
| A_Autobahn | B_Autobahn | 15.18067 | 5.145171 | 1.28456 | 29.07678 | 0.0326* |
| C_Autobahn | B_Autobahn | 9.25303 | 4.874657 | -3.91248 | 22.41853 | 0.1851 |
| A_Autobahn | C_Autobahn | 5.92764 | 5.154483 | -7.99362 | 19.84890 | 0.5052 |

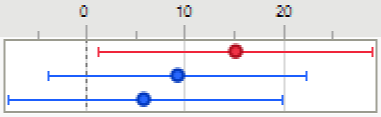

###### Least Squares Means Plot

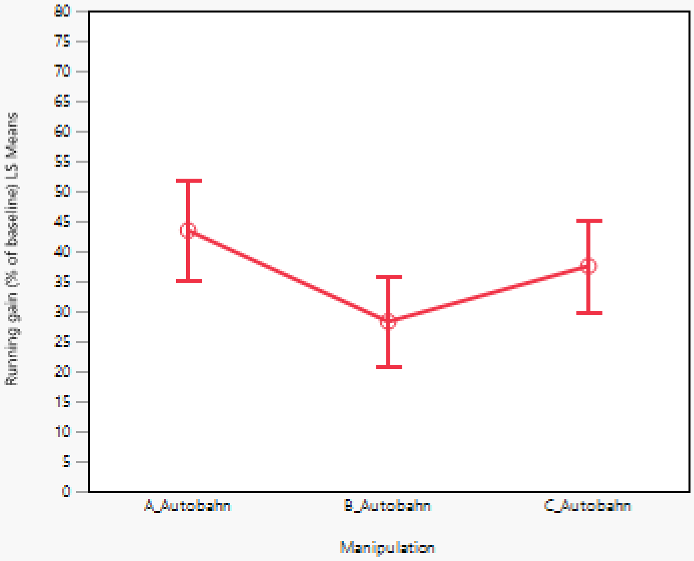

##### Running lost (% of baseline)

##### Manipulation*Running lost (% of baseline)

#### Response Running gain (% of baseline)

##### Regression Plot

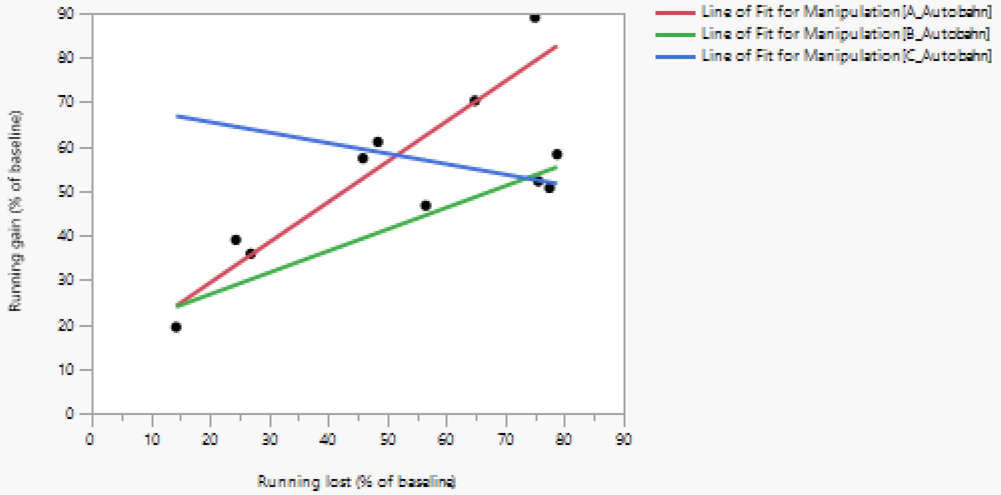

##### Effect Summary

| Source | Logworth | PValue |
|---|---|---|
| Running lost (% of baseline) | 1.133 | 0.07370 |
| Manipulation | 0.927 | 0.11831 |
| Manipulation*Running lost (% of baseline) | 0.824 | 0.14982 |

Remove Add Edit FDR

##### Summary of Fit

| | |
|---|---|
| RSquare | 0.862221 |
| RSquare Adj | 0.724442 |
| Root Mean Square Error | 9.642918 |
| Mean of Response | 52.73316 |
| Observations (or Sum Wgts) | 11 |

##### Analysis of Variance

| Source | DF | Sum of Squares | Mean Square | F Ratio |
|---|---|---|---|---|
| Model | 5 | 2909.5335 | 581.907 | 6.2580 |
| Error | 5 | 464.9293 | 92.986 | Prob > F |
| C. Total | 10 | 3374.4628 | | 0.0328* |

##### Parameter Estimates

| Term | Estimate | Std Error | t Ratio | Prob>\|t\| |
|---|---|---|---|---|
| Intercept | 32.807834 | 9.961421 | 3.29 | 0.0216* |
| Manipulation[A_Autobahn] | 6.2463062 | 4.085073 | 1.53 | 0.1868 |
| Manipulation[B_Autobahn] | -10.40576 | 4.096036 | -2.54 | 0.0519 |
| Running lost (% of baseline) | 0.3865671 | 0.171335 | 2.26 | 0.0737 |
| Manipulation[A_Autobahn]*(Running lost (% of baseline)-53.4581) | 0.5214297 | 0.225704 | 2.31 | 0.0689 |
| Manipulation[B_Autobahn]*(Running lost (% of baseline)-53.4581) | 0.101397 | 0.196234 | 0.52 | 0.6274 |

##### Effect Tests

| Source | Nparm | DF | Sum of Squares | F Ratio | Prob > F |
|---|---|---|---|---|---|
| Manipulation | 2 | 2 | 626.95793 | 3.3713 | 0.1183 |
| Running lost (% of baseline) | 1 | 1 | 473.34123 | 5.0905 | 0.0737 |
| Manipulation*Running lost (% of baseline) | 2 | 2 | 528.55810 | 2.8421 | 0.1498 |

#### Effect Details

##### Manipulation

###### Least Squares Means Table

| Level | Least Sq Mean | Std Error | Mean |
|---|---|---|---|
| A_Autobahn | 59.719283 | 4.8414863 | 61.2889 |
| B_Autobahn | 43.067214 | 4.8691946 | 41.0643 |
| C_Autobahn | 57.632433 | 5.7202578 | 56.8840 |

###### LSMeans Differences Tukey HSD

α= 0.050 Q= 3.25386

LSMean[j]

| LSMean[i] | Mean[i]-Mean[j] / Std Err Dif / Lower CL Dif / Upper CL Dif | A_Autobahn | B_Autobahn | C_Autobahn |
|---|---|---|---|---|
| A_Autobahn | | 0 | 16.65207 | 2.08685 |
| | | 0 | 6.866516 | 7.494087 |
| | | 0 | -5.69063 | -22.2979 |
| | | 0 | 38.99477 | 26.47158 |
| B_Autobahn | | -16.6521 | 0 | -14.5652 |
| | | 6.866516 | 0 | 7.512017 |
| | | -38.9948 | 0 | -39.0083 |
| | | 5.690634 | 0 | 9.877857 |
| C_Autobahn | | -2.08685 | 14.56522 | 0 |
| | | 7.494087 | 7.512017 | 0 |
| | | -26.4716 | -9.87786 | 0 |
| | | 22.29788 | 39.00829 | 0 |

| Level | | Least Sq Mean |
|---|---|---|
| A_Autobahn | A | 59.719283 |
| C_Autobahn | A | 57.632433 |
| B_Autobahn | A | 43.067214 |

Levels not connected by same letter are significantly different.

| Level | - Level | Difference | Std Err Dif | Lower CL | Upper CL | p-Value |
|---|---|---|---|---|---|---|
| A_Autobahn | B_Autobahn | 16.65207 | 6.866516 | -5.6906 | 38.99477 | 0.1263 |
| C_Autobahn | B_Autobahn | 14.56522 | 7.512017 | -9.8779 | 39.00829 | 0.2225 |
| A_Autobahn | C_Autobahn | 2.08685 | 7.494087 | -22.2979 | 26.47158 | 0.9585 |

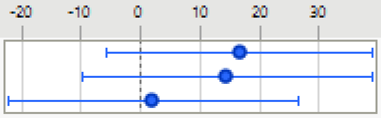

###### Least Squares Means Plot

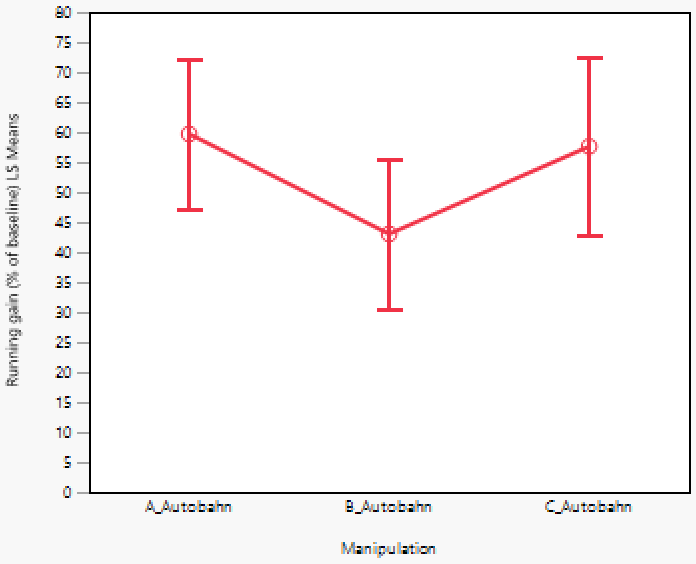

##### Running lost (% of baseline)

##### Manipulation*Running lost (% of baseline)

**3I**

#### Response Max gain rate

##### Whole Model

###### Regression Plot

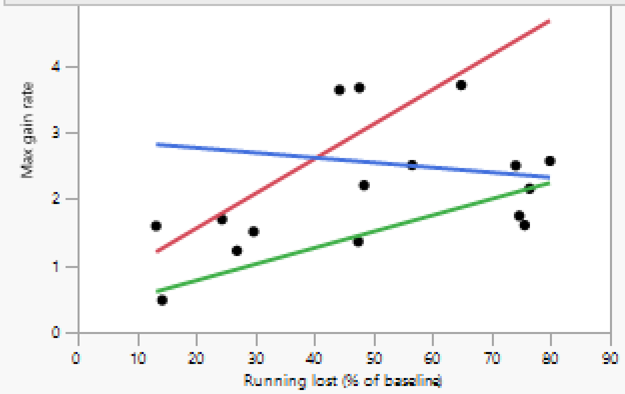

###### Actual by Predicted Plot

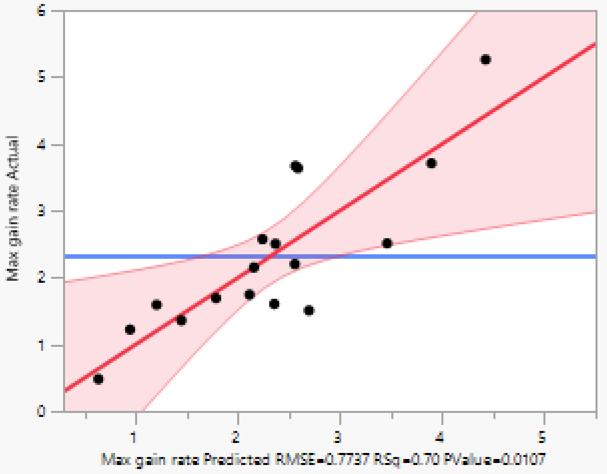

###### Effect Summary

| Source | Logworth | | PValue |
|---|---|---|---|
| Manipulation | 1.815 | | 0.01533 |
| Running lost (% of baseline) | 1.575 | | 0.02660 |
| Manipulation*Running lost (% of baseline) | 1.072 | | 0.08474 |

Remove Add Edit ☐ FDR

###### Residual by Predicted Plot

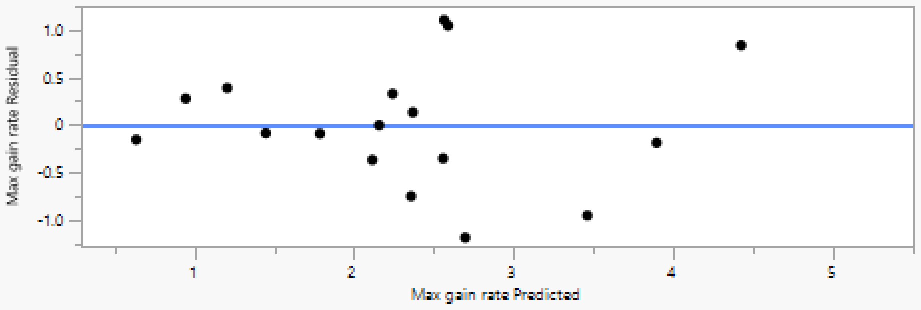

#### Summary of Fit

| | |
|---|---|
| RSquare | 0.703556 |
| RSquare Adj | 0.568809 |
| Root Mean Square Error | 0.773669 |
| Mean of Response | 2.31962 |
| Observations (or Sum Wgts) | 17 |

#### Analysis of Variance

| Source | DF | Sum of Squares | Mean Square | F Ratio |
|---|---|---|---|---|
| Model | 5 | 15.626422 | 3.12528 | 5.2213 |
| Error | 11 | 6.584201 | 0.59856 | Prob > F |
| C. Total | 16 | 22.210623 | | 0.0107* |

#### Parameter Estimates

| Term | Estimate | Std Error | t Ratio | Prob>\|t\| |
|---|---|---|---|---|
| Intercept | 1.2383278 | 0.501109 | 2.47 | 0.0311* |
| Manipulation[A_Autobahn] | 0.7677887 | 0.278441 | 2.76 | 0.0186* |
| Manipulation[B_Autobahn] | -0.879779 | 0.263819 | -3.33 | 0.0067* |
| Running lost (% of baseline) | 0.0230735 | 0.009019 | 2.56 | 0.0266* |
| Manipulation[A_Autobahn]*(Running lost (% of baseline)-51.3584) | 0.029045 | 0.01231 | 2.36 | 0.0378* |
| Manipulation[B_Autobahn]*(Running lost (% of baseline)-51.3584) | 0.0014239 | 0.011486 | 0.12 | 0.9036 |

#### Effect Tests

| Source | Nparm | DF | Sum of Squares | F Ratio | Prob > F |
|---|---|---|---|---|---|
| Manipulation | 2 | 2 | 7.4901473 | 6.2568 | 0.0153* |
| Running lost (% of baseline) | 1 | 1 | 3.9175116 | 6.5449 | 0.0266* |
| Manipulation*Running lost (% of baseline) | 2 | 2 | 3.7291313 | 3.1151 | 0.0847 |

#### Manipulation

##### Least Squares Means Table

| Level | Least Sq Mean | Std Error | Mean |
|---|---|---|---|
| A_Autobahn | 3.1911329 | 0.35226139 | 2.95356 |
| B_Autobahn | 1.5435657 | 0.31670239 | 1.58977 |
| C_Autobahn | 2.5353341 | 0.31798320 | 2.52119 |

##### LSMeans Differences Tukey HSD

α= 0.050 Q= 2.70081

LSMean[j]

| LSMean[i] | Mean[i]-Mean[j] / Std Err Dif / Lower CL Dif / Upper CL Dif | A_Autobahn | B_Autobahn | C_Autobahn |
|---|---|---|---|---|
| A_Autobahn | | 0 | 1.647567 | 0.655799 |
| | | 0 | 0.473697 | 0.474554 |
| | | 0 | 0.368205 | -0.62588 |
| | | 0 | 2.92693 | 1.937477 |
| B_Autobahn | | -1.64757 | 0 | -0.99177 |
| | | 0.473697 | 0 | 0.448791 |
| | | -2.92693 | 0 | -2.20387 |
| | | -0.3682 | 0 | 0.22033 |
| C_Autobahn | | -0.6558 | 0.991768 | 0 |
| | | 0.474554 | 0.448791 | 0 |
| | | -1.93748 | -0.22033 | 0 |
| | | 0.625879 | 2.203867 | 0 |

| Level | | | Least Sq Mean |
|---|---|---|---|
| A_Autobahn | A | | 3.1911329 |
| C_Autobahn | A | B | 2.5353341 |
| B_Autobahn | | B | 1.5435657 |

Levels not connected by same letter are significantly different.

| Level | - Level | Difference | Std Err Dif | Lower CL | Upper CL | p-Value |
|---|---|---|---|---|---|---|
| A_Autobahn | B_Autobahn | 1.647567 | 0.4736966 | 0.368205 | 2.926930 | 0.0132* |
| C_Autobahn | B_Autobahn | 0.991768 | 0.4487914 | -0.220330 | 2.203867 | 0.1133 |
| A_Autobahn | C_Autobahn | 0.655799 | 0.4745539 | -0.625879 | 1.937477 | 0.3829 |

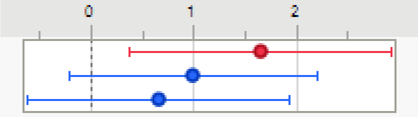

**3J**

#### Response Gain rate 0-11

##### Regression Plot

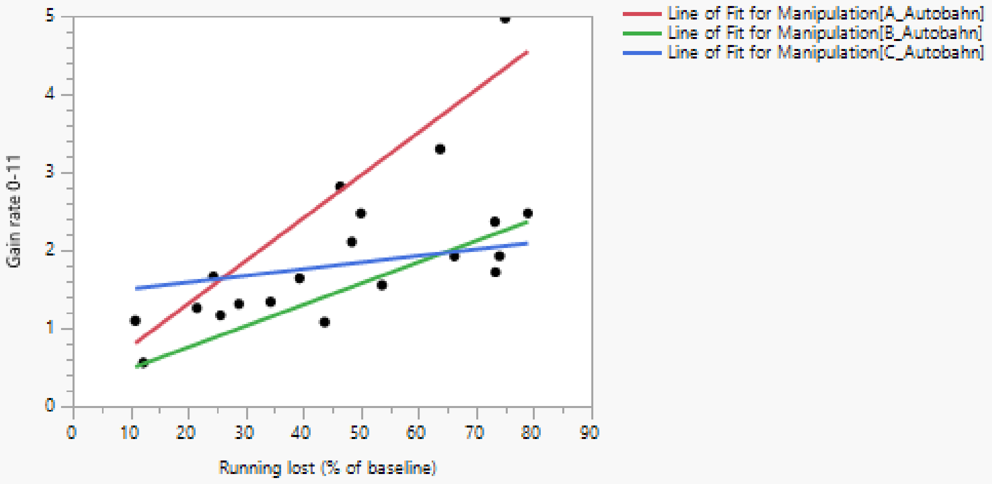

##### Effect Summary

| Source | Logworth | PValue |
|---|---|---|
| Running lost (% of baseline) | 4.737 | 0.00002 |
| Manipulation | 3.456 | 0.00035 |
| Running lost (% of baseline)*Manipulation | 2.244 | 0.00570 |

Remove Add Edit FDR

##### Summary of Fit

| | |
|---|---|
| RSquare | 0.854342 |
| RSquare Adj | 0.802322 |
| Root Mean Square Error | 0.434285 |
| Mean of Response | 1.93541 |
| Observations (or Sum Wgts) | 20 |

##### Analysis of Variance

| Source | DF | Sum of Squares | Mean Square | F Ratio |
|---|---|---|---|---|
| Model | 5 | 15.487284 | 3.09746 | 16.4231 |
| Error | 14 | 2.640444 | 0.18860 | Prob > F |
| C. Total | 19 | 18.127729 | | <.0001* |

##### Parameter Estimates

| Term | Estimate | Std Error | t Ratio | Prob>\|t\| |
|---|---|---|---|---|
| Intercept | 0.6122073 | 0.246265 | 2.49 | 0.0262* |
| Running lost (% of baseline) | 0.0302378 | 0.004771 | 6.34 | <.0001* |
| Manipulation[A_Autobahn] | 0.7647603 | 0.145122 | 5.27 | 0.0001* |
| Manipulation[B_Autobahn] | -0.543327 | 0.137859 | -3.94 | 0.0015* |
| (Running lost (% of baseline)-47.1919)*Manipulation[A_Autobahn] | 0.0247003 | 0.006458 | 3.82 | 0.0019* |
| (Running lost (% of baseline)-47.1919)*Manipulation[B_Autobahn] | -0.002936 | 0.006301 | -0.47 | 0.6484 |

##### Effect Tests

| Source | Nparm | DF | Sum of Squares | F Ratio | Prob > F |
|---|---|---|---|---|---|
| Running lost (% of baseline) | 1 | 1 | 7.5769576 | 40.1741 | <.0001* |
| Manipulation | 2 | 2 | 5.5897721 | 14.8189 | 0.0003* |
| Running lost (% of baseline)*Manipulation | 2 | 2 | 2.8833221 | 7.6439 | 0.0057* |

#### Effect Details

##### Running lost (% of baseline)

##### Manipulation

###### Least Squares Means Table

| Level | Least Sq Mean | Std Error | Mean |
|---|---|---|---|
| A_Autobahn | 2.8039477 | 0.18349731 | 2.45928 |
| B_Autobahn | 1.4958600 | 0.16584784 | 1.58665 |
| C_Autobahn | 1.8177547 | 0.16538573 | 1.83514 |

###### LSMeans Differences Tukey HSD

α= 0.050 Q= 2.61728

LSMean[j]

| LSMean[i] | Mean[i]-Mean[j]<br>Std Err Dif<br>Lower CL Dif<br>Upper CL Dif | A_Autobahn | B_Autobahn | C_Autobahn |
|---|---|---|---|---|
| | A_Autobahn | 0 | 1.308088 | 0.986193 |
| | | 0 | 0.247339 | 0.24703 |
| | | 0 | 0.660731 | 0.339647 |
| | | 0 | 1.955444 | 1.632739 |
| | B_Autobahn | -1.30809 | 0 | -0.32189 |
| | | 0.247339 | 0 | 0.234218 |
| | | -1.95544 | 0 | -0.93491 |
| | | -0.66073 | 0 | 0.291119 |
| | C_Autobahn | -0.98619 | 0.321895 | 0 |
| | | 0.24703 | 0.234218 | 0 |
| | | -1.63274 | -0.29112 | 0 |
| | | -0.33965 | 0.934908 | 0 |

| Level | | | Least Sq Mean |
|---|---|---|---|
| A_Autobahn | A | | 2.8039477 |
| C_Autobahn | | B | 1.8177547 |
| B_Autobahn | | B | 1.4958600 |

Levels not connected by same letter are significantly different.

| Level | - Level | Difference | Std Err Dif | Lower CL | Upper CL | p-Value |
|---|---|---|---|---|---|---|
| A_Autobahn | B_Autobahn | 1.308088 | 0.2473394 | 0.660731 | 1.955444 | 0.0003* |
| A_Autobahn | C_Autobahn | 0.986193 | 0.2470298 | 0.339647 | 1.632739 | 0.0036* |
| C_Autobahn | B_Autobahn | 0.321895 | 0.2342177 | -0.291119 | 0.934908 | 0.3801 |

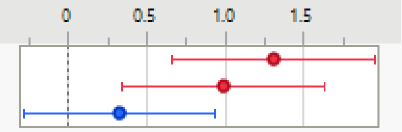

##### Running lost (% of baseline)*Manipulation

#### Response Gain rate 11-18

##### Whole Model

###### Regression Plot

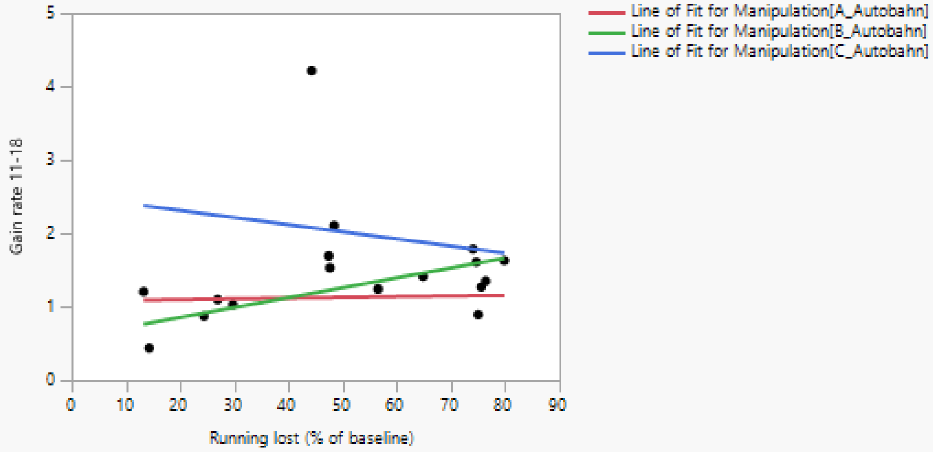

###### Actual by Predicted Plot

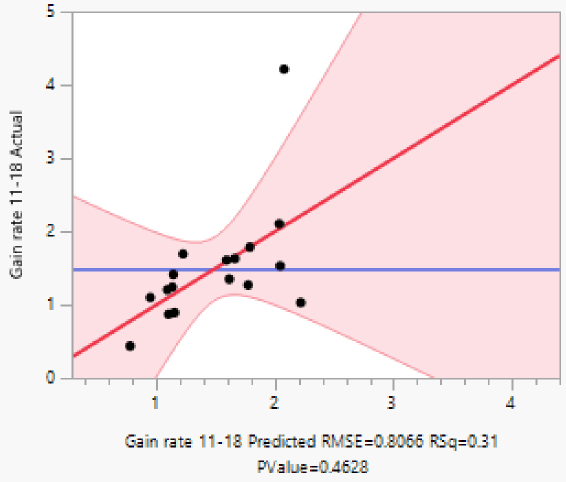

###### Effect Summary

| Source | Logworth | PValue | |
|---|---|---|---|
| Manipulation | 0.712 | 0.19411 | |
| Running lost (% of baseline)*Manipulation | 0.216 | 0.60782 | |
| Running lost (% of baseline) | 0.061 | 0.86905 | ^ |

Remove Add Edit FDR ('^' denotes effects with containing effects above them)

###### Residual by Predicted Plot

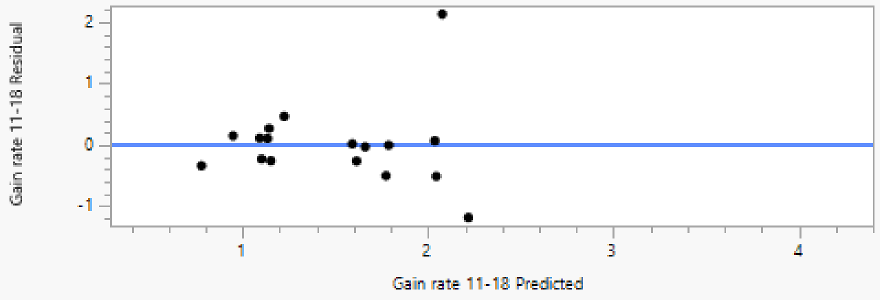

#### Summary of Fit

| | |
|---|---|
| RSquare | 0.311905 |
| RSquare Adj | -0.00087 |
| Root Mean Square Error | 0.806576 |
| Mean of Response | 1.492839 |
| Observations (or Sum Wgts) | 17 |

#### Analysis of Variance

| Source | DF | Sum of Squares | Mean Square | F Ratio |
|---|---|---|---|---|
| Model | 5 | 3.243822 | 0.648764 | 0.9972 |
| Error | 11 | 7.156206 | 0.650564 | Prob > F |
| C. Total | 16 | 10.400027 | | 0.4628 |

#### Parameter Estimates

| Term | Estimate | Std Error | t Ratio | Prob>\|t\| |
|---|---|---|---|---|
| Intercept | 1.3901048 | 0.522423 | 2.66 | 0.0222* |
| Running lost (% of baseline) | 0.0015868 | 0.009403 | 0.17 | 0.8691 |
| Manipulation[A_Autobahn] | -0.342339 | 0.290284 | -1.18 | 0.2631 |
| Manipulation[B_Autobahn] | -0.194021 | 0.27504 | -0.71 | 0.4952 |
| (Running lost (% of baseline)-51.3584)*Manipulation[A_Autobahn] | -0.000603 | 0.012833 | -0.05 | 0.9634 |
| (Running lost (% of baseline)-51.3584)*Manipulation[B_Autobahn] | 0.0118841 | 0.011974 | 0.99 | 0.3423 |

#### Effect Tests

| Source | Nparm | DF | Sum of Squares | F Ratio | Prob > F |
|---|---|---|---|---|---|
| Running lost (% of baseline) | 1 | 1 | 0.0185270 | 0.0285 | 0.8691 |
| Manipulation | 2 | 2 | 2.4849488 | 1.9098 | 0.1941 |
| Running lost (% of baseline)*Manipulation | 2 | 2 | 0.6780308 | 0.5211 | 0.6078 |

#### Response Gain rate 18-35

##### Whole Model

###### Regression Plot

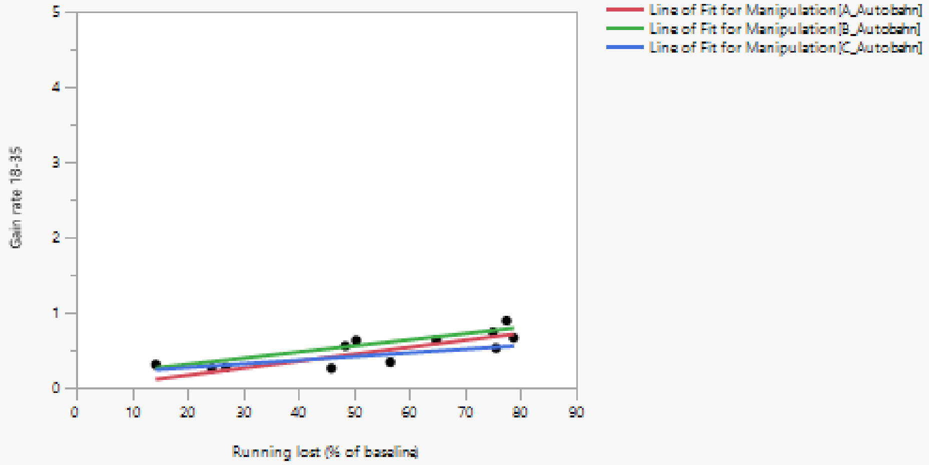

###### Actual by Predicted Plot

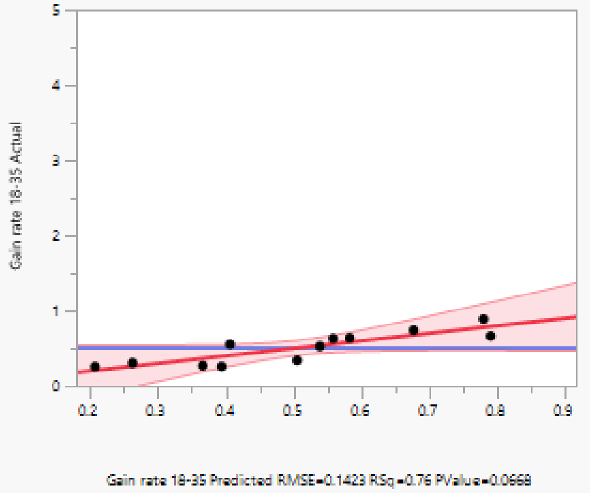

###### Effect Summary

| Source | Logworth | PValue |
|---|---|---|
| Running lost (% of baseline) | 1.592 | 0.02559 |
| Manipulation | 0.435 | 0.36718 |
| Running lost (% of baseline)*Manipulation | 0.080 | 0.83243 |

Remove Add Edit ☐ FDR

###### Residual by Predicted Plot

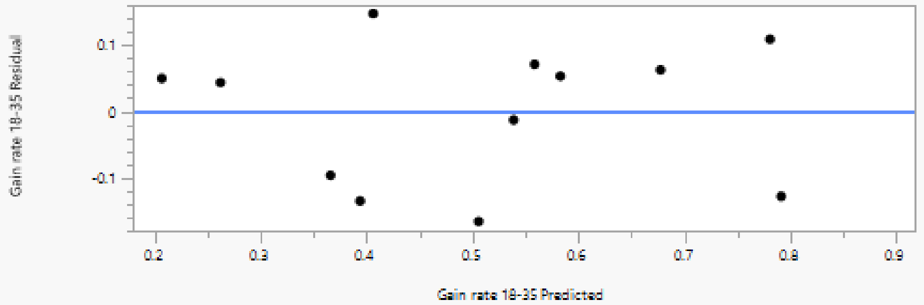

#### Summary of Fit

| | |
|---|---|
| RSquare | 0.760841 |
| RSquare Adj | 0.561542 |
| Root Mean Square Error | 0.142316 |
| Mean of Response | 0.505851 |
| Observations (or Sum Wgts) | 12 |

#### Analysis of Variance

| Source | DF | Sum of Squares | Mean Square | F Ratio |
|---|---|---|---|---|
| Model | 5 | 0.38660467 | 0.077321 | 3.8176 |
| Error | 6 | 0.12152335 | 0.020254 | Prob > F |
| C. Total | 11 | 0.50812801 | | 0.0668 |

#### Parameter Estimates

| Term | Estimate | Std Error | t Ratio | Prob>\|t\| |
|---|---|---|---|---|
| Intercept | 0.0983652 | 0.146678 | 0.67 | 0.5274 |
| Running lost (% of baseline) | 0.0074612 | 0.002529 | 2.95 | 0.0256* |
| Manipulation[A_Autobahn] | -0.020585 | 0.059439 | -0.35 | 0.7409 |
| Manipulation[B_Autobahn] | 0.0862335 | 0.056599 | 1.52 | 0.1784 |
| (Running lost (% of baseline)-53.2008)*Manipulation[A_Autobahn] | 0.001836 | 0.003331 | 0.55 | 0.6014 |
| (Running lost (% of baseline)-53.2008)*Manipulation[B_Autobahn] | 0.0007401 | 0.002896 | 0.26 | 0.8068 |

#### Effect Tests

| Source | Nparm | DF | Sum of Squares | F Ratio | Prob > F |
|---|---|---|---|---|---|
| Running lost (% of baseline) | 1 | 1 | 0.17634183 | 8.7066 | 0.0256* |
| Manipulation | 2 | 2 | 0.04818371 | 1.1895 | 0.3672 |
| Running lost (% of baseline)*Manipulation | 2 | 2 | 0.00766111 | 0.1891 | 0.8324 |

**Figure 4**

**4A**

**Contrast**

**Test Detail**

| | |
|---|---|
| A_Autobahn | 1 |
| B_Autobahn | -1 |
| C_Autobahn | 0 |
| Estimate | 5.7273 |
| Std Error | 4.3412 |
| t Ratio | 1.3193 |
| Prob>\|t\| | 0.2082 |
| SS | 58.428 |
| Lower 95% | -3.584 |
| Upper 95% | 15.038 |

| SS | NumDF | DenDF | F Ratio | Prob > F |
|---|---|---|---|---|
| 58.43 | 1 | 14 | 1.7405 | 0.2082 |

Running lost (% of baseline) 25

**Parameter Function**

**Contrast**

**Test Detail**

| | |
|---|---|
| A_Autobahn | 1 |
| B_Autobahn | 0 |
| C_Autobahn | -1 |
| Estimate | -1.549 |
| Std Error | 4.8081 |
| t Ratio | -0.322 |
| Prob>\|t\| | 0.7521 |
| SS | 3.4844 |
| Lower 95% | -11.86 |
| Upper 95% | 8.7632 |

| SS | NumDF | DenDF | F Ratio | Prob > F |
|---|---|---|---|---|
| 3.484 | 1 | 14 | 0.1038 | 0.7521 |

Running lost (% of baseline) 25

**Parameter Function**

**Contrast**

**Test Detail**

| | |
|---|---|
| A_Autobahn | 0 |
| B_Autobahn | 1 |
| C_Autobahn | -1 |
| Estimate | -7.276 |
| Std Error | 5.0642 |
| t Ratio | -1.437 |
| Prob>\|t\| | 0.1727 |
| SS | 69.3 |
| Lower 95% | -18.14 |
| Upper 95% | 3.5854 |

| SS | NumDF | DenDF | F Ratio | Prob > F |
|---|---|---|---|---|
| 69.3 | 1 | 14 | 2.0644 | 0.1727 |

Running lost (% of baseline) 25

**Parameter Function**

**Contrast**

**Test Detail**

| | |
|---|---|
| A_Autobahn | 1 |
| B_Autobahn | -1 |
| C_Autobahn | 0 |
| Estimate | 27.089 |
| Std Error | 5.2504 |
| t Ratio | 5.1595 |
| Prob>\|t\| | 0.0001 |
| SS | 893.61 |
| Lower 95% | 15.828 |
| Upper 95% | 38.35 |

| SS | NumDF | DenDF | F Ratio | Prob > F |
|---|---|---|---|---|
| 893.6 | 1 | 14 | 26.6202 | 0.0001* |

Running lost (% of baseline) 75

**Parameter Function**

**Contrast**

**Test Detail**

| | |
|---|---|
| A_Autobahn | 1 |
| B_Autobahn | 0 |
| C_Autobahn | -1 |
| Estimate | 27.591 |
| Std Error | 5.7977 |
| t Ratio | 4.759 |
| Prob>\|t\| | 0.0003 |
| SS | 760.27 |
| Lower 95% | 15.156 |
| Upper 95% | 40.026 |

| SS | NumDF | DenDF | F Ratio | Prob > F |
|---|---|---|---|---|
| 760.3 | 1 | 14 | 22.6481 | 0.0003* |

Running lost (% of baseline) 75

**Parameter Function**

**Contrast**

**Test Detail**

| | |
|---|---|
| A_Autobahn | 0 |
| B_Autobahn | 1 |
| C_Autobahn | -1 |
| Estimate | 0.5021 |
| Std Error | 5.1464 |
| t Ratio | 0.0976 |
| Prob>\|t\| | 0.9237 |
| SS | 0.3195 |
| Lower 95% | -10.54 |
| Upper 95% | 11.54 |

| SS | NumDF | DenDF | F Ratio | Prob > F |
|---|---|---|---|---|
| 0.319 | 1 | 14 | 0.0095 | 0.9237 |

Running lost (% of baseline) 75

**Parameter Function**

| | | | |
|---|---|---|---|
| 11d | Autobahn A | 25 | 18.69489 |
| 11d | Autobahn A | 75 | 52.66114 |
| 11d | Autobahn B | 25 | 12.9676 |
| 11d | Autobahn B | 75 | 25.57202 |
| 11d | Autobahn C | 25 | 20.24393 |
| 11d | Autobahn C | 75 | 25.06997 |

**4B**

| | | | | | |
|---|---|---|---|---|---|
| **Autobahn A** | | | | | |
| 44-49 dpc | Lost | Gain | | | |
| <30% loss | 24.39024 | 39.02439 | | | |
| 50-90% loss | 75 | 89.0625 | | | |
| | 64.83516 | 70.32967 | | | |
| | 56.52174 | 46.73913 | | > 50% loss | |
| | | | | paired t te | 0.686313 |
| | | | | unpaired t | 0.819321 |

**Figure 5**

**5C**

##### Oneway Analysis of Running lost (% of baseline) By Manipulation 2

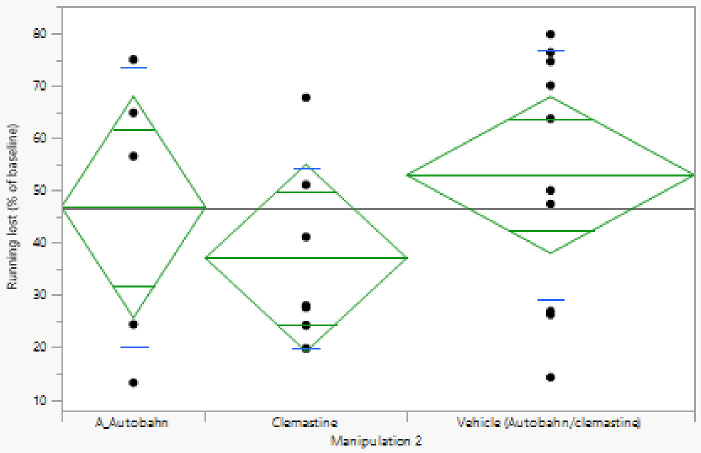

##### Oneway Anova

###### Summary of Fit

| | |
|---|---|
| Rsquare | 0.096689 |
| Adj Rsquare | 0.001603 |
| Root Mean Square Error | 22.60031 |
| Mean of Response | 46.49873 |
| Observations (or Sum Wgts) | 22 |

###### Analysis of Variance

| Source | DF | Sum of Squares | Mean Square | F Ratio | Prob > F |
|---|---|---|---|---|---|
| Manipulation 2 | 2 | 1038.772 | 519.386 | 1.0169 | 0.3806 |
| Error | 19 | 9704.705 | 510.774 | | |
| C. Total | 21 | 10743.477 | | | |

###### Means for Oneway Anova

| Level | Number | Mean | Std Error | Lower 95% | Upper 95% |
|---|---|---|---|---|---|
| A_Autobahn | 5 | 46.8000 | 10.107 | 25.645 | 67.955 |
| Clemastine | 7 | 37.0698 | 8.542 | 19.191 | 54.949 |
| Vehicle (Autobahn/clemastine) | 10 | 52.9484 | 7.147 | 37.990 | 67.907 |

Std Error uses a pooled estimate of error variance

##### Tests that the Variances are Equal

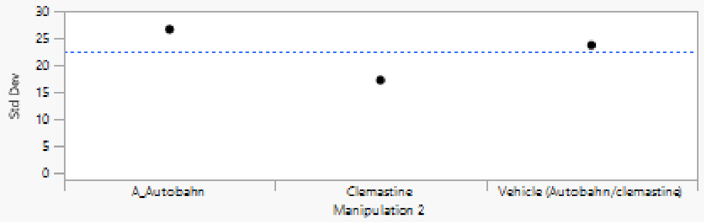

| Level | Count | Std Dev | MeanAbsDif to Mean | MeanAbsDif to Median |
|---|---|---|---|---|
| A_Autobahn | 5 | 26.65810 | 22.38272 | 20.43838 |
| Clemastine | 7 | 17.26694 | 13.91211 | 12.61244 |
| Vehicle (Autobahn/clemastine) | 10 | 23.74214 | 19.97089 | 19.97089 |

| Test | F Ratio | DFNum | DFDen | Prob > F |
|---|---|---|---|---|
| O'Brien[.5] | 1.0694 | 2 | 19 | 0.3630 |
| Brown-Forsythe | 0.6904 | 2 | 19 | 0.5135 |
| Levene | 1.2494 | 2 | 19 | 0.3092 |
| Bartlett | 0.4734 | 2 | . | 0.6229 |

Warning: Small sample sizes. Use Caution.

###### Welch's Test

Welch Anova testing Means Equal, allowing Std Devs Not Equal

| F Ratio | DFNum | DFDen | Prob > F |
|---|---|---|---|
| 1.2167 | 2 | 9.8481 | 0.3371 |

Excluded Rows 670

**5D**

#### Response Gain rate 0-11

##### Whole Model

###### Regression Plot

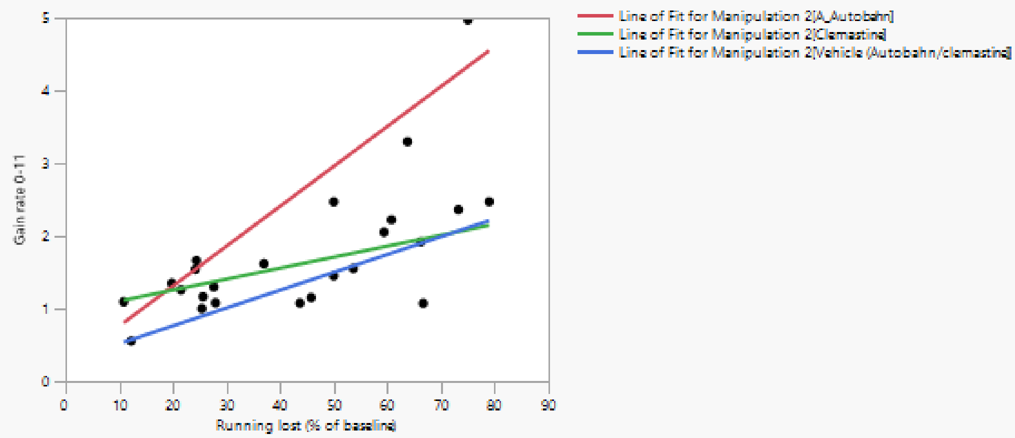

###### Actual by Predicted Plot

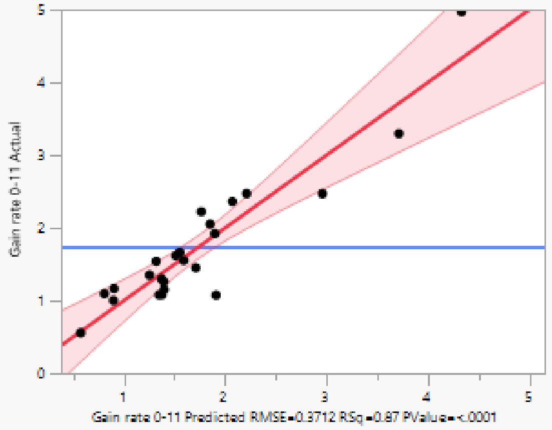

###### Effect Summary

| Source | Logworth | | PValue |
|---|---|---|---|
| Running lost (% of baseline) | 5.870 | | 0.00000 |
| Manipulation 2 | 4.830 | | 0.00001 |
| Running lost (% of baseline)*Manipulation 2 | 2.591 | | 0.00256 |

Remove Add Edit ☐ FDR

###### Residual by Predicted Plot

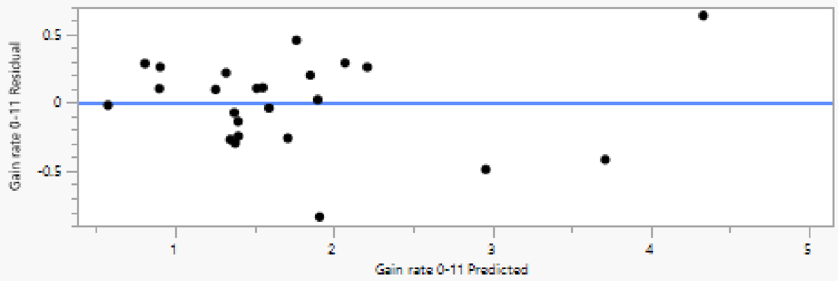

#### Summary of Fit

| | |
|---|---|
| RSquare | 0.874752 |
| RSquare Adj | 0.839961 |
| Root Mean Square Error | 0.371158 |
| Mean of Response | 1.737663 |
| Observations (or Sum Wgts) | 24 |

#### Analysis of Variance

| Source | DF | Sum of Squares | Mean Square | F Ratio |
|---|---|---|---|---|
| Model | 5 | 17.318282 | 3.46366 | 25.1431 |
| Error | 18 | 2.479643 | 0.13776 | Prob > F |
| C. Total | 23 | 19.797924 | | <.0001* |

#### Parameter Estimates

<table>
<tr><th>Term</th><th>Estimate</th><th>Std Error</th><th>t Ratio</th><th>Prob> |t|</th></tr>
<tr><td>Intercept</td><td>0.4793435</td><td>0.191962</td><td>2.50</td><td>0.0224*</td></tr>
<tr><td>Running lost (% of baseline)</td><td>0.0315311</td><td>0.004458</td><td>7.07</td><td><.0001*</td></tr>
<tr><td>Manipulation 2[A_Autobahn]</td><td>0.7498461</td><td>0.12202</td><td>6.15</td><td><.0001*</td></tr>
<tr><td>Manipulation 2[Clemastine]</td><td>-0.2397</td><td>0.127358</td><td>-1.88</td><td>0.0761</td></tr>
<tr><td>(Running lost (% of baseline)-43.486)*Manipulation 2[A_Autobahn]</td><td>0.023407</td><td>0.005806</td><td>4.03</td><td>0.0008*</td></tr>
<tr><td>(Running lost (% of baseline)-43.486)*Manipulation 2[Clemastine]</td><td>-0.016412</td><td>0.007478</td><td>-2.19</td><td>0.0416*</td></tr>
</table>

#### Effect Tests

| Source | Nparm | DF | Sum of Squares | F Ratio | Prob > F |
|---|---|---|---|---|---|
| Running lost (% of baseline) | 1 | 1 | 6.8928430 | 50.0359 | <.0001* |
| Manipulation 2 | 2 | 2 | 6.0527376 | 21.9687 | <.0001* |
| Running lost (% of baseline)*Manipulation 2 | 2 | 2 | 2.3318911 | 8.4637 | 0.0026* |

#### Manipulation 2

##### Leverage Plot

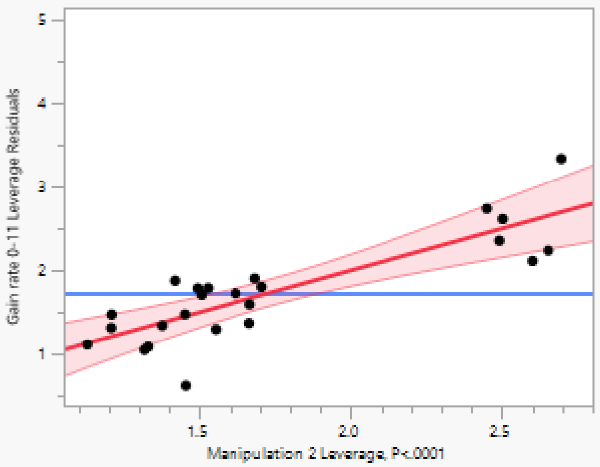

##### Least Squares Means Table

| Level | Least Sq Mean | Std Error | Mean |
|---|---|---|---|
| A_Autobahn | 2.6003510 | 0.15242518 | 2.45928 |
| Clemastine | 1.6108047 | 0.16500698 | 1.48451 |
| Vehicle (Autobahn/clemastine) | 1.3403590 | 0.11763643 | 1.50516 |

##### LSMeans Differences Tukey HSD

α= 0.050 Q= 2.55216

LSMean[j]

| LSMean[i] | Mean[i]-Mean[j] / Std Err Dif / Lower CL Dif / Upper CL Dif | A_Autobahn | Clemastine | Vehicle (Autobahn/clemastine) |
|---|---|---|---|---|
| A_Autobahn | Mean[i]-Mean[j] | 0 | 0.989546 | 1.259992 |
| | Std Err Dif | 0 | 0.224635 | 0.19254 |
| | Lower CL Dif | 0 | 0.416242 | 0.768598 |
| | Upper CL Dif | 0 | 1.562851 | 1.751386 |
| Clemastine | Mean[i]-Mean[j] | -0.98955 | 0 | 0.270446 |
| | Std Err Dif | 0.224635 | 0 | 0.202647 |
| | Lower CL Dif | -1.56285 | 0 | -0.24674 |
| | Upper CL Dif | -0.41624 | 0 | 0.787633 |
| Vehicle (Autobahn/clemastine) | Mean[i]-Mean[j] | -1.25999 | -0.27045 | 0 |
| | Std Err Dif | 0.19254 | 0.202647 | 0 |
| | Lower CL Dif | -1.75139 | -0.78763 | 0 |
| | Upper CL Dif | -0.7686 | 0.246741 | 0 |

| Level | | | Least Sq Mean |
|---|---|---|---|
| A_Autobahn | A | | 2.6003510 |
| Clemastine | | B | 1.6108047 |
| Vehicle (Autobahn/clemastine) | | B | 1.3403590 |

Levels not connected by same letter are significantly different.

| Level | - Level | Difference | Std Err Dif | Lower CL | Upper CL | p-Value |
|---|---|---|---|---|---|---|
| A_Autobahn | Vehicle (Autobahn/clemastine) | 1.259992 | 0.1925403 | 0.768598 | 1.751386 | <.0001* |
| A_Autobahn | Clemastine | 0.989546 | 0.2246347 | 0.416242 | 1.562851 | 0.0009* |
| Clemastine | Vehicle (Autobahn/clemastine) | 0.270446 | 0.2026466 | -0.246741 | 0.787633 | 0.3951 |

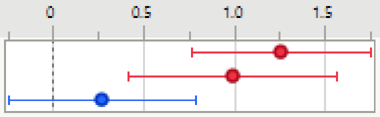

#### Response Gain rate 11-18

##### Whole Model

###### Regression Plot

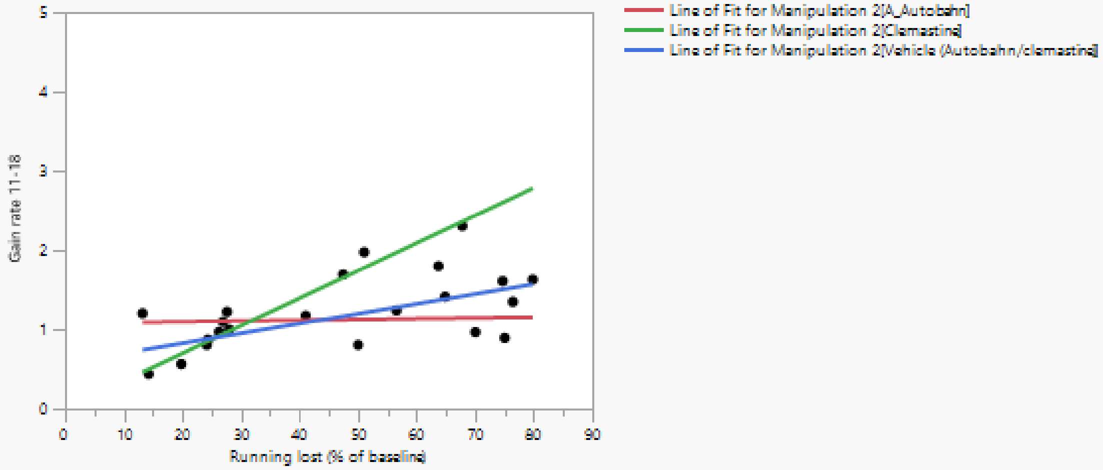

###### Actual by Predicted Plot

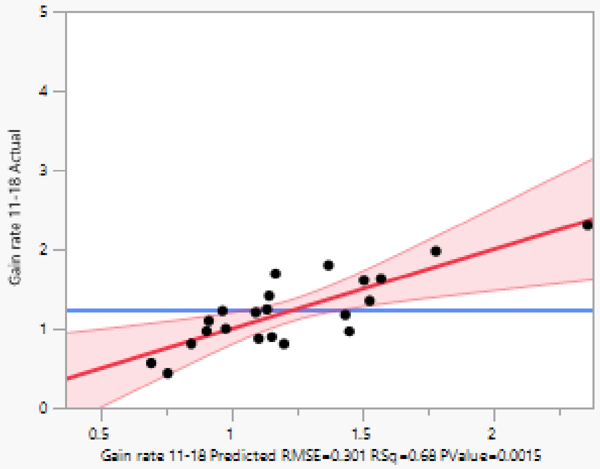

###### Effect Summary

| Source | Logworth | | PValue | |
|---|---|---|---|---|
| Running lost (% of baseline) | 3.717 | | 0.00019 | |
| Running lost (% of baseline)*Manipulation 2 | 2.175 | | 0.00669 | |
| Manipulation 2 | 1.628 | | 0.02353 | ^ |

Remove Add Edit ☐ FDR ('^' denotes effects with containing effects above them)

###### Residual by Predicted Plot

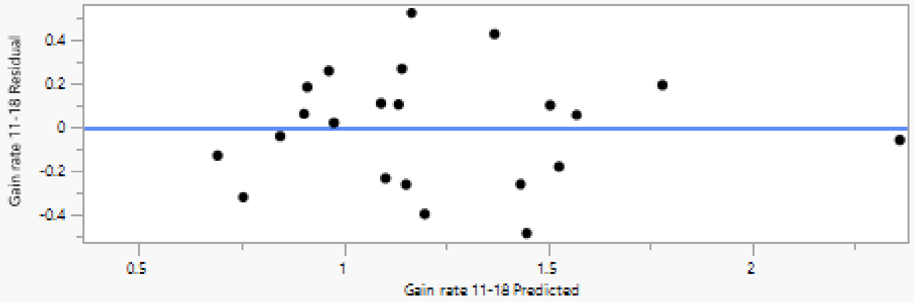

#### Summary of Fit

| | |
|---|---|
| RSquare | 0.67679 |
| RSquare Adj | 0.575787 |
| Root Mean Square Error | 0.300989 |
| Mean of Response | 1.228555 |
| Observations (or Sum Wgts) | 22 |

#### Analysis of Variance

| Source | DF | Sum of Squares | Mean Square | F Ratio |
|---|---|---|---|---|
| Model | 5 | 3.0352259 | 0.607045 | 6.7007 |
| Error | 16 | 1.4495139 | 0.090595 | Prob > F |
| C. Total | 21 | 4.4847398 | | 0.0015* |

#### Parameter Estimates

| Term | Estimate | Std Error | t Ratio | Prob>\|t\| |
|---|---|---|---|---|
| Intercept | 0.5530254 | 0.159715 | 3.46 | 0.0032* |
| Running lost (% of baseline) | 0.0160687 | 0.00334 | 4.81 | 0.0002* |
| Manipulation 2[A_Autobahn] | -0.175723 | 0.105269 | -1.67 | 0.1145 |
| Manipulation 2[Clemastine] | 0.3206442 | 0.104191 | 3.08 | 0.0072* |
| (Running lost (% of baseline)-46.4987)*Manipulation 2[A_Autobahn] | -0.015084 | 0.004666 | -3.23 | 0.0052* |
| (Running lost (% of baseline)-46.4987)*Manipulation 2[Clemastine] | 0.0187206 | 0.005295 | 3.54 | 0.0027* |

#### Effect Tests

| Source | Nparm | DF | Sum of Squares | F Ratio | Prob > F |
|---|---|---|---|---|---|
| Running lost (% of baseline) | 1 | 1 | 2.0974844 | 23.1524 | 0.0002* |
| Manipulation 2 | 2 | 2 | 0.8666943 | 4.7834 | 0.0235* |
| Running lost (% of baseline)*Manipulation 2 | 2 | 2 | 1.2609691 | 6.9594 | 0.0067* |

#### Manipulation 2

##### Leverage Plot

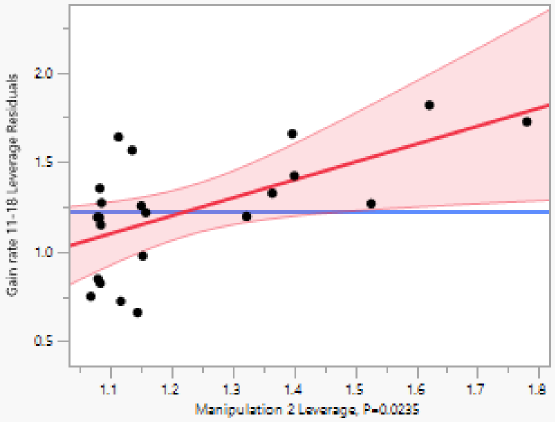

##### Least Squares Means Table

| Level | Least Sq Mean | Std Error | Mean |
|---|---|---|---|
| A_Autobahn | 1.1244765 | 0.13461730 | 1.12477 |
| Clemastine | 1.6208432 | 0.13207781 | 1.29282 |
| Vehicle (Autobahn/clemastine) | 1.1552774 | 0.09900653 | 1.23546 |

##### LSMeans Differences Tukey HSD

α= 0.050 Q= 2.58033

LSMean[j]

| LSMean[i] | Mean[i]-Mean[j] / Std Err Dif / Lower CL Dif / Upper CL Dif | A_Autobahn | Clemastine | Vehicle (Autobahn/clemastine) |
|---|---|---|---|---|
| A_Autobahn | Mean[i]-Mean[j] | 0 | -0.49637 | -0.0308 |
| | Std Err Dif | 0 | 0.18859 | 0.167105 |
| | Lower CL Dif | 0 | -0.98299 | -0.46199 |
| | Upper CL Dif | 0 | -0.00974 | 0.400386 |
| Clemastine | Mean[i]-Mean[j] | 0.496367 | 0 | 0.465566 |
| | Std Err Dif | 0.18859 | 0 | 0.165066 |
| | Lower CL Dif | 0.009741 | 0 | 0.03964 |
| | Upper CL Dif | 0.982993 | 0 | 0.891491 |
| Vehicle (Autobahn/clemastine) | Mean[i]-Mean[j] | 0.030801 | -0.46557 | 0 |
| | Std Err Dif | 0.167105 | 0.165066 | 0 |
| | Lower CL Dif | -0.40039 | -0.89149 | 0 |
| | Upper CL Dif | 0.461987 | -0.03964 | 0 |

| Level | | | Least Sq Mean |
|---|---|---|---|
| Clemastine | A | | 1.6208432 |
| Vehicle (Autobahn/clemastine) | | B | 1.1552774 |
| A_Autobahn | | B | 1.1244765 |

Levels not connected by same letter are significantly different.

| Level | - Level | Difference | Std Err Dif | Lower CL | Upper CL | p-Value |
|---|---|---|---|---|---|---|
| Clemastine | A_Autobahn | 0.4963667 | 0.1885905 | 0.009741 | 0.9829925 | 0.0453* |
| Clemastine | Vehicle (Autobahn/clemastine) | 0.4655658 | 0.1650662 | 0.039640 | 0.8914912 | 0.0313* |
| Vehicle (Autobahn/clemastine) | A_Autobahn | 0.0308009 | 0.1671051 | -0.400386 | 0.4619873 | 0.9815 |

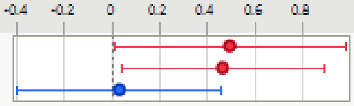

#### Response Gain rate 18-35

##### Whole Model

###### Regression Plot

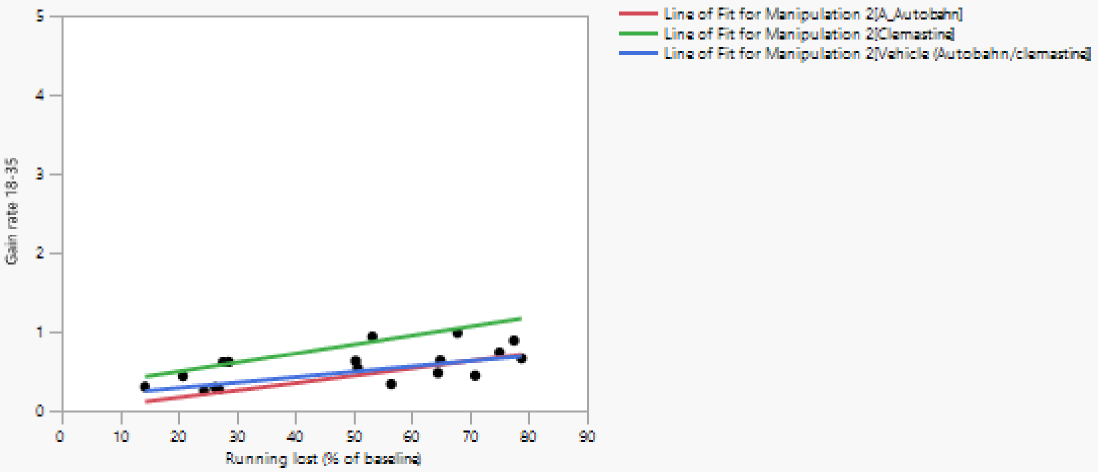

###### Actual by Predicted Plot

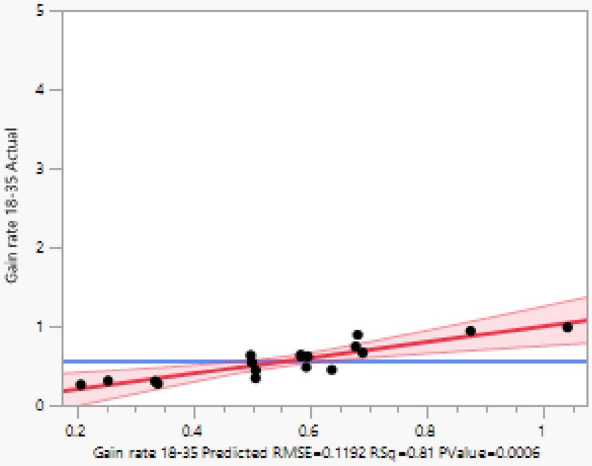

###### Effect Summary

| Source | Logworth | | PValue |
|---|---|---|---|
| Running lost (% of baseline) | 4.113 | | 0.00008 |
| Manipulation 2 | 3.058 | | 0.00087 |
| Running lost (% of baseline)*Manipulation 2 | 0.385 | | 0.41210 |

Remove Add Edit ☐ FDR

###### Residual by Predicted Plot

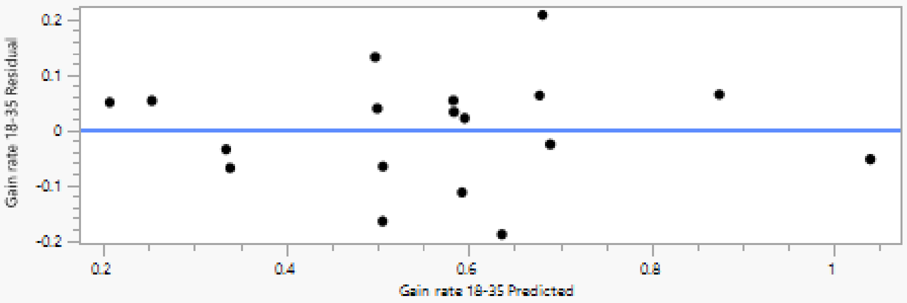

#### Summary of Fit

| | |
|---|---|
| RSquare | 0.807553 |
| RSquare Adj | 0.727366 |
| Root Mean Square Error | 0.119174 |
| Mean of Response | 0.560746 |
| Observations (or Sum Wgts) | 18 |

#### Analysis of Variance

| Source | DF | Sum of Squares | Mean Square | F Ratio |
|---|---|---|---|---|
| Model | 5 | 0.71515963 | 0.143032 | 10.0710 |
| Error | 12 | 0.17042909 | 0.014202 | Prob > F |
| C. Total | 17 | 0.88558872 | | 0.0006* |

#### Parameter Estimates

| Term | Estimate | Std Error | t Ratio | Prob>\|t\| |
|---|---|---|---|---|
| Intercept | 0.1354153 | 0.081763 | 1.66 | 0.1236 |
| Running lost (% of baseline) | 0.0091433 | 0.00156 | 5.86 | <.0001* |
| Manipulation 2[A_Autobahn] | -0.147796 | 0.048301 | -3.06 | 0.0099* |
| Manipulation 2[Clemastine] | 0.2427273 | 0.047069 | 5.16 | 0.0002* |
| (Running lost (% of baseline)-48.8252)*Manipulation 2[A_Autobahn] | 0.0001539 | 0.002394 | 0.06 | 0.9498 |
| (Running lost (% of baseline)-48.8252)*Manipulation 2[Clemastine] | 0.0022234 | 0.002325 | 0.96 | 0.3577 |

#### Effect Tests

| Source | Nparm | DF | Sum of Squares | F Ratio | Prob > F |
|---|---|---|---|---|---|
| Running lost (% of baseline) | 1 | 1 | 0.48790498 | 34.3536 | <.0001* |
| Manipulation 2 | 2 | 2 | 0.38065843 | 13.4012 | 0.0009* |
| Running lost (% of baseline)*Manipulation 2 | 2 | 2 | 0.02713611 | 0.9553 | 0.4121 |

#### Manipulation 2

##### Leverage Plot

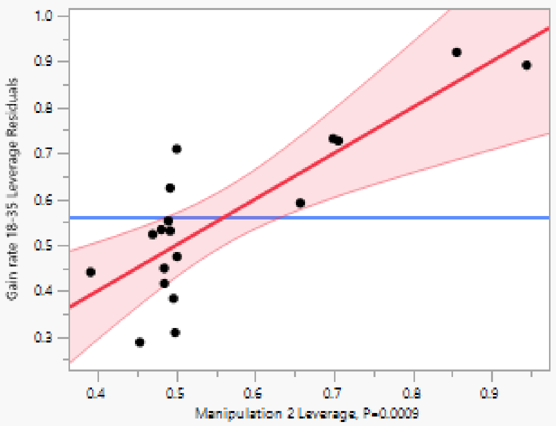

##### Least Squares Means Table

| Level | Least Sq Mean | Std Error | Mean |
|---|---|---|---|
| A_Autobahn | 0.43404253 | 0.06285622 | 0.493188 |
| Clemastine | 0.82456565 | 0.05998758 | 0.719742 |
| Vehicle (Autobahn/clemastine) | 0.48690694 | 0.03992961 | 0.502441 |

##### LSMeans Differences Tukey HSD

α= 0.050 Q= 2.66776

LSMean[j]

| LSMean[i] | Mean[i]-Mean[j] / Std Err Dif / Lower CL Dif / Upper CL Dif | A_Autobahn | Clemastine | Vehicle (Autobahn/clemastin |
|---|---|---|---|---|
| A_Autobahn | Mean[i]-Mean[j] | 0 | -0.39052 | -0.05286 |
| | Std Err Dif | 0 | 0.086887 | 0.074467 |
| | Lower CL Dif | 0 | -0.62232 | -0.25152 |
| | Upper CL Dif | 0 | -0.15873 | 0.145794 |
| Clemastine | Mean[i]-Mean[j] | 0.390523 | 0 | 0.337659 |
| | Std Err Dif | 0.086887 | 0 | 0.072062 |
| | Lower CL Dif | 0.158729 | 0 | 0.145416 |
| | Upper CL Dif | 0.622317 | 0 | 0.529902 |
| Vehicle (Autobahn/clemastine) | Mean[i]-Mean[j] | 0.052864 | -0.33766 | 0 |
| | Std Err Dif | 0.074467 | 0.072062 | 0 |
| | Lower CL Dif | -0.14579 | -0.5299 | 0 |
| | Upper CL Dif | 0.251523 | -0.14542 | 0 |

| Level | | | Least Sq Mean |
|---|---|---|---|
| Clemastine | A | | 0.82456565 |
| Vehicle (Autobahn/clemastine) | | B | 0.48690694 |
| A_Autobahn | | B | 0.43404253 |

Levels not connected by same letter are significantly different.

| Level | - Level | Difference | Std Err Dif | Lower CL | Upper CL | p-Value |
|---|---|---|---|---|---|---|
| Clemastine | A_Autobahn | 0.3905231 | 0.0868874 | 0.158729 | 0.6223175 | 0.0020* |
| Clemastine | Vehicle (Autobahn/clemastine) | 0.3376587 | 0.0720617 | 0.145416 | 0.5299017 | 0.0014* |
| Vehicle (Autobahn/clemastine) | A_Autobahn | 0.0528644 | 0.0744666 | -0.145794 | 0.2515233 | 0.7624 |

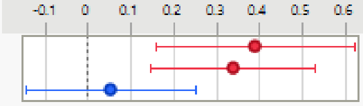

**5E**

#### Response Max gain rate

##### Whole Model

###### Regression Plot

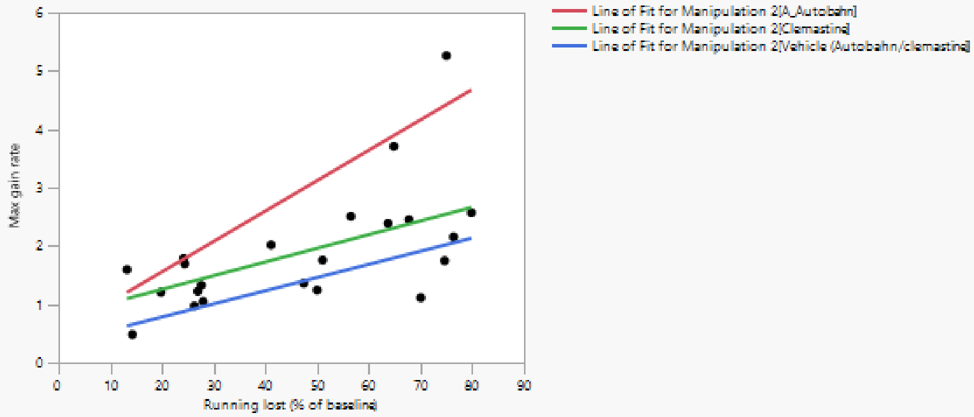

###### Actual by Predicted Plot

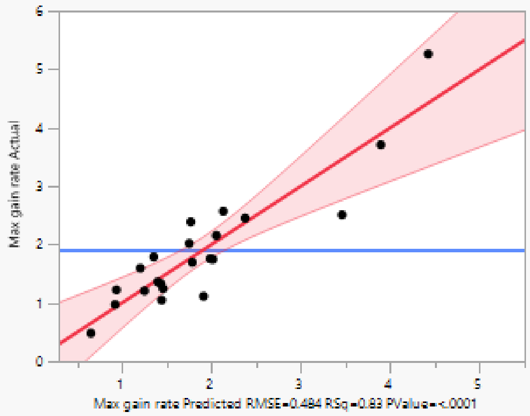

###### Effect Summary

| Source | Logworth | PValue |
|---|---|---|
| Running lost (% of baseline) | 4.808 | 0.00002 |
| Manipulation 2 | 3.934 | 0.00012 |
| Running lost (% of baseline)*Manipulation 2 | 1.317 | 0.04822 |

Remove Add Edit ☐ FDR

###### Residual by Predicted Plot

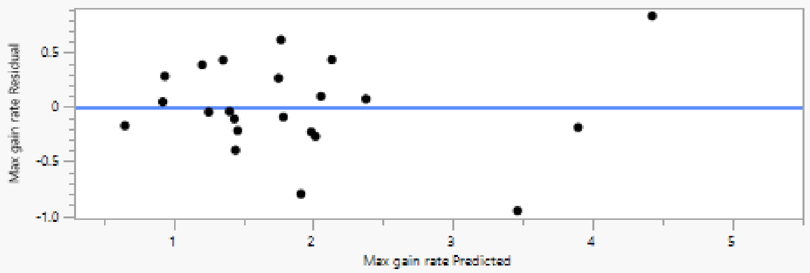

#### Summary of Fit

| | |
|---|---|
| RSquare | 0.832698 |
| RSquare Adj | 0.780417 |
| Root Mean Square Error | 0.484028 |
| Mean of Response | 1.891915 |
| Observations (or Sum Wgts) | 22 |

#### Analysis of Variance

| Source | DF | Sum of Squares | Mean Square | F Ratio |
|---|---|---|---|---|
| Model | 5 | 18.657268 | 3.73145 | 15.9271 |
| Error | 16 | 3.748523 | 0.23428 | **Prob > F** |
| C. Total | 21 | 22.405791 | | <.0001* |

#### Parameter Estimates

| Term | Estimate | Std Error | t Ratio | Prob>\|t\| |
|---|---|---|---|---|
| Intercept | 0.5438426 | 0.256841 | 2.12 | 0.0502 |
| Running lost (% of baseline) | 0.0327205 | 0.00537 | 6.09 | <.0001* |
| Manipulation 2[A_Autobahn] | 0.8725512 | 0.169285 | 5.15 | <.0001* |
| Manipulation 2[Clemastine] | -0.187192 | 0.167552 | -1.12 | 0.2804 |
| (Running lost (% of baseline)-46.4987)*Manipulation 2[A_Autobahn] | 0.019398 | 0.007504 | 2.58 | 0.0199* |
| (Running lost (% of baseline)-46.4987)*Manipulation 2[Clemastine] | -0.009261 | 0.008514 | -1.09 | 0.2929 |

#### Effect Tests

| Source | Nparm | DF | Sum of Squares | F Ratio | Prob > F |
|---|---|---|---|---|---|
| Running lost (% of baseline) | 1 | 1 | 8.6971520 | 37.1225 | <.0001* |
| Manipulation 2 | 2 | 2 | 7.8832593 | 16.8242 | 0.0001* |
| Running lost (% of baseline)*Manipulation 2 | 2 | 2 | 1.7273550 | 3.6865 | 0.0482* |

#### Manipulation 2

##### Leverage Plot

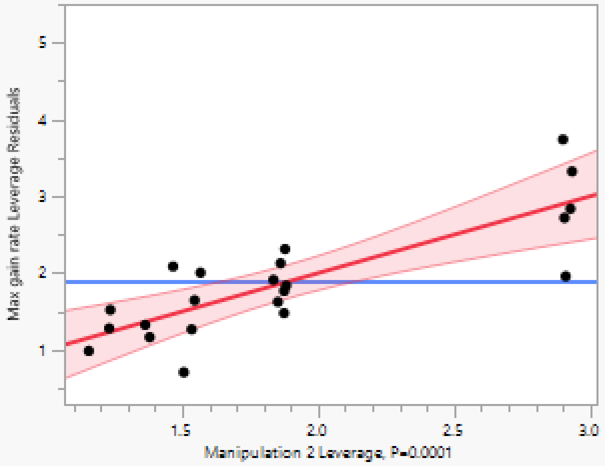

##### Least Squares Means Table

| Level | Least Sq Mean | Std Error | Mean |
|---|---|---|---|
| A_Autobahn | 2.9378543 | 0.21648100 | 2.95356 |
| Clemastine | 1.8781114 | 0.21239719 | 1.65691 |
| Vehicle (Autobahn/clemastine) | 1.3799438 | 0.15921456 | 1.52560 |

##### LSMeans Differences Tukey HSD

α= 0.050 Q= 2.58033

LSMean[j]

| Mean[i]-Mean[j]<br>Std Err Dif<br>Lower CL Dif<br>Upper CL Dif | A_Autobahn | Clemastine | Vehicle (Autobahn/clemastin |
|---|---|---|---|
| A_Autobahn | 0 | 1.059743 | 1.557911 |
| | 0 | 0.303276 | 0.268725 |
| | 0 | 0.277189 | 0.86451 |
| | 0 | 1.842297 | 2.251311 |
| Clemastine | -1.05974 | 0 | 0.498168 |
| | 0.303276 | 0 | 0.265446 |
| | -1.8423 | 0 | -0.18677 |
| | -0.27719 | 0 | 1.183107 |
| Vehicle (Autobahn/clemastine) | -1.55791 | -0.49817 | 0 |
| | 0.268725 | 0.265446 | 0 |
| | -2.25131 | -1.18311 | 0 |
| | -0.86451 | 0.186772 | 0 |

LSMean[i]

| Level | | | Least Sq Mean |
|---|---|---|---|
| A_Autobahn | A | | 2.9378543 |
| Clemastine | | B | 1.8781114 |
| Vehicle (Autobahn/clemastine) | | B | 1.3799438 |

Levels not connected by same letter are significantly different.

| Level | - Level | Difference | Std Err Dif | Lower CL | Upper CL | p-Value |
|---|---|---|---|---|---|---|
| A_Autobahn | Vehicle (Autobahn/clemastine) | 1.557911 | 0.2687253 | 0.864510 | 2.251311 | <.0001* |
| A_Autobahn | Clemastine | 1.059743 | 0.3032764 | 0.277189 | 1.842297 | 0.0080* |
| Clemastine | Vehicle (Autobahn/clemastine) | 0.498168 | 0.2654465 | -0.186772 | 1.183107 | 0.1776 |

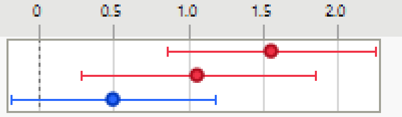

**5F**

#### Response Running gain (% of baseline)

##### Regression Plot

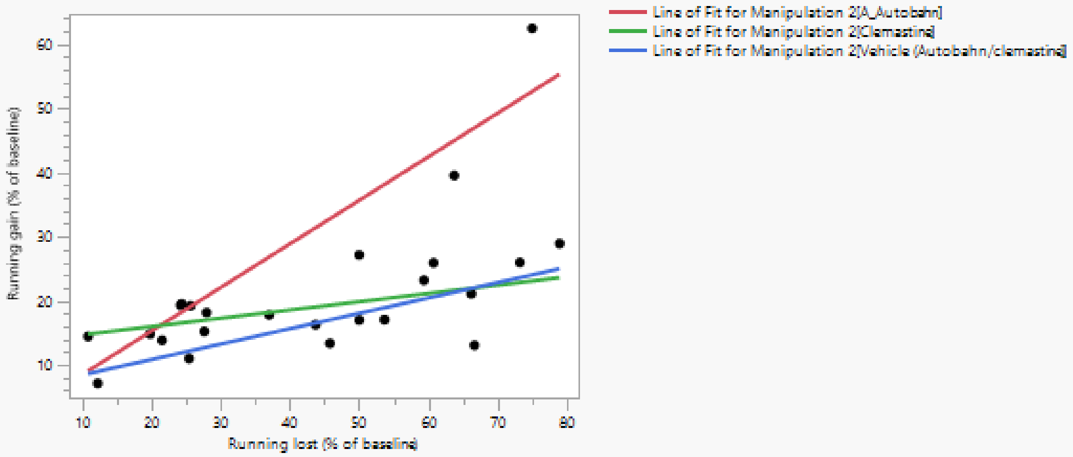

##### Effect Summary

| Source | Logworth | | PValue |
|---|---|---|---|
| Running lost (% of baseline) | 4.712 | | 0.00002 |
| Manipulation 2 | 4.052 | | 0.00009 |
| Manipulation 2*Running lost (% of baseline) | 2.738 | | 0.00183 |

Remove Add Edit FDR

##### Summary of Fit

| | |
|---|---|
| RSquare | 0.839095 |
| RSquare Adj | 0.794399 |
| Root Mean Square Error | 5.072984 |
| Mean of Response | 20.91404 |
| Observations (or Sum Wgts) | 24 |

##### Analysis of Variance

| Source | DF | Sum of Squares | Mean Square | F Ratio |
|---|---|---|---|---|
| Model | 5 | 2415.6869 | 483.137 | 18.7734 |
| Error | 18 | 463.2331 | 25.735 | Prob > F |
| C. Total | 23 | 2878.9200 | | <.0001* |

##### Parameter Estimates

| Term | Estimate | Std Error | t Ratio | Prob>\|t\| |
|---|---|---|---|---|
| Intercept | 7.0624843 | 2.623731 | 2.69 | 0.0149* |
| Manipulation 2[A_Autobahn] | 8.9914153 | 1.667767 | 5.39 | <.0001* |
| Manipulation 2[Clemastine] | -3.227536 | 1.740733 | -1.85 | 0.0802 |
| Running lost (% of baseline) | 0.3495142 | 0.060926 | 5.74 | <.0001* |
| Manipulation 2[A_Autobahn]*(Running lost (% of baseline)-43.486) | 0.3298109 | 0.079356 | 4.16 | 0.0006* |
| Manipulation 2[Clemastine]*(Running lost (% of baseline)-43.486) | -0.220297 | 0.102213 | -2.16 | 0.0449* |

##### Effect Tests

| Source | Nparm | DF | Sum of Squares | F Ratio | Prob > F |
|---|---|---|---|---|---|
| Manipulation 2 | 2 | 2 | 842.92267 | 16.3769 | <.0001* |
| Running lost (% of baseline) | 1 | 1 | 846.93297 | 32.9096 | <.0001* |
| Manipulation 2*Running lost (% of baseline) | 2 | 2 | 469.99150 | 9.1313 | 0.0018* |

#### Effect Details

##### Manipulation 2

###### Least Squares Means Table

| Level | Least Sq Mean | Std Error | Mean |
|---|---|---|---|
| A_Autobahn | 31.252860 | 2.0833485 | 29.5084 |
| Clemastine | 19.033908 | 2.2553165 | 17.9545 |
| Vehicle (Autobahn/clemastine) | 16.497566 | 1.6078555 | 18.1095 |

###### LSMeans Differences Tukey HSD

α= 0.050 Q= 2.55216

LSMean[j]

| LSMean[i] | Mean[i]-Mean[j]<br>Std Err Dif<br>Lower CL Dif<br>Upper CL Dif | A_Autobahn | Clemastine | Vehicle (Autobahn/clemastin |
|---|---|---|---|---|
| A_Autobahn | | 0 | 12.21895 | 14.75529 |
| | | 0 | 3.070308 | 2.631642 |
| | | 0 | 4.383023 | 8.038913 |
| | | 0 | 20.05488 | 21.47168 |
| Clemastine | | -12.219 | 0 | 2.536342 |
| | | 3.070308 | 0 | 2.769775 |
| | | -20.0549 | 0 | -4.53258 |
| | | -4.38302 | 0 | 9.60526 |
| Vehicle (Autobahn/clemastine) | | -14.7553 | -2.53634 | 0 |
| | | 2.631642 | 2.769775 | 0 |
| | | -21.4717 | -9.60526 | 0 |
| | | -8.03891 | 4.532575 | 0 |

| Level | | | Least Sq Mean |
|---|---|---|---|
| A_Autobahn | A | | 31.252860 |
| Clemastine | | B | 19.033908 |
| Vehicle (Autobahn/clemastine) | | B | 16.497566 |

Levels not connected by same letter are significantly different.

| Level | - Level | Difference | Std Err Dif | Lower CL | Upper CL | p-Value |
|---|---|---|---|---|---|---|
| A_Autobahn | Vehicle (Autobahn/clemastine) | 14.75529 | 2.631642 | 8.03891 | 21.47168 | <.0001* |
| A_Autobahn | Clemastine | 12.21895 | 3.070308 | 4.38302 | 20.05488 | 0.0024* |
| Clemastine | Vehicle (Autobahn/clemastine) | 2.53634 | 2.769775 | -4.53258 | 9.60526 | 0.6376 |

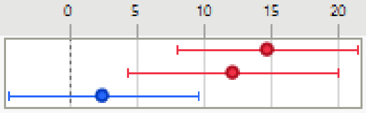

**Figure 6**

**6D**

##### Oneway Analysis of Latency_combined By mouse_condition

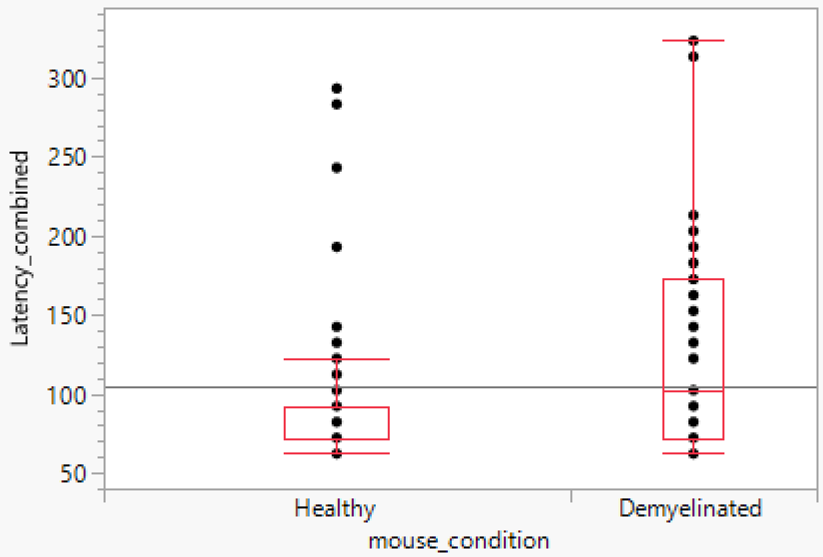

##### Quantiles

| Level | Minimum | 10% | 25% | Median | 75% | 90% | Maximum |
|---|---|---|---|---|---|---|---|
| Healthy | 62.20884 | 62.20884 | 72.249 | 72.249 | 92.32932 | 142.5301 | 293.1325 |
| Demyelinated | 62.20884 | 62.20884 | 72.249 | 102.3695 | 172.6506 | 206.7871 | 323.253 |

##### Wilcoxon / Kruskal-Wallis Tests (Rank Sums)

| Level | Count | Score Sum | Expected Score | Score Mean | (Mean-Mean0)/Std0 |
|---|---|---|---|---|---|
| Healthy | 104 | 7122.00 | 8320.00 | 68.481 | -4.371 |
| Demyelinated | 55 | 5598.00 | 4400.00 | 101.782 | 4.371 |

###### Wilcoxon Two-Sample Test, Normal Approximation

| S | Z | Prob>\|Z\| |
|---|---|---|
| 5598 | 4.37145 | <.0001* |

###### Kruskal-Wallis Test, ChiSquare Approximation

| ChiSquare | DF | Prob>ChiSq |
|---|---|---|
| 19.1255 | 1 | <.0001* |

Missing Rows 1921

Excluded Rows 7252

**6E**

#### Oneway Analysis of Latency_combined By mouse_condition

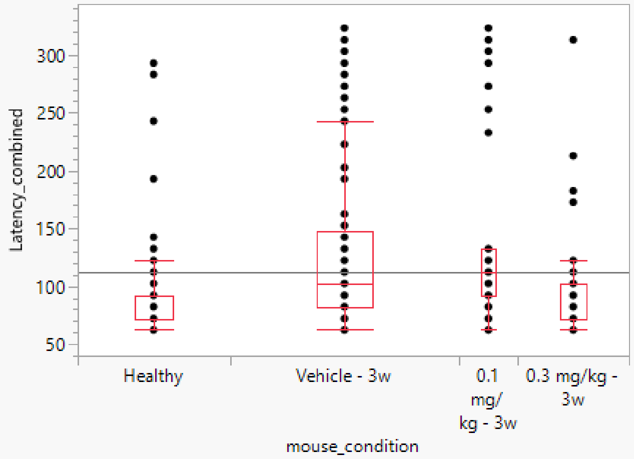

##### Quantiles

| Level | Minimum | 10% | 25% | Median | 75% | 90% | Maximum |
|---|---|---|---|---|---|---|---|
| Healthy | 62.20884 | 62.20884 | 72.249 | 72.249 | 92.32932 | 142.5301 | 293.1325 |
| Vehicle - 3w | 62.20884 | 62.20884 | 82.28916 | 102.3695 | 147.5502 | 244.9398 | 323.253 |
| 0.1 mg/kg - 3w | 62.20884 | 73.25301 | 92.32932 | 112.4096 | 132.49 | 302.1687 | 323.253 |
| 0.3 mg/kg - 3w | 62.20884 | 62.20884 | 72.249 | 72.249 | 102.3695 | 172.6506 | 313.2129 |

##### Wilcoxon / Kruskal-Wallis Tests (Rank Sums)

| Level | Count | Score Sum | Expected Score | Score Mean | (Mean-Mean0)/Std0 |
|---|---|---|---|---|---|
| Healthy | 104 | 15027.0 | 19656.0 | 144.490 | -4.945 |
| Vehicle - 3w | 157 | 33915.0 | 29673.0 | 216.019 | 4.108 |
| 0.1 mg/kg - 3w | 40 | 9867.00 | 7560.00 | 246.675 | 3.576 |
| 0.3 mg/kg - 3w | 76 | 12444.0 | 14364.0 | 163.737 | -2.285 |

###### Kruskal-Wallis Test, ChiSquare Approximation

| ChiSquare | DF | Prob>ChiSq |
|---|---|---|
| 43.1675 | 3 | <.0001* |

##### Nonparametric Comparisons For All Pairs Using Steel-Dwass Method

| q* | Alpha |
|---|---|
| 2.56903 | 0.05 |

| Level | - Level | Score Mean Difference | Std Err Dif | Z | p-Value | Hodges-Lehmann | Lower CL | Upper CL | Difference Plot |
|---|---|---|---|---|---|---|---|---|---|
| Vehicle - 3w | Healthy | 48.5698 | 9.45418 | 5.13740 | <.0001* | 20.0803 | 10.0402 | 30.1205 | |
| 0.1 mg/kg - 3w | Healthy | 38.6827 | 7.65583 | 5.05271 | <.0001* | 30.1205 | 20.0803 | 40.1606 | |
| 0.1 mg/kg - 3w | Vehicle - 3w | 15.3867 | 10.04298 | 1.53209 | 0.4182 | 10.0402 | -10.0402 | 30.1205 | |
| 0.3 mg/kg - 3w | Healthy | 10.7376 | 7.63045 | 1.40720 | 0.4948 | 0.0000 | 0.0000 | 10.0402 | |
| 0.3 mg/kg - 3w | 0.1 mg/kg - 3w | -26.6151 | 6.46798 | -4.11490 | 0.0002* | -20.0803 | -40.1606 | -10.0402 | |
| 0.3 mg/kg - 3w | Vehicle - 3w | -33.0695 | 9.33466 | -3.54266 | 0.0022* | -10.0402 | -30.1205 | 0.0000 | |

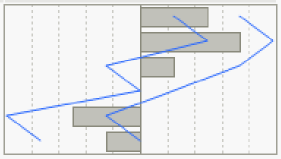

Missing Rows 4377

Excluded Rows 4578

**6G**

#### Oneway Analysis of n70_latency (ms) By condition

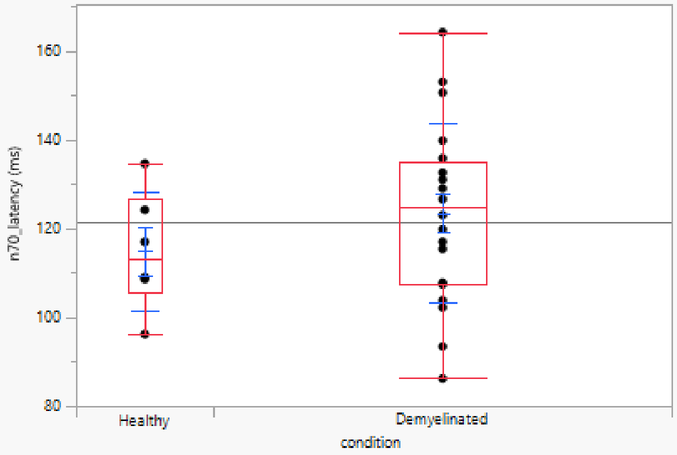

##### Quantiles

| Level | Minimum | 10% | 25% | Median | 75% | 90% | Maximum |
|---|---|---|---|---|---|---|---|
| Healthy | 96.11845 | 96.11845 | 105.4222 | 112.9252 | 126.7307 | 134.5338 | 134.5338 |
| Demyelinated | 86.11445 | 94.19768 | 107.423 | 124.7299 | 134.934 | 152.7011 | 164.1457 |

##### Means and Std Deviations

| Level | Number | Mean | Std Dev | Std Err Mean | Lower 95% | Upper 95% | Std Dev Lower 95% | Std Dev Upper 95% |
|---|---|---|---|---|---|---|---|---|
| Healthy | 6 | 114.85928 | 13.445418 | 5.4890688 | 100.74918 | 128.96938 | 8.3927374 | 32.97642 |
| Demyelinated | 20 | 123.40936 | 20.112865 | 4.4973734 | 113.99625 | 132.82247 | 15.295641 | 29.37628 |

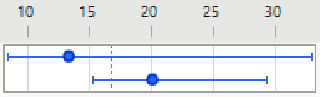

##### Wilcoxon / Kruskal-Wallis Tests (Rank Sums)

| Level | Count | Score Sum | Expected Score | Score Mean | (Mean-Mean0)/Std0 |
|---|---|---|---|---|---|
| Healthy | 6 | 67.500 | 81.000 | 11.2500 | -0.791 |
| Demyelinated | 20 | 283.500 | 270.000 | 14.1750 | 0.791 |

###### Wilcoxon Two-Sample Test, Normal Approximation

| S | Z | Prob>\|Z\| |
|---|---|---|
| 67.5 | -0.79143 | 0.4287 |

###### Kruskal-Wallis Test, ChiSquare Approximation

| ChiSquare | DF | Prob>ChiSq |
|---|---|---|
| 0.6755 | 1 | 0.4112 |

Excluded Rows 118

## 6H

##### Oneway Analysis of n70_latency (ms) By condition

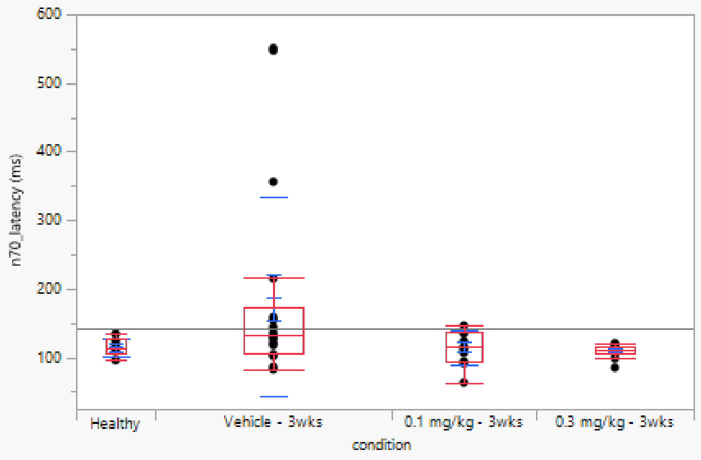

##### Quantiles

| Level | Minimum | 10% | 25% | Median | 75% | 90% | Maximum |
|---|---|---|---|---|---|---|---|
| Healthy | 96.11845 | 96.11845 | 105.4222 | 112.9252 | 126.7307 | 134.5338 | 134.5338 |
| Vehicle - 3wks | 82.91317 | 86.15446 | 105.122 | 133.5334 | 173.0492 | 547.459 | 550.7003 |
| 0.1 mg/kg - 3wks | 63.70548 | 69.30772 | 94.11765 | 115.7263 | 137.3349 | 145.018 | 146.5386 |
| 0.3 mg/kg - 3wks | 85.31413 | 89.39576 | 106.2225 | 111.7247 | 116.8267 | 120.3281 | 120.9284 |

##### Means and Std Deviations

| Level | Number | Mean | Std Dev | Std Err Mean | Lower 95% | Upper 95% | Std Dev Lower 95% | Std Dev Upper 95% |
|---|---|---|---|---|---|---|---|---|
| Healthy | 6 | 114.85928 | 13.445418 | 5.4890688 | 100.74918 | 128.96938 | 8.3927374 | 32.97642 |
| Vehicle - 3wks | 18 | 188.13303 | 144.88112 | 34.148808 | 116.08534 | 260.18072 | 108.717 | 217.19766 |
| 0.1 mg/kg - 3wks | 11 | 115.07148 | 24.946553 | 7.5216686 | 98.312161 | 131.83081 | 17.430582 | 43.779542 |
| 0.3 mg/kg - 3wks | 12 | 109.75724 | 9.9221187 | 2.864269 | 103.45302 | 116.06145 | 7.0287808 | 16.846547 |

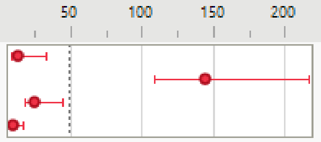

##### Wilcoxon / Kruskal-Wallis Tests (Rank Sums)

| Level | Count | Score Sum | Expected Score | Score Mean | (Mean-Mean0)/Std0 |
|---|---|---|---|---|---|
| Healthy | 6 | 129.500 | 144.000 | 21.5833 | -0.446 |
| Vehicle - 3wks | 18 | 538.000 | 432.000 | 29.8889 | 2.309 |
| 0.1 mg/kg - 3wks | 11 | 247.000 | 264.000 | 22.4545 | -0.415 |
| 0.3 mg/kg - 3wks | 12 | 213.500 | 288.000 | 17.7917 | -1.806 |

###### Kruskal-Wallis Test, ChiSquare Approximation

| ChiSquare | DF | Prob>ChiSq |
|---|---|---|
| 6.1077 | 3 | 0.1065 |

Excluded Rows 97

**6I**

#### Oneway Analysis of correlation_baseline By condition

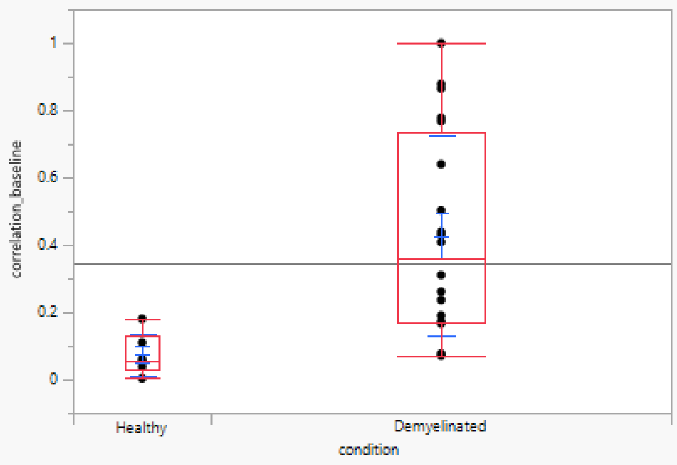

##### Quantiles

| Level | Minimum | 10% | 25% | Median | 75% | 90% | Maximum |
|---|---|---|---|---|---|---|---|
| Healthy | 0.004105 | 0.004105 | 0.029768 | 0.056664 | 0.128007 | 0.180706 | 0.180706 |
| Demyelinated | 0.071835 | 0.086583 | 0.169594 | 0.359778 | 0.735754 | 0.876869 | 0.999668 |

##### Means and Std Deviations

| Level | Number | Mean | Std Dev | Std Err Mean | Lower 95% | Upper 95% | Std Dev Lower 95% | Std Dev Upper 95% |
|---|---|---|---|---|---|---|---|---|
| Healthy | 6 | 0.0744834 | 0.0624481 | 0.0254943 | 0.0089481 | 0.1400186 | 0.0389806 | 0.153161 |
| Demyelinated | 20 | 0.4267095 | 0.2957816 | 0.0661388 | 0.2882795 | 0.5651396 | 0.2249391 | 0.4320102 |

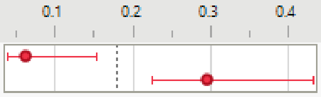

##### Wilcoxon / Kruskal-Wallis Tests (Rank Sums)

| Level | Count | Score Sum | Expected Score | Score Mean | (Mean-Mean0)/Std0 |
|---|---|---|---|---|---|
| Healthy | 6 | 29.000 | 81.000 | 4.8333 | -3.134 |
| Demyelinated | 20 | 322.000 | 270.000 | 16.1000 | 3.134 |

###### Wilcoxon Two-Sample Test, Normal Approximation

| S | Z | Prob>\|Z\| |
|---|---|---|
| 29 | -3.13419 | 0.0017* |

###### Kruskal-Wallis Test, ChiSquare Approximation

| ChiSquare | DF | Prob>ChiSq |
|---|---|---|
| 10.0148 | 1 | 0.0016* |

Excluded Rows 118

**6J**

#### Oneway Analysis of correlation_baseline By condition

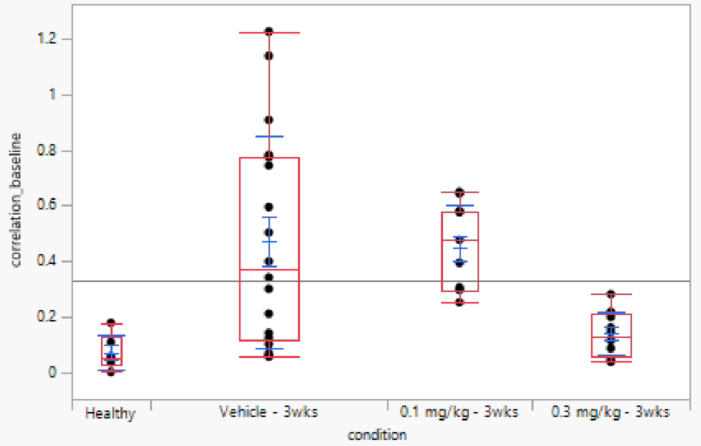

##### Quantiles

| Level | Minimum | 10% | 25% | Median | 75% | 90% | Maximum |
|---|---|---|---|---|---|---|---|
| Healthy | 0.004105 | 0.004105 | 0.029768 | 0.056664 | 0.128007 | 0.180706 | 0.180706 |
| Vehicle - 3wks | 0.059003 | 0.061838 | 0.120903 | 0.372192 | 0.776546 | 1.146616 | 1.225352 |
| 0.1 mg/kg - 3wks | 0.253301 | 0.253762 | 0.297566 | 0.477763 | 0.581293 | 0.649025 | 0.650203 |
| 0.3 mg/kg - 3wks | 0.040074 | 0.042838 | 0.059533 | 0.133317 | 0.211771 | 0.264369 | 0.283427 |

##### Means and Std Deviations

| Level | Number | Mean | Std Dev | Std Err Mean | Lower 95% | Upper 95% | Std Dev Lower 95% | Std Dev Upper 95% |
|---|---|---|---|---|---|---|---|---|
| Healthy | 6 | 0.0744834 | 0.0624481 | 0.0254943 | 0.0089481 | 0.1400186 | 0.0389806 | 0.153161 |
| Vehicle - 3wks | 18 | 0.4721251 | 0.3803425 | 0.0896476 | 0.2829852 | 0.661265 | 0.2854043 | 0.5701881 |
| 0.1 mg/kg - 3wks | 11 | 0.4470598 | 0.1536102 | 0.0463152 | 0.343863 | 0.5502565 | 0.1073301 | 0.2695757 |
| 0.3 mg/kg - 3wks | 12 | 0.1412302 | 0.0779792 | 0.0225107 | 0.0916846 | 0.1907759 | 0.0552401 | 0.1323992 |

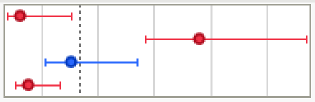

##### Wilcoxon / Kruskal-Wallis Tests (Rank Sums)

| Level | Count | Score Sum | Expected Score | Score Mean | (Mean-Mean0)/Std0 |
|---|---|---|---|---|---|
| Healthy | 6 | 50.000 | 144.000 | 8.3333 | -2.981 |
| Vehicle - 3wks | 18 | 528.000 | 432.000 | 29.3333 | 2.090 |
| 0.1 mg/kg - 3wks | 11 | 366.000 | 264.000 | 33.2727 | 2.550 |
| 0.3 mg/kg - 3wks | 12 | 184.000 | 288.000 | 15.3333 | -2.525 |

###### Kruskal-Wallis Test, ChiSquare Approximation

| ChiSquare | DF | Prob>ChiSq |
|---|---|---|
| 20.3820 | 3 | 0.0001* |

##### Nonparametric Comparisons For All Pairs Using Steel-Dwass Method

| q* | Alpha |
|---|---|
| 2.56903 | 0.05 |

| Level | - Level | Score Mean Difference | Std Err Dif | Z | p-Value | Hodges-Lehmann | Lower CL | Upper CL |
|---|---|---|---|---|---|---|---|---|
| Vehicle - 3wks | Healthy | 9.4444 | 3.333333 | 2.83333 | 0.0238* | 0.301618 | 0.009145 | 0.869803 |
| 0.1 mg/kg - 3wks | Healthy | 8.3712 | 2.562846 | 3.26637 | 0.0060* | 0.379547 | 0.145163 | 0.591304 |
| 0.3 mg/kg - 3wks | Healthy | 4.3750 | 2.669270 | 1.63903 | 0.3566 | 0.061800 | -0.060939 | 0.181579 |
| 0.1 mg/kg - 3wks | Vehicle - 3wks | 0.6591 | 3.258633 | 0.20226 | 0.9971 | 0.053461 | -0.430074 | 0.336584 |
| 0.3 mg/kg - 3wks | Vehicle - 3wks | -7.9861 | 3.280837 | -2.43417 | 0.0708 | -0.252655 | -0.686310 | 0.015979 |
| 0.3 mg/kg - 3wks | 0.1 mg/kg - 3wks | -11.0644 | 2.831104 | -3.90815 | 0.0005* | -0.302565 | -0.486822 | -0.134185 |

Difference Plot

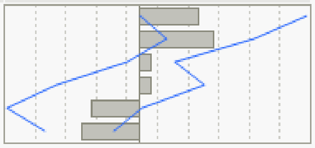

Excluded Rows 97

**Figure 7**

**7A**

**Oneway Analysis of Latency_delay_combined By mouse_condition**

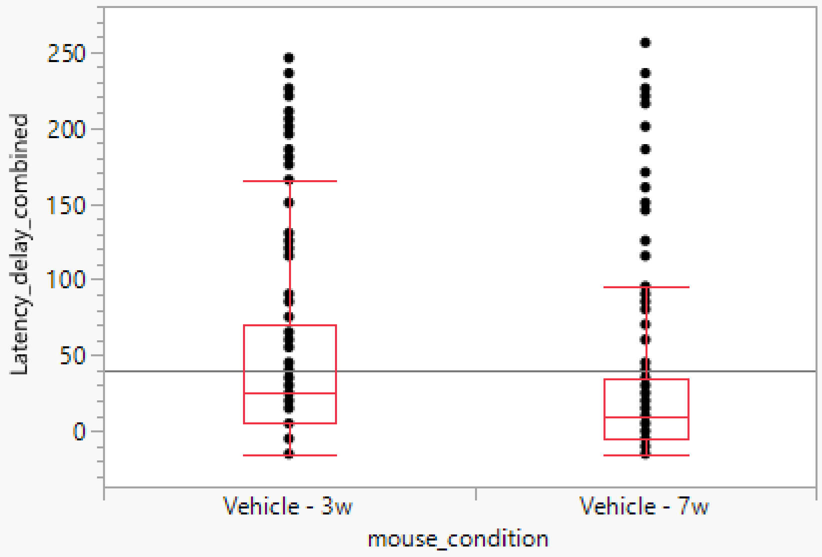

**Quantiles**

| Level | Minimum | 10% | 25% | Median | 75% | 90% | Maximum |
|---|---|---|---|---|---|---|---|
| Vehicle - 3w | -15.0602 | -15.0602 | 5.020157 | 25.10048 | 70.2812 | 167.6708 | 245.984 |
| Vehicle - 7w | -15.0602 | -10.0402 | -5.02 | 10.04016 | 35.14064 | 120.482 | 256.0242 |

**Wilcoxon / Kruskal-Wallis Tests (Rank Sums)**

| Level | Count | Score Sum | Expected Score | Score Mean | (Mean-Mean0)/Std0 |
|---|---|---|---|---|---|
| Vehicle - 3w | 157 | 25437.0 | 23707.0 | 162.019 | 2.299 |
| Vehicle - 7w | 144 | 20014.0 | 21744.0 | 138.986 | -2.299 |

**Wilcoxon Two-Sample Test, Normal Approximation**

| S | Z | Prob>\|Z\| |
|---|---|---|
| 20014 | -2.29910 | 0.0215* |

**Kruskal-Wallis Test, ChiSquare Approximation**

| ChiSquare | DF | Prob>ChiSq |
|---|---|---|
| 5.2889 | 1 | 0.0215* |

Missing Rows 3347

Excluded Rows 5684

#### Oneway Analysis of Latency_delay_combined By mouse_condition

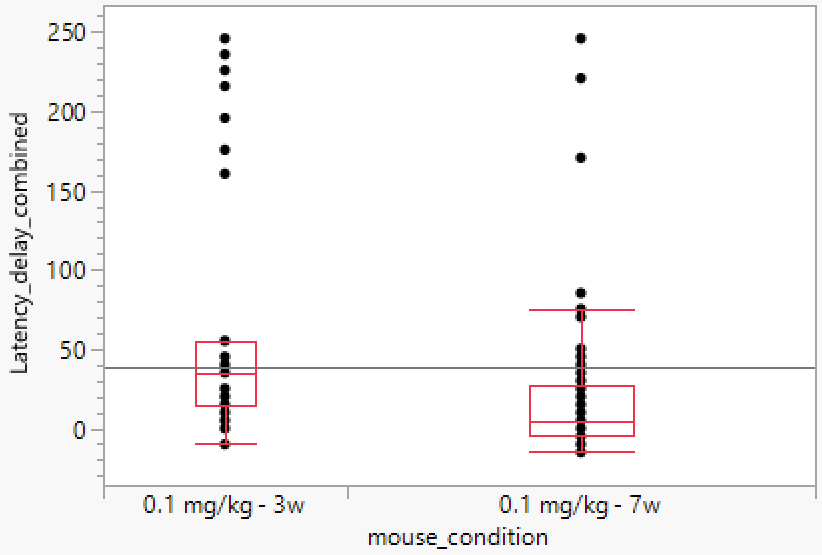

##### Quantiles

| Level | Minimum | 10% | 25% | Median | 75% | 90% | Maximum |
|---|---|---|---|---|---|---|---|
| 0.1 mg/kg - 3w | -10.0402 | 0.502012 | 15.06032 | 35.14064 | 55.22096 | 224.8997 | 245.984 |
| 0.1 mg/kg - 7w | -15.0602 | -15.0602 | -5.02 | 5.020157 | 27.61048 | 77.30931 | 245.984 |

##### Wilcoxon / Kruskal-Wallis Tests (Rank Sums)

| Level | Count | Score Sum | Expected Score | Score Mean | (Mean-Mean0)/Std0 |
|---|---|---|---|---|---|
| 0.1 mg/kg - 3w | 40 | 3064.00 | 2360.00 | 76.6000 | 4.053 |
| 0.1 mg/kg - 7w | 77 | 3839.00 | 4543.00 | 49.8571 | -4.053 |

###### Wilcoxon Two-Sample Test, Normal Approximation

| S | Z | Prob>\|Z\| |
|---|---|---|
| 3064 | 4.05274 | <.0001* |

###### Kruskal-Wallis Test, ChiSquare Approximation

| ChiSquare | DF | Prob>ChiSq |
|---|---|---|
| 16.4480 | 1 | <.0001* |

Missing Rows 1809

Excluded Rows 7406

#### Oneway Analysis of Latency_delay_combined By mouse_condition

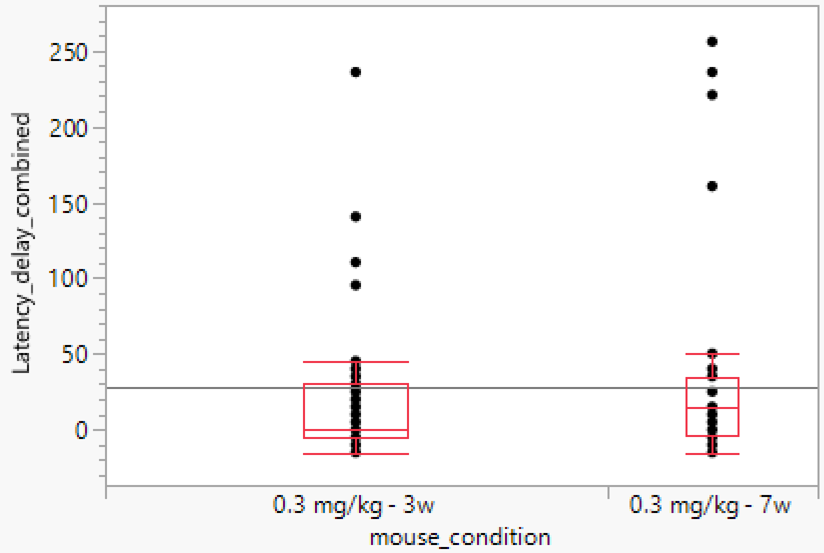

##### Quantiles

| Level | Minimum | 10% | 25% | Median | 75% | 90% | Maximum |
|---|---|---|---|---|---|---|---|
| 0.3 mg/kg - 3w | -15.0602 | -10.0402 | -5.02 | -4.02e-6 | 30.12048 | 95.3816 | 235.9439 |
| 0.3 mg/kg - 7w | -15.0602 | -10.0402 | -3.765 | 15.06032 | 35.14064 | 202.8112 | 256.0242 |

##### Wilcoxon / Kruskal-Wallis Tests (Rank Sums)

| Level | Count | Score Sum | Expected Score | Score Mean | (Mean-Mean0)/Std0 |
|---|---|---|---|---|---|
| 0.3 mg/kg - 3w | 76 | 3978.00 | 4142.00 | 52.3421 | -1.106 |
| 0.3 mg/kg - 7w | 32 | 1908.00 | 1744.00 | 59.6250 | 1.106 |

###### Wilcoxon Two-Sample Test, Normal Approximation

| S | Z | Prob>\|Z\| |
|---|---|---|
| 1908 | 1.10623 | 0.2686 |

###### Kruskal-Wallis Test, ChiSquare Approximation

| ChiSquare | DF | Prob>ChiSq |
|---|---|---|
| 1.2312 | 1 | 0.2672 |

Missing Rows 1570

Excluded Rows 7654

**7B**

#### Oneway Analysis of Latency_combined By mouse_condition

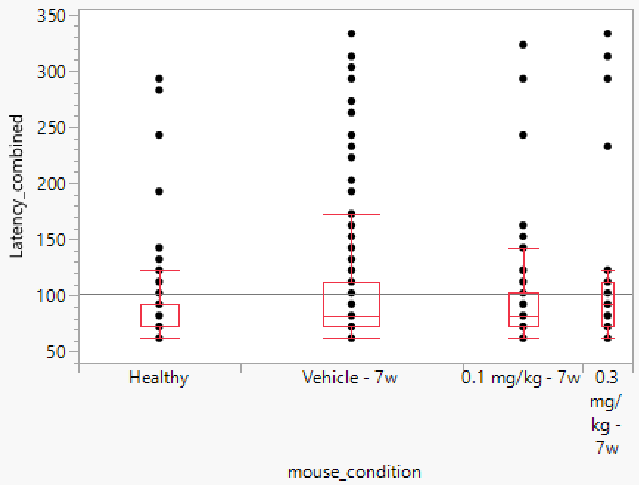

##### Quantiles

| Level | Minimum | 10% | 25% | Median | 75% | 90% | Maximum |
|---|---|---|---|---|---|---|---|
| Healthy | 62.20884 | 62.20884 | 72.249 | 72.249 | 92.32932 | 142.5301 | 293.1325 |
| Vehicle - 7w | 62.20884 | 62.20884 | 72.249 | 82.28916 | 112.4096 | 197.751 | 333.2932 |
| 0.1 mg/kg - 7w | 62.20884 | 62.20884 | 72.249 | 82.28916 | 102.3695 | 154.5783 | 323.253 |
| 0.3 mg/kg - 7w | 62.20884 | 62.20884 | 72.249 | 92.32932 | 112.4096 | 275.0602 | 333.2932 |

##### Wilcoxon / Kruskal-Wallis Tests (Rank Sums)

| Level | Count | Score Sum | Expected Score | Score Mean | (Mean-Mean0)/Std0 |
|---|---|---|---|---|---|
| Healthy | 104 | 15950.0 | 18616.0 | 153.365 | -3.045 |
| Vehicle - 7w | 144 | 27869.0 | 25776.0 | 193.535 | 2.214 |
| 0.1 mg/kg - 7w | 77 | 13688.0 | 13783.0 | 177.766 | -0.119 |
| 0.3 mg/kg - 7w | 32 | 6396.00 | 5728.00 | 199.875 | 1.213 |

###### Kruskal-Wallis Test, ChiSquare Approximation

| ChiSquare | DF | Prob>ChiSq |
|---|---|---|
| 10.8514 | 3 | 0.0126* |

##### Nonparametric Comparisons For All Pairs Using Steel-Dwass Method

| q* | Alpha |
|---|---|
| 2.56903 | 0.05 |

| Level | - Level | Score Mean Difference | Std Err Dif | Z | p-Value | Hodges-Lehmann | Lower CL | Upper CL |
|---|---|---|---|---|---|---|---|---|
| Vehicle - 7w | Healthy | 27.5638 | 9.110969 | 3.02535 | 0.0133* | 10.04016 | 0.0000 | 10.04016 |
| 0.3 mg/kg - 7w | Healthy | 16.5300 | 7.813189 | 2.11566 | 0.1480 | 10.04016 | 0.0000 | 30.12048 |
| 0.1 mg/kg - 7w | Healthy | 13.4597 | 7.730710 | 1.74107 | 0.3023 | 0.00000 | 0.0000 | 10.04016 |
| 0.3 mg/kg - 7w | 0.1 mg/kg - 7w | 7.5645 | 6.559891 | 1.15315 | 0.6565 | 10.04016 | -10.0402 | 20.08032 |
| 0.3 mg/kg - 7w | Vehicle - 7w | 3.4757 | 9.883303 | 0.35167 | 0.9851 | 0.00000 | -10.0402 | 20.08032 |
| 0.1 mg/kg - 7w | Vehicle - 7w | -10.3444 | 8.939696 | -1.15713 | 0.6540 | 0.00000 | -10.0402 | 0.00000 |

Difference Plot

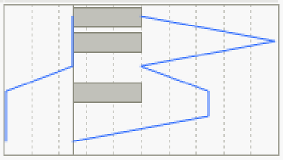

Missing Rows 3821

Excluded Rows 5154

**7C**

#### Oneway Analysis of n70_latency (ms) By condition

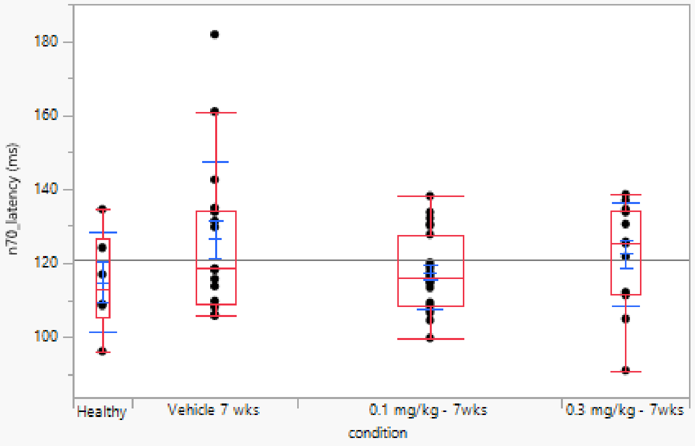

##### Quantiles

| Level | Minimum | 10% | 25% | Median | 75% | 90% | Maximum |
|---|---|---|---|---|---|---|---|
| Healthy | 96.11845 | 96.11845 | 105.4222 | 112.9252 | 126.7307 | 134.5338 | 134.5338 |
| Vehicle 7 wks | 105.7223 | 106.0424 | 108.9236 | 118.5274 | 134.3337 | 165.106 | 181.7527 |
| 0.1 mg/kg - 7wks | 99.71989 | 104.5218 | 108.5234 | 116.1265 | 127.7311 | 132.453 | 138.1353 |
| 0.3 mg/kg - 7wks | 90.91637 | 96.51861 | 111.7247 | 125.3301 | 134.1337 | 137.8952 | 138.5354 |

##### Means and Std Deviations

| Level | Number | Mean | Std Dev | Std Err Mean | Lower 95% | Upper 95% | Std Dev Lower 95% | Std Dev Upper 95% |
|---|---|---|---|---|---|---|---|---|
| Healthy | 6 | 114.85928 | 13.445418 | 5.4890688 | 100.74918 | 128.96938 | 8.3927374 | 32.97642 |
| Vehicle 7 wks | 17 | 126.48353 | 20.757222 | 5.0343658 | 115.81116 | 137.15591 | 15.459353 | 31.591017 |
| 0.1 mg/kg - 7wks | 27 | 117.50478 | 10.129622 | 1.9494467 | 113.49763 | 121.51192 | 7.977244 | 13.881955 |
| 0.3 mg/kg - 7wks | 13 | 122.43667 | 14.044757 | 3.8953148 | 113.94951 | 130.92383 | 10.071299 | 23.184174 |

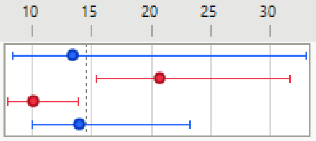

##### Wilcoxon / Kruskal-Wallis Tests (Rank Sums)

| Level | Count | Score Sum | Expected Score | Score Mean | (Mean-Mean0)/Std0 |
|---|---|---|---|---|---|
| Healthy | 6 | 156.000 | 192.000 | 26.0000 | -0.831 |
| Vehicle 7 wks | 17 | 609.500 | 544.000 | 35.8529 | 1.007 |
| 0.1 mg/kg - 7wks | 27 | 770.000 | 864.000 | 28.5185 | -1.299 |
| 0.3 mg/kg - 7wks | 13 | 480.500 | 416.000 | 36.9615 | 1.087 |

###### Kruskal-Wallis Test, ChiSquare Approximation

| ChiSquare | DF | Prob>ChiSq |
|---|---|---|
| 3.3222 | 3 | 0.3446 |

Excluded Rows 81

**7D**

#### Oneway Analysis of correlation_baseline By condition

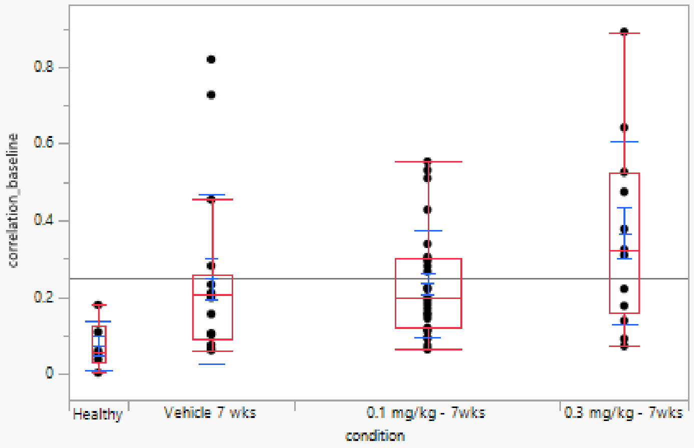

##### Quantiles

| Level | Minimum | 10% | 25% | Median | 75% | 90% | Maximum |
|---|---|---|---|---|---|---|---|
| Healthy | 0.004105 | 0.004105 | 0.029768 | 0.056664 | 0.128007 | 0.180706 | 0.180706 |
| Vehicle 7 wks | 0.062297 | 0.066081 | 0.089697 | 0.205705 | 0.258374 | 0.745635 | 0.819639 |
| 0.1 mg/kg - 7wks | 0.064952 | 0.089624 | 0.121091 | 0.200588 | 0.301223 | 0.514441 | 0.553502 |
| 0.3 mg/kg - 7wks | 0.073988 | 0.081462 | 0.158984 | 0.324872 | 0.527018 | 0.791625 | 0.891375 |

##### Means and Std Deviations

| Level | Number | Mean | Std Dev | Std Err Mean | Lower 95% | Upper 95% | Std Dev Lower 95% | Std Dev Upper 95% |
|---|---|---|---|---|---|---|---|---|
| Healthy | 6 | 0.0744834 | 0.0624481 | 0.0254943 | 0.0089481 | 0.1400186 | 0.0389806 | 0.153161 |
| Vehicle 7 wks | 17 | 0.2495023 | 0.221468 | 0.0537139 | 0.1356339 | 0.3633706 | 0.1649427 | 0.3370586 |
| 0.1 mg/kg - 7wks | 27 | 0.2358758 | 0.1381764 | 0.0265921 | 0.1812151 | 0.2905366 | 0.1088162 | 0.1893613 |
| 0.3 mg/kg - 7wks | 13 | 0.3678726 | 0.2394254 | 0.0664047 | 0.2231893 | 0.5125559 | 0.1716886 | 0.395228 |

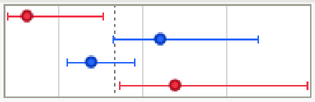

##### Wilcoxon / Kruskal-Wallis Tests (Rank Sums)

| Level | Count | Score Sum | Expected Score | Score Mean | (Mean-Mean0)/Std0 |
|---|---|---|---|---|---|
| Healthy | 6 | 55.000 | 192.000 | 9.1667 | -3.196 |
| Vehicle 7 wks | 17 | 524.000 | 544.000 | 30.8235 | -0.302 |
| 0.1 mg/kg - 7wks | 27 | 893.000 | 864.000 | 33.0741 | 0.396 |
| 0.3 mg/kg - 7wks | 13 | 544.000 | 416.000 | 41.8462 | 2.165 |

###### Kruskal-Wallis Test, ChiSquare Approximation

| ChiSquare | DF | Prob>ChiSq |
|---|---|---|
| 13.2237 | 3 | 0.0042* |

##### Nonparametric Comparisons For All Pairs Using Steel-Dwass Method

| q* | Alpha |
|---|---|
| 2.56903 | 0.05 |

| Level | - Level | Score Mean Difference | Std Err Dif | Z | p-Value | Hodges-Lehmann | Lower CL | Upper CL |
|---|---|---|---|---|---|---|---|---|
| 0.1 mg/kg - 7wks | Healthy | 13.34259 | 4.364206 | 3.057278 | 0.0120* | 0.1326487 | 0.021173 | 0.3009645 |
| Vehicle 7 wks | Healthy | 8.45588 | 3.220644 | 2.625525 | 0.0430* | 0.1205890 | -0.003575 | 0.4502969 |
| 0.3 mg/kg - 7wks | Healthy | 7.91667 | 2.777350 | 2.850439 | 0.0227* | 0.2696844 | 0.029499 | 0.5889915 |
| 0.3 mg/kg - 7wks | 0.1 mg/kg - 7wks | 6.38177 | 3.946460 | 1.617086 | 0.3689 | 0.1038481 | -0.052990 | 0.3199678 |
| 0.3 mg/kg - 7wks | Vehicle 7 wks | 5.15837 | 3.243511 | 1.590367 | 0.3841 | 0.1110033 | -0.093341 | 0.3681243 |
| 0.1 mg/kg - 7wks | Vehicle 7 wks | 1.82135 | 3.977058 | 0.457964 | 0.9681 | 0.0182528 | -0.099792 | 0.1194281 |

Difference Plot

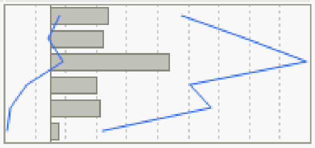

Excluded Rows 81

**7E**

#### Response Restoration of myelin (% from baseline)

##### Regression Plot

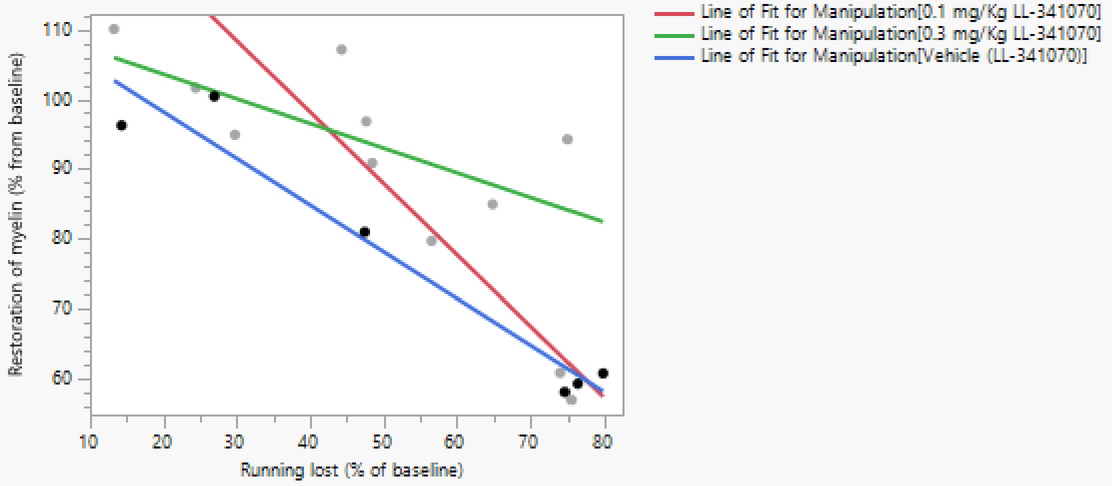

##### Effect Summary

| Source | Logworth | PValue |
|---|---|---|
| Running lost (% of baseline) | 4.585 | 0.00003 |
| Manipulation | 1.428 | 0.03731 |
| Manipulation*Running lost (% of baseline) | 1.119 | 0.07603 |

Remove Add Edit FDR

##### Summary of Fit

| | |
|---|---|
| RSquare | 0.856293 |
| RSquare Adj | 0.790971 |
| Root Mean Square Error | 8.492712 |
| Mean of Response | 84.32663 |
| Observations (or Sum Wgts) | 17 |

##### Analysis of Variance

| Source | DF | Sum of Squares | Mean Square | F Ratio |
|---|---|---|---|---|
| Model | 5 | 4727.4740 | 945.495 | 13.1089 |
| Error | 11 | 793.3878 | 72.126 | **Prob > F** |
| C. Total | 16 | 5520.8618 | | 0.0003* |

##### Parameter Estimates

| Term | Estimate | Std Error | t Ratio | Prob>\|t\| |
|---|---|---|---|---|
| Intercept | 120.47351 | 5.500771 | 21.90 | <.0001* |
| Manipulation[0.1 mg/Kg LL-341070] | 1.1185775 | 2.901621 | 0.39 | 0.7072 |
| Manipulation[0.3 mg/Kg LL-341070] | 7.0974333 | 3.056503 | 2.32 | 0.0404* |
| Running lost (% of baseline) | -0.682792 | 0.099004 | -6.90 | <.0001* |
| Manipulation[0.1 mg/Kg LL-341070]*(Running lost (% of baseline)-51.3584) | -0.341628 | 0.157018 | -2.18 | 0.0523 |
| Manipulation[0.3 mg/Kg LL-341070]*(Running lost (% of baseline)-51.3584) | 0.3284971 | 0.135128 | 2.43 | 0.0334* |

##### Effect Tests

| Source | Nparm | DF | Sum of Squares | F Ratio | Prob > F |
|---|---|---|---|---|---|
| Manipulation | 2 | 2 | 649.2337 | 4.5007 | 0.0373* |
| Running lost (% of baseline) | 1 | 1 | 3430.5300 | 47.5629 | <.0001* |
| Manipulation*Running lost (% of baseline) | 2 | 2 | 474.0832 | 3.2865 | 0.0760 |

#### Effect Details

##### Manipulation

###### Least Squares Means Table

| Level | Least Sq Mean | Std Error | Mean |
|---|---|---|---|
| 0.1 mg/Kg LL-341070 | 86.524989 | 3.4905622 | 84.5657 |
| 0.3 mg/Kg LL-341070 | 92.503845 | 3.8668404 | 94.1189 |
| Vehicle (LL-341070) | 77.190401 | 3.4765026 | 75.9274 |

###### Least Squares Means Plot

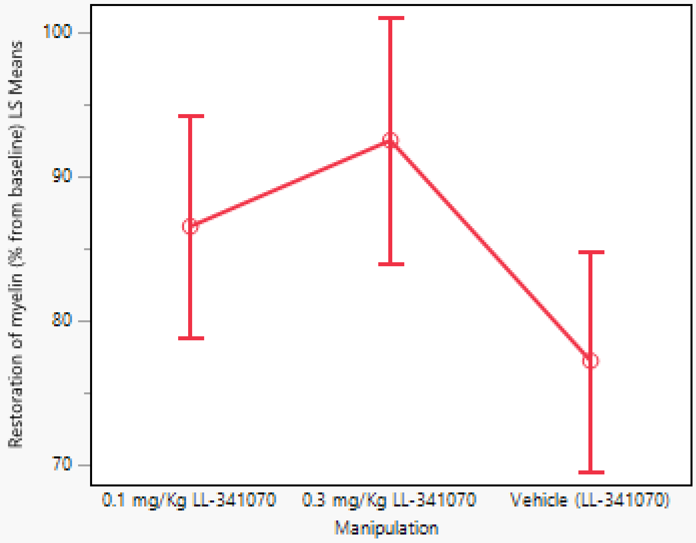

##### Running lost (% of baseline)

##### Manipulation*Running lost (% of baseline)

**7F**

#### Response Restoration of myelin (% from baseline)

##### Regression Plot

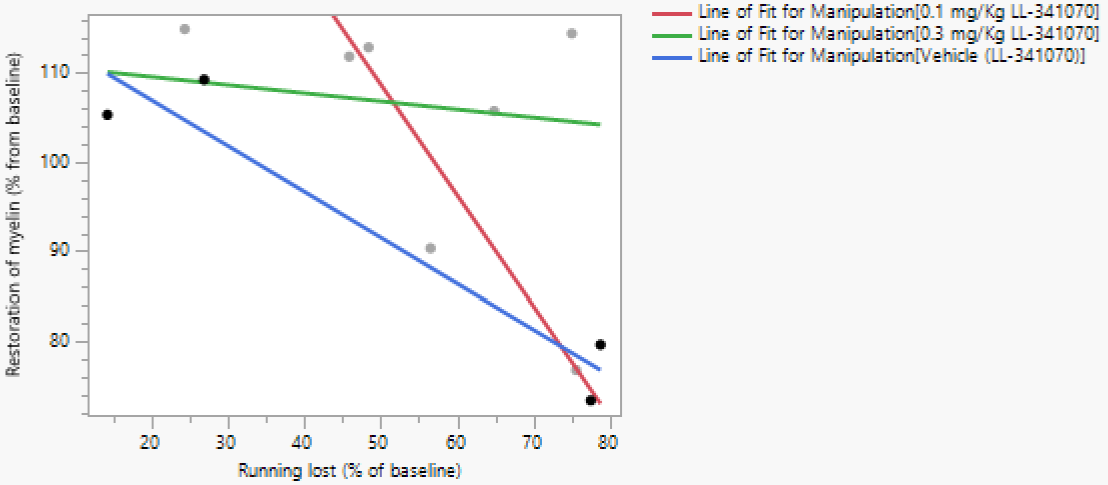

##### Effect Summary

| Source | Logworth | PValue |
|---|---|---|
| Running lost (% of baseline) | 1.794 | 0.01606 |
| Manipulation | 0.924 | 0.11913 |
| Manipulation*Running lost (% of baseline) | 0.823 | 0.15034 |

Remove Add Edit FDR

##### Summary of Fit

| | |
|---|---|
| RSquare | 0.820975 |
| RSquare Adj | 0.641951 |
| Root Mean Square Error | 9.720737 |
| Mean of Response | 99.49512 |
| Observations (or Sum Wgts) | 11 |

##### Analysis of Variance

| Source | DF | Sum of Squares | Mean Square | F Ratio |
|---|---|---|---|---|
| Model | 5 | 2166.6370 | 433.327 | 4.5858 |
| Error | 5 | 472.4637 | 94.493 | Prob > F |
| C. Total | 10 | 2639.1006 | | 0.0600 |

##### Parameter Estimates

| Term | Estimate | Std Error | t Ratio | Prob>\|t\| |
|---|---|---|---|---|
| Intercept | 133.19503 | 10.04181 | 13.26 | <.0001* |
| Manipulation[0.1 mg/Kg LL-341070] | 4.1983132 | 4.483545 | 0.94 | 0.3921 |
| Manipulation[0.3 mg/Kg LL-341070] | 6.2693702 | 4.11804 | 1.52 | 0.1884 |
| Running lost (% of baseline) | -0.616444 | 0.172718 | -3.57 | 0.0161* |
| Manipulation[0.1 mg/Kg LL-341070]*(Running lost (% of baseline)-53.4581) | -0.627079 | 0.296798 | -2.11 | 0.0883 |
| Manipulation[0.3 mg/Kg LL-341070]*(Running lost (% of baseline)-53.4581) | 0.5249235 | 0.227525 | 2.31 | 0.0692 |

##### Effect Tests

| Source | Nparm | DF | Sum of Squares | F Ratio | Prob > F |
|---|---|---|---|---|---|
| Manipulation | 2 | 2 | 634.0628 | 3.3551 | 0.1191 |
| Running lost (% of baseline) | 1 | 1 | 1203.6835 | 12.7384 | 0.0161* |
| Manipulation*Running lost (% of baseline) | 2 | 2 | 535.7096 | 2.8347 | 0.1503 |

Effect Details

Manipulation

Least Squares Means Table

| Level | Least Sq Mean | Std Error | Mean |
|---|---|---|---|
| 0.1 mg/Kg LL-341070 | 104.43940 | 5.7664211 | 100.500 |
| 0.3 mg/Kg LL-341070 | 106.51045 | 4.8805578 | 106.352 |
| Vehicle (LL-341070) | 89.77340 | 4.9084897 | 91.884 |

Least Squares Means Plot

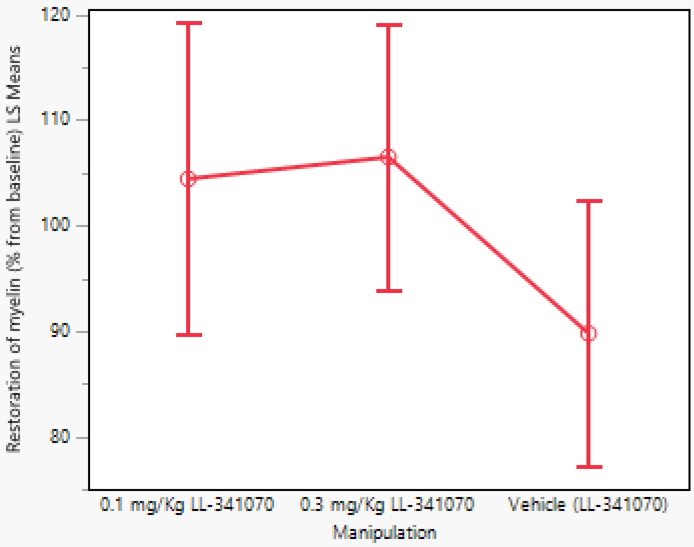

Running lost (% of baseline)

Manipulation*Running lost (% of baseline)

**Supplementary figures**

**Figure S4**

**S4A**

See Fig 3H

**S4B**

See Fig 3H

**S4E**

See Fig 3I

**S4F**

See Fig 3J

**S4G**

See Fig 3J

**S4H**

See Fig 3J

#### Figure S5

### S5B

##### Oneway Analysis of Running lost (% of baseline) By Manipulation

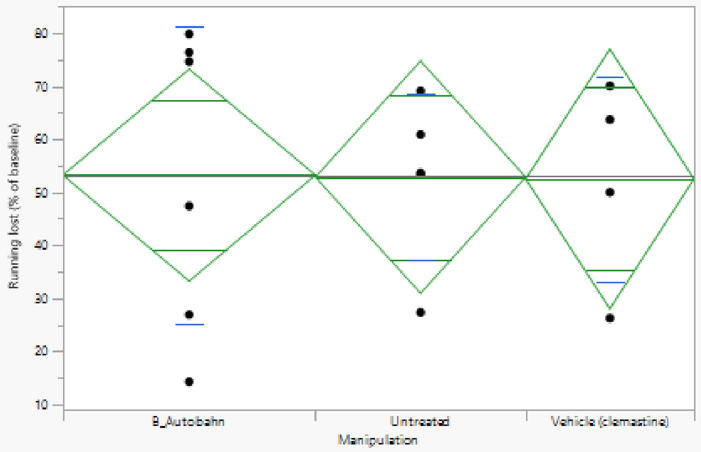

##### Oneway Anova

###### Summary of Fit

| | |
|---|---|
| Rsquare | 0.000221 |
| Adj Rsquare | -0.16641 |
| Root Mean Square Error | 22.45752 |
| Mean of Response | 52.91898 |
| Observations (or Sum Wgts) | 15 |

###### Analysis of Variance

| Source | DF | Sum of Squares | Mean Square | F Ratio | Prob > F |
|---|---|---|---|---|---|
| Manipulation | 2 | 1.3407 | 0.670 | 0.0013 | 0.9987 |
| Error | 12 | 6052.0842 | 504.340 | | |
| C. Total | 14 | 6053.4249 | | | |

###### Means for Oneway Anova

| Level | Number | Mean | Std Error | Lower 95% | Upper 95% |
|---|---|---|---|---|---|
| B_Autobahn | 6 | 53.2444 | 9.168 | 33.269 | 73.220 |
| Untreated | 5 | 52.8602 | 10.043 | 30.978 | 74.743 |
| Vehicle (clemastine) | 4 | 52.5043 | 11.229 | 28.039 | 76.970 |

Std Error uses a pooled estimate of error variance

##### Tests that the Variances are Equal

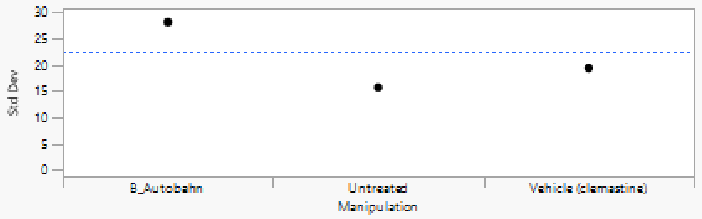

| Level | Count | Std Dev | MeanAbsDif to Mean | MeanAbsDif to Median |
|---|---|---|---|---|
| B_Autobahn | 6 | 28.08714 | 23.70570 | 23.70570 |
| Untreated | 5 | 15.65403 | 10.20069 | 9.87364 |
| Vehicle (clemastine) | 4 | 19.38601 | 14.36868 | 14.36868 |

| Test | F Ratio | DFNum | DFDen | Prob > F |
|---|---|---|---|---|
| O'Brien[.5] | 1.4990 | 2 | 12 | 0.2624 |
| Brown-Forsythe | 1.8209 | 2 | 12 | 0.2039 |
| Levene | 2.3718 | 2 | 12 | 0.1355 |
| Bartlett | 0.6863 | 2 | . | 0.5034 |

Warning: Small sample sizes. Use Caution.

###### Welch's Test

Welch Anova testing Means Equal, allowing Std Devs Not Equal

| F Ratio | DFNum | DFDen | Prob > F |
|---|---|---|---|
| 0.0011 | 2 | 7.4116 | 0.9989 |

Excluded Rows 677

## S5C

##### Response Max gain rate

##### Whole Model

###### Regression Plot

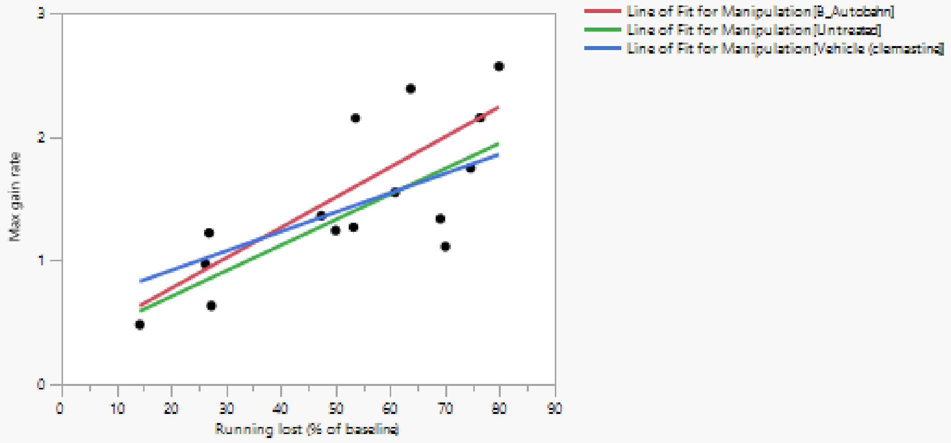

###### Actual by Predicted Plot

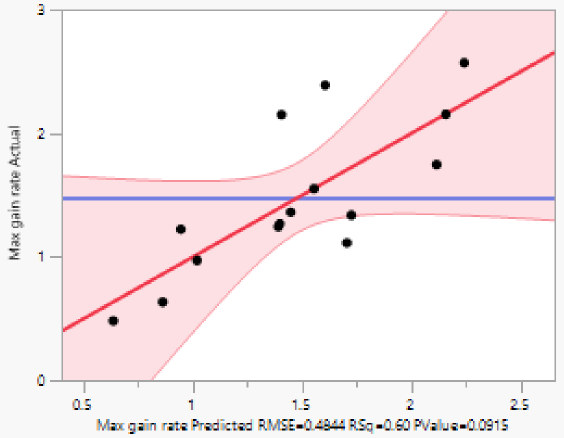

###### Effect Summary

| Source | Logworth | | PValue |
|---|---|---|---|
| Running lost (% of baseline) | 1.614 | | 0.02434 |
| Manipulation | 0.100 | | 0.79490 |
| Running lost (% of baseline)*Manipulation | 0.065 | | 0.86112 |

Remove Add Edit ☐ FDR

###### Residual by Predicted Plot

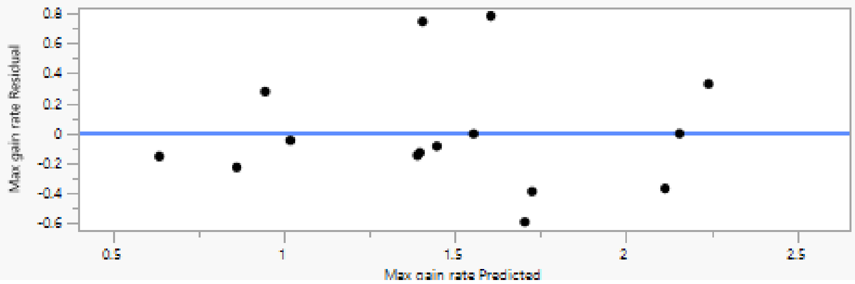

#### Summary of Fit

| | |
|---|---|
| RSquare | 0.601433 |
| RSquare Adj | 0.380007 |
| Root Mean Square Error | 0.484384 |
| Mean of Response | 1.479971 |
| Observations (or Sum Wgts) | 15 |

#### Analysis of Variance

| Source | DF | Sum of Squares | Mean Square | F Ratio |
|---|---|---|---|---|
| Model | 5 | 3.1864562 | 0.637291 | 2.7162 |
| Error | 9 | 2.1116489 | 0.234628 | **Prob > F** |
| C. Total | 14 | 5.2981051 | | 0.0915 |

#### Parameter Estimates

| Term | Estimate | Std Error | t Ratio | Prob>\|t\| |
|---|---|---|---|---|
| Intercept | 0.3962927 | 0.415778 | 0.95 | 0.3654 |
| Running lost (% of baseline) | 0.0202743 | 0.007505 | 2.70 | 0.0243* |
| Manipulation[B_Autobahn] | 0.1126081 | 0.17064 | 0.66 | 0.5258 |
| Manipulation[Untreated] | -0.079256 | 0.17811 | -0.44 | 0.6668 |
| (Running lost (% of baseline)-52.919)*Manipulation[B_Autobahn] | 0.0042231 | 0.008727 | 0.48 | 0.6400 |
| (Running lost (% of baseline)-52.919)*Manipulation[Untreated] | 0.0003911 | 0.011667 | 0.03 | 0.9740 |

#### Effect Tests

| Source | Nparm | DF | Sum of Squares | F Ratio | Prob > F |
|---|---|---|---|---|---|
| Running lost (% of baseline) | 1 | 1 | 1.7121544 | 7.2973 | 0.0243* |
| Manipulation | 2 | 2 | 0.1105047 | 0.2355 | 0.7949 |
| Running lost (% of baseline)*Manipulation | 2 | 2 | 0.0713417 | 0.1520 | 0.8611 |

**S5D**

#### Response Gain rate 0-11

##### Whole Model

###### Regression Plot

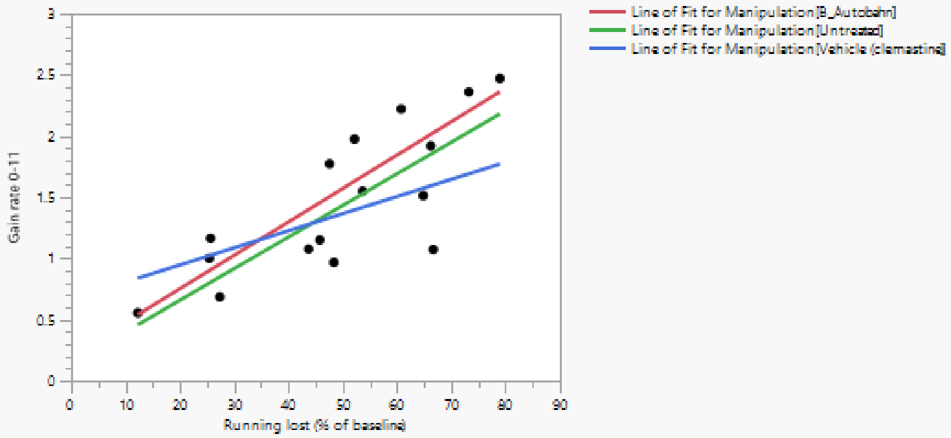

###### Actual by Predicted Plot

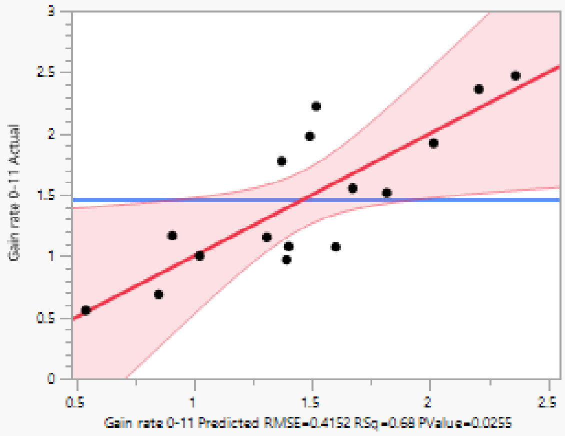

###### Effect Summary

| Source | Logworth | | PValue | |
|---|---|---|---|---|
| Running lost (% of baseline) | 1.986 | | 0.01032 | |
| Running lost (% of baseline)*Manipulation | 0.172 | | 0.67224 | |
| Manipulation | 0.140 | | 0.72401 | ^ |

Remove Add Edit ☐ FDR ('^' denotes effects with containing effects above them)

###### Residual by Predicted Plot

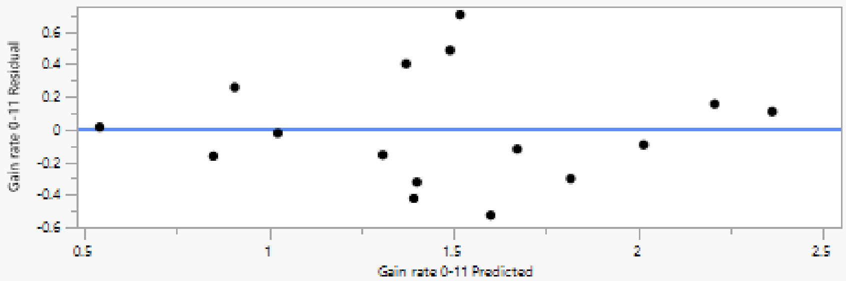

#### Summary of Fit

| | |
|---|---|
| RSquare | 0.677763 |
| RSquare Adj | 0.516645 |
| Root Mean Square Error | 0.415225 |
| Mean of Response | 1.467299 |
| Observations (or Sum Wgts) | 16 |

#### Analysis of Variance

| Source | DF | Sum of Squares | Mean Square | F Ratio |
|---|---|---|---|---|
| Model | 5 | 3.6263555 | 0.725271 | 4.2066 |
| Error | 10 | 1.7241212 | 0.172412 | Prob > F |
| C. Total | 15 | 5.3504767 | | 0.0255* |

#### Parameter Estimates

| Term | Estimate | Std Error | t Ratio | Prob>\|t\| |
|---|---|---|---|---|
| Intercept | 0.3396067 | 0.363282 | 0.93 | 0.3719 |
| Running lost (% of baseline) | 0.0223693 | 0.0071 | 3.15 | 0.0103* |
| Manipulation[B_Autobahn] | 0.1121407 | 0.140162 | 0.80 | 0.4423 |
| Manipulation[Untreated] | -0.025417 | 0.151944 | -0.17 | 0.8705 |
| (Running lost (% of baseline)-49.531)*Manipulation[B_Autobahn] | 0.0049323 | 0.008118 | 0.61 | 0.5570 |
| (Running lost (% of baseline)-49.531)*Manipulation[Untreated] | 0.0034546 | 0.011381 | 0.30 | 0.7677 |

#### Effect Tests

| Source | Nparm | DF | Sum of Squares | F Ratio | Prob > F |
|---|---|---|---|---|---|
| Running lost (% of baseline) | 1 | 1 | 1.7112752 | 9.9255 | 0.0103* |
| Manipulation | 2 | 2 | 0.1150359 | 0.3336 | 0.7240 |
| Running lost (% of baseline)*Manipulation | 2 | 2 | 0.1425288 | 0.4133 | 0.6722 |

**S5E**

#### Response Gain rate 11-18

##### Whole Model

###### Regression Plot

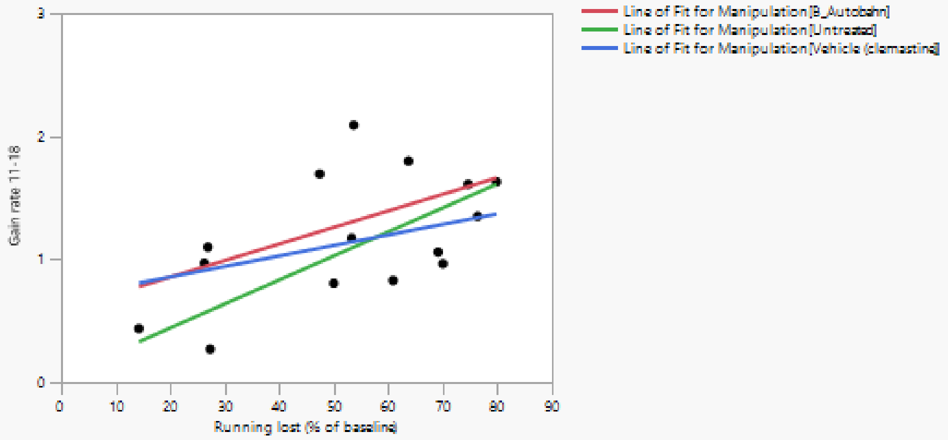

###### Actual by Predicted Plot

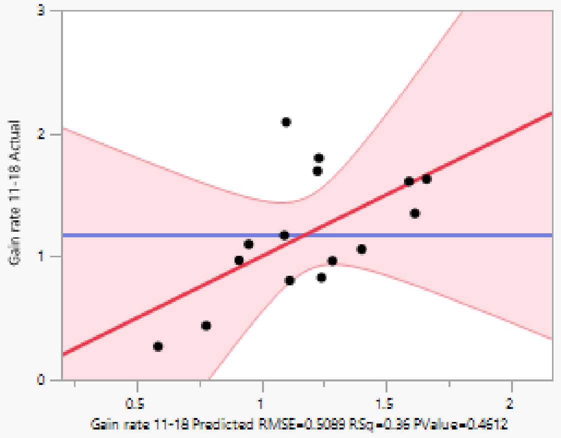

###### Effect Summary

| Source | Logworth | PValue |
| --- | --- | --- |
| Running lost (% of baseline) | 0.947 | 0.11290 |
| Manipulation | 0.111 | 0.77376 |
| Running lost (% of baseline)*Manipulation | 0.052 | 0.88649 |

Remove Add Edit ☐ FDR

###### Residual by Predicted Plot

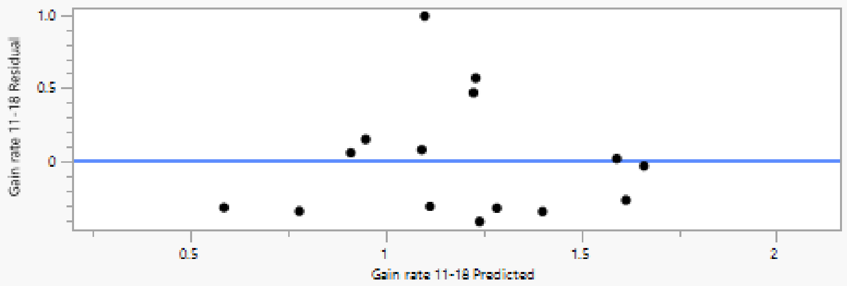

#### Summary of Fit

| | |
|---|---|
| RSquare | 0.361101 |
| RSquare Adj | 0.006157 |
| Root Mean Square Error | 0.508893 |
| Mean of Response | 1.184806 |
| Observations (or Sum Wgts) | 15 |

#### Analysis of Variance

| Source | DF | Sum of Squares | Mean Square | F Ratio |
|---|---|---|---|---|
| Model | 5 | 1.3173220 | 0.263464 | 1.0173 |
| Error | 9 | 2.3307492 | 0.258972 | **Prob > F** |
| C. Total | 14 | 3.6480712 | | 0.4612 |

#### Parameter Estimates

| Term | Estimate | Std Error | t Ratio | Prob>\|t\| |
|---|---|---|---|---|
| Intercept | 0.4407709 | 0.436816 | 1.01 | 0.3393 |
| Running lost (% of baseline) | 0.0138492 | 0.007885 | 1.76 | 0.1129 |
| Manipulation[B_Autobahn] | 0.1249444 | 0.179274 | 0.70 | 0.5034 |
| Manipulation[Untreated] | -0.089017 | 0.187122 | -0.48 | 0.6456 |
| (Running lost (% of baseline)-52.919)*Manipulation[B_Autobahn] | -0.000378 | 0.009168 | -0.04 | 0.9680 |
| (Running lost (% of baseline)-52.919)*Manipulation[Untreated] | 0.0056786 | 0.012257 | 0.46 | 0.6542 |

#### Effect Tests

| Source | Nparm | DF | Sum of Squares | F Ratio | Prob > F |
|---|---|---|---|---|---|
| Running lost (% of baseline) | 1 | 1 | 0.79891279 | 3.0849 | 0.1129 |
| Manipulation | 2 | 2 | 0.13670588 | 0.2639 | 0.7738 |
| Running lost (% of baseline)*Manipulation | 2 | 2 | 0.06324822 | 0.1221 | 0.8865 |

**S5F**

#### Response Gain rate 18-35

##### Whole Model

###### Regression Plot

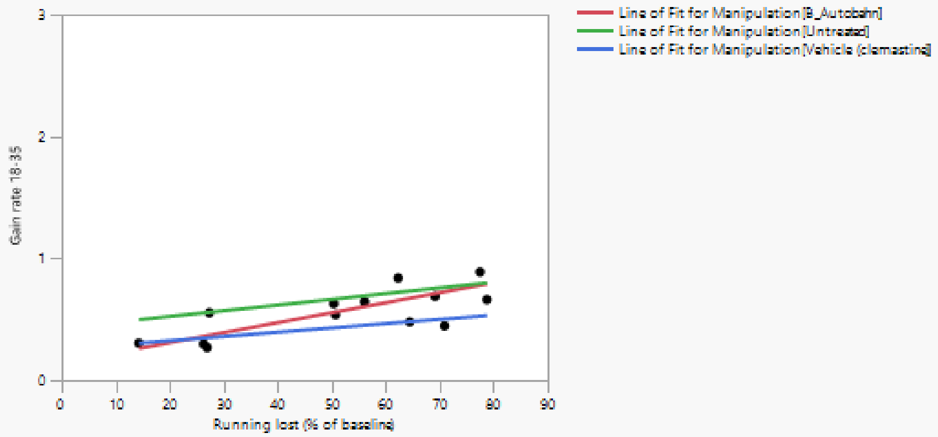

###### Actual by Predicted Plot

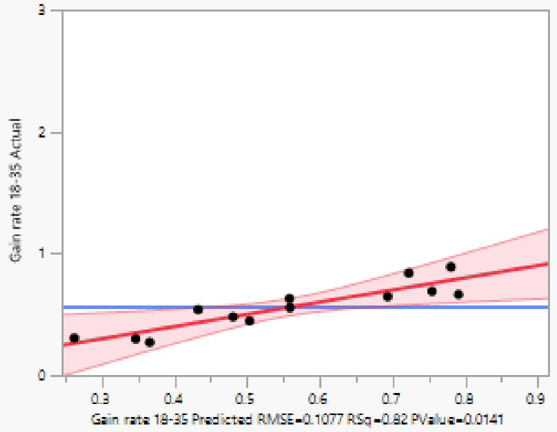

###### Effect Summary

| Source | Logworth | | PValue |
|---|---|---|---|
| Running lost (% of baseline) | 1.877 | | 0.01328 |
| Manipulation | 1.319 | | 0.04798 |
| Running lost (% of baseline)*Manipulation | 0.387 | | 0.40976 |

Remove Add Edit FDR

###### Residual by Predicted Plot

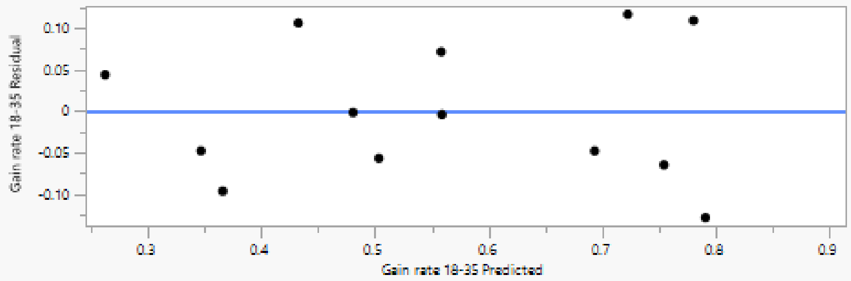

#### Summary of Fit

| | |
|---|---|
| RSquare | 0.824382 |
| RSquare Adj | 0.69894 |
| Root Mean Square Error | 0.107729 |
| Mean of Response | 0.557807 |
| Observations (or Sum Wgts) | 13 |

#### Analysis of Variance

| Source | DF | Sum of Squares | Mean Square | F Ratio |
|---|---|---|---|---|
| Model | 5 | 0.38134909 | 0.076270 | 6.5718 |
| Error | 7 | 0.08123899 | 0.011606 | Prob > F |
| C. Total | 12 | 0.46258808 | | 0.0141* |

#### Parameter Estimates

| Term | Estimate | Std Error | t Ratio | Prob>\|t\| |
|---|---|---|---|---|
| Intercept | 0.2769316 | 0.092836 | 2.98 | 0.0204* |
| Running lost (% of baseline) | 0.0054637 | 0.00166 | 3.29 | 0.0133* |
| Manipulation[B_Autobahn] | 0.0104382 | 0.041116 | 0.25 | 0.8069 |
| Manipulation[Untreated] | 0.1133135 | 0.043474 | 2.61 | 0.0351* |
| (Running lost (% of baseline)-51.9231)*Manipulation[B_Autobahn] | 0.0027376 | 0.001974 | 1.39 | 0.2081 |
| (Running lost (% of baseline)-51.9231)*Manipulation[Untreated] | -0.000785 | 0.002565 | -0.31 | 0.7684 |

#### Effect Tests

| Source | Nparm | DF | Sum of Squares | F Ratio | Prob > F |
|---|---|---|---|---|---|
| Running lost (% of baseline) | 1 | 1 | 0.12569524 | 10.8306 | 0.0133* |
| Manipulation | 2 | 2 | 0.11222498 | 4.8350 | 0.0480* |
| Running lost (% of baseline)*Manipulation | 2 | 2 | 0.02358734 | 1.0162 | 0.4098 |

### Manipulation

#### Leverage Plot

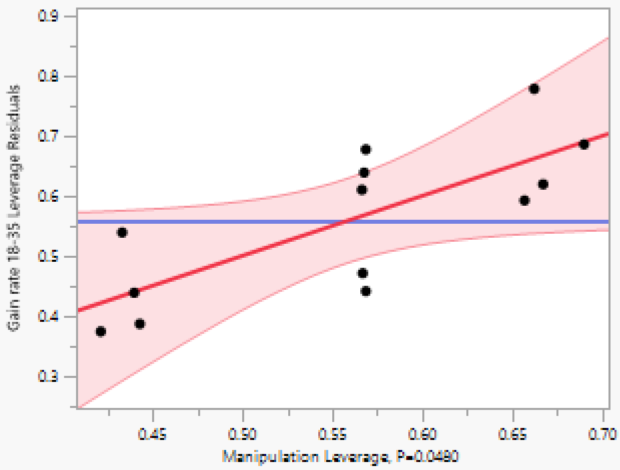

#### Least Squares Means Table

| Level | Least Sq Mean | Std Error | Mean |
|---|---|---|---|
| B_Autobahn | 0.57106300 | 0.04837662 | 0.551656 |
| Untreated | 0.67393826 | 0.05421006 | 0.682381 |
| Vehicle (clemastine) | 0.43687312 | 0.05398686 | 0.440923 |

#### LSMeans Differences Tukey HSD

α= 0.050 Q= 2.94498

LSMean[j]

| LSMean[i] | Mean[i]-Mean[j] / Std Err Dif / Lower CL Dif / Upper CL Dif | B_Autobahn | Untreated | Vehicle (clemastine) |
|---|---|---|---|---|
| B_Autobahn | Mean[i]-Mean[j] | 0 | -0.10288 | 0.13419 |
| | Std Err Dif | 0 | 0.072657 | 0.072491 |
| | Lower CL Dif | 0 | -0.31685 | -0.07929 |
| | Upper CL Dif | 0 | 0.111098 | 0.347673 |
| Untreated | Mean[i]-Mean[j] | 0.102875 | 0 | 0.237065 |
| | Std Err Dif | 0.072657 | 0 | 0.076507 |
| | Lower CL Dif | -0.1111 | 0 | 0.011754 |
| | Upper CL Dif | 0.316848 | 0 | 0.462376 |
| Vehicle (clemastine) | Mean[i]-Mean[j] | -0.13419 | -0.23707 | 0 |
| | Std Err Dif | 0.072491 | 0.076507 | 0 |
| | Lower CL Dif | -0.34767 | -0.46238 | 0 |
| | Upper CL Dif | 0.079293 | -0.01175 | 0 |

| Level | | | Least Sq Mean |
|---|---|---|---|
| Untreated | A | | 0.67393826 |
| B_Autobahn | A | B | 0.57106300 |
| Vehicle (clemastine) | | B | 0.43687312 |

Levels not connected by same letter are significantly different.

| Level | - Level | Difference | Std Err Dif | Lower CL | Upper CL | p-Value |
|---|---|---|---|---|---|---|
| Untreated | Vehicle (clemastine) | 0.2370651 | 0.0765069 | 0.011754 | 0.4623763 | 0.0406* |
| B_Autobahn | Vehicle (clemastine) | 0.1341899 | 0.0724905 | -0.079293 | 0.3476729 | 0.2223 |
| Untreated | B_Autobahn | 0.1028753 | 0.0726569 | -0.111098 | 0.3168482 | 0.3840 |

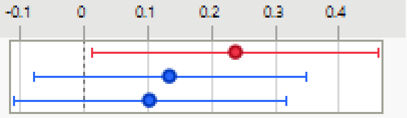

**S5G**

Response Running gain (% of baseline)

Whole Model

Regression Plot

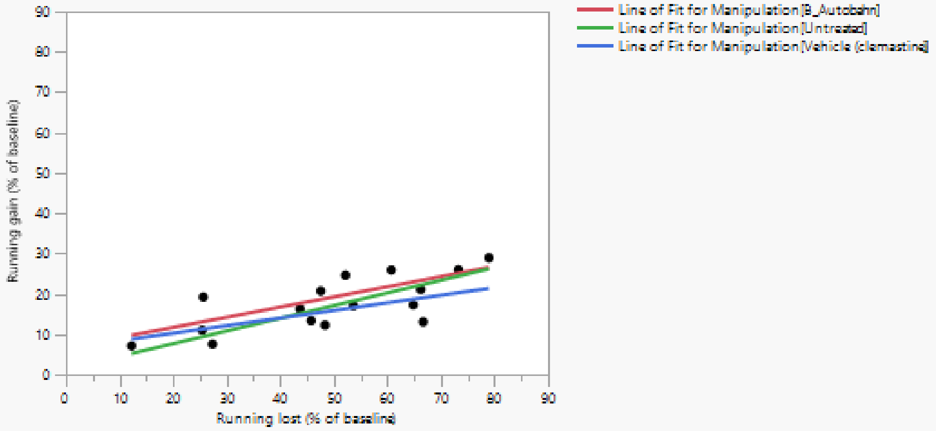

Actual by Predicted Plot

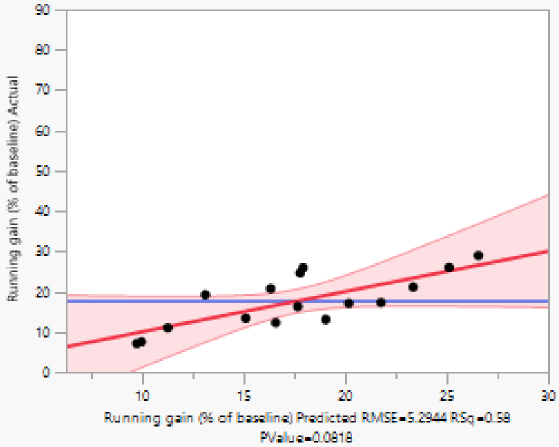

Effect Summary

| Source | Logworth | PValue |
|---|---|---|
| Running lost (% of baseline) | 1.712 | 0.01943 |
| Manipulation | 0.230 | 0.58857 |
| Running lost (% of baseline)*Manipulation | 0.053 | 0.88597 |

Remove Add Edit FDR

Residual by Predicted Plot

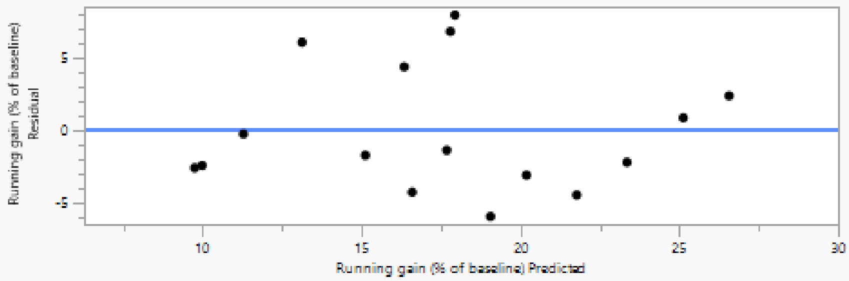

#### Summary of Fit

| | |
|---|---|
| RSquare | 0.578498 |
| RSquare Adj | 0.367746 |
| Root Mean Square Error | 5.294387 |
| Mean of Response | 17.60627 |
| Observations (or Sum Wgts) | 16 |

#### Analysis of Variance

| Source | DF | Sum of Squares | Mean Square | F Ratio |
|---|---|---|---|---|
| Model | 5 | 384.70947 | 76.9419 | 2.7449 |
| Error | 10 | 280.30531 | 28.0305 | Prob > F |
| C. Total | 15 | 665.01478 | | 0.0818 |

#### Parameter Estimates

| Term | Estimate | Std Error | t Ratio | Prob>\|t\| |
|---|---|---|---|---|
| Intercept | 4.8453555 | 4.632073 | 1.05 | 0.3202 |
| Running lost (% of baseline) | 0.2517447 | 0.090533 | 2.78 | 0.0194* |
| Manipulation[B_Autobahn] | 1.8370562 | 1.787151 | 1.03 | 0.3282 |
| Manipulation[Untreated] | -0.350659 | 1.937379 | -0.18 | 0.8600 |
| (Running lost (% of baseline)-49.531)*Manipulation[B_Autobahn] | 0.0003436 | 0.103513 | 0.00 | 0.9974 |
| (Running lost (% of baseline)-49.531)*Manipulation[Untreated] | 0.0628613 | 0.145109 | 0.43 | 0.6741 |

#### Effect Tests

| Source | Nparm | DF | Sum of Squares | F Ratio | Prob > F |
|---|---|---|---|---|---|
| Running lost (% of baseline) | 1 | 1 | 216.73818 | 7.7322 | 0.0194* |
| Manipulation | 2 | 2 | 31.34756 | 0.5592 | 0.5886 |
| Running lost (% of baseline)*Manipulation | 2 | 2 | 6.87006 | 0.1225 | 0.8860 |

## S5H

##### Response Running gain (% of baseline)

##### Whole Model

###### Regression Plot

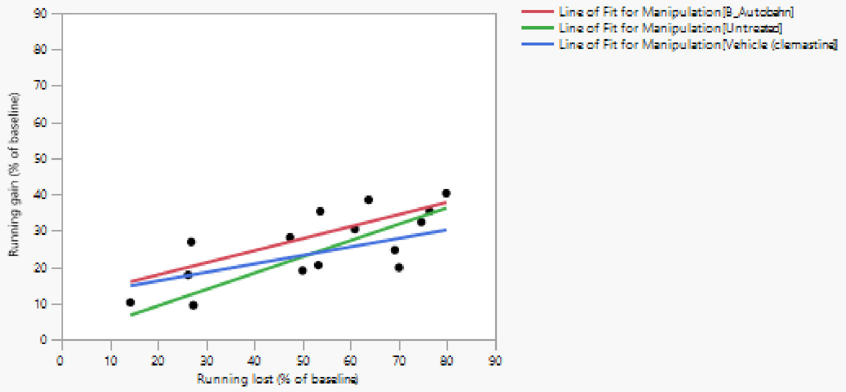

###### Actual by Predicted Plot

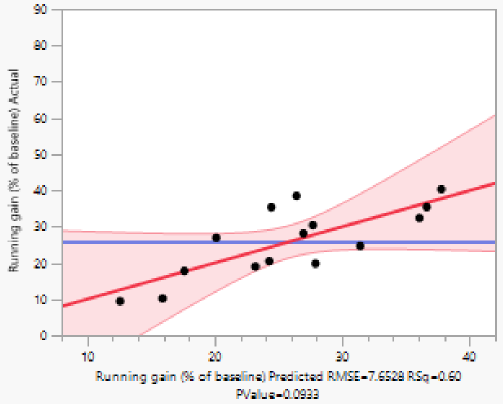

###### Effect Summary

| Source | Logworth | | PValue |
|---|---|---|---|
| Running lost (% of baseline) | 1.729 | | 0.01864 |
| Manipulation | 0.285 | | 0.51913 |
| Running lost (% of baseline)*Manipulation | 0.089 | | 0.81530 |

Remove Add Edit FDR

###### Residual by Predicted Plot

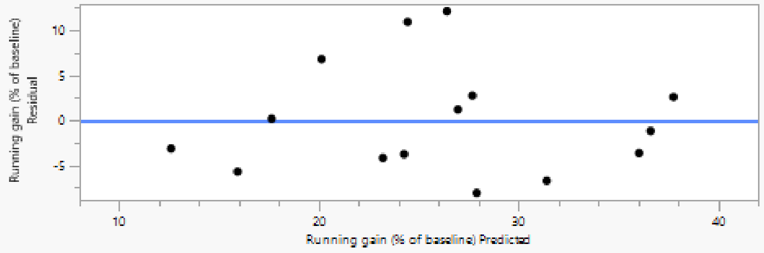

#### Summary of Fit

| | |
|---|---|
| RSquare | 0.59932 |
| RSquare Adj | 0.37672 |
| Root Mean Square Error | 7.652767 |
| Mean of Response | 25.93545 |
| Observations (or Sum Wgts) | 15 |

#### Analysis of Variance

| Source | DF | Sum of Squares | Mean Square | F Ratio |
|---|---|---|---|---|
| Model | 5 | 788.3886 | 157.678 | 2.6924 |
| Error | 9 | 527.0836 | 58.565 | Prob > F |
| C. Total | 14 | 1315.4722 | | 0.0933 |

#### Parameter Estimates

| Term | Estimate | Std Error | t Ratio | Prob>\|t\| |
|---|---|---|---|---|
| Intercept | 7.6250492 | 6.568871 | 1.16 | 0.2756 |
| Running lost (% of baseline) | 0.3396563 | 0.118575 | 2.86 | 0.0186* |
| Manipulation[B_Autobahn] | 3.2011141 | 2.69594 | 1.19 | 0.2655 |
| Manipulation[Untreated] | -1.489274 | 2.813958 | -0.53 | 0.6094 |
| (Running lost (% of baseline)-52.919)*Manipulation[B_Autobahn] | -0.006254 | 0.137874 | -0.05 | 0.9648 |
| (Running lost (% of baseline)-52.919)*Manipulation[Untreated] | 0.1111268 | 0.184326 | 0.60 | 0.5615 |

#### Effect Tests

| Source | Nparm | DF | Sum of Squares | F Ratio | Prob > F |
|---|---|---|---|---|---|
| Running lost (% of baseline) | 1 | 1 | 480.54159 | 8.2053 | 0.0186* |
| Manipulation | 2 | 2 | 82.66650 | 0.7058 | 0.5191 |
| Running lost (% of baseline)*Manipulation | 2 | 2 | 24.46804 | 0.2089 | 0.8153 |

**S5I**

#### Response Running gain (% of baseline)

##### Whole Model

###### Regression Plot

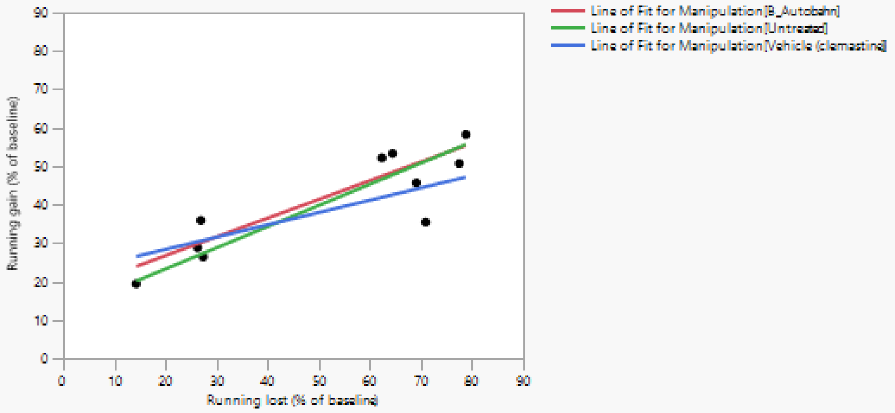

###### Actual by Predicted Plot

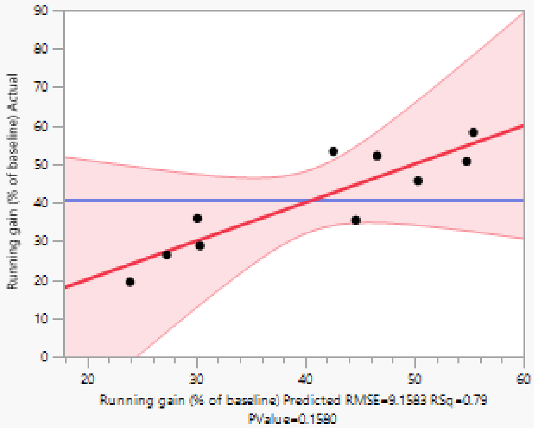

###### Effect Summary

| Source | Logworth | | PValue | |
|---|---|---|---|---|
| Running lost (% of baseline) | 1.483 | | 0.03290 | |
| Running lost (% of baseline)*Manipulation | 0.083 | | 0.82584 | |
| Manipulation | 0.059 | | 0.87318 | ^ |

Remove Add Edit ☐ FDR ('^' denotes effects with containing effects above them)

###### Residual by Predicted Plot

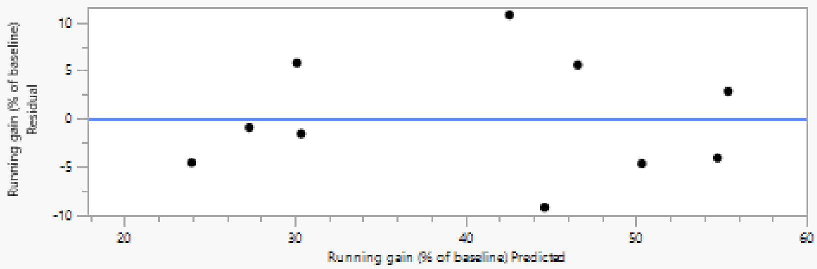

#### Summary of Fit

| | |
|---|---|
| RSquare | 0.786978 |
| RSquare Adj | 0.520702 |
| Root Mean Square Error | 9.158251 |
| Mean of Response | 40.61151 |
| Observations (or Sum Wgts) | 10 |

#### Analysis of Variance

| Source | DF | Sum of Squares | Mean Square | F Ratio |
|---|---|---|---|---|
| Model | 5 | 1239.4370 | 247.887 | 2.9555 |
| Error | 4 | 335.4943 | 83.874 | Prob > F |
| C. Total | 9 | 1574.9313 | | 0.1580 |

#### Parameter Estimates

| Term | Estimate | Std Error | t Ratio | Prob>\|t\| |
|---|---|---|---|---|
| Intercept | 17.063064 | 8.030587 | 2.12 | 0.1008 |
| Running lost (% of baseline) | 0.4530227 | 0.141564 | 3.20 | 0.0329* |
| Manipulation[B_Autobahn] | 1.7278787 | 3.955286 | 0.44 | 0.6848 |
| Manipulation[Untreated] | 0.2643503 | 4.238219 | 0.06 | 0.9533 |
| (Running lost (% of baseline)-51.7829)*Manipulation[B_Autobahn] | 0.0349414 | 0.168213 | 0.21 | 0.8456 |
| (Running lost (% of baseline)-51.7829)*Manipulation[Untreated] | 0.0980835 | 0.218787 | 0.45 | 0.6771 |

#### Effect Tests

| Source | Nparm | DF | Sum of Squares | F Ratio | Prob > F |
|---|---|---|---|---|---|
| Running lost (% of baseline) | 1 | 1 | 858.93465 | 10.2408 | 0.0329* |
| Manipulation | 2 | 2 | 23.53729 | 0.1403 | 0.8732 |
| Running lost (% of baseline)*Manipulation | 2 | 2 | 33.68558 | 0.2008 | 0.8258 |

**Figure S6**

**S6A**

See Figure 5D

**S6B**

See Figure 5D

**S6C**

See Figure 5D

**S6F**

See Figure 5E

**S6G**

See Figure 5F

**S6H**

See Figure 5F

**S6I**

See Figure 5F

**Figure S8**

**S8A**

**Oneway Analysis of dark_latency (ms) By mouse_condition**

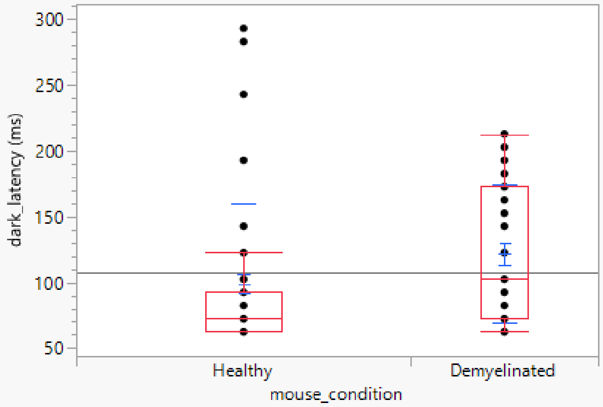

**Quantiles**

| Level | Minimum | 10% | 25% | Median | 75% | 90% | Maximum |
|---|---|---|---|---|---|---|---|
| Healthy | 62.20884 | 62.20884 | 62.20884 | 72.249 | 92.32932 | 197.751 | 293.1325 |
| Demyelinated | 62.20884 | 62.20884 | 72.249 | 102.3695 | 172.6506 | 202.7711 | 212.8112 |

**Means and Std Deviations**

| Level | Number | Mean | Std Dev | Std Err Mean | Lower 95% | Upper 95% | Std Dev Lower 95% | Std Dev Upper 95% |
|---|---|---|---|---|---|---|---|---|
| Healthy | 68 | 98.825892 | 61.32427 | 7.4366601 | 83.982256 | 113.66953 | 52.47027 | 73.800978 |
| Demyelinated | 38 | 121.65715 | 52.476456 | 8.5128053 | 104.40857 | 138.90574 | 42.782155 | 67.891272 |

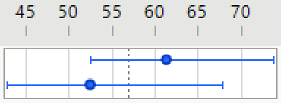

**Wilcoxon / Kruskal-Wallis Tests (Rank Sums)**

| Level | Count | Score Sum | Expected Score | Score Mean | (Mean-Mean0)/Std0 |
|---|---|---|---|---|---|
| Healthy | 68 | 3127.00 | 3638.00 | 45.9853 | -3.391 |
| Demyelinated | 38 | 2544.00 | 2033.00 | 66.9474 | 3.391 |

**Wilcoxon Two-Sample Test, Normal Approximation**

| S | Z | Prob>\|Z\| |
|---|---|---|
| 2544 | 3.39145 | 0.0007* |

**Kruskal-Wallis Test, ChiSquare Approximation**

| ChiSquare | DF | Prob>ChiSq |
|---|---|---|
| 11.5245 | 1 | 0.0007* |

Excluded Rows 9226

**S8B**

#### Oneway Analysis of dark_latency (ms) By mouse_condition

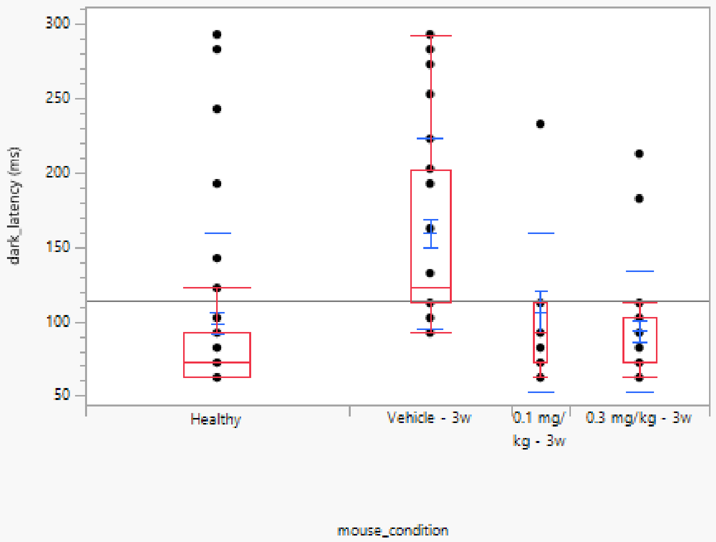

##### Quantiles

| Level | Minimum | 10% | 25% | Median | 75% | 90% | Maximum |
|---|---|---|---|---|---|---|---|
| Healthy | 62.20884 | 62.20884 | 62.20884 | 72.249 | 92.32932 | 197.751 | 293.1325 |
| Vehicle - 3w | 92.32932 | 102.3695 | 112.4096 | 122.4498 | 202.7711 | 267.0281 | 293.1325 |
| 0.1 mg/kg - 3w | 62.20884 | 62.20884 | 72.249 | 92.32932 | 112.4096 | 232.8916 | 232.8916 |
| 0.3 mg/kg - 3w | 62.20884 | 62.20884 | 72.249 | 72.249 | 102.3695 | 182.6908 | 212.8112 |

##### Means and Std Deviations

| Level | Number | Mean | Std Dev | Std Err Mean | Lower 95% | Upper 95% | Std Dev Lower 95% | Std Dev Upper 95% |
|---|---|---|---|---|---|---|---|---|
| Healthy | 68 | 98.825892 | 61.32427 | 7.4366601 | 83.982256 | 113.66953 | 52.47027 | 73.800978 |
| Vehicle - 3w | 42 | 159.26372 | 63.718583 | 9.8319908 | 139.40761 | 179.11983 | 52.427971 | 81.251745 |
| 0.1 mg/kg - 3w | 15 | 106.38554 | 53.907876 | 13.918954 | 76.532355 | 136.23873 | 39.467389 | 85.018058 |
| 0.3 mg/kg - 3w | 36 | 93.444891 | 40.572883 | 6.7621471 | 79.717002 | 107.17278 | 32.907918 | 52.924812 |

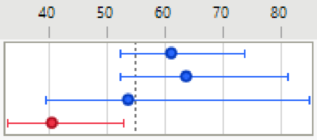

##### Wilcoxon / Kruskal-Wallis Tests (Rank Sums)

| Level | Count | Score Sum | Expected Score | Score Mean | (Mean-Mean0)/Std0 |
|---|---|---|---|---|---|
| Healthy | 68 | 4241.00 | 5508.00 | 62.368 | -4.380 |
| Vehicle - 3w | 42 | 5149.50 | 3402.00 | 122.607 | 6.796 |
| 0.1 mg/kg - 3w | 15 | 1218.50 | 1215.00 | 81.233 | 0.018 |
| 0.3 mg/kg - 3w | 36 | 2432.00 | 2916.00 | 67.556 | -1.982 |

###### Kruskal-Wallis Test, ChiSquare Approximation

| ChiSquare | DF | Prob>ChiSq |
|---|---|---|
| 48.2995 | 3 | <.0001* |

##### Nonparametric Comparisons For All Pairs Using Steel-Dwass Method

| q* | Alpha |
|---|---|
| 2.56903 | 0.05 |

| Level | - Level | Score Mean Difference | Std Err Dif | Z | p-Value | Hodges-Lehmann | Lower CL | Upper CL | Difference Plot |
|---|---|---|---|---|---|---|---|---|---|
| Vehicle - 3w | Healthy | 38.1495 | 6.214371 | 6.13892 | <.0001* | 50.2008 | 40.161 | 90.3614 | |
| 0.1 mg/kg - 3w | Healthy | 11.1887 | 6.735922 | 1.66105 | 0.3445 | 10.0402 | -10.040 | 30.1205 | |
| 0.3 mg/kg - 3w | Healthy | 5.8415 | 6.042313 | 0.96677 | 0.7684 | 0.0000 | -10.040 | 10.0402 | |
| 0.3 mg/kg - 3w | 0.1 mg/kg - 3w | -5.7139 | 4.449220 | -1.28425 | 0.5730 | -10.0402 | -30.120 | 10.0402 | |
| 0.1 mg/kg - 3w | Vehicle - 3w | -17.6429 | 4.938521 | -3.57250 | 0.0020* | -40.1606 | -100.402 | -20.0803 | |
| 0.3 mg/kg - 3w | Vehicle - 3w | -28.9147 | 5.097344 | -5.67250 | <.0001* | -40.1606 | -90.361 | -30.1205 | |

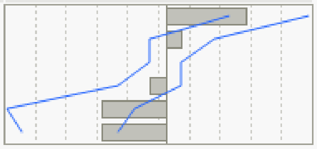

Excluded Rows 9171

**S8C**

#### Oneway Analysis of dark_latency (ms) By mouse_condition

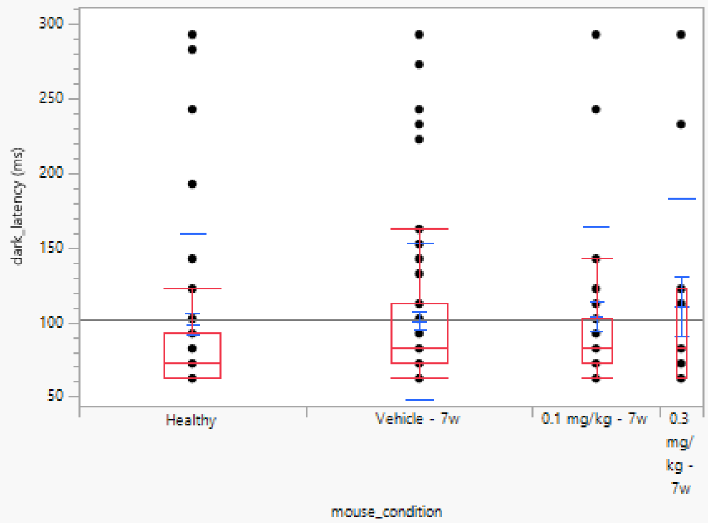

##### Quantiles

| Level | Minimum | 10% | 25% | Median | 75% | 90% | Maximum |
|---|---|---|---|---|---|---|---|
| Healthy | 62.20884 | 62.20884 | 62.20884 | 72.249 | 92.32932 | 197.751 | 293.1325 |
| Vehicle - 7w | 62.20884 | 62.20884 | 72.249 | 82.28916 | 112.4096 | 168.6345 | 293.1325 |
| 0.1 mg/kg - 7w | 62.20884 | 62.20884 | 72.249 | 82.28916 | 102.3695 | 242.9317 | 293.1325 |
| 0.3 mg/kg - 7w | 62.20884 | 62.20884 | 62.20884 | 82.28916 | 122.4498 | 269.0361 | 293.1325 |

##### Means and Std Deviations

| Level | Number | Mean | Std Dev | Std Err Mean | Lower 95% | Upper 95% | Std Dev Lower 95% | Std Dev Upper 95% |
|---|---|---|---|---|---|---|---|---|
| Healthy | 68 | 98.825892 | 61.32427 | 7.4366601 | 83.982256 | 113.66953 | 52.47027 | 73.800978 |
| Vehicle - 7w | 68 | 101.04063 | 52.583977 | 6.3767438 | 88.312599 | 113.76867 | 44.991901 | 63.282432 |
| 0.1 mg/kg - 7w | 38 | 104.21898 | 60.595184 | 9.8298369 | 84.30184 | 124.13612 | 49.40106 | 78.394853 |
| 0.3 mg/kg - 7w | 13 | 110.865 | 72.150292 | 20.01089 | 67.265014 | 154.46498 | 51.737963 | 119.10102 |

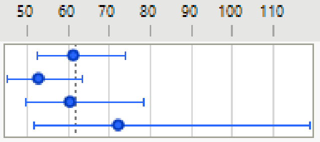

##### Wilcoxon / Kruskal-Wallis Tests (Rank Sums)

| Level | Count | Score Sum | Expected Score | Score Mean | (Mean-Mean0)/Std0 |
|---|---|---|---|---|---|
| Healthy | 68 | 5785.00 | 6392.00 | 85.074 | -1.729 |
| Vehicle - 7w | 68 | 6704.50 | 6392.00 | 98.596 | 0.889 |
| 0.1 mg/kg - 7w | 38 | 3842.00 | 3572.00 | 101.105 | 0.918 |
| 0.3 mg/kg - 7w | 13 | 1246.50 | 1222.00 | 95.885 | 0.129 |

###### Kruskal-Wallis Test, ChiSquare Approximation

| ChiSquare | DF | Prob>ChiSq |
|---|---|---|
| 3.1012 | 3 | 0.3763 |

Excluded Rows 9145

**S8D**

#### Oneway Analysis of bright_latency (ms) By mouse_condition

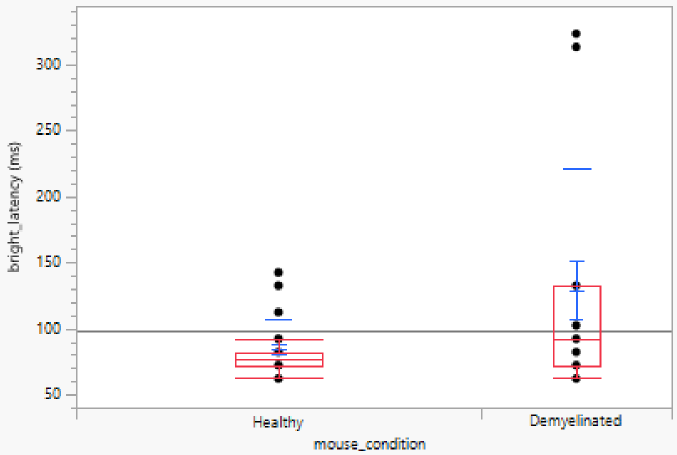

##### Quantiles

| Level | Minimum | 10% | 25% | Median | 75% | 90% | Maximum |
|---|---|---|---|---|---|---|---|
| Healthy | 62.20884 | 62.20884 | 72.249 | 77.26908 | 82.28916 | 132.49 | 142.5301 |
| Demyelinated | 62.20884 | 62.20884 | 72.249 | 92.32932 | 132.49 | 315.2209 | 323.253 |

##### Means and Std Deviations

| Level | Number | Mean | Std Dev | Std Err Mean | Lower 95% | Upper 95% | Std Dev Lower 95% | Std Dev Upper 95% |
|---|---|---|---|---|---|---|---|---|
| Healthy | 36 | 84.520303 | 22.141894 | 3.6903157 | 77.028564 | 92.012043 | 17.958882 | 28.882729 |
| Demyelinated | 17 | 128.94637 | 91.809925 | 22.267178 | 81.742066 | 176.15068 | 68.377263 | 139.72818 |

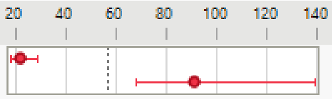

##### Wilcoxon / Kruskal-Wallis Tests (Rank Sums)

| Level | Count | Score Sum | Expected Score | Score Mean | (Mean-Mean0)/Std0 |
|---|---|---|---|---|---|
| Healthy | 36 | 838.000 | 972.000 | 23.2778 | -2.578 |
| Demyelinated | 17 | 593.000 | 459.000 | 34.8824 | 2.578 |

###### Wilcoxon Two-Sample Test, Normal Approximation

| S | Z | Prob>\|Z\| |
|---|---|---|
| 593 | 2.57816 | 0.0099* |

###### Kruskal-Wallis Test, ChiSquare Approximation

| ChiSquare | DF | Prob>ChiSq |
|---|---|---|
| 6.6968 | 1 | 0.0097* |

Excluded Rows 9279

**S8E**

#### Oneway Analysis of bright_latency (ms) By mouse_condition

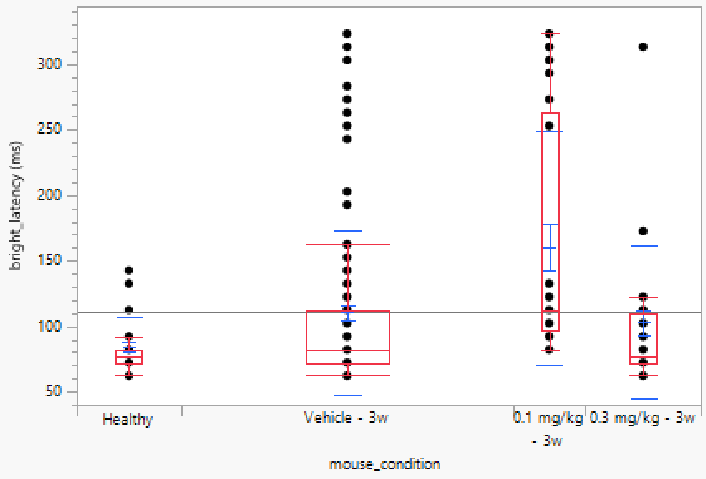

##### Quantiles

| Level | Minimum | 10% | 25% | Median | 75% | 90% | Maximum |
|---|---|---|---|---|---|---|---|
| Healthy | 62.20884 | 62.20884 | 72.249 | 77.26908 | 82.28916 | 132.49 | 142.5301 |
| Vehicle - 3w | 62.20884 | 62.20884 | 72.249 | 82.28916 | 112.4096 | 202.7711 | 323.253 |
| 0.1 mg/kg - 3w | 82.28916 | 88.31325 | 97.3494 | 112.4096 | 263.012 | 317.2289 | 323.253 |
| 0.3 mg/kg - 3w | 62.20884 | 63.21285 | 72.249 | 77.26908 | 109.8996 | 172.6506 | 313.2129 |

##### Means and Std Deviations

| Level | Number | Mean | Std Dev | Std Err Mean | Lower 95% | Upper 95% | Std Dev Lower 95% | Std Dev Upper 95% |
|---|---|---|---|---|---|---|---|---|
| Healthy | 36 | 84.520303 | 22.141894 | 3.6903157 | 77.028564 | 92.012043 | 17.958882 | 28.882729 |
| Vehicle - 3w | 115 | 110.48891 | 62.628756 | 5.8401617 | 98.919597 | 122.05823 | 55.447544 | 71.963775 |
| 0.1 mg/kg - 3w | 25 | 160.2008 | 89.299125 | 17.859825 | 123.33994 | 197.06167 | 69.72729 | 124.22867 |
| 0.3 mg/kg - 3w | 40 | 102.87149 | 58.27597 | 9.21424 | 84.233926 | 121.50905 | 47.737403 | 74.828414 |

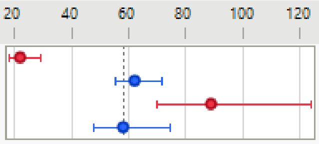

##### Wilcoxon / Kruskal-Wallis Tests (Rank Sums)

| Level | Count | Score Sum | Expected Score | Score Mean | (Mean-Mean0)/Std0 |
|---|---|---|---|---|---|
| Healthy | 36 | 3025.00 | 3906.00 | 84.028 | -2.601 |
| Vehicle - 3w | 115 | 12447.0 | 12477.5 | 108.235 | -0.066 |
| 0.1 mg/kg - 3w | 25 | 4058.00 | 2712.50 | 162.320 | 4.629 |
| 0.3 mg/kg - 3w | 40 | 3906.00 | 4340.00 | 97.650 | -1.229 |

###### Kruskal-Wallis Test, ChiSquare Approximation

| ChiSquare | DF | Prob>ChiSq |
|---|---|---|
| 25.8452 | 3 | <.0001* |

##### Nonparametric Comparisons For All Pairs Using Steel-Dwass Method

| q* | Alpha |
|---|---|
| 2.56903 | 0.05 |

| Level | - Level | Score Mean Difference | Std Err Dif | Z | p-Value | Hodges-Lehmann | Lower CL | Upper CL |
|---|---|---|---|---|---|---|---|---|
| 0.1 mg/kg - 3w | Vehicle - 3w | 34.7687 | 8.885126 | 3.91313 | 0.0005* | 30.1205 | 10.0402 | 50.2008 |
| 0.1 mg/kg - 3w | Healthy | 23.0106 | 4.567494 | 5.03790 | <.0001* | 40.1606 | 20.0803 | 60.2410 |
| Vehicle - 3w | Healthy | 16.9419 | 8.235572 | 2.05716 | 0.1674 | 10.0402 | 0.0000 | 20.0803 |
| 0.3 mg/kg - 3w | Healthy | 3.9847 | 4.894002 | 0.81421 | 0.8478 | 0.0000 | -10.0402 | 20.0803 |
| 0.3 mg/kg - 3w | Vehicle - 3w | -7.3625 | 8.144081 | -0.90403 | 0.8027 | 0.0000 | -10.0402 | 10.0402 |
| 0.3 mg/kg - 3w | 0.1 mg/kg - 3w | -18.8825 | 4.772623 | -3.95642 | 0.0004* | -30.1205 | -60.2410 | -10.0402 |

Difference Plot

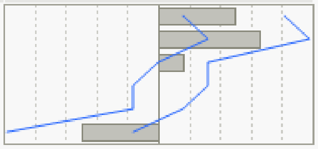

Excluded Rows 9116

**S8F**

#### Oneway Analysis of bright_latency (ms) By mouse_condition

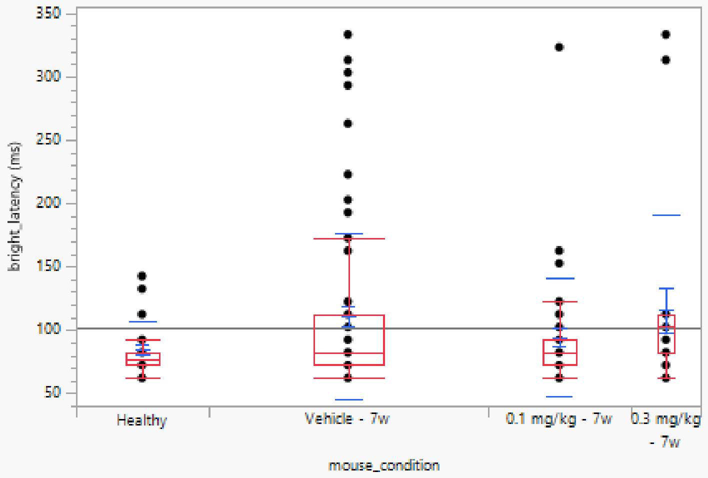

##### Quantiles

| Level | Minimum | 10% | 25% | Median | 75% | 90% | Maximum |
|---|---|---|---|---|---|---|---|
| Healthy | 62.20884 | 62.20884 | 72.249 | 77.26908 | 82.28916 | 132.49 | 142.5301 |
| Vehicle - 7w | 62.20884 | 69.23695 | 72.249 | 82.28916 | 112.4096 | 208.7952 | 333.2932 |
| 0.1 mg/kg - 7w | 62.20884 | 62.20884 | 72.249 | 82.28916 | 92.32932 | 152.5703 | 323.253 |
| 0.3 mg/kg - 7w | 62.20884 | 62.20884 | 82.28916 | 102.3695 | 112.4096 | 313.2129 | 333.2932 |

##### Means and Std Deviations

| Level | Number | Mean | Std Dev | Std Err Mean | Lower 95% | Upper 95% | Std Dev Lower 95% | Std Dev Upper 95% |
|---|---|---|---|---|---|---|---|---|
| Healthy | 36 | 84.520303 | 22.141894 | 3.6903157 | 77.028564 | 92.012043 | 17.958882 | 28.882729 |
| Vehicle - 7w | 76 | 110.95646 | 65.69885 | 7.5361749 | 95.943627 | 125.96929 | 56.659587 | 78.196708 |
| 0.1 mg/kg - 7w | 39 | 94.131397 | 46.689264 | 7.4762656 | 78.996489 | 109.26631 | 38.15661 | 60.172107 |
| 0.3 mg/kg - 7w | 19 | 115.58022 | 74.98839 | 17.203517 | 79.436968 | 151.72346 | 56.662175 | 110.89467 |

##### Wilcoxon / Kruskal-Wallis Tests (Rank Sums)

| Level | Count | Score Sum | Expected Score | Score Mean | (Mean-Mean0)/Std0 |
|---|---|---|---|---|---|
| Healthy | 36 | 2486.00 | 3078.00 | 69.056 | -2.289 |
| Vehicle - 7w | 76 | 7146.00 | 6498.00 | 94.026 | 2.059 |
| 0.1 mg/kg - 7w | 39 | 2992.00 | 3334.50 | 76.718 | -1.286 |
| 0.3 mg/kg - 7w | 19 | 1911.00 | 1624.50 | 100.579 | 1.435 |

##### Kruskal-Wallis Test, ChiSquare Approximation

| ChiSquare | DF | Prob>ChiSq |
|---|---|---|
| 9.6015 | 3 | 0.0223* |

##### Nonparametric Comparisons For All Pairs Using Steel-Dwass Method

| q* | Alpha |
|---|---|
| 2.56903 | 0.05 |

| Level | - Level | Score Mean Difference | Std Err Dif | Z | p-Value | Hodges-Lehmann | Lower CL | Upper CL |
|---|---|---|---|---|---|---|---|---|
| Vehicle - 7w | Healthy | 16.2310 | 6.462348 | 2.51162 | 0.0581 | 10.0402 | 0.0000 | 20.08032 |
| 0.3 mg/kg - 7w | Healthy | 10.0914 | 4.449307 | 2.26808 | 0.1056 | 20.0803 | 0.0000 | 30.12048 |
| 0.3 mg/kg - 7w | 0.1 mg/kg - 7w | 8.3752 | 4.649996 | 1.80111 | 0.2727 | 10.0402 | -10.0402 | 30.12048 |
| 0.1 mg/kg - 7w | Healthy | 3.6592 | 4.904044 | 0.74616 | 0.8783 | 0.0000 | -10.0402 | 10.04016 |
| 0.3 mg/kg - 7w | Vehicle - 7w | 3.4539 | 7.005137 | 0.49306 | 0.9607 | 0.0000 | -10.0402 | 20.08032 |
| 0.1 mg/kg - 7w | Vehicle - 7w | -11.7755 | 6.481515 | -1.81678 | 0.2653 | -10.0402 | -20.0803 | 0.00000 |

Excluded Rows 9162

#### Figure S9

## S9A

##### Oneway Analysis of n70_latency (ms) By condition

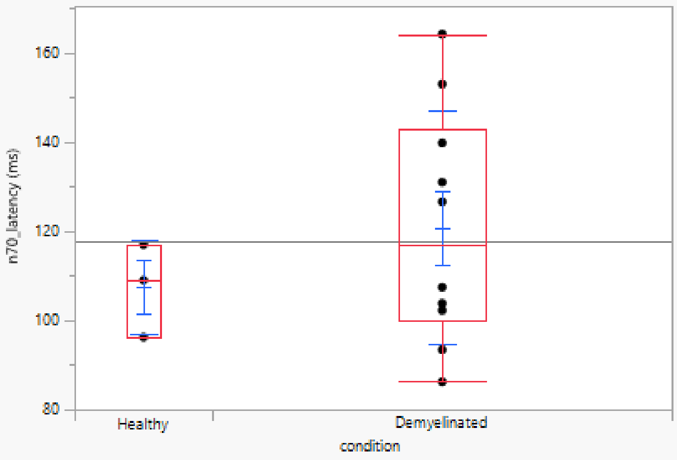

##### Quantiles

| Level | Minimum | 10% | 25% | Median | 75% | 90% | Maximum |
|---|---|---|---|---|---|---|---|
| Healthy | 96.11845 | 96.11845 | 96.11845 | 108.9236 | 116.9268 | 116.9268 | 116.9268 |
| Demyelinated | 86.11445 | 86.83473 | 99.91997 | 116.9268 | 143.0372 | 163.0252 | 164.1457 |

##### Means and Std Deviations

| Level | Number | Mean | Std Dev | Std Err Mean | Lower 95% | Upper 95% | Std Dev Lower 95% | Std Dev Upper 95% |
|---|---|---|---|---|---|---|---|---|
| Healthy | 3 | 107.32293 | 10.4961 | 6.0599262 | 81.249171 | 133.39669 | 5.4648813 | 65.965204 |
| Demyelinated | 10 | 120.68828 | 26.218001 | 8.2908599 | 101.93305 | 139.4435 | 18.033664 | 47.86385 |

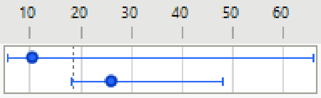

##### Wilcoxon / Kruskal-Wallis Tests (Rank Sums)

| Level | Count | Score Sum | Expected Score | Score Mean | (Mean-Mean0)/Std0 |
|---|---|---|---|---|---|
| Healthy | 3 | 18.000 | 21.000 | 6.00000 | -0.423 |
| Demyelinated | 10 | 73.000 | 70.000 | 7.30000 | 0.423 |

###### Wilcoxon Two-Sample Test, Normal Approximation

| S | Z | Prob>\|Z\| |
|---|---|---|
| 18 | -0.42258 | 0.6726 |

###### Kruskal-Wallis Test, ChiSquare Approximation

| ChiSquare | DF | Prob>ChiSq |
|---|---|---|
| 0.2571 | 1 | 0.6121 |

Excluded Rows 131

**S9B**

#### Oneway Analysis of n70_latency (ms) By condition

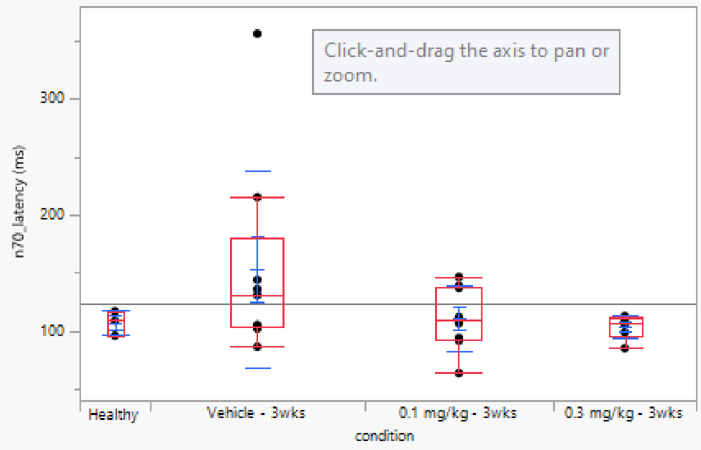

##### Quantiles

| Level | Minimum | 10% | 25% | Median | 75% | 90% | Maximum |
|---|---|---|---|---|---|---|---|
| Healthy | 96.11845 | 96.11845 | 96.11845 | 108.9236 | 116.9268 | 116.9268 | 116.9268 |
| Vehicle - 3wks | 86.51461 | 86.51461 | 103.3213 | 130.9324 | 179.7519 | 356.2225 | 356.2225 |
| 0.1 mg/kg - 3wks | 63.70548 | 63.70548 | 92.31693 | 109.5238 | 138.5354 | 146.5386 | 146.5386 |
| 0.3 mg/kg - 3wks | 85.31413 | 85.31413 | 95.51821 | 106.7227 | 111.1244 | 112.9252 | 112.9252 |

##### Means and Std Deviations

| Level | Number | Mean | Std Dev | Std Err Mean | Lower 95% | Upper 95% | Std Dev Lower 95% | Std Dev Upper 95% |
|---|---|---|---|---|---|---|---|---|
| Healthy | 3 | 107.32293 | 10.4961 | 6.0599262 | 81.249171 | 133.39669 | 5.4648813 | 65.965204 |
| Vehicle - 3wks | 9 | 153.47472 | 84.903744 | 28.301248 | 88.211928 | 218.73752 | 57.348831 | 162.65612 |
| 0.1 mg/kg - 3wks | 8 | 111.42457 | 28.402708 | 10.041874 | 87.679312 | 135.16983 | 18.779136 | 57.807239 |
| 0.3 mg/kg - 3wks | 6 | 103.52141 | 10.12571 | 4.1338039 | 92.895127 | 114.14769 | 6.3205493 | 24.834459 |

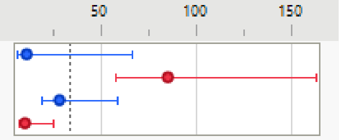

##### Wilcoxon / Kruskal-Wallis Tests (Rank Sums)

| Level | Count | Score Sum | Expected Score | Score Mean | (Mean-Mean0)/Std0 |
|---|---|---|---|---|---|
| Healthy | 3 | 38.000 | 40.500 | 12.6667 | -0.161 |
| Vehicle - 3wks | 9 | 143.000 | 121.500 | 15.8889 | 1.132 |
| 0.1 mg/kg - 3wks | 8 | 105.000 | 108.000 | 13.1250 | -0.139 |
| 0.3 mg/kg - 3wks | 6 | 65.000 | 81.000 | 10.8333 | -0.943 |

###### Kruskal-Wallis Test, ChiSquare Approximation

| ChiSquare | DF | Prob>ChiSq |
|---|---|---|
| 1.6622 | 3 | 0.6454 |

Excluded Rows 118

**S9C**

#### Oneway Analysis of n70_latency (ms) By condition

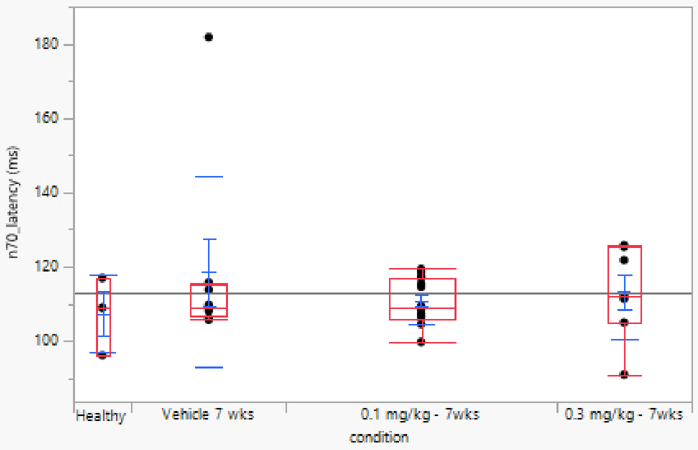

##### Quantiles

| Level | Minimum | 10% | 25% | Median | 75% | 90% | Maximum |
|---|---|---|---|---|---|---|---|
| Healthy | 96.11845 | 96.11845 | 96.11845 | 108.9236 | 116.9268 | 116.9268 | 116.9268 |
| Vehicle 7 wks | 105.7223 | 105.7223 | 106.6226 | 108.9236 | 115.2261 | 181.7527 | 181.7527 |
| 0.1 mg/kg - 7wks | 99.71989 | 102.1208 | 106.0224 | 108.9236 | 117.0268 | 118.9276 | 119.3277 |
| 0.3 mg/kg - 7wks | 90.91637 | 90.91637 | 104.922 | 112.1248 | 125.3301 | 125.7303 | 125.7303 |

##### Means and Std Deviations

| Level | Number | Mean | Std Dev | Std Err Mean | Lower 95% | Upper 95% | Std Dev Lower 95% | Std Dev Upper 95% |
|---|---|---|---|---|---|---|---|---|
| Healthy | 3 | 107.32293 | 10.4961 | 6.0599262 | 81.249171 | 133.39669 | 5.4648813 | 65.965204 |
| Vehicle 7 wks | 8 | 118.62745 | 25.746321 | 9.1026991 | 97.102988 | 140.15191 | 17.022802 | 52.400769 |
| 0.1 mg/kg - 7wks | 14 | 110.78146 | 6.148592 | 1.6432803 | 107.23136 | 114.33155 | 4.4574459 | 9.9056425 |
| 0.3 mg/kg - 7wks | 7 | 113.15383 | 12.560729 | 4.7475092 | 101.5371 | 124.77057 | 8.0940508 | 27.659555 |

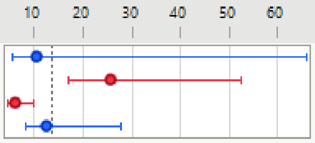

##### Wilcoxon / Kruskal-Wallis Tests (Rank Sums)

| Level | Count | Score Sum | Expected Score | Score Mean | (Mean-Mean0)/Std0 |
|---|---|---|---|---|---|
| Healthy | 3 | 41.500 | 49.500 | 13.8333 | -0.485 |
| Vehicle 7 wks | 8 | 130.000 | 132.000 | 16.2500 | -0.065 |
| 0.1 mg/kg - 7wks | 14 | 222.500 | 231.000 | 15.8929 | -0.304 |
| 0.3 mg/kg - 7wks | 7 | 134.000 | 115.500 | 19.1429 | 0.821 |

###### Kruskal-Wallis Test, ChiSquare Approximation

| ChiSquare | DF | Prob>ChiSq |
|---|---|---|
| 0.8630 | 3 | 0.8344 |

Excluded Rows 112

**S9D**

#### Oneway Analysis of n70_latency (ms) By condition

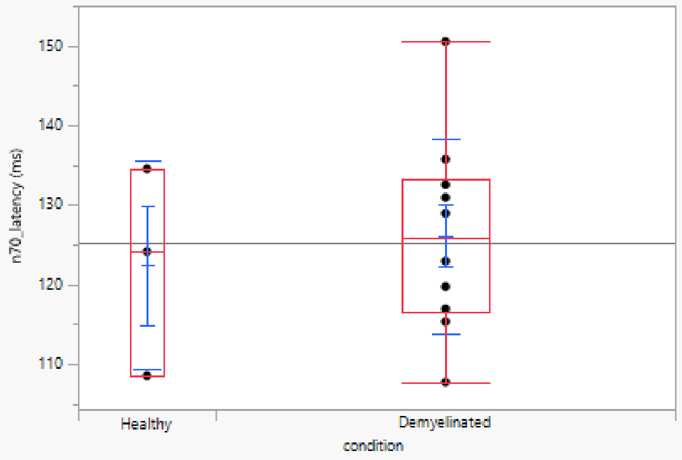

##### Quantiles

| Level | Minimum | 10% | 25% | Median | 75% | 90% | Maximum |
|---|---|---|---|---|---|---|---|
| Healthy | 108.5234 | 108.5234 | 108.5234 | 124.1297 | 134.5338 | 134.5338 | 134.5338 |
| Demyelinated | 107.7231 | 108.4834 | 116.5266 | 125.9304 | 133.3333 | 149.0596 | 150.5402 |

##### Means and Std Deviations

| Level | Number | Mean | Std Dev | Std Err Mean | Lower 95% | Upper 95% | Std Dev Lower 95% | Std Dev Upper 95% |
|---|---|---|---|---|---|---|---|---|
| Healthy | 3 | 122.39562 | 13.091616 | 7.5584482 | 89.874247 | 154.917 | 6.8162583 | 82.277335 |
| Demyelinated | 10 | 126.13045 | 12.254165 | 3.8751071 | 117.36435 | 134.89655 | 8.4288459 | 22.371328 |

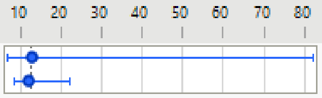

##### Wilcoxon / Kruskal-Wallis Tests (Rank Sums)

| Level | Count | Score Sum | Expected Score | Score Mean | (Mean-Mean0)/Std0 |
|---|---|---|---|---|---|
| Healthy | 3 | 20.000 | 21.000 | 6.66667 | -0.085 |
| Demyelinated | 10 | 71.000 | 70.000 | 7.10000 | 0.085 |

###### Wilcoxon Two-Sample Test, Normal Approximation

| S | Z | Prob>\|Z\| |
|---|---|---|
| 20 | -0.08452 | 0.9326 |

###### Kruskal-Wallis Test, ChiSquare Approximation

| ChiSquare | DF | Prob>ChiSq |
|---|---|---|
| 0.0286 | 1 | 0.8658 |

Excluded Rows 131

**S9E**

#### Oneway Analysis of n70_latency (ms) By condition

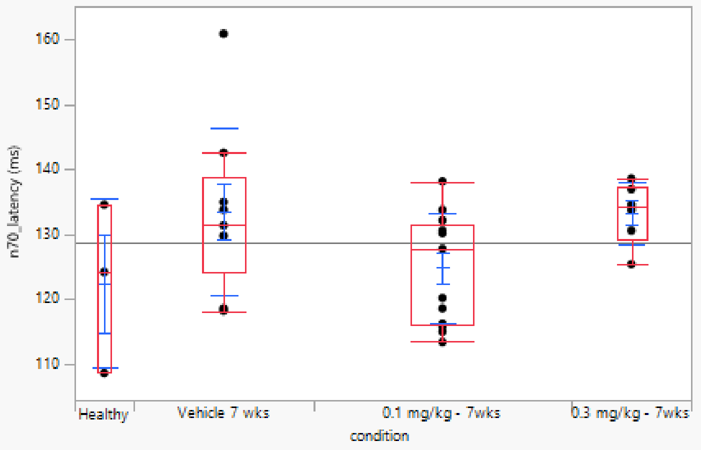

##### Quantiles

| Level | Minimum | 10% | 25% | Median | 75% | 90% | Maximum |
|---|---|---|---|---|---|---|---|
| Healthy | 108.5234 | 108.5234 | 108.5234 | 124.1297 | 134.5338 | 134.5338 | 134.5338 |
| Vehicle 7 wks | 118.1273 | 118.1273 | 124.1297 | 131.3325 | 138.7355 | 160.9444 | 160.9444 |
| 0.1 mg/kg - 7wks | 113.3253 | 113.9656 | 115.9264 | 127.7311 | 131.3325 | 136.3745 | 138.1353 |
| 0.3 mg/kg - 7wks | 125.3301 | 125.3301 | 129.2317 | 134.1337 | 137.3349 | 138.5354 | 138.5354 |

##### Means and Std Deviations

| Level | Number | Mean | Std Dev | Std Err Mean | Lower 95% | Upper 95% | Std Dev Lower 95% | Std Dev Upper 95% |
|---|---|---|---|---|---|---|---|---|
| Healthy | 3 | 122.39562 | 13.091616 | 7.5584482 | 89.874247 | 154.917 | 6.8162583 | 82.277335 |
| Vehicle 7 wks | 9 | 133.46672 | 12.850373 | 4.2834576 | 123.58905 | 143.34439 | 8.6798747 | 24.61837 |
| 0.1 mg/kg - 7wks | 13 | 124.74528 | 8.4754862 | 2.3506769 | 119.6236 | 129.86697 | 6.0776524 | 13.990783 |
| 0.3 mg/kg - 7wks | 6 | 133.26664 | 4.7645381 | 1.9451145 | 128.26656 | 138.26672 | 2.9740628 | 11.685573 |

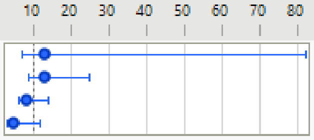

##### Wilcoxon / Kruskal-Wallis Tests (Rank Sums)

| Level | Count | Score Sum | Expected Score | Score Mean | (Mean-Mean0)/Std0 |
|---|---|---|---|---|---|
| Healthy | 3 | 35.500 | 48.000 | 11.8333 | -0.803 |
| Vehicle 7 wks | 9 | 172.500 | 144.000 | 19.1667 | 1.220 |
| 0.1 mg/kg - 7wks | 13 | 158.500 | 208.000 | 12.1923 | -1.964 |
| 0.3 mg/kg - 7wks | 6 | 129.500 | 96.000 | 21.5833 | 1.652 |

###### Kruskal-Wallis Test, ChiSquare Approximation

| ChiSquare | DF | Prob>ChiSq |
|---|---|---|
| 6.2783 | 3 | 0.0988 |

Excluded Rows 113

**S9F**

#### Oneway Analysis of n70_latency (ms) By condition

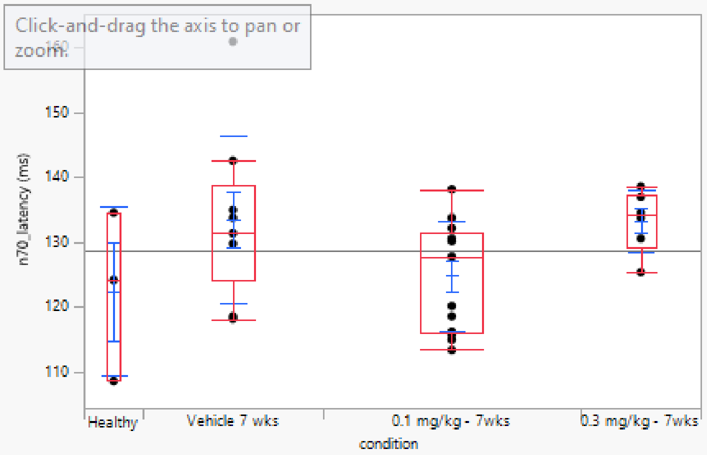

##### Quantiles

| Level | Minimum | 10% | 25% | Median | 75% | 90% | Maximum |
|---|---|---|---|---|---|---|---|
| Healthy | 108.5234 | 108.5234 | 108.5234 | 124.1297 | 134.5338 | 134.5338 | 134.5338 |
| Vehicle 7 wks | 118.1273 | 118.1273 | 124.1297 | 131.3325 | 138.7355 | 160.9444 | 160.9444 |
| 0.1 mg/kg - 7wks | 113.3253 | 113.9656 | 115.9264 | 127.7311 | 131.3325 | 136.3745 | 138.1353 |
| 0.3 mg/kg - 7wks | 125.3301 | 125.3301 | 129.2317 | 134.1337 | 137.3349 | 138.5354 | 138.5354 |

##### Means and Std Deviations

| Level | Number | Mean | Std Dev | Std Err Mean | Lower 95% | Upper 95% | Std Dev Lower 95% | Std Dev Upper 95% |
|---|---|---|---|---|---|---|---|---|
| Healthy | 3 | 122.39562 | 13.091616 | 7.5584482 | 89.874247 | 154.917 | 6.8162583 | 82.277335 |
| Vehicle 7 wks | 9 | 133.46672 | 12.850373 | 4.2834576 | 123.58905 | 143.34439 | 8.6798747 | 24.61837 |
| 0.1 mg/kg - 7wks | 13 | 124.74528 | 8.4754862 | 2.3506769 | 119.6236 | 129.86697 | 6.0776524 | 13.990783 |
| 0.3 mg/kg - 7wks | 6 | 133.26664 | 4.7645381 | 1.9451145 | 128.26656 | 138.26672 | 2.9740628 | 11.685573 |

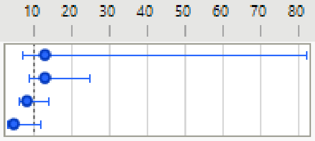

##### Wilcoxon / Kruskal-Wallis Tests (Rank Sums)

| Level | Count | Score Sum | Expected Score | Score Mean | (Mean-Mean0)/Std0 |
|---|---|---|---|---|---|
| Healthy | 3 | 35.500 | 48.000 | 11.8333 | -0.803 |
| Vehicle 7 wks | 9 | 172.500 | 144.000 | 19.1667 | 1.220 |
| 0.1 mg/kg - 7wks | 13 | 158.500 | 208.000 | 12.1923 | -1.964 |
| 0.3 mg/kg - 7wks | 6 | 129.500 | 96.000 | 21.5833 | 1.652 |

###### Kruskal-Wallis Test, ChiSquare Approximation

| ChiSquare | DF | Prob>ChiSq |
|---|---|---|
| 6.2783 | 3 | 0.0988 |

Excluded Rows 113

**S9G**

#### Oneway Analysis of correlation_baseline By condition

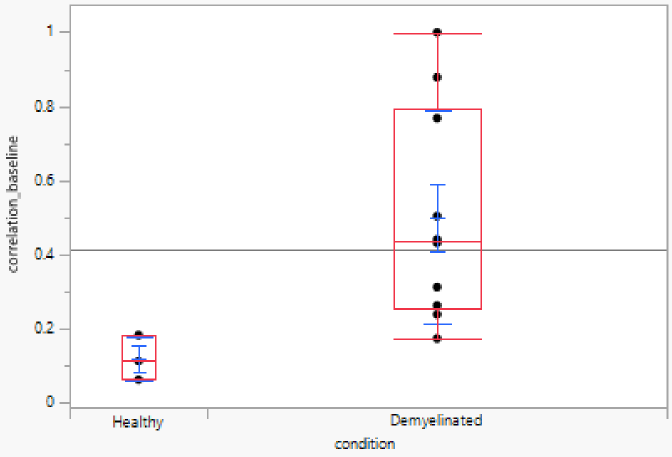

##### Quantiles

| Level | Minimum | 10% | 25% | Median | 75% | 90% | Maximum |
|---|---|---|---|---|---|---|---|
| Healthy | 0.060319 | 0.060319 | 0.060319 | 0.11044 | 0.180706 | 0.180706 | 0.180706 |
| Demyelinated | 0.171548 | 0.178071 | 0.25505 | 0.434862 | 0.79531 | 0.987519 | 0.999668 |

##### Means and Std Deviations

| Level | Number | Mean | Std Dev | Std Err Mean | Lower 95% | Upper 95% | Std Dev Lower 95% | Std Dev Upper 95% |
|---|---|---|---|---|---|---|---|---|
| Healthy | 3 | 0.117155 | 0.060474 | 0.0349147 | -0.033071 | 0.2673807 | 0.0314863 | 0.380063 |
| Demyelinated | 10 | 0.4998148 | 0.2873136 | 0.0908565 | 0.294283 | 0.7053466 | 0.1976244 | 0.5245226 |

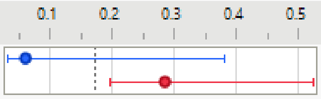

##### Wilcoxon / Kruskal-Wallis Tests (Rank Sums)

| Level | Count | Score Sum | Expected Score | Score Mean | (Mean-Mean0)/Std0 |
|---|---|---|---|---|---|
| Healthy | 3 | 7.000 | 21.000 | 2.33333 | -2.282 |
| Demyelinated | 10 | 84.000 | 70.000 | 8.40000 | 2.282 |

###### Wilcoxon Two-Sample Test, Normal Approximation

| S | Z | Prob>\|Z\| |
|---|---|---|
| 7 | -2.28192 | 0.0225* |

###### Kruskal-Wallis Test, ChiSquare Approximation

| ChiSquare | DF | Prob>ChiSq |
|---|---|---|
| 5.6000 | 1 | 0.0180* |

Excluded Rows 131

**S9H**

#### Oneway Analysis of correlation_baseline By condition

Save Script to Data Table

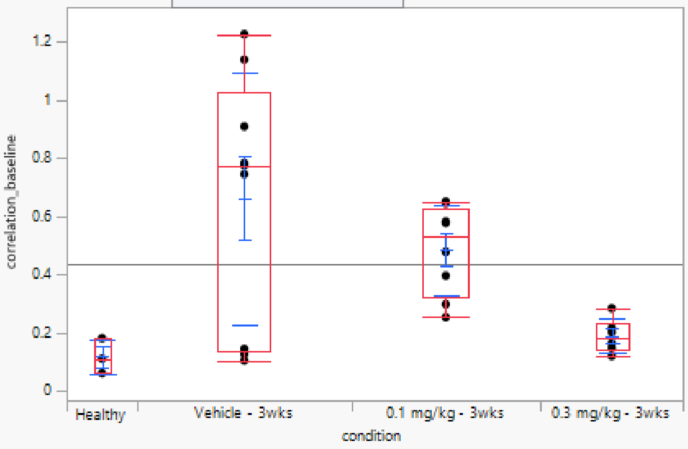

##### Quantiles

| Level | Minimum | 10% | 25% | Median | 75% | 90% | Maximum |
|---|---|---|---|---|---|---|---|
| Healthy | 0.060319 | 0.060319 | 0.060319 | 0.11044 | 0.180706 | 0.180706 | 0.180706 |
| Vehicle - 3wks | 0.103788 | 0.103788 | 0.135375 | 0.774634 | 1.022996 | 1.225352 | 1.225352 |
| 0.1 mg/kg - 3wks | 0.253301 | 0.253301 | 0.322071 | 0.527529 | 0.628558 | 0.650203 | 0.650203 |
| 0.3 mg/kg - 3wks | 0.119767 | 0.119767 | 0.140092 | 0.182398 | 0.232274 | 0.283427 | 0.283427 |

##### Means and Std Deviations

| Level | Number | Mean | Std Dev | Std Err Mean | Lower 95% | Upper 95% | Std Dev Lower 95% | Std Dev Upper 95% |
|---|---|---|---|---|---|---|---|---|
| Healthy | 3 | 0.117155 | 0.060474 | 0.0349147 | -0.033071 | 0.2673807 | 0.0314863 | 0.380063 |
| Vehicle - 3wks | 9 | 0.6608038 | 0.4336982 | 0.1445661 | 0.3274338 | 0.9941737 | 0.2929445 | 0.8308664 |
| 0.1 mg/kg - 3wks | 8 | 0.4846653 | 0.154608 | 0.0546622 | 0.3554098 | 0.6139208 | 0.1022228 | 0.3146693 |
| 0.3 mg/kg - 3wks | 6 | 0.1883466 | 0.058276 | 0.0237911 | 0.1271897 | 0.2495036 | 0.0363764 | 0.1429286 |

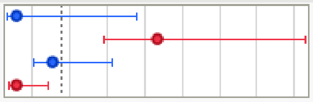

##### Wilcoxon / Kruskal-Wallis Tests (Rank Sums)

| Level | Count | Score Sum | Expected Score | Score Mean | (Mean-Mean0)/Std0 |
|---|---|---|---|---|---|
| Healthy | 3 | 13.000 | 40.500 | 4.3333 | -2.167 |
| Vehicle - 3wks | 9 | 154.000 | 121.500 | 17.1111 | 1.725 |
| 0.1 mg/kg - 3wks | 8 | 131.000 | 108.000 | 16.3750 | 1.250 |
| 0.3 mg/kg - 3wks | 6 | 53.000 | 81.000 | 8.8333 | -1.674 |

###### Kruskal-Wallis Test, ChiSquare Approximation

| ChiSquare | DF | Prob>ChiSq |
|---|---|---|
| 9.6792 | 3 | 0.0215* |

##### Nonparametric Comparisons For All Pairs Using Steel-Dwass Method

| q* | Alpha |
|---|---|
| 2.56903 | 0.05 |

| Level | - Level | Score Mean Difference | Std Err Dif | Z | p-Value | Hodges-Lehmann | Lower CL | Upper CL |
|---|---|---|---|---|---|---|---|---|
| 0.1 mg/kg - 3wks | Healthy | 5.27083 | 2.245366 | 2.34743 | 0.0876 | 0.398589 | . | . |
| Vehicle - 3wks | Healthy | 4.00000 | 2.403701 | 1.66410 | 0.3428 | 0.664194 | . | . |
| 0.3 mg/kg - 3wks | Healthy | 2.75000 | 1.936492 | 1.42009 | 0.4867 | 0.072998 | . | . |
| 0.1 mg/kg - 3wks | Vehicle - 3wks | -2.71528 | 2.453739 | -1.10659 | 0.6855 | -0.231454 | -0.66010 | 0.473508 |
| 0.3 mg/kg - 3wks | Vehicle - 3wks | -2.91667 | 2.357023 | -1.23744 | 0.6030 | -0.580961 | -1.02394 | 0.111435 |
| 0.3 mg/kg - 3wks | 0.1 mg/kg - 3wks | -6.56250 | 2.259240 | -2.90474 | 0.0193* | -0.322639 | -0.50334 | -0.038078 |

Difference Plot

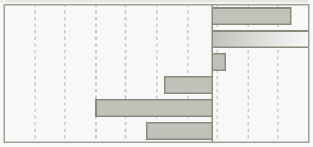

Excluded Rows 118

## S9I

##### Oneway Analysis of correlation_baseline By condition

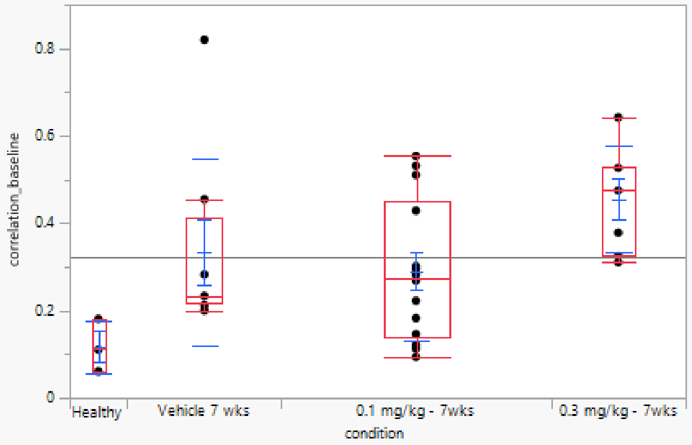

##### Quantiles

| Level | Minimum | 10% | 25% | Median | 75% | 90% | Maximum |
|---|---|---|---|---|---|---|---|
| Healthy | 0.060319 | 0.060319 | 0.060319 | 0.11044 | 0.180706 | 0.180706 | 0.180706 |
| Vehicle 7 wks | 0.199829 | 0.199829 | 0.217467 | 0.233916 | 0.41135 | 0.819639 | 0.819639 |
| 0.1 mg/kg - 7wks | 0.093485 | 0.102956 | 0.13952 | 0.274842 | 0.44889 | 0.542582 | 0.553502 |
| 0.3 mg/kg - 7wks | 0.310181 | 0.310181 | 0.324872 | 0.474989 | 0.527755 | 0.642 | 0.642 |

##### Means and Std Deviations

| Level | Number | Mean | Std Dev | Std Err Mean | Lower 95% | Upper 95% | Std Dev Lower 95% | Std Dev Upper 95% |
|---|---|---|---|---|---|---|---|---|
| Healthy | 3 | 0.117155 | 0.060474 | 0.0349147 | -0.033071 | 0.2673807 | 0.0314863 | 0.380063 |
| Vehicle 7 wks | 8 | 0.3336657 | 0.2124854 | 0.0751249 | 0.1560235 | 0.5113079 | 0.1404898 | 0.4324655 |
| 0.1 mg/kg - 7wks | 14 | 0.2890919 | 0.1597682 | 0.0426998 | 0.1968445 | 0.3813393 | 0.1158246 | 0.2573933 |
| 0.3 mg/kg - 7wks | 7 | 0.4548611 | 0.122192 | 0.0461842 | 0.3418523 | 0.5678699 | 0.0787397 | 0.2690749 |

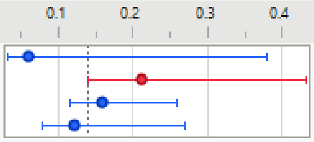

##### Wilcoxon / Kruskal-Wallis Tests (Rank Sums)

| Level | Count | Score Sum | Expected Score | Score Mean | (Mean-Mean0)/Std0 |
|---|---|---|---|---|---|
| Healthy | 3 | 11.000 | 49.500 | 3.6667 | -2.457 |
| Vehicle 7 wks | 8 | 131.000 | 132.000 | 16.3750 | -0.022 |
| 0.1 mg/kg - 7wks | 14 | 212.000 | 231.000 | 15.1429 | -0.703 |
| 0.3 mg/kg - 7wks | 7 | 174.000 | 115.500 | 24.8571 | 2.644 |

###### Kruskal-Wallis Test, ChiSquare Approximation

| ChiSquare | DF | Prob>ChiSq |
|---|---|---|
| 11.4646 | 3 | 0.0095* |

##### Nonparametric Comparisons For All Pairs Using Steel-Dwass Method

| q* | Alpha |
|---|---|
| 2.56903 | 0.05 |

| Level | - Level | Score Mean Difference | Std Err Dif | Z | p-Value | Hodges-Lehmann | Lower CL | Upper CL |
|---|---|---|---|---|---|---|---|---|
| 0.3 mg/kg - 7wks | 0.1 mg/kg - 7wks | 6.32143 | 2.872281 | 2.20084 | 0.1230 | 0.191148 | -0.050526 | 0.3815043 |
| 0.1 mg/kg - 7wks | Healthy | 6.27381 | 3.212698 | 1.95282 | 0.2062 | 0.139792 | . | . |
| Vehicle 7 wks | Healthy | 5.27083 | 2.245366 | 2.34743 | 0.0876 | 0.145705 | . | . |
| 0.3 mg/kg - 7wks | Healthy | 4.76190 | 2.089277 | 2.27921 | 0.1029 | 0.345575 | . | . |
| 0.3 mg/kg - 7wks | Vehicle 7 wks | 4.68750 | 2.314550 | 2.02523 | 0.1786 | 0.144705 | -0.293358 | 0.4074480 |
| 0.1 mg/kg - 7wks | Vehicle 7 wks | -0.88393 | 2.877975 | -0.30714 | 0.9900 | -0.015841 | -0.287978 | 0.2494641 |

Difference Plot

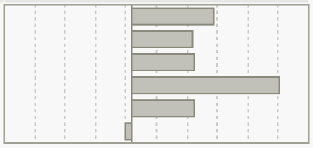

Excluded Rows 112

**S9J**

#### Oneway Analysis of correlation_baseline By condition

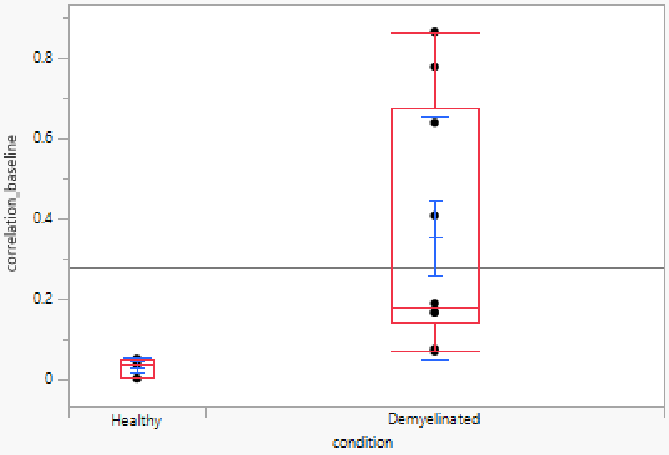

##### Quantiles

| Level | Minimum | 10% | 25% | Median | 75% | 90% | Maximum |
|---|---|---|---|---|---|---|---|
| Healthy | 0.004105 | 0.004105 | 0.004105 | 0.038322 | 0.053009 | 0.053009 | 0.053009 |
| Demyelinated | 0.071835 | 0.072418 | 0.144539 | 0.179606 | 0.674529 | 0.85635 | 0.865029 |

##### Means and Std Deviations

| Level | Number | Mean | Std Dev | Std Err Mean | Lower 95% | Upper 95% | Std Dev Lower 95% | Std Dev Upper 95% |
|---|---|---|---|---|---|---|---|---|
| Healthy | 3 | 0.0318117 | 0.0250936 | 0.0144878 | -0.030524 | 0.0941477 | 0.0130652 | 0.1577067 |
| Demyelinated | 10 | 0.3536042 | 0.3004476 | 0.0950099 | 0.138677 | 0.5685315 | 0.2066584 | 0.5485002 |

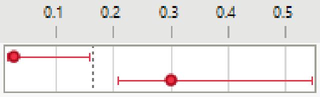

##### Wilcoxon / Kruskal-Wallis Tests (Rank Sums)

| Level | Count | Score Sum | Expected Score | Score Mean | (Mean-Mean0)/Std0 |
|---|---|---|---|---|---|
| Healthy | 3 | 6.000 | 21.000 | 2.00000 | -2.451 |
| Demyelinated | 10 | 85.000 | 70.000 | 8.50000 | 2.451 |

###### Wilcoxon Two-Sample Test, Normal Approximation

| S | Z | Prob>\|Z\| |
|---|---|---|
| 6 | -2.45095 | 0.0142* |

###### Kruskal-Wallis Test, ChiSquare Approximation

| ChiSquare | DF | Prob>ChiSq |
|---|---|---|
| 6.4286 | 1 | 0.0112* |

Excluded Rows 131

## S9K

##### Oneway Analysis of correlation_baseline By condition

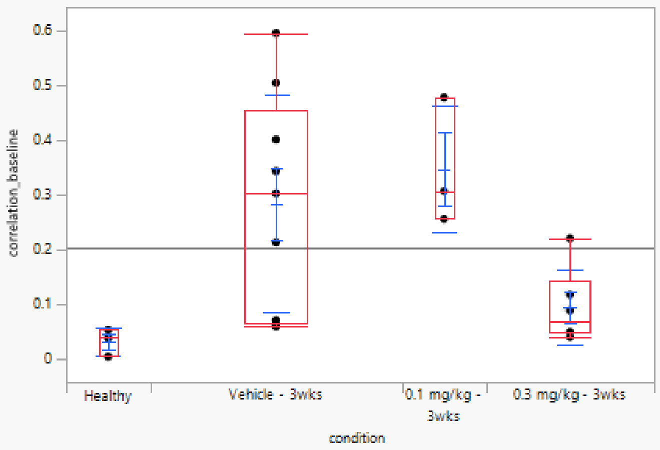

##### Quantiles

| Level | Minimum | 10% | 25% | Median | 75% | 90% | Maximum |
|---|---|---|---|---|---|---|---|
| Healthy | 0.004105 | 0.004105 | 0.004105 | 0.038322 | 0.053009 | 0.053009 | 0.053009 |
| Vehicle - 3wks | 0.059003 | 0.059003 | 0.066269 | 0.302267 | 0.452715 | 0.595474 | 0.595474 |
| 0.1 mg/kg - 3wks | 0.255603 | 0.255603 | 0.255603 | 0.306681 | 0.478051 | 0.478051 | 0.478051 |
| 0.3 mg/kg - 3wks | 0.040074 | 0.040074 | 0.046984 | 0.06913 | 0.142846 | 0.2199 | 0.2199 |

##### Means and Std Deviations

| Level | Number | Mean | Std Dev | Std Err Mean | Lower 95% | Upper 95% | Std Dev Lower 95% | Std Dev Upper 95% |
|---|---|---|---|---|---|---|---|---|
| Healthy | 3 | 0.0318117 | 0.0250936 | 0.0144878 | -0.030524 | 0.0941477 | 0.0130652 | 0.1577067 |
| Vehicle - 3wks | 9 | 0.2834465 | 0.1980152 | 0.0660051 | 0.1312385 | 0.4356544 | 0.1337508 | 0.3793518 |
| 0.1 mg/kg - 3wks | 3 | 0.3467784 | 0.1165184 | 0.0672719 | 0.0573305 | 0.6362262 | 0.0606663 | 0.7322874 |
| 0.3 mg/kg - 3wks | 6 | 0.0941138 | 0.0682179 | 0.0278498 | 0.0225236 | 0.1657041 | 0.0425821 | 0.1673121 |

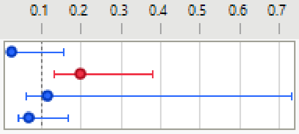

##### Wilcoxon / Kruskal-Wallis Tests (Rank Sums)

| Level | Count | Score Sum | Expected Score | Score Mean | (Mean-Mean0)/Std0 |
|---|---|---|---|---|---|
| Healthy | 3 | 9.000 | 33.000 | 3.0000 | -2.362 |
| Vehicle - 3wks | 9 | 127.000 | 99.000 | 14.1111 | 1.954 |
| 0.1 mg/kg - 3wks | 3 | 49.000 | 33.000 | 16.3333 | 1.558 |
| 0.3 mg/kg - 3wks | 6 | 46.000 | 66.000 | 7.6667 | -1.518 |

###### Kruskal-Wallis Test, ChiSquare Approximation

| ChiSquare | DF | Prob>ChiSq |
|---|---|---|
| 11.1977 | 3 | 0.0107* |

Small sample sizes. Refer to statistical tables for tests, rather than large-sample approximations.

##### Nonparametric Comparisons For All Pairs Using Steel-Dwass Method

| q* | Alpha |
|---|---|
| 2.56903 | 0.05 |

| Level | - Level | Score Mean Difference | Std Err Dif | Z | p-Value | Hodges-Lehmann | Lower CL | Upper CL |
|---|---|---|---|---|---|---|---|---|
| Vehicle - 3wks | Healthy | 5.77778 | 2.403701 | 2.40370 | 0.0764 | 0.263945 | . | . |
| 0.3 mg/kg - 3wks | Healthy | 2.75000 | 1.936492 | 1.42009 | 0.4867 | 0.045507 | . | . |
| 0.1 mg/kg - 3wks | Healthy | 2.66667 | 1.527525 | 1.74574 | 0.3000 | 0.268359 | . | . |
| 0.1 mg/kg - 3wks | Vehicle - 3wks | 0.88889 | 2.403701 | 0.36980 | 0.9827 | 0.077063 | . | . |
| 0.3 mg/kg - 3wks | 0.1 mg/kg - 3wks | -4.25000 | 1.936492 | -2.19469 | 0.1247 | -0.237551 | . | . |
| 0.3 mg/kg - 3wks | Vehicle - 3wks | -4.58333 | 2.357023 | -1.94454 | 0.2095 | -0.183097 | -0.507150 | 0.0581578 |

Difference Plot

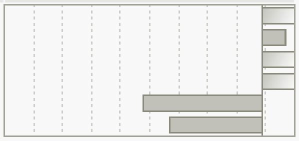

Excluded Rows 123

**S9L**

#### Oneway Analysis of correlation_baseline By condition

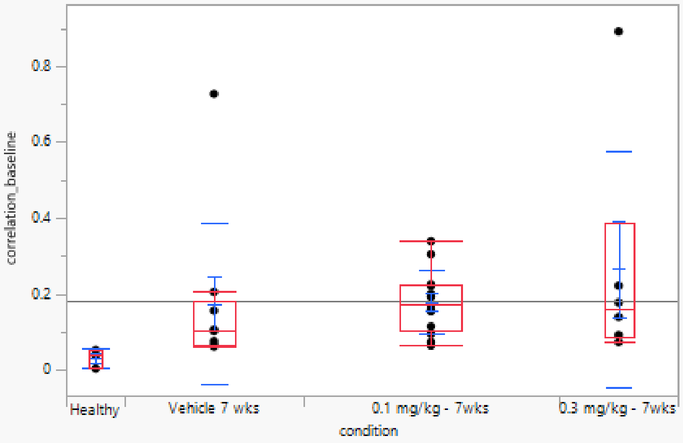

##### Quantiles

| Level | Minimum | 10% | 25% | Median | 75% | 90% | Maximum |
|---|---|---|---|---|---|---|---|
| Healthy | 0.004105 | 0.004105 | 0.004105 | 0.038322 | 0.053009 | 0.053009 | 0.053009 |
| Vehicle 7 wks | 0.062297 | 0.062297 | 0.067043 | 0.102838 | 0.181218 | 0.727134 | 0.727134 |
| 0.1 mg/kg - 7wks | 0.064952 | 0.068644 | 0.105566 | 0.171295 | 0.223917 | 0.325541 | 0.339189 |
| 0.3 mg/kg - 7wks | 0.073988 | 0.073988 | 0.088001 | 0.158984 | 0.389578 | 0.891375 | 0.891375 |

##### Means and Std Deviations

| Level | Number | Mean | Std Dev | Std Err Mean | Lower 95% | Upper 95% | Std Dev Lower 95% | Std Dev Upper 95% |
|---|---|---|---|---|---|---|---|---|
| Healthy | 3 | 0.0318117 | 0.0250936 | 0.0144878 | -0.030524 | 0.0941477 | 0.0130652 | 0.1577067 |
| Vehicle 7 wks | 9 | 0.1746904 | 0.2126264 | 0.0708755 | 0.0112512 | 0.3381295 | 0.14362 | 0.4073435 |
| 0.1 mg/kg - 7wks | 13 | 0.1785662 | 0.0827798 | 0.022959 | 0.1285428 | 0.2285895 | 0.0593603 | 0.1366476 |
| 0.3 mg/kg - 7wks | 6 | 0.266386 | 0.31098 | 0.1269571 | -0.059968 | 0.5927396 | 0.1941162 | 0.762714 |

##### Wilcoxon / Kruskal-Wallis Tests (Rank Sums)

| Level | Count | Score Sum | Expected Score | Score Mean | (Mean-Mean0)/Std0 |
|---|---|---|---|---|---|
| Healthy | 3 | 6.000 | 48.000 | 2.0000 | -2.773 |
| Vehicle 7 wks | 9 | 126.000 | 144.000 | 14.0000 | -0.762 |
| 0.1 mg/kg - 7wks | 13 | 251.000 | 208.000 | 19.3077 | 1.701 |
| 0.3 mg/kg - 7wks | 6 | 113.000 | 96.000 | 18.8333 | 0.825 |

##### Kruskal-Wallis Test, ChiSquare Approximation

| ChiSquare | DF | Prob>ChiSq |
|---|---|---|
| 9.8516 | 3 | 0.0199* |

##### Nonparametric Comparisons For All Pairs Using Steel-Dwass Method

| q* | Alpha |
|---|---|
| 2.56903 | 0.05 |

| Level | - Level | Score Mean Difference | Std Err Dif | Z | p-Value | Hodges-Lehmann | Lower CL | Upper CL |
|---|---|---|---|---|---|---|---|---|
| 0.1 mg/kg - 7wks | Healthy | 7.79487 | 3.049450 | 2.55616 | 0.0517 | 0.147579 | . | . |
| Vehicle 7 wks | Healthy | 5.77778 | 2.403701 | 2.40370 | 0.0764 | 0.064516 | . | . |
| 0.3 mg/kg - 7wks | Healthy | 4.25000 | 1.936492 | 2.19469 | 0.1247 | 0.130428 | . | . |
| 0.1 mg/kg - 7wks | Vehicle 7 wks | 3.94872 | 2.815808 | 1.40234 | 0.4978 | 0.052591 | -0.082550 | 0.1549223 |
| 0.3 mg/kg - 7wks | Vehicle 7 wks | 2.63889 | 2.357023 | 1.11959 | 0.6774 | 0.035088 | -0.549104 | 0.8148186 |
| 0.3 mg/kg - 7wks | 0.1 mg/kg - 7wks | -0.36538 | 2.777350 | -0.13156 | 0.9992 | -0.003362 | -0.147994 | 0.6907874 |

Excluded Rows 113

#### Figure S10

### S10A

**Oneway Analysis of p100-n70_amplitude_by_baseline By condition**

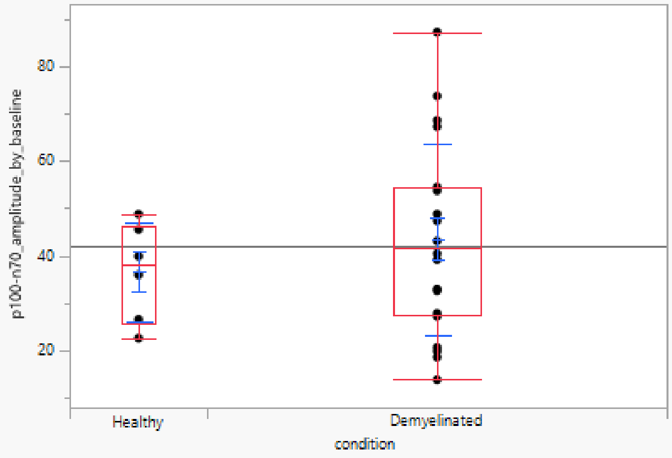

**Quantiles**

| Level | Minimum | 10% | 25% | Median | 75% | 90% | Maximum |
|---|---|---|---|---|---|---|---|
| Healthy | 22.53129 | 22.53129 | 25.50807 | 37.89695 | 46.35021 | 48.77696 | 48.77696 |
| Demyelinated | 13.74972 | 18.65283 | 27.30586 | 41.76553 | 54.31898 | 73.21806 | 87.25876 |

**Means and Std Deviations**

| Level | Number | Mean | Std Dev | Std Err Mean | Lower 95% | Upper 95% | Std Dev Lower 95% | Std Dev Upper 95% |
|---|---|---|---|---|---|---|---|---|
| Healthy | 6 | 36.523963 | 10.384556 | 4.239477 | 25.62604 | 47.421885 | 6.4821227 | 25.469306 |
| Demyelinated | 20 | 43.536898 | 20.218959 | 4.5210966 | 34.074134 | 52.999662 | 15.376324 | 29.531238 |

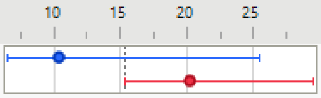

**Wilcoxon / Kruskal-Wallis Tests (Rank Sums)**

| Level | Count | Score Sum | Expected Score | Score Mean | (Mean-Mean0)/Std0 |
|---|---|---|---|---|---|
| Healthy | 6 | 69.000 | 81.000 | 11.5000 | -0.700 |
| Demyelinated | 20 | 282.000 | 270.000 | 14.1000 | 0.700 |

**Wilcoxon Two-Sample Test, Normal Approximation**

| S | Z | Prob>\|Z\| |
|---|---|---|
| 69 | -0.69987 | 0.4840 |

**Kruskal-Wallis Test, ChiSquare Approximation**

| ChiSquare | DF | Prob>ChiSq |
|---|---|---|
| 0.5333 | 1 | 0.4652 |

Excluded Rows 118

## S10B

##### Oneway Analysis of p100-n70_amplitude_by_baseline By condition

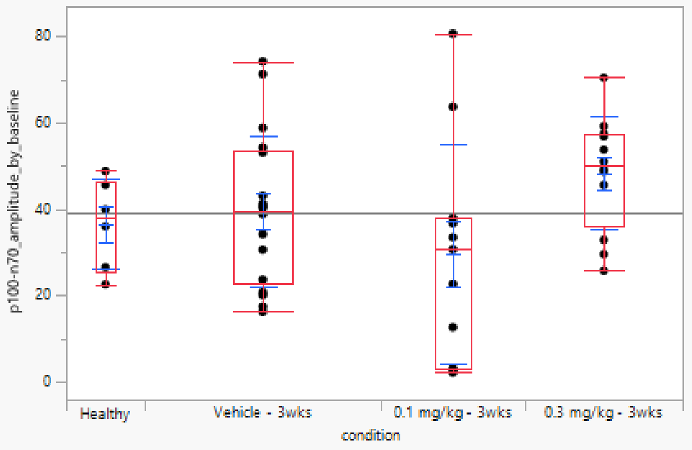

##### Quantiles

| Level | Minimum | 10% | 25% | Median | 75% | 90% | Maximum |
|---|---|---|---|---|---|---|---|
| Healthy | 22.53129 | 22.53129 | 25.50807 | 37.89695 | 46.35021 | 48.77696 | 48.77696 |
| Vehicle - 3wks | 16.21827 | 17.27619 | 22.87906 | 39.52324 | 53.38876 | 71.50978 | 74.15498 |
| 0.1 mg/kg - 3wks | 2.083847 | 2.188597 | 3.091341 | 30.65442 | 37.86811 | 77.22972 | 80.61881 |
| 0.3 mg/kg - 3wks | 25.73802 | 26.87651 | 36.01172 | 50.0602 | 57.26638 | 67.02968 | 70.40214 |

##### Means and Std Deviations

| Level | Number | Mean | Std Dev | Std Err Mean | Lower 95% | Upper 95% | Std Dev Lower 95% | Std Dev Upper 95% |
|---|---|---|---|---|---|---|---|---|
| Healthy | 6 | 36.523963 | 10.384556 | 4.239477 | 25.62604 | 47.421885 | 6.4821227 | 25.469306 |
| Vehicle - 3wks | 18 | 39.379602 | 17.492521 | 4.1230268 | 30.680776 | 48.078428 | 13.126171 | 26.223808 |
| 0.1 mg/kg - 3wks | 11 | 29.630651 | 25.346109 | 7.6421394 | 12.602904 | 46.658399 | 17.709758 | 44.480737 |
| 0.3 mg/kg - 3wks | 12 | 48.326637 | 13.166871 | 3.8009482 | 39.960807 | 56.692468 | 9.3273475 | 22.355741 |

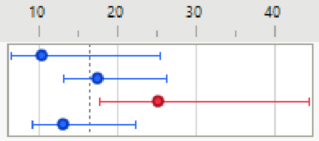

##### Wilcoxon / Kruskal-Wallis Tests (Rank Sums)

| Level | Count | Score Sum | Expected Score | Score Mean | (Mean-Mean0)/Std0 |
|---|---|---|---|---|---|
| Healthy | 6 | 132.000 | 144.000 | 22.0000 | -0.367 |
| Vehicle - 3wks | 18 | 428.000 | 432.000 | 23.7778 | -0.077 |
| 0.1 mg/kg - 3wks | 11 | 191.000 | 264.000 | 17.3636 | -1.822 |
| 0.3 mg/kg - 3wks | 12 | 377.000 | 288.000 | 31.4167 | 2.159 |

###### Kruskal-Wallis Test, ChiSquare Approximation

| ChiSquare | DF | Prob>ChiSq |
|---|---|---|
| 6.2204 | 3 | 0.1014 |

Excluded Rows 97

**S10C**

#### Oneway Analysis of p100-n70_amplitude_by_baseline By condition

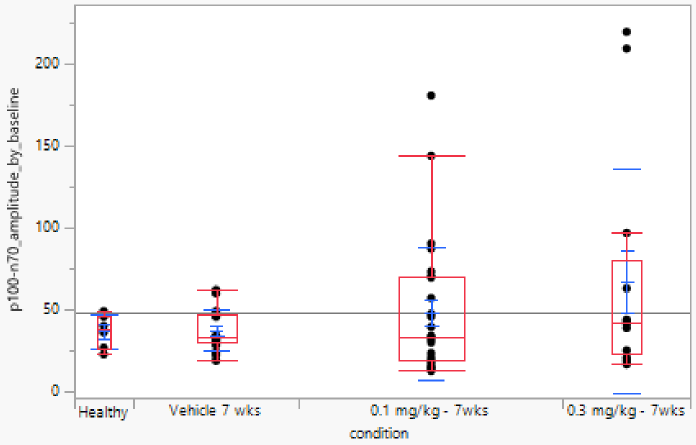

##### Quantiles

| Level | Minimum | 10% | 25% | Median | 75% | 90% | Maximum |
|---|---|---|---|---|---|---|---|
| Healthy | 22.53129 | 22.53129 | 25.50807 | 37.89695 | 46.35021 | 48.77696 | 48.77696 |
| Vehicle 7 wks | 18.69962 | 21.34592 | 29.42997 | 32.48892 | 46.82736 | 60.10557 | 61.89275 |
| 0.1 mg/kg - 7wks | 12.36293 | 13.9297 | 18.67283 | 33.07366 | 69.4975 | 100.9979 | 180.798 |
| 0.3 mg/kg - 7wks | 16.37258 | 17.21007 | 22.63155 | 41.40866 | 79.7495 | 215.6271 | 219.7059 |

##### Means and Std Deviations

| Level | Number | Mean | Std Dev | Std Err Mean | Lower 95% | Upper 95% | Std Dev Lower 95% | Std Dev Upper 95% |
|---|---|---|---|---|---|---|---|---|
| Healthy | 6 | 36.523963 | 10.384556 | 4.239477 | 25.62604 | 47.421885 | 6.4821227 | 25.469306 |
| Vehicle 7 wks | 17 | 36.948469 | 12.524484 | 3.0376336 | 30.508974 | 43.387965 | 9.3278579 | 19.061375 |
| 0.1 mg/kg - 7wks | 27 | 47.696774 | 40.437485 | 7.7821975 | 31.700238 | 63.69331 | 31.845183 | 55.416807 |
| 0.3 mg/kg - 7wks | 13 | 67.40275 | 68.715293 | 19.058193 | 25.878514 | 108.92699 | 49.274774 | 113.43075 |

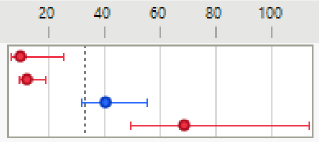

##### Wilcoxon / Kruskal-Wallis Tests (Rank Sums)

| Level | Count | Score Sum | Expected Score | Score Mean | (Mean-Mean0)/Std0 |
|---|---|---|---|---|---|
| Healthy | 6 | 192.000 | 192.000 | 32.0000 | 0.000 |
| Vehicle 7 wks | 17 | 531.000 | 544.000 | 31.2353 | -0.194 |
| 0.1 mg/kg - 7wks | 27 | 826.000 | 864.000 | 30.5926 | -0.521 |
| 0.3 mg/kg - 7wks | 13 | 467.000 | 416.000 | 35.9231 | 0.858 |

###### Kruskal-Wallis Test, ChiSquare Approximation

| ChiSquare | DF | Prob>ChiSq |
|---|---|---|
| 0.7842 | 3 | 0.8532 |

Excluded Rows 81

**S10D**

Oneway Analysis of p100-n70_amplitude_by_baseline By condition

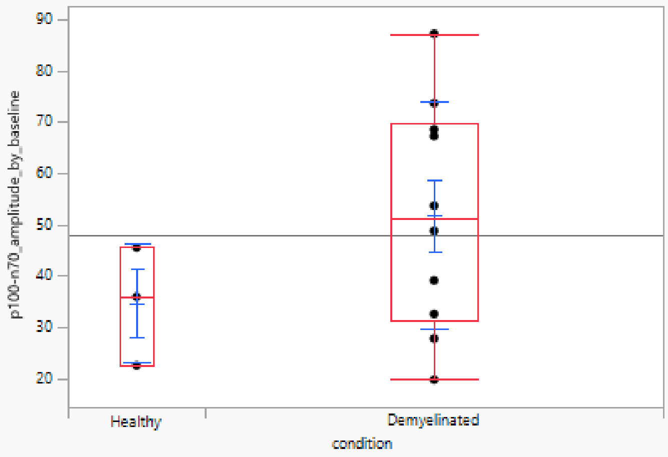

##### Quantiles

| Level | Minimum | 10% | 25% | Median | 75% | 90% | Maximum |
|---|---|---|---|---|---|---|---|
| Healthy | 22.53129 | 22.53129 | 22.53129 | 35.93375 | 45.5413 | 45.5413 | 45.5413 |
| Demyelinated | 19.78191 | 20.5836 | 31.37069 | 51.25411 | 69.86958 | 85.90621 | 87.25876 |

##### Means and Std Deviations

| Level | Number | Mean | Std Dev | Std Err Mean | Lower 95% | Upper 95% | Std Dev Lower 95% | Std Dev Upper 95% |
|---|---|---|---|---|---|---|---|---|
| Healthy | 3 | 34.668778 | 11.557045 | 6.6724631 | 5.9594867 | 63.378069 | 6.017271 | 72.632962 |
| Demyelinated | 10 | 51.863467 | 22.120377 | 6.9950774 | 36.039502 | 67.687431 | 15.215174 | 40.383185 |

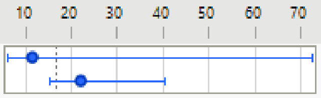

##### Wilcoxon / Kruskal-Wallis Tests (Rank Sums)

| Level | Count | Score Sum | Expected Score | Score Mean | (Mean-Mean0)/Std0 |
|---|---|---|---|---|---|
| Healthy | 3 | 14.000 | 21.000 | 4.66667 | -1.099 |
| Demyelinated | 10 | 77.000 | 70.000 | 7.70000 | 1.099 |

###### Wilcoxon Two-Sample Test, Normal Approximation

| S | Z | Prob>\|Z\| |
|---|---|---|
| 14 | -1.09870 | 0.2719 |

###### Kruskal-Wallis Test, ChiSquare Approximation

| ChiSquare | DF | Prob>ChiSq |
|---|---|---|
| 1.4000 | 1 | 0.2367 |

Excluded Rows 131

## S10E

##### Oneway Analysis of p100-n70_amplitude_by_baseline By condition

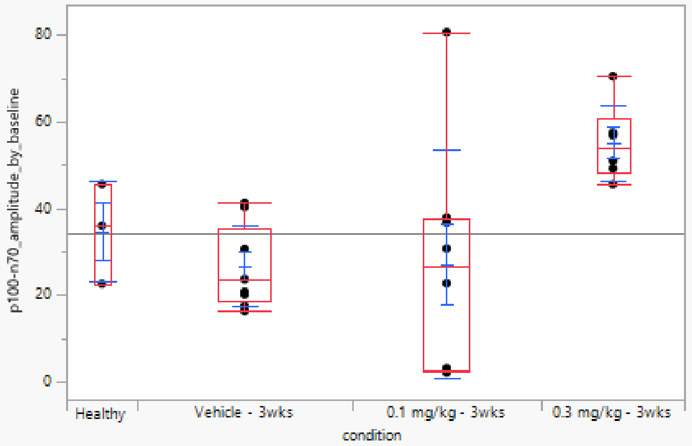

##### Quantiles

| Level | Minimum | 10% | 25% | Median | 75% | 90% | Maximum |
|---|---|---|---|---|---|---|---|
| Healthy | 22.53129 | 22.53129 | 22.53129 | 35.93375 | 45.5413 | 45.5413 | 45.5413 |
| Vehicle - 3wks | 16.21827 | 16.21827 | 18.71508 | 23.60928 | 35.4526 | 41.19528 | 41.19528 |
| 0.1 mg/kg - 3wks | 2.083847 | 2.083847 | 2.728534 | 26.64728 | 37.57482 | 80.61881 | 80.61881 |
| 0.3 mg/kg - 3wks | 45.53058 | 45.53058 | 48.26071 | 53.86758 | 60.67054 | 70.40214 | 70.40214 |

##### Means and Std Deviations

| Level | Number | Mean | Std Dev | Std Err Mean | Lower 95% | Upper 95% | Std Dev Lower 95% | Std Dev Upper 95% |
|---|---|---|---|---|---|---|---|---|
| Healthy | 3 | 34.668778 | 11.557045 | 6.6724631 | 5.9594867 | 63.378069 | 6.017271 | 72.632962 |
| Vehicle - 3wks | 9 | 26.733324 | 9.4273423 | 3.1424474 | 19.486827 | 33.97982 | 6.3677647 | 18.060628 |
| 0.1 mg/kg - 3wks | 8 | 27.032401 | 26.476048 | 9.3606965 | 4.897871 | 49.166931 | 17.505278 | 53.885962 |
| 0.3 mg/kg - 3wks | 6 | 55.044219 | 8.7902937 | 3.5886224 | 45.819372 | 64.269067 | 5.4869717 | 21.559197 |

##### Wilcoxon / Kruskal-Wallis Tests (Rank Sums)

| Level | Count | Score Sum | Expected Score | Score Mean | (Mean-Mean0)/Std0 |
|---|---|---|---|---|---|
| Healthy | 3 | 42.000 | 40.500 | 14.0000 | 0.080 |
| Vehicle - 3wks | 9 | 90.000 | 121.500 | 10.0000 | -1.671 |
| 0.1 mg/kg - 3wks | 8 | 85.000 | 108.000 | 10.6250 | -1.250 |
| 0.3 mg/kg - 3wks | 6 | 134.000 | 81.000 | 22.3333 | 3.195 |

###### Kruskal-Wallis Test, ChiSquare Approximation

| ChiSquare | DF | Prob>ChiSq |
|---|---|---|
| 11.0306 | 3 | 0.0116* |

##### Nonparametric Comparisons For All Pairs Using Steel-Dwass Method

| q* | Alpha |
|---|---|
| 2.56903 | 0.05 |

| Level | - Level | Score Mean Difference | Std Err Dif | Z | p-Value | Hodges-Lehmann | Lower CL | Upper CL |
|---|---|---|---|---|---|---|---|---|
| 0.3 mg/kg - 3wks | Vehicle - 3wks | 7.36111 | 2.357023 | 3.12305 | 0.0097* | 28.8083 | 8.9086 | 49.71374 |
| 0.3 mg/kg - 3wks | 0.1 mg/kg - 3wks | 5.10417 | 2.259240 | 2.25924 | 0.1077 | 30.4218 | -29.6692 | 67.31080 |
| 0.3 mg/kg - 3wks | Healthy | 3.75000 | 1.936492 | 1.93649 | 0.2127 | 21.1723 | . | . |
| 0.1 mg/kg - 3wks | Vehicle - 3wks | -0.35417 | 2.453739 | -0.14434 | 0.9989 | -2.8606 | -28.4695 | 40.35670 |
| 0.1 mg/kg - 3wks | Healthy | -1.14583 | 2.245366 | -0.51031 | 0.9567 | -11.0700 | . | . |
| Vehicle - 3wks | Healthy | -2.66667 | 2.403701 | -1.10940 | 0.6837 | -5.3804 | . | . |

Excluded Rows 118

**S10F**

#### Oneway Analysis of p100-n70_amplitude_by_baseline By condition

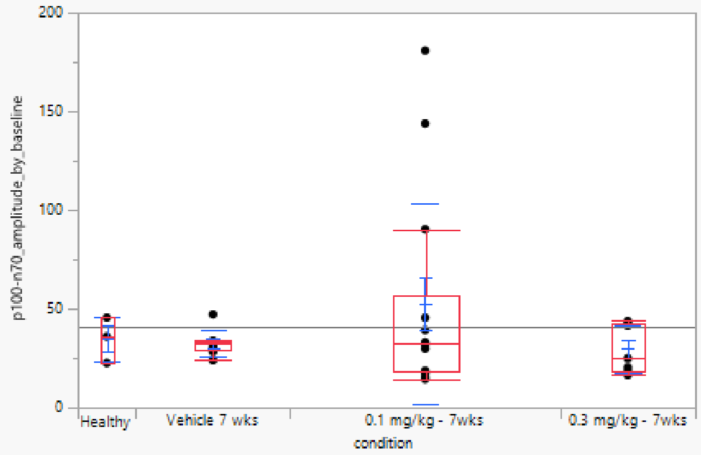

##### Quantiles

| Level | Minimum | 10% | 25% | Median | 75% | 90% | Maximum |
|---|---|---|---|---|---|---|---|
| Healthy | 22.53129 | 22.53129 | 22.53129 | 35.93375 | 45.5413 | 45.5413 | 45.5413 |
| Vehicle 7 wks | 23.88322 | 23.88322 | 28.82397 | 32.37542 | 33.73568 | 47.19582 | 47.19582 |
| 0.1 mg/kg - 7wks | 14.40551 | 14.87993 | 18.32449 | 32.49144 | 56.87208 | 162.3195 | 180.798 |
| 0.3 mg/kg - 7wks | 16.37258 | 16.37258 | 18.4663 | 24.93371 | 42.53937 | 43.77948 | 43.77948 |

##### Means and Std Deviations

| Level | Number | Mean | Std Dev | Std Err Mean | Lower 95% | Upper 95% | Std Dev Lower 95% | Std Dev Upper 95% |
|---|---|---|---|---|---|---|---|---|
| Healthy | 3 | 34.668778 | 11.557045 | 6.6724631 | 5.9594867 | 63.378069 | 6.017271 | 72.632962 |
| Vehicle 7 wks | 8 | 32.738772 | 6.6996346 | 2.3686785 | 27.137737 | 38.339806 | 4.4296252 | 13.635579 |
| 0.1 mg/kg - 7wks | 14 | 52.56519 | 50.792789 | 13.574944 | 23.238306 | 81.892073 | 36.822432 | 81.829338 |
| 0.3 mg/kg - 7wks | 7 | 29.689929 | 12.345816 | 4.6662798 | 18.271954 | 41.107904 | 7.9555624 | 27.186303 |

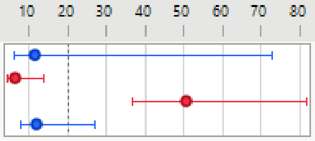

##### Wilcoxon / Kruskal-Wallis Tests (Rank Sums)

| Level | Count | Score Sum | Expected Score | Score Mean | (Mean-Mean0)/Std0 |
|---|---|---|---|---|---|
| Healthy | 3 | 56.000 | 49.500 | 18.6667 | 0.388 |
| Vehicle 7 wks | 8 | 135.000 | 132.000 | 16.8750 | 0.109 |
| 0.1 mg/kg - 7wks | 14 | 240.000 | 231.000 | 17.1429 | 0.323 |
| 0.3 mg/kg - 7wks | 7 | 97.000 | 115.500 | 13.8571 | -0.821 |

##### Kruskal-Wallis Test, ChiSquare Approximation

| ChiSquare | DF | Prob>ChiSq |
|---|---|---|
| 0.7942 | 3 | 0.8509 |

Excluded Rows 112

**S10G**

Oneway Analysis of p100-n70_amplitude_by_baseline By condition

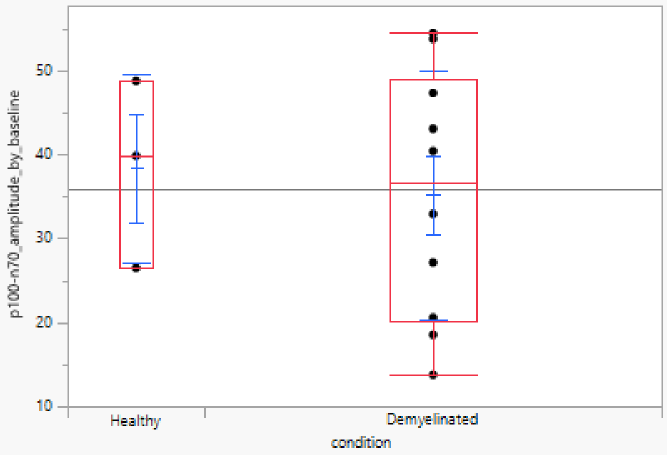

##### Quantiles

| Level | Minimum | 10% | 25% | Median | 75% | 90% | Maximum |
|---|---|---|---|---|---|---|---|
| Healthy | 26.50033 | 26.50033 | 26.50033 | 39.86015 | 48.77696 | 48.77696 | 48.77696 |
| Demyelinated | 13.74972 | 14.22748 | 20.06805 | 36.69164 | 48.95303 | 54.42424 | 54.49442 |

##### Means and Std Deviations

| Level | Number | Mean | Std Dev | Std Err Mean | Lower 95% | Upper 95% | Std Dev Lower 95% | Std Dev Upper 95% |
|---|---|---|---|---|---|---|---|---|
| Healthy | 3 | 38.379147 | 11.211916 | 6.4732029 | 10.527203 | 66.231091 | 5.8375769 | 70.463919 |
| Demyelinated | 10 | 35.210329 | 14.820725 | 4.6867247 | 24.608222 | 45.812437 | 10.194216 | 27.056866 |

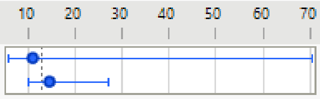

##### Wilcoxon / Kruskal-Wallis Tests (Rank Sums)

| Level | Count | Score Sum | Expected Score | Score Mean | (Mean-Mean0)/Std0 |
|---|---|---|---|---|---|
| Healthy | 3 | 22.000 | 21.000 | 7.33333 | 0.085 |
| Demyelinated | 10 | 69.000 | 70.000 | 6.90000 | -0.085 |

###### Wilcoxon Two-Sample Test, Normal Approximation

| S | Z | Prob>\|Z\| |
|---|---|---|
| 22 | 0.08452 | 0.9326 |

###### Kruskal-Wallis Test, ChiSquare Approximation

| ChiSquare | DF | Prob>ChiSq |
|---|---|---|
| 0.0286 | 1 | 0.8658 |

Excluded Rows 131

## S10H

##### Oneway Analysis of p100-n70_amplitude_by_baseline By condition

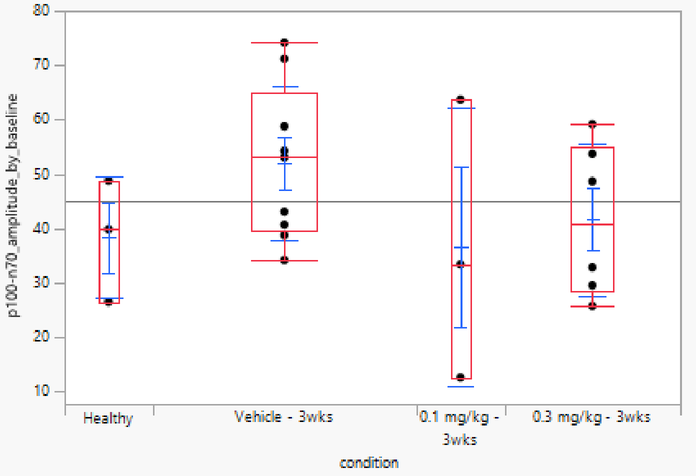

##### Quantiles

| Level | Minimum | 10% | 25% | Median | 75% | 90% | Maximum |
|---|---|---|---|---|---|---|---|
| Healthy | 26.50033 | 26.50033 | 26.50033 | 39.86015 | 48.77696 | 48.77696 | 48.77696 |
| Vehicle - 3wks | 34.1748 | 34.1748 | 39.7345 | 53.09648 | 64.9922 | 74.15498 | 74.15498 |
| 0.1 mg/kg - 3wks | 12.59552 | 12.59552 | 12.59552 | 33.40909 | 63.67334 | 63.67334 | 63.67334 |
| 0.3 mg/kg - 3wks | 25.73802 | 25.73802 | 28.58424 | 40.74979 | 55.0825 | 59.1606 | 59.1606 |

##### Means and Std Deviations

| Level | Number | Mean | Std Dev | Std Err Mean | Lower 95% | Upper 95% | Std Dev Lower 95% | Std Dev Upper 95% |
|---|---|---|---|---|---|---|---|---|
| Healthy | 3 | 38.379147 | 11.211916 | 6.4732029 | 10.527203 | 66.231091 | 5.8375769 | 70.463919 |
| Vehicle - 3wks | 9 | 52.02588 | 14.195476 | 4.7318255 | 41.114271 | 62.93749 | 9.5884344 | 27.195281 |
| 0.1 mg/kg - 3wks | 3 | 36.559318 | 25.684217 | 14.82879 | -27.24381 | 100.36245 | 13.3727 | 161.41849 |
| 0.3 mg/kg - 3wks | 6 | 41.609056 | 13.9941 | 5.7130674 | 26.923148 | 56.294963 | 8.7352292 | 34.322125 |

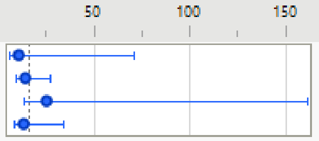

##### Wilcoxon / Kruskal-Wallis Tests (Rank Sums)

| Level | Count | Score Sum | Expected Score | Score Mean | (Mean-Mean0)/Std0 |
|---|---|---|---|---|---|
| Healthy | 3 | 25.000 | 33.000 | 8.3333 | -0.754 |
| Vehicle - 3wks | 9 | 124.000 | 99.000 | 13.7778 | 1.741 |
| 0.1 mg/kg - 3wks | 3 | 26.000 | 33.000 | 8.6667 | -0.653 |
| 0.3 mg/kg - 3wks | 6 | 56.000 | 66.000 | 9.3333 | -0.740 |

###### Kruskal-Wallis Test, ChiSquare Approximation

| ChiSquare | DF | Prob>ChiSq |
|---|---|---|
| 3.2150 | 3 | 0.3596 |

Small sample sizes. Refer to statistical tables for tests, rather than large-sample approximations.

Excluded Rows 123

## S10I

##### Oneway Analysis of p100-n70_amplitude_by_baseline By condition

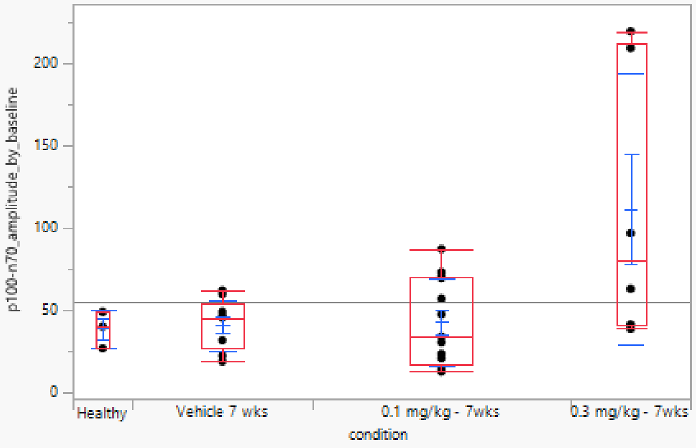

##### Quantiles

| Level | Minimum | 10% | 25% | Median | 75% | 90% | Maximum |
|---|---|---|---|---|---|---|---|
| Healthy | 26.50033 | 26.50033 | 26.50033 | 39.86015 | 48.77696 | 48.77696 | 48.77696 |
| Vehicle 7 wks | 18.69962 | 18.69962 | 26.79486 | 45.38125 | 54.25259 | 61.89275 | 61.89275 |
| 0.1 mg/kg - 7wks | 12.36293 | 12.9229 | 17.24421 | 33.75439 | 69.90352 | 81.49329 | 87.12507 |
| 0.3 mg/kg - 7wks | 38.52495 | 38.52495 | 40.50692 | 79.7495 | 212.0581 | 219.7059 | 219.7059 |

##### Means and Std Deviations

| Level | Number | Mean | Std Dev | Std Err Mean | Lower 95% | Upper 95% | Std Dev Lower 95% | Std Dev Upper 95% |
|---|---|---|---|---|---|---|---|---|
| Healthy | 3 | 38.379147 | 11.211916 | 6.4732029 | 10.527203 | 66.231091 | 5.8375769 | 70.463919 |
| Vehicle 7 wks | 9 | 40.690423 | 15.52343 | 5.1744768 | 28.758058 | 52.622787 | 10.48541 | 29.739336 |
| 0.1 mg/kg - 7wks | 13 | 42.453864 | 26.278923 | 7.2884618 | 26.57367 | 58.334058 | 18.844247 | 43.379541 |
| 0.3 mg/kg - 7wks | 6 | 111.40104 | 82.675374 | 33.75208 | 24.638557 | 198.16353 | 51.60663 | 202.77078 |

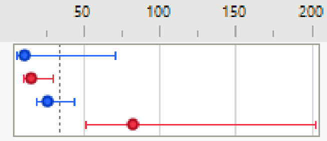

##### Wilcoxon / Kruskal-Wallis Tests (Rank Sums)

| Level | Count | Score Sum | Expected Score | Score Mean | (Mean-Mean0)/Std0 |
|---|---|---|---|---|---|
| Healthy | 3 | 41.000 | 48.000 | 13.6667 | -0.434 |
| Vehicle 7 wks | 9 | 129.000 | 144.000 | 14.3333 | -0.631 |
| 0.1 mg/kg - 7wks | 13 | 184.000 | 208.000 | 14.1538 | -0.941 |
| 0.3 mg/kg - 7wks | 6 | 142.000 | 96.000 | 23.6667 | 2.275 |

##### Kruskal-Wallis Test, ChiSquare Approximation

| ChiSquare | DF | Prob>ChiSq |
|---|---|---|
| 5.3021 | 3 | 0.1510 |

Excluded Rows 113
